# Historic breeding practices contribute to germplasm divergence in leaf specialized metabolism and ecophysiology in cultivated sunflower (*Helianthus annuus*)

**DOI:** 10.1101/2024.02.09.579651

**Authors:** Jordan A. Dowell, Alan W. Bowsher, Amna Jamshad, Rahul Shah, John M. Burke, Lisa A. Donovan, Chase M. Mason

## Abstract

The use of hybrid breeding systems to increase crop yields has been the cornerstone of modern agriculture and is exemplified in the breeding and improvement of cultivated sunflower (*Helianthus annuus)*. However, it is poorly understood what effect supporting separate breeding pools in such systems, combined with continued selection for yield, may have on leaf ecophysiology and specialized metabolite variation. Here, we analyze 288 cultivated *H. annuus* lines to examine the genomic basis of several specialized metabolites and agronomically important traits across major heterotic groups. Heterotic group identity supports phenotypic divergences between fertility restoring and cytoplasmic male-sterility maintainer lines in leaf ecophysiology and specialized metabolism. However, the divergence is not associated with physical linkage to nuclear genes that support current hybrid breeding systems in cultivated *H. annuus*. Further, we identified four genomic regions associated with variation in leaf ecophysiology and specialized metabolism that co-localize with previously identified QTLs in cultivated *H. annuus* for quantitative self-compatibility traits and with SPH-proteins, a recently discovered family of proteins associated with self-incompatibility and self/nonself recognition in *Papaver rhoeas* (common poppy) with suggested conserved downstream mechanisms among eudicots. Self-compatibility is a derived trait in cultivated *H. annuus* with quantitative variation in selfing success, suggesting that trait linkage to divergent phenotypic traits may have partially arisen as a potential unintended consequence of historical breeding practices. Further work is necessary to confirm the self-incompatibility mechanisms in cultivated *H. annuus* and their relationship to the integrative and polygenic architecture of leaf ecophysiology and specialized metabolism in cultivated sunflower.

## Introduction

The mid-twentieth century’s green revolution transformed agriculture worldwide, leading to significant increases in yield for many crops (Evenson, 2003). Adopting hybrid breeding practices was one of many contributing factors to the success of the Green Revolution for many crops (Flint-Garcia, 2013). Hybrid breeding takes advantage of heterosis, more colloquially termed ‘hybrid vigor,’ where the combination of two disparate genotypes often produces larger, higher-yielding, and more disease-resistant offspring (Bohra et al., 2016; Meena et al., 2017). Since the green revolution, our understanding of the genetic basis of traits and how traits contribute to yield has improved significantly (Slafer, 2003). Alongside improvements in genotype-to-phenotype predictions, breeding targets have shifted away from total yield towards yield regularity to combat year-to-year biotic and abiotic stress variability (Davey and Jan, 2010; Vincourt, 2014; Vear, 2016; Radanović et al., 2018). While the adoption of heterotic breeding practices and shifts among breeding targets aims to feed a growing world under threat from the effects of climate change, few studies have explicitly examined how post-green-revolution breeding practices have shaped the current phenotypic diversity within crop germplasm pools (Condón et al., 2009; Flint-Garcia, 2013; Fu, 2015; Gage et al., 2017; Mascher et al., 2019).

### Hybrid breeding

Hybrid breeding programs are simplified by securing genetic sources of male sterility and complementary sources of fertility-restoring alleles or genes (Bohra et al., 2016). Cytoplasmic male sterility (CMS) is leveraged as the source of male sterility in crops such as sunflower, maize, and rice, where the presence or absence of particular mitochondrial or nuclear alleles either disable or restore mitochondrial processes during pollen formation, allowing for controlled inhibition of successful pollen production (Onemli and Gucer, 2010; Meena et al., 2017). Three types of parental lines are needed for hybrid breeding programs in cultivated sunflower: CMS lines, HA (*Helianthus annuus*)-maintainer lines, and RHA (restorer *Helianthus annuus*)-fertility-restoring (RHA-restorer) lines (Bohra et al., 2016). CMS lines and HA-maintainers are typically isogenic, homozygous recessive for nuclear-encoded fertility restoring alleles, and differ in mitochondrial cytotype; HA-maintainers produce pollen through their lack of CMS-conferring mitochondrial alleles. RHA-restorer lines are homozygous dominant for fertility restoring alleles, and the F_1_ hybrid seed used by most growers is produced by crossing CMS lines with RHA-restorer lines. In contrast, the generation and maintenance of CMS lines require breeding HA-maintainers with CMS lines (Bohra et al., 2016). Experimental crosses leveraging male-sterile lines prevent self-fertilization due to the lack of pollen formation in the pollen-receptive line, ensuring successful outcrosses (Bohra et al., 2016; Radanović et al., 2018). Because of the ease of use, there is an observed bias toward using male-sterile and maintainer lines in breeding programs sunflowercompared to fertility restoring lines among many crops (Bohra et al., 2016; Radanović et al., 2018).

In cultivated sunflower, CMS was initially discovered by P. Leclercq in France, followed by the discovery of an associated fertility restoration source in the USA by M. L. Kinman (Vear, 2016; Radanović et al., 2018). Currently, there are over 70 reported sources of CMS and 30 sources of complementary nuclear fertility restoration alleles in cultivated sunflower (Davey and Jan, 2010; Onemli and Gucer, 2010). However, most CMS sources in cultivated sunflower are unstable and thus not viable for use in heterotic breeding, especially at an industrial scale (Davey and Jan, 2010; Onemli and Gucer, 2010). Therefore, the adoption of CMS in cultivated sunflower breeding programs comes from a few genetic sources. Thus, there is potential that the transition to CMS-based breeding contributed to observed genetic bottlenecks in cultivated sunflower, further reducing the genetic diversity of cultivated sunflower lines associated with modern breeding and improvement (Burke et al., 2002; Liu and Burke, 2006; Wills and Burke, 2007; Davey and Jan, 2010; Onemli and Gucer, 2010; Mandel et al., 2013).

### Self-incompatibility in sunflower

As with many fruit and seed crops, self-compatibility is a vital contributor to total yield as it helps assure seed production even without pollinator activity (Anita-Sari et al., 2017; Muñoz-Sanz et al., 2020). Like many other crop systems, early breeding targets for cultivated sunflower were aimed at increasing total yield (Vear, 2016). Interestingly, the wild progenitor of cultivated sunflower is self-incompatible, while modern cultivated lines are largely self-compatible(Gandhi et al., 2005; Wills and Burke, 2007; Baack et al., 2008; Mandel et al., 2013). Self-compatibility is also observed in many landraces of sunflower, and genomic information suggests that the likely sole origin of self-compatibility occurred during the early domestication of sunflower by indigenous North Americans (Radanović et al., 2018).

While the explicit molecular mechanisms of *Helianthus* self-incompatibility are unknown, early studies identified several QTLs for traits such as the quantity and quality of selfed seeds (Burke et al., 2002; Gandhi et al., 2005; Wills and Burke, 2007). All QTLs linked to self-compatibility traits in sunflower are hypothesized to be incompletely dominant (Burke et al., 2002; Gandhi et al., 2005; Wills and Burke, 2007), suggesting that there are combinatorial elements to self-compatibility that result in the variable success of self-pollination. Recent work on wild sunflower incompatibility suggests that one previously identified self-incompatibility QTL colocalizes with a large S-locus gene candidate on chromosome 17 of the XRQ sunflower genome assembly between 6.4 cM and 2.1 cM (Gandhi et al., 2005; Badouin et al., 2021). Validations of this S-locus gene candidate display a binary phenotype of self-incompatibility and expression in the pistil, suggesting that other mechanisms are at play and responsible for the combinatory nature of self-compatibility present in cultivated sunflower (Badouin et al., 2021). Further, the gene (Ha7650b) implicated is a serine/threonine kinase receptor-like protein with leucine-rich repeats (LRRs) and a malectin extracellular domain, and no ligand has been identified or suggested (Badouin et al., 2021). In exploring potential complementary elements of self-incompatibility, approximately 20 Mb upstream from the suggested S-locus, there are seven annotated genes (putative self-incompatibility S1) belonging to the SPH (S-protein homolog)-protein family (Hübner et al., 2019).

The SPH-protein family is a family of small disulfide-bonded secreted proteins present in eudicots and bryophytes, but noticeably absent in monocots (Ride et al., 1999; Rajasekar et al., 2019). SPH proteins share a highly temperature stable β-sandwich structure with 8–9 β-sheets that exhibit sequence variation in the loops connecting each sheet (Rajasekar et al., 2019). While research on the explicit functionality of these proteins across species is missing, the most charismatic examples of the SPH-protein family are found in poppy (*Papavera rhoas*). In poppy, SPH-proteins act as ligands and receptors (Wheeler et al., 2009, 2010; Rajasekar et al., 2019). However, proteins in this family are predicted to interact with a wide variety of receptor types (Wheeler et al., 2010; Rajasekar et al., 2019). In poppy, when two members of the SPH protein family (PrsS1 and PrpS1) are cognate, they confer sporophytic self-incompatibility (Foote et al., 1994; Wheeler et al., 2009; de Graaf et al., 2012). Combinations of allelic variants in SPH-proteins confer varying degrees of, or quantitative, self-incompatibility reminiscent of early QTLs for self-compatibility traits identified in sunflower (Burke et al., 2002; Gandhi et al., 2005; Wills and Burke, 2007) as opposed to binary phenotypes observed in the recently proposed sunflower S-locus gene (Badouin et al., 2021). Transgenic studies of poppy-specific SPH-proteins demonstrate combinations of cognate proteins trigger self-recognition and associated programmed cell death in reproductive and vegetative tissue of *Arabidopsis thaliana*, pointing toward the potential for conserved downstream cell programming (de Graaf et al., 2012; Lin et al., 2020). Further mutagenic studies of *Arabidopsis thaliana* and poppy SPH proteins support the robustness of the functionality of this protein family to sequence variation, and that sequence variation can confer degrees of self-incompatibility (de Graaf et al., 2012; Rajasekar et al., 2019; Lin et al., 2020). In-silico work on the SPH family suggests these proteins can act as a modifiable scaffold to support several peptides in the loops between β-sheets, with the potential for each peptide to interact with different receptors within and outside of this protein family, indicating broad potential for cell signaling outside of self-incompatibility (Rajasekar et al., 2019).

In the HA412HO sunflower genome assembly version 2.0, there are 51 annotated SPH-protein homologs, many of which align with previously identified QTL for self-incompatibility traits (Burke et al., 2002; Hübner et al., 2019). In comparison, the earlier *H. annuus* XRQ 2.0 sunflower genome based on the HanXRQr2.0-SUNRISE assembly has 53 SPH-protein homologs (Badouin et al., 2017). Among annotated homologs, only one SPH protein has documented expression in relevant expression in pollen and stamens of cultivated sunflower (locus tag: HanXRQr2_Chr17g0790791) (Badouin et al., 2017). This locus is within 1 Mb of a cluster of 7 SPH-proteins and 20Mb from the suggested binary sunflower S-gene candidate expressed in the pistil (Hübner et al., 2019; Badouin et al., 2021). In addition, it is important to note that, to our knowledge, no study has established quantitative expression needs for the induction of programmed cell death due to SPH proteins (de Graaf et al., 2012; Lin et al., 2020), indicating that expression values observed from RNAseq data are not necessarily indicative of functionality (Oshlack and Wakefield, 2009; Mandelboum et al., 2019).

It is fully expected that self-incompatibility genes would not be expressed in cultivated sunflower, as high self-compatibility was a key part of the domestication syndrome alongside reduced branching and seed shattering (Burke et al., 2002, 2005; Gandhi et al., 2005; Wills and Burke, 2007; Baack et al., 2008). However, the degree of selfing is known to be a continuous trait (Burke et al., 2002; Gandhi et al., 2005; Wills and Burke, 2007), such that selection for higher total yield has likely involved selection for improved seed set under self-pollination conditions. We, therefore, hypothesize that in the context of modern sunflower breeding, selection for increased yield may have acted in part on genes related to self-incompatibility or self/nonself-recognition systems and that this may have had unintended consequences on unrelated genes and plant phenotypes (Burke et al., 2005; Huang et al., 2023). We further predict that this pattern may manifest as allelic variants associated with plant functional traits colocalizing with genes with potential involvement with self-incompatibility and self/nonself-recognition (Burke et al., 2005; Hartfield and Bataillon, 2018).

### Breeding target history

As with most crops, the economics of production drive sunflower breeding targets, and the development of a more robust understanding of the mechanisms contributing to abiotic stress resistance has led to a breeding target shift towards the inclusion of yield regularity (Fu, 2015). In the 1990s, there was a large overproduction of sunflower seed, leading to a shift in breeding targets from total yield toward yield regularity to maintain yields at market needs while reducing resource inputs (Vear, 2016). For instance, French yields have changed little since the 1970s (2.5 tonnes/hectare to current 2.1 tonnes/hectare); however, agricultural practices for sunflower production have transitioned to little or no irrigation (Vear, 2016). The incorporation of drought resistance into breeding programs may have led to the homogenization of ecophysiological traits across heterotic groups. For example, the maintenance of heterotic groups of individuals with drought-resistant yield stability for use in breeding would necessitate the presence of drought-resistance alleles within and among groups if drought-resistance loci do not exhibit dominance when in the heterozygous F1 state. Given worldwide sunflower breeding has focused primarily on improving abiotic stress resistance (Vear, 2016) and a general bias toward the use of HA-maintainer lines in breeding programs (Bohra et al., 2016; Radanović et al., 2018), we hypothesize HA-maintainer lines will have phenotypic characteristics, aligned with current breeding trends, which contribute to yield regularity, in comparison to RHA-restorer lines which will be more similar to previous breeding trends of total yield.

Improving biotic stress resistance in cultivated sunflower has mainly been a North American goal, given that sunflowers and their ancestral pests and pathogens are native to North America (Vear, 2016). For example, two alleles maintained the resistance to downy mildew in Europe until the early 2000s; however, this resistance was overcome in the US as early as the 1980s (Vear, 2016). Breeding efforts for biotic stress resistance have focused on a few alleles that confer qualitative defenses (R-genes, hypersensitive-like responses, etc.) rather than quantitative defenses usually conferred by specialized metabolites (e.g., phenylpropanoids, terpenes, etc.) (Mestries et al., 1998; Micic et al., 2004, 2005; Gong et al., 2013; Vear, 2016; Seiler et al., 2017). Given the recent breeding focus on biotic stress resistance coupled with the bias in using male-sterile and maintainer lines in new breeding line production, we hypothesize greater specialized metabolite variation, underlying biotic stress resistance, among HA-maintainer lines in comparison to RHA-restorer, fertility restoring lines.

To examine the potential effects of post-green revolution breeding practices on crop sunflower, we here focus on a mapping panel of 288 cultivated sunflower lines as a model system to (1) examine phenotypic variation among heterotic breeding groups (HA-maintainer and RHA-restorer) in leaf ecophysiology and specialized metabolites, and (2) assess the genetic architecture of leaf ecophysiology and specialized metabolites.

## Materials and Methods

### Association Mapping Population

We used the Sunflower Association Mapping (SAM) panel described by Mandel et al. (2011, 2013). The SAM panel consists of 288 inbred cultivated sunflower lines selected based on microsatellites to capture approximately 90% of the allelic diversity present within the United States Department of Agriculture (USDA) and the French Institut National de la Recherche Agronomique (INRA) germplasm collections (Mandel et al., 2011). The SAM panel includes representatives of both heterotic groups (HA-maintainer and RHA-restorer) and market types (oil vs. non-oil [i.e., confectionery]), as well as an assortment of other genotypes outside these main breeding pools. The average introduction date for lines in this panel is 1990 (±7 years SD), although a few landraces and open-pollinated lines (OPV) with much older introduction dates are included. The SAM panel allows for robust mapping of phenotypes to the sunflower genome, with broad relevance across cultivated sunflower germplasm (Mandel et al., 2011, 2013; Masalia et al., 2018; Dowell et al., 2019; Temme et al., 2020). The SAM panel has the reference genome line, and the other 287 genotypes were previously subjected to whole-genome shotgun resequencing (Hübner et al., 2019). From the resequencing data, Todesco et al. (2020) developed a set of 1.4 million single nucleotide polymorphisms (SNPs) across all 17 *H. annuus* chromosomes. These SNPs served as the basis for our analyses, and the dataset was filtered based on the 288 genotypes used in this study to remove SNPs with a minor allele frequency of less than 5% or heterozygosity over 10%.

### Plant Cultivation

For each of the lines, growth conditions and parameters are described in detail by Dowell et al. (2019). We grew 3-4 replicate plants of each genotype in climate-controlled growth chambers at the University of Georgia Plant Biology Greenhouses in Athens, GA, United States. The full panel was randomly divided into eight groups grown in rapid succession from early June through late December 2014. Individuals within each group were arranged in a randomized spatial design in a plant growth chamber (Conviron, Winnipeg, Canada) to exclude pests and pathogens. Thus, plants represent constitutive variation in leaf ecophysiology and specialized metabolism. The chamber temperature was maintained at 25°C, and metal halide lighting was provided with a 14-hour day length. Pots were 1.3 L in volume, with a well-draining substrate (clean river sand), and received water and nutrient availability conducive to rapid growth. Pots were watered daily to field capacity, supplemented with 5ml of Osmocote 15-9-12 slow-release pelleted fertilizer (Scotts, Marysville, OH), as well as fertigated weekly to field capacity with a dilute liquid fertilizer (Jack’s 20-10-20, 5ml in 3 L, JR Peters Inc., Allentown, PA). Once plants were established with 4-6 fully expanded leaf pairs, each individual replicate plant was phenotyped for the present study before moving to a greenhouse bay for further growth and then reproductive phenotyping (Schneiter and Miller, 1981; Dowell et al., 2019).

### Ecophysiological Phenotyping

For phenotyping, we used a modified set of protocols previously applied to wild *Helianthus* (Mason and Donovan, 2015; Mason et al., 2016). For each replicate of every genotype, we identified the pair of most recently fully expanded leaves. For this leaf pair, one of the two leaves was used to assess ecophysiological traits: leaf fresh mass, dry mass, water content, leaf area, perimeter, aspect ratio, circularity, solidity, leaf mass per area (LMA), lamina thickness, lamina toughness, lamina density, and chlorophyll content on an area and mass basis.

From the selected ecophysiology leaf, we first took chlorophyll content measurements in SPAD units with a Konica Minolta SPAD-502-Plus (Konica Minolta Inc., Tokyo, Japan) and then clipped the leaf at the base of the petiole and weighed it for fresh mass. We scanned the leaf with a digital scanner to later derive leaf area, perimeter, aspect ratio, circularity, and solidity via ImageJ (Schneider et al., 2012). Following scanning, we measured lamina thickness with digital calipers at the midpoint of the leaf’s length, avoiding major veins. Lamina toughness was assessed using a digital penetrometer (FG-3000, Shimpo Inc.), averaging three measurements of force required to penetrate the leaf lamina with a millimeter-wide flat-tipped needle. Leaves were then dried at 60°C in a forced-air oven for 96 hours and weighed for dry mass. SPAD scores were converted into an area-basis (μmol m^-2^) using the approach of Markwell et al. (1995). Dry mass was used to calculate other traits, including water content, LMA, and mass-based chlorophyll content. In an unfortunate turn of events, dried leaves were accidentally destroyed for a subset of lines (N=125) while in storage awaiting processing, resulting in missing data for these lines for dry mass and the three traits derived using it.

### Coarse Defense Phenotyping and Sampling for Analytical Chemistry

We took the opposing leaf to phenotype coarse defense traits and sample tissue for analytical chemistry analysis of secondary metabolism. We split this leaf down the midrib, and one half was rolled, placed into a microcentrifuge tube, and snap-frozen in liquid nitrogen. This leaf sample was stored at −80°C for later analysis. The other half of the leaf was dried at 60°C for at least 36 h. This dried leaf material was used to assess four coarse leaf defense metrics: trichome density, ash content, total phenolics, and total flavonoids. Trichome density was calculated using a dissecting scope to count the trichomes present in a 0.25 cm^2^ region (Mason et al., 2016). The dried leaf tissue was then ground into a fine, homogenous powder. This material was used to estimate total flavonoids via the aluminum complexation assay and estimate total phenolics via the Folin-Ciocalteu assay (Singleton et al., 1999; Pękal and Pyrzynska, 2014; Webber and Mason, 2016). To estimate total ash content, leaf powder was combusted in crucibles in a muffle furnace at 600°C for 12 hours, and the pre-and post-combustion mass was compared (Moles et al., 2013).

### Analysis of Leaf Secondary Metabolism

To characterize broad metabolomic variation across cultivars, we used an untargeted approach. For each genotype, replicate frozen leaf samples were pooled in even proportion by mass, and these samples were ground to homogeneity under liquid nitrogen. A total of 10 mg of homogenized frozen material was extracted in 0.4 ml of 1:1 methanol:chloroform (v/v. in a sonicated ice-water bath for 30 min. Post-extraction, 0.2 ml of HPLC-grade water was added to each sample, and the mixture was vortexed and centrifuged. The aqueous layer was collected and further vortexed and centrifuged. The resulting aqueous layer extract was used for analysis by high-performance liquid chromatography (HPLC).

Two complementary approaches were used to: (1) identify secondary metabolites in sunflower, and (2) screen the large number of genotypes under study, using HPLC-MS and HPLC-UV, respectively. The first approach leveraged 12 lines within the SAM panel that represent approximately 50% of the allelic diversity present within the panel (Mandel et al., 2013). For genotypes within this ‘Core 12’, 3μl of each sample was injected into an Agilent 1200 HPLC with a 6220 accurate time-of-flight mass spectrometer with dual electrospray ionization and an ultraviolet diode array detector (Agilent Technologies, Santa Clara, CA, USA). This configuration coupled the HPLC to the mass spectrometer. Separation was achieved using an Agilent ZORBAX Rapid Resolution Eclipse XDB-C18 column (4.6 ⋅ 50 mm 1.8 μm), with mobile phase solvents of water:acetonitrile:formic acid = 97:3:0.1 (A) and 3:97:0.1 (B). The flow rate was set to 1 ml min^-1^, and the elution gradient was as follows: 3% B from 0-1 min, linear gradient to 17% B over 2 min, isocratic at 17% B for 2 min, linear gradient to 60% B over 4 min, then to 98% B over 2 min. Mass spectrometer acquisition was in negative ionization mode in m/z range of 100 to 1500 with the following settings: gas temperature, 350°C; drying gas flow, 13 l min-1; nebulizer pressure, 60 psig; capillary voltage, 3500 V; and fragmentor voltage, 125 V. For each extract analyzed via HPLC-MS, we analyzed the mass spectra of each retention time peak for matches to the NIST reference library, to create a putative list of metabolites within the ‘Core 12’ (Table S1) for comparison of retention-times to HPLC-UV data obtained for all samples, described below (Lemmon et al., 2017).

In the second approach, 3 µl of extract from each sample (including the ‘Core 12’ lines) was injected into the same Agilent 1200 HPLC making use of the same parameters, although coupled to an ultraviolet diode array detector set at 260, 270, 280, 310, and 350 nm, as opposed to a mass spectrometer. We chose only the 350nm wavelength for analyses based on the high number of peaks detected via the diode array detector at this wavelength versus the others. We linearly calibrated retention time and peak area based on the averages of retention time (RT) and peak areas for two common peaks (RT 4.11 and 7.05) found among all samples. We further used manual binning of peaks based on retention time similarity, producing a list of metabolites present among at least 200 of 288 lines. For each peak detected via HPLC-UV, if a corresponding peak was identified via HPLC-MS in the ‘Core 12’, we considered this a putative identification as the same compound. Hereafter, we use the following naming convention listing retention time (RT) from HPLC-UV data with the putative class identification from HPLC-MS (e.g., RT XX-SQTL or RT XX-UNK).

### Principal Components Analysis of Ecophysiology and Secondary Metabolism

To explore the presence of trait syndromes or chemotypes, we employed singular value decomposition to derive principal components (PC) based on groups of traits. We selected leaf traits of fresh mass, dry mass, water content, leaf area, perimeter, solidity, aspect ratio, circularity, LMA, lamina thickness, lamina density, lamina toughness, chlorophyll content per unit area and mass, trichome density, and ash content as representatives of leaf ecophysiological variation, including the leaf economic spectrum (Mason and Donovan, 2015). We used the eigenstructure created from 150 lines with complete dry mass data to impute missing mass-based trait data. From the imputed data, we derived PCs for use in a genome-wide association. However, the genome-wide association of individual traits was only conducted with the non-imputed data.

In addition, we separately analyzed the relationship between total phenolics and total flavonoids. As flavonoids are typically the most abundant type of phenolic compounds in plants, we wanted an indirect measure of the relationship between the production of flavonoid and non-flavonoid phenolics. We analyzed flavonoids and phenolics in a PCA approach where PC1(FLAVPHEN1) explains the correlation between total flavonoids and total phenolics, and PC2(FLAVPHEN2) explains the residual variance.

For secondary metabolism data derived from HPLC, we performed four PC analyses. The first two were based on: (1) peak area for all metabolites detected, and (2) the subset of metabolites found in at least 200 lines. The third and fourth analyses were based on relative ratios of peaks within each profile for: (3) all metabolites detected, and (4) the subset of metabolites found in at least 200 lines. We selected 200 lines as an arbitrary cutoff to include 70% of the lines in the panel.

For each analysis, we retained any PC with an eigenvalue greater than one for investigation. For each PC retained, we conducted two analyses evaluating mean differences between HA-maintainer (N = 125) and RHA-restorer (N=100) lines and Oil (N=134) and Nonoil (N=83) varieties leveraging Bayesian factor analyses, using the BayesFactor package in R version 4.0.2 (Morey et al., 2015; R Core Team, 2020). We report the mean and credible interval of the effect of HA-maintainer relative to RHA-restorer as the effect size estimated from sampling from the posterior distribution 10,000x and interpreted evidence support based on the criteria of Kass and Raftery (1995) (3-20 positive evidence, 20-150 strong evidence, BF >150 very strong evidence).

### Genome-Wide Association Mapping

We performed all association analyses using a modified pipeline described in Temme et al. (2020). for each trait collected and derived. Briefly, we performed all association analyses using GEMMA v0.98.1 (Zhou and Stephens, 2012). We only assessed models with corrections for kinship and population structure for each trait association. We leveraged GEMMA to compute kinship and used SNPRelate (Zheng et al., 2012) to estimate population structure from a subset of 24k independently inherited SNPs; the first four principal components from this latter analysis were used as the population structure metrics. We estimated the effective number of independent markers in the genome using ordered SNPs to construct a haplotype map from observed linkage disequilibrium (LD) patterns with PLINK v1.9 (Purcell et al., 2007). We identified haplotype blocks using D’ with a standard confidence interval of 0.7005 - 0.98, an informative fraction of 0.9, and a maximum block span of 100Mb to accommodate the largest observed linkage block in the genome (Gabriel et al., 2002). We calculated narrow-sense heritability (*h*^2^_SNP_) based on the estimated kinship expressed as the additive genetic variance divided by the observed phenotypic variance for each trait (Tropf et al., 2017).

To correct our significance thresholds for multiple comparisons, we used the approach of Gao (2011) using the number of multi-SNP haplotype blocks identified as the effective number of independent tests. While reducing the conservative nature of a Bonferroni multiple-testing correction, this type remains a highly conservative significance threshold dependent on genome assembly accuracy (Temme et al., 2020). For our threshold, we used the HA412HOv2 genome (Todesco et al., 2020). We identified genomic regions in LD for each significant marker-trait association based on the previous haplotype analysis. For interpretation, we assumed significantly associated SNPs indicative of the associated genomic region.

For model interpretation, we chose the maximum β value across all significantly associated SNPs in the corresponding associated region as the magnitude of the associated region’s effect on each trait of interest. We expressed relative effect size as (2\**β*)/(trait range) for each associated region (Masalia et al., 2018). We summed the relative effect size values across all associated regions per trait to estimate the significantly associated genomic regions’ relative effect size (RES). We considered regions associated with multiple traits as co-localized, whereby LD between associated SNPs is observed. Given our conservative threshold and other studies examining the conservative nature of the Gao (2011) approach and Bonferroni thresholds, we also examined suggestive colocalization cases based on occurrences of significantly associated regions for each trait containing SNPs with p-values < 0.0001 for any other trait (Gao et al., 2008, 2009; Gao, 2011). Furthermore, we extracted a list of genes within the region and one flanking gene from each side of the region based on the annotation of the HA412-HO genome for every associated region (Todesco et al. 2020).

In the trait associated regions with multiple genes, we used a gene set enrichment of GO terms approach to describe gene trends in the region. We used a Kolmogorov-Smirnov (KS) test to compute enrichment based on gene scores (Ackermann and Strimmer, 2009). In this instance, because for each trait, we had sets of suggestive (p < 0.001) and/or significant (p < gao) SNPs, we binarized the p-value of the mixed linear model results such that significant SNPs had a gene score of 1 and suggestive SNPs had a gene score of 0. We visualized the top 10 GO categories with the lowest KS *p*-value. Further, we summarized the results of the top 20 nodes reporting the number of significant genes associated with each GO term, and the KS p-value adjusted for multiple testing corrections based on the assumption that each gene is independent, as a conservative threshold. We conducted a secondary gene set enrichment analysis to assess if there were common GO categories across our entire analysis, which may indicate underlying integrative processes. In the secondary analysis, we combined all genes associated with our GWA of traits into a single data set, where all GO terms were analyzed in the same framework as individual traits.

Finally, all regions identified as significantly associated with various ecophysiological and specialized metabolic traits were cross-referenced with the 51 SPH-protein family genes in the annotated HA412-HO genome (Appendix S1). Code for our modified GWAS pipeline can be found at https://github.com/jordandowell/SAM_HPLC_LES_HeteoticGrouping.git and https://github.com/jordandowell/GWAS_pipeline.git. All analyses were completed in R 4.0.2 (R Core Team, 2020).

## Results

### Ecophysiology and Coarse Defense Trait Variation

Assessment of leaf ecophysiology by principal components analysis produced two easily interpretable axes (Figure 1, Appendix S2). The first axis (35.3% variance explained) represents a traditional leaf economic spectrum, while the second axis (20.2% variance explained) consists of a spectrum predictive of leaf size (Figure 1, Appendix S2). Positive values of PC1 indicate high LMA, lamina density, and toughness, while negative values indicate high water content, ash content, and chlorophyll content on a mass basis. This can be interpreted as more positive values indicating leaves built with relatively more carbon-based (and combustible) structural components, like cellulose or surface waxes, contributing to a high LMA, density, and toughness. At the other end of this spectrum, negative values indicate a relatively higher reliance on turgor and inorganic minerals for leaf structural support (as indicated by high water content and ash content), and likely higher nutrient content and productivity (as indicated by both ash content and mass-based chlorophyll content). With respect to the orthogonal second axis, positive values of PC2 indicate smaller leaves with lower mass and area, while negative values indicate larger leaves with higher mass and area.

**Figure 1.**
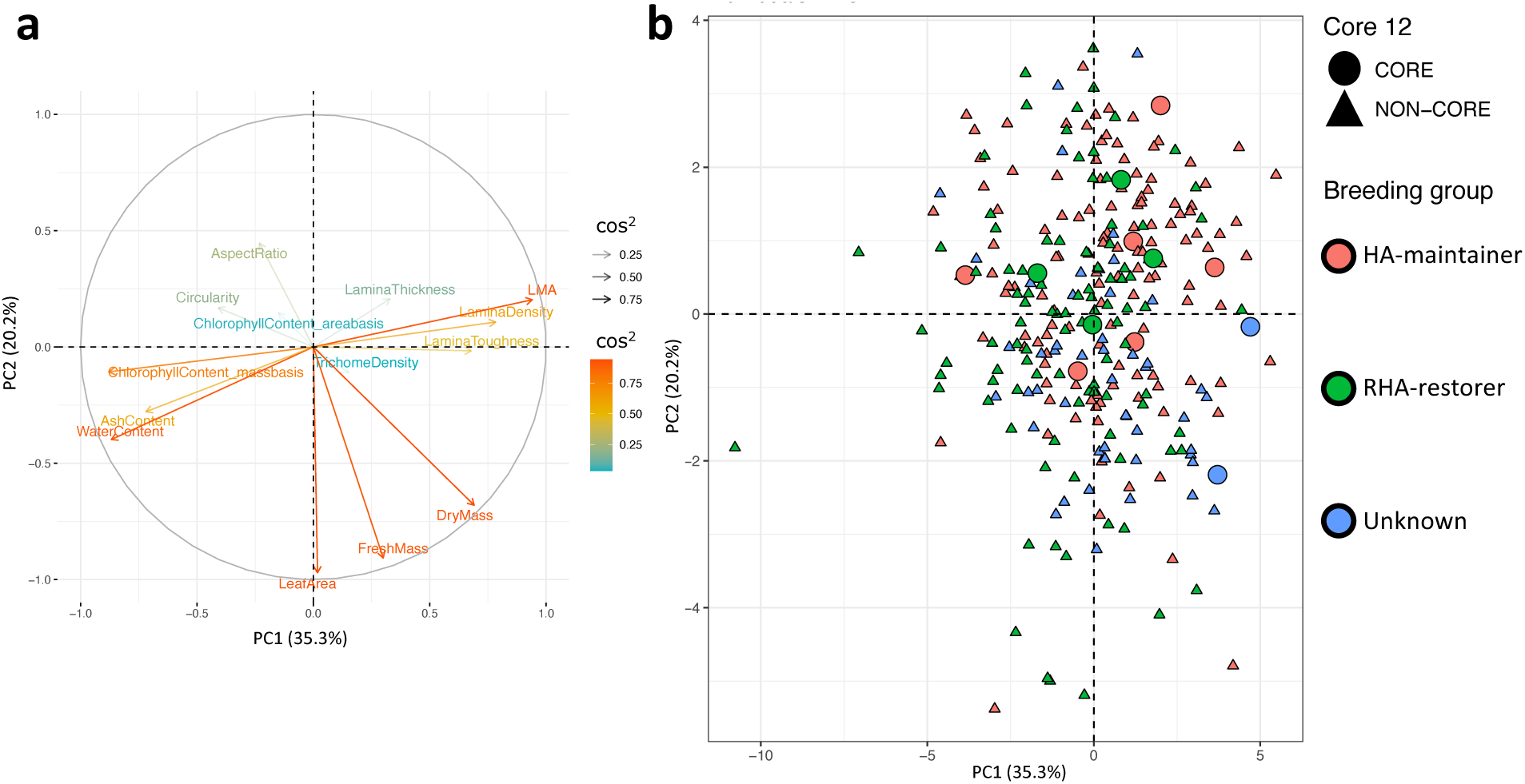
Principal component analysis of ecophysiological traits as a proxy for the leaf economic spectrum (LES) and leaf size across 288 cultivated *Helianthus annuus* genotypes. a) Variable loading plot for traits on the first two PCs. Vectors are color-coded from red to blue based on variable importance expressed as cos2 of the variable loadings along the first two axes. b) Individual loadings of cultivated *Helianthus annuus* genotypes along PC 1 and PC2. Colors indicate breeding groups, and shapes indicate whether individuals are a part of the ‘Core 12’ set of lines (circles), representing 50% of the genetic diversity of the SAM panel or not (triangles). Cos2 indicates variable importance derived from the vector loadings of traits on principal components 1 and 2.

Bayes factor analyses indicate strong support for differences between HA-maintainer (N = 125) and RHA-restorer (N =100) lines in leaf economics, as indicated by LES PC1 (BF 56) (Figure 1, Appendix S1, S2). HA-maintainer lines had on average more positive values along LES PC1 with a relative effect size of 10% (4 to 15%; 95% CI) for HA-maintainer identity compared to RHA-restorer (Appendix S1, S2). There was low support for differences between HA-maintainer and RHA-restorer lines in leaf size, as indicated by LES PC2 (BF 3) (Appendix S2). HA-maintainer identity had a relative effect size of 8% (2 to 15%; 95% CI) and more positive values along LES PC2 when compared to RHA-restorer (Appendix S1, S2). In contrast to RHA-restorer and HA-maintainer variation, Bayes factor analyses suggest strong differences along LES PC2 (BF=30) and low support for differences in LES PC1 (BF=1.1) between oil and confectionary (Nonoil) lines (Appendix S1, S2). Oil variety identity has a relative effect size of 13% (5 to 22%; 97.5% CI) for LES PC2 indicating that Oil varieties have on average larger leaves (Appendix S1, S2).. From the PCA individual loading plots, the positions of ‘Core 12’ lines separate based on RHA-restorer vs. HA-maintainer heterotic group identity; however, examination of all lines shows interspersion between RHA-restorer and HA-maintainer groups (Figure 1).

### Specialized Metabolite Variation

Among the HPLC-MS data, we identified 23 different chromatographic peaks present in 6 or more of the ‘Core 12’ lines, with putative identities belonging to 15 different classes (Appendix S2). Although we used the same liquid-chromatograph coupled to different detectors, and the data between detectors should be highly correlative, we erred on the side of caution in ascribing identity to HPLC-UV data based on putative HPLC-MS identity. From the HPLC-UV data, we identified a total of 106 peaks across 288 lines, with 32 ‘common’ metabolites present in at least 70% of lines (200 of 288 lines). All peaks found in the HPLC-MS data have corresponding peaks in HPLC-UV data based on retention time similarity. However, only 11 of the 23 peaks (6 of 15 classes) identified with HPLC-MS were recovered across at least 70% of lines with HPLC-UV (Appendix S2).

Principal component analyses of relative ratios of metabolites compared to analyses of peak area demonstrated no noticeable differences in resulting analyses, apart from variation in rotation (Figure 2, Appendix S2). However, after trimming the dataset to common metabolites observed in at least 70% of lines (≥200), we observed a more than doubling of variance explained by the first PC from 19.4% to 42.6% in analyses of both relative ratios and peak area of metabolites (Figure 2, Appendix S2). From the PCA individual loading plots, lines separated based on RHA-restorer or HA-maintainer heterotic group identity (Figure 2, Appendix S2).

**Figure 2.**
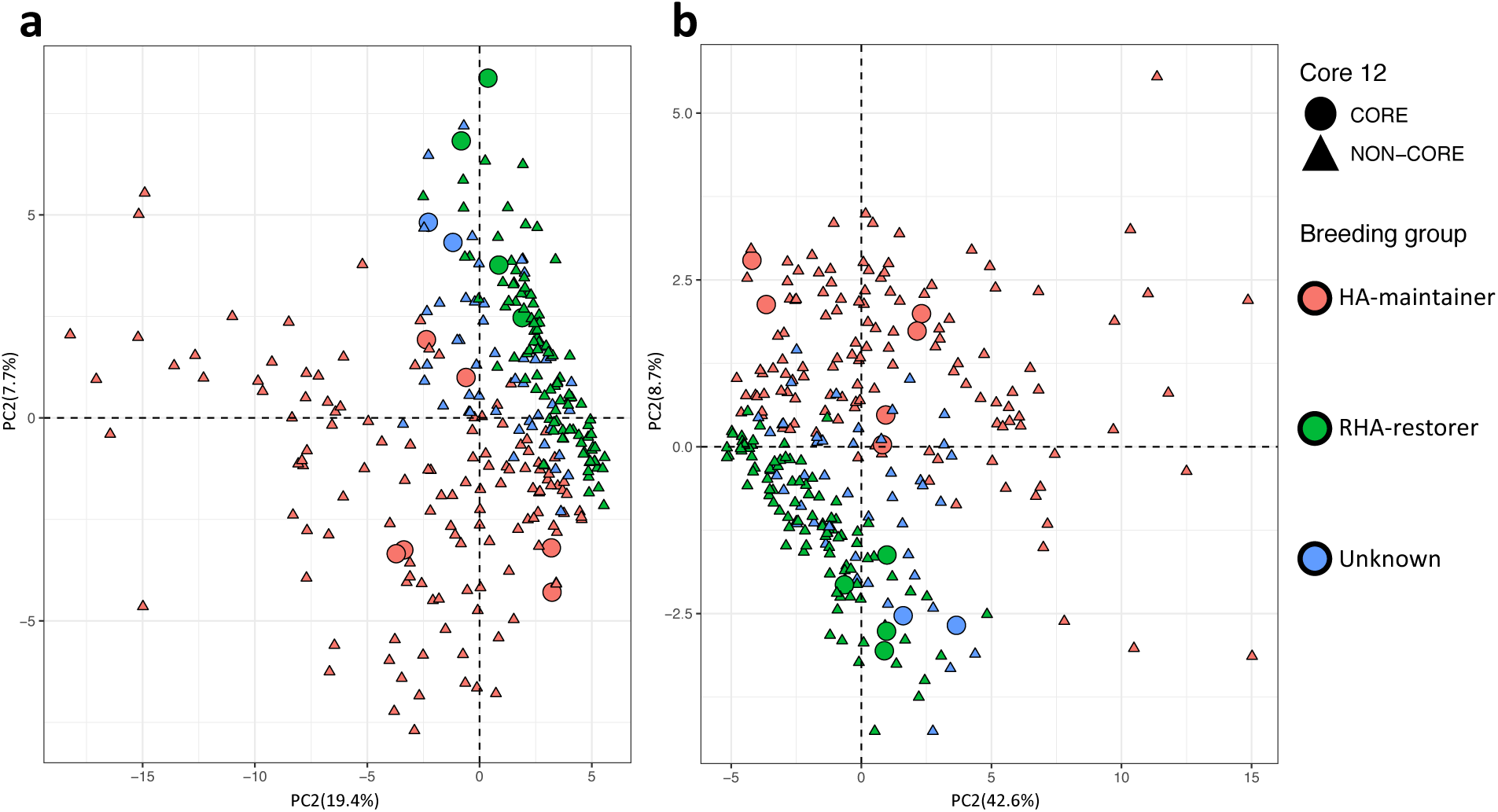
Principal component analysis of specialized metabolites peak area of 288 cultivated *Helianthus annuus* genotypes. a) Individual loadings of cultivated *Helianthus annuus* genotypes along PC 1 and PC2 of the 106 metabolites identified, including unique metabolites. b) Individual loadings of cultivated *Helianthus annuus* genotypes along PC 1 and PC2 of the 32 metabolites found across all genotypes. Colors indicate breeding groups, and shapes indicate whether individuals are a part of the ‘Core 12’ set of lines (circles), representing 50% of the genetic diversity of the SAM population or not (triangles).

Examination of all lines shows greater dispersion among USDA lines than INRA lines (Figure 2). Lines with the oldest plant introduction dates (landraces and open-pollinated lines) show an intermediate phenotype between RHA-restorer and HA-maintainer lines (Figure 2). In addition, INRA lines are more similar to the intermediate phenotype in comparison to USDA lines (Figure 2). Variable loadings indicate differences between PC 1 and 2 among principal component analyses of all and common metabolites. Considering all metabolites, 90% of variable loadings for PC 1 were negative, whereas when considering only common metabolites 99% of variables had positive loading values. In view of the observed rotational differences between the PCAs of all metabolites and common metabolites, this indicates PC 1 corresponds to quantitative variation in overall concentrations of specialized metabolites. Among PCAs, the individual loadings indicating high compound concentrations were exclusively among HA-maintainer lines and the lowest concentrations among RHA-restorer lines. PC 2 in all metabolite analyses showed more nuanced variation, although PC 2 explains less than 10% of the variation in all PCs. In considering all metabolites, positive values of PC 2 indicate high concentrations of flavonoids (e.g., Desmethoxysudachitin, Apigenin, 6,4’-dimethoxy, Jaceosidin), and negative values indicate high concentrations of sesquiterpene-lactones (e.g., Eupasserin trihydroxyangelate, Tifruticin, 1-hydroxy, Cumambranolide 2-hydroxyethyl acrylate, and Cumambranolide angelate) (Figure 2). The opposite pattern was observed when considering the PCA performed on common metabolites, in line with the rotational differences observed (Figure 2).

Bayesian factor analyses indicate strong support for differences in secondary metabolism between HA-maintainer and RHA-restorer lines, using analyses of metabolite PCs (BF >1000 for all PCs). With respect to HA-maintainer identity compared to RHA-restorer identity, in the all metabolites PCA there was a relative effect size of −24% (−17% to −31%; 95% CI) for PC1 and - 23% (−16% to −30%; 95% CI) for PC2 (Appendix S1, S2). Among common metabolites, HA-maintainer identity had a relative effect size of 17% (10% to 24%; 95% CI) for PC1 and 24% (19% to 29%; 95% CI for PC2 (Appendix S1, S2). This indicates that varieties in the HA-maintainer pool have on average higher concentrations of specialized metabolites than the RHA-restorer pool, and also indicates that the pools have a difference in relative balance of metabolite classes, with the HA-maintainer pool skewed toward more sesquiterpene lactones and the RHA-restorer pool skewed toward more flavonoids. Like breeding pools, Bayes factor analyses indicate strong support for difference between Oil and Nonoil varieties (BF > 1000). Nonoil identity has a relative effect of −23% (−16 to −29%; 95% CI), indicating that Nonoil varieties have, on average, a greater concentration of specialized metabolites than Oil varieties (Appendix S1, S2). There was no support for qualitative differences in specialized metabolite variation between market types (BF = 0.33) (Appendix S1, S2). Line means and Bayes factor results can be found in Appendix S1.

### Genome-Wide Association Mapping

We included kinship as a random effect for our mixed linear model approach and 4 PCs denoting population structure as fixed effects. We iteratively ran this model for each of the 1.4 million SNPs across the sunflower genome for all 128 measured and/or derived traits, including leaf ecophysiology and specialized metabolism.. In total, we identified 96 linkage blocks significantly associated with one or more traits across all chromosomes other than chromosome 1. These regions contained 887 unique significantly associated SNPs and 3,707 annotated genes. Among significant SNPs, 27 were associated with more than one trait, these traits being exclusively specialized metabolites (Appendix S2). Among linkage blocks, significant blocks had, on average, seven associated traits, and the relative effect size (RES) for any given linkage block varied from 0.04 to 0.45 (Appendix S2). Water content in linkage block 17-01 had the highest RES, although it was not significantly associated above our suggestive threshold (*p*<0.0001). Region 17-01 was significantly associated with leaf dry mass (RES = 0.19), aspect ratio (RES = 0.20), and RT 8.81-UNK (RES = 0.31). The directionality of the effects for aspect ratio and dry mass were inverse such that as leaves increased in width relative to length, their dry mass was reduced, alongside the quantity of RT 8.81-UNK (Figure 4). Among linkage blocks with significant SNPs, the additive relative effect sizes for individual traits ranged from 0.07 to 3.39. A few specialized metabolites (RT 7.71-HCA, 7.47-UNK, 4.21-UNK, 3.52-UNK, and 8.81-UNK) had large individual relative effect sizes that summed to greater than one (Appendix S2). Large relative effect sizes may occur if the effect of multiple regions compensates for opposing effects of population structure or kinship on the trait or if multiple regions have large effects. A broad range of narrow-sense heritability values was identified (*h*^2^ 0.00-0.43), with the traits of lamina thickness (0.43), chlorophyll content (0.40), and leaf area (0.40) having among the highest heritability values (Appendix S2). Among traits, heritability values were uncorrelated to RES (*p* > 0.01) (Appendix S2). Among All traits, the directionality of effects coincided with expectations based on relationships identified in PCAs of ecophysiological traits and specialized metabolites (Figure 1,2,3,4).

**Figure 3.**
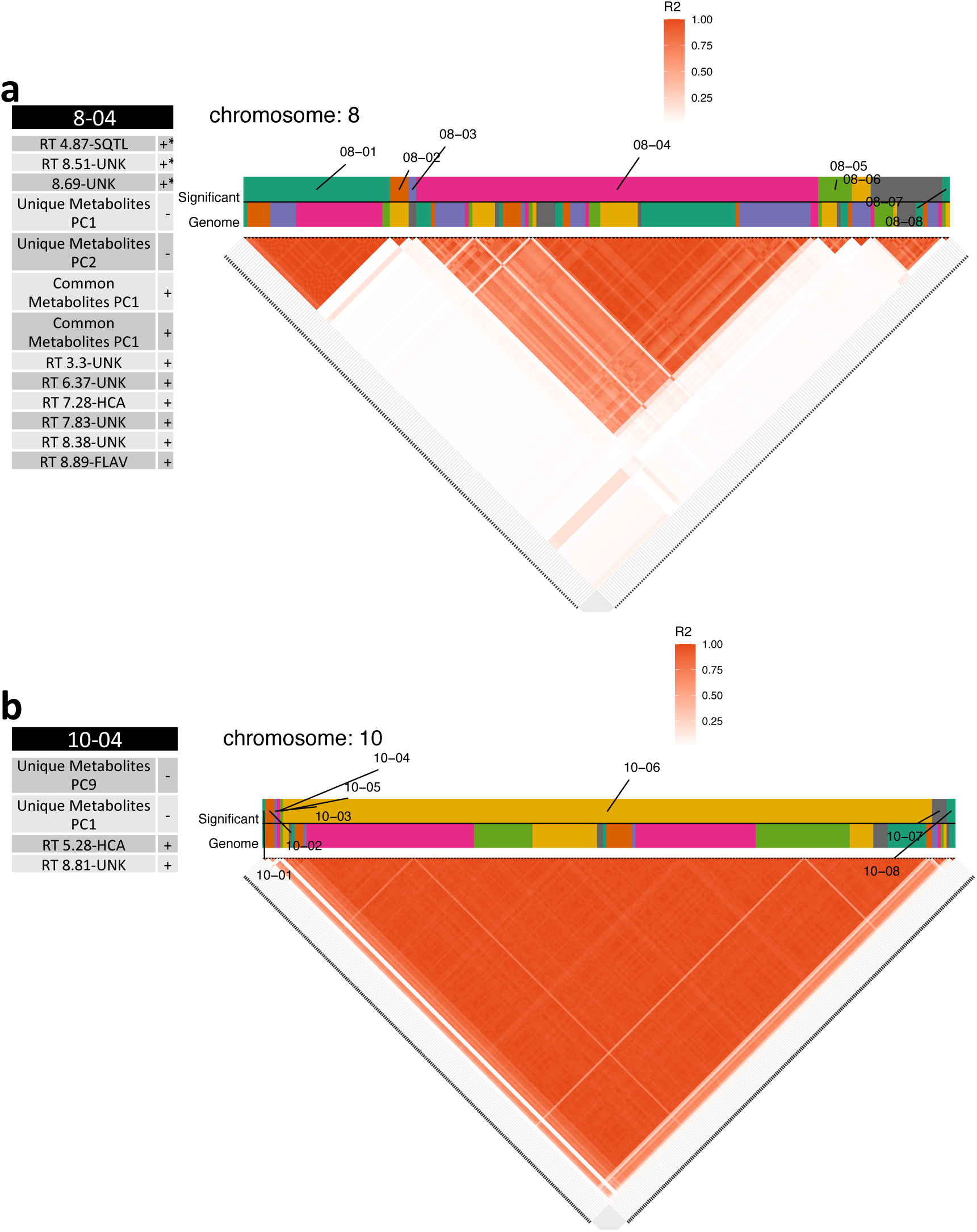
Sunflower GWAS and trait colocalization analyses for chromosomal haplotype blocks with annotated self-compatibility genes. a) Significant SNPs across all traits on chromosome 8 correspond to 48 unique haplotype blocks. However, after reassessing haplotype blocks using only significant SNPs, eight blocks remained based on observed patterns of LD, and only block 08-04 contained a self-compatibility protein homolog (SPH) and a significantly associated SNP. Trait colocalization in haplotype block 8-04 (left) has three instances of genomic overlap among traits with significant SNPs (*) and ten instances of overlap that are suggestive for a trait. A plus (+) or minus (−) indicates the sign of the effect of the minor allele on the trait. b) significant SNPs across all traits on chromosome 10 correspond to 29 unique haplotype blocks. However, after reassessing haplotype blocks using only significant SNPs, eight blocks remained based on observed patterns of LD, and only block 10-04 contains a self-compatibility protein homolog (SPH) and a significantly associated SNP. Trait colocalization in haplotype block 10-04 (right) has one instance of genomic overlap among traits with significant SNPs (*) and three instances of overlap that are suggestive for a trait. A plus (+). or minus (−) indicates the sign of the effect of the minor allele on the trait. Adjacent blocks are colored alternatingly, repeating the same seven colors. R^2^ is based on linkage disequilibrium between SNPs

**Figure 4.**
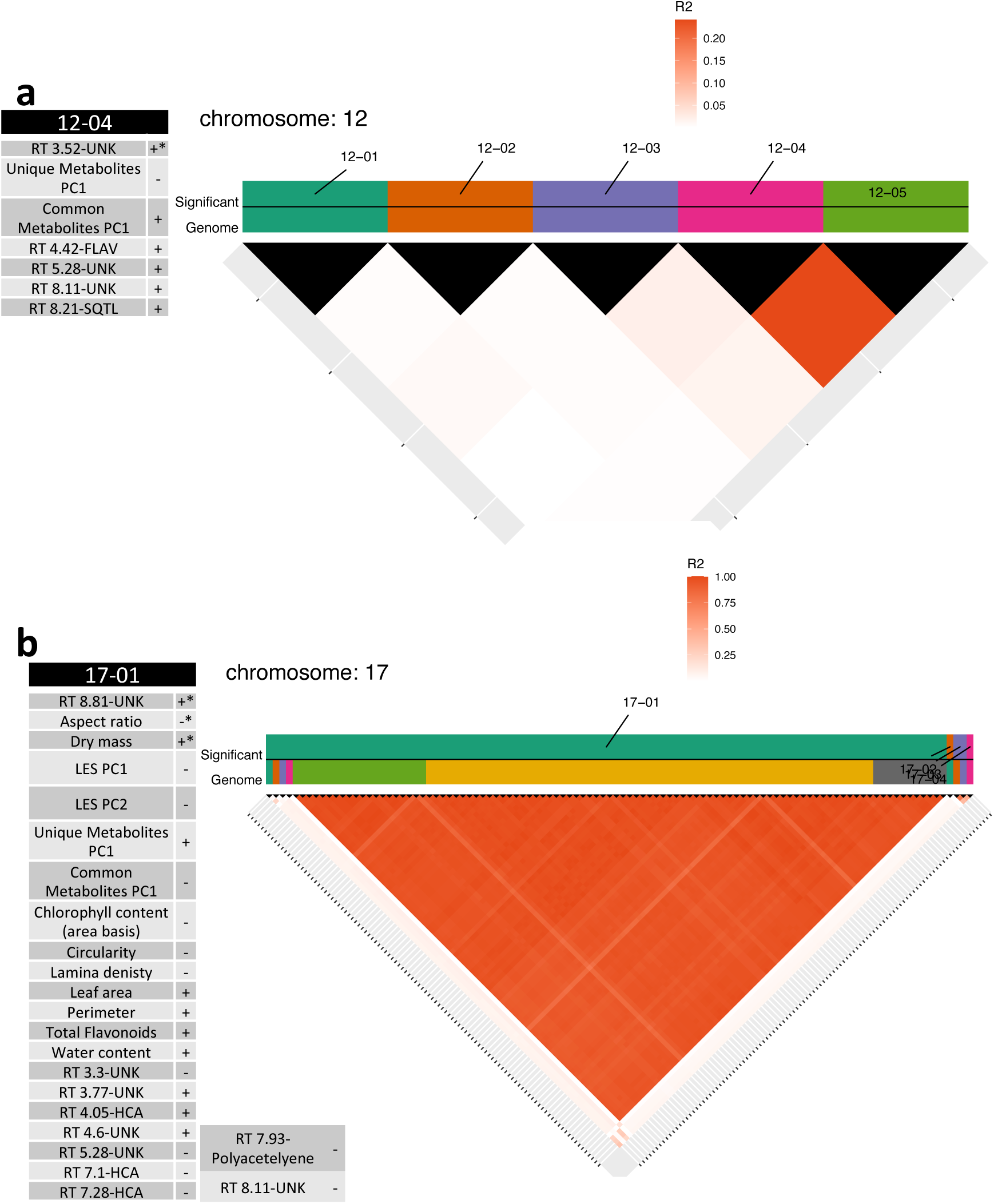
Sunflower GWAS and trait colocalization analyses for chromosomal haplotype blocks with annotated self-compatibility genes. a). Significant SNPs across all traits on chromosome 12 correspond to 5 unique haplotype blocks. After reassessing haplotype blocks using only significant SNPs, five blocks remained based on observed patterns of LD, and block 12-04 contains a self-compatibility protein homolog (SPH) and a significantly associated SNP. Trait colocalization in haplotype block 12-04 (left) has one instance of genomic overlap among traits with significant SNPs (*) and six instances of overlap that are suggestive for a trait. A plus (+) or minus (−) indicates the sign of the effect of the minor allele on the trait. b) Significant SNPs across all traits on chromosome 17 correspond to 11 unique haplotype blocks. However, after reassessing haplotype blocks using only significant SNPs, four blocks remained based on observed patterns of LD, and only block 17-01 contains a self-compatibility protein homolog (SPH) and a significantly associated SNP. Trait colocalization in haplotype block 17-01 (right) has three instances of genomic overlap among traits with significant SNPs (*) and 21 instances of overlap suggestive for a trait. A plus (+) or minus (−) indicates the sign of the effect of the minor allele on the trait. Adjacent blocks are colored alternatingly, repeating the same seven colors. R^2^ is based on linkage disequilibrium between SNPs.

From the annotation of the HA412-HO genome, there are 51 total annotated SPH-protein homologs (Hübner et al., 2019). Our analyses found linkage between significantly associated regions and 11 of the 51 SPH-protein homologs. We identified these associations on chromosomes 8, 10, 12, and 17. The linkage block for chromosome 17 (17-01) contains the most SPH-protein homologs at 7, while linkage blocks on chromosomes 12 (12-04), 10 (10-04), and 8 (8-04) have 2, 1, and 1 SPH-protein homologs, respectively. Region 8-04 (1 SPH) was significantly associated with metabolites RT 8.69-UNK, RT 8.51-UNK, and RT 4.87-UNK, with a max RES for each compound of 0.19, 0.17, and 0.11 (Figure 3). Region 10-04 was significantly associated with PeakareasPC9, driven by variation in RT 8.81-UNK. Region 10-04 (1 SPH) is also suggested for RT 8.81-UNK with a RES of 0.26 and PC1 which describes quantitative variation in specialized metabolites (Figure 3). Region 12-04 (2 SPH) is significantly associated with RT 3.52-UNK with a RES of 0.15 (Figure 4). The final region is 17-01 with 7 SPH-protein homologs, is associated with RT 8.81-UNK, aspect ratio, and dry mass (RES; 0.31, 0.2, 0.19), and suggestively associated (Gao threshold < p < 0.0001) with 46 other chemical and ecophysiological traits (Figure 4).

Among specific traits, the genes associated with leaf dry mass were significantly enriched (*p*<0.01) for the biological processes of hydrogen peroxide catabolism (GO:0042744), response to oxidative stress (GO:0006979), response to far red light (GO:0010218), oligopeptide transmembrane transport (GO:0035672), lignin catabolic processes, regulation of transcription (GO:0006355), and various cellular responses to small molecules (GO:0048522, GO:1901701, GO:0071310, and GO:0010243). Similarly, molecular functions were enriched (*p*< 0.01) for polysaccharide binding (GO:0030247), acetyl-CoA carboxylation (GO:0003989), hydroquinone:oxygen oxidoreductase activity (GO:0052716), calcium (GO:0005509), copper (GO:0005507), and flavin mononucleotide binding (GO:0010181), oligopeptide transmembrane transport (GO:0035673), and peptide:proton symporter activity (GO:001533). Further, all 32 common metabolites assessed had significantly associated regions across the genome with individual region RES values ranging from 0.05 to 0.37. In each of these regions, there were several occurrences of annotated genes known from previous work to be related to plant immunity and pathogen perception, such as receptor-like protein kinases, Gnk2 proteins, wd40 proteins, xenobiotic transporters, and generalized disease resistance genes (Miller et al., 2016; Borrelli et al., 2018; Delgado-Cerrone et al., 2018). Among metabolites able to be given a putative identity through MS data, we found no significant associations between metabolites and regions containing terminal and/or precursor enzymes necessary for their production. In examining the GO results for regions associated with metabolites, many were enriched for redox reactions, metal ion binding, and transport. For instance, genes associated with RT 8.81-UNK had enriched molecular functions (p<0.001) for peroxidase activity (GO:004601), alcohol-forming fatty acyl-CoA reductase activity (GO:0080019 and GO:0102965), heme binding (GO:0020037), and iron and manganese ion transmembrane transporter activity (GO:0005381 and GO:0005384). Similarly for biological processes of RT 8.81-UNK, redox reactions (GO:0055114), response to oxidative stress (GO:0006979), hydrogen peroxide catabolism (GO:0047244), wax biosynthesis (GO:00010025), and general cellular homeostasis (GO:0019725) were enriched in genes associated with RT 8.81-UNK (*p*<0.01). Among all genes associated with traits, biological processes were enriched for fatty acid biosynthesis (GO:0006633), lignin catabolic processes (GO:0046274), protein ubiquitination (GO:0016567), positive regulation of transcription (GO:0045893), chromatin organization (GO:0006325), and response to oxidative stress (GO:0006979). Molecular functions were enriched (*p*<0.01) for U5 (GO:0030623) and U6 snRNA binding (GO:0017070), amino acid transmembrane transport (GO:0015171), RNA-directed DNA polymerase activity (GO:0003964), metalloendopeptidase activity (GO:0004222), oxioreductase activity (GO:0016717), hydroquinone:oxygen oxidoreductase activity (GO:0052716), peroxidase activity (GO:004601), metal ion binding (GO:0046872), and polysaccharide binding (GO:0030247). Although each of these traits, as mentioned earlier, have other significantly associated regions across the genome, we only find annotated terminal enzymes for specialized metabolic products such as vinorine (indole-alkaloid), kaempferol (flavonoid), and secologanin (secoiridoid monoterpene) in regions 8-04, 10-04, 12-04, and 17-01. All results can be found in supplementary appendices S1 and S2.

## Discussion

### Heterotic group variation

Leaf ecophysiological traits showed a divergence between HA-maintainer and RHA-restorer heterotic groups (Figure 1, Appendix S2). On average, HA-maintainer lines exhibited a ‘slower’ leaf economic strategy compared to RHA-restorer lines’ ‘faster’ leaf economic strategy (Figure 1, Appendix S2). "Fast" leaf economic strategies are reminiscent of initial breeding targets for total yield; resource-acquisitive trait strategies support fast plant growth rates and large plant sizes needed for high seed output. In comparison, HA-maintainer lines exhibited a ‘slower’ leaf economic strategy, with a larger investment into carbon-based structural or defensive elements. Plants that exhibit slower leaf economic strategies are predicted to be more resistant to abiotic and biotic stressors (Wright et al., 2004; Mason et al., 2016; Züst and Agrawal, 2017), in line with breeding targets towards yield regularity. The slow transition of breeding targets from yield quantity to yield regularity alongside the male-sterile and maintainer bias in breeding programs may be manifesting as a pattern of ‘phenotypic lag,’ with RHA-restorer lines exhibiting trait syndromes indicative of older total yield breeding targets and HA-maintainer lines exhibiting traits consistent with more recent breeding targets of yield regularity and stability (Adeboye et al., 2014; Salazar et al., 2016; Radanović et al., 2018). The existence of ecophysiological trait variation within and among heterotic groups, coupled with the retention of functional genetic diversity, demonstrates the potential for improvement in ecophysiology-mediated performance leveraging current cultivated genetic resources.

In comparison to leaf ecophysiological traits, there is a more robust separation between HA-maintainer and RHA-restorer breeding pools with respect to leaf specialized metabolite profiles (Figure 2, Appendix S1, S2). The pattern is consistent for both common and unique metabolites, representing quantitative and qualitative differences between specialized metabolite profiles of maintainer and fertility-restoring lines (Figure 2, Appendix S1, S2). Our findings that HA-maintainer lines exhibit both higher concentrations of specialized metabolites and a more "resource conservative" leaf economic strategy further supports the idea that quantitative chemical defenses may be an important contributor to observed leaf economic strategy (Mason and Donovan, 2015).

HA-maintainer lines demonstrate a greater specialized metabolite diversity among both unique and common metabolites, observable by the greater share of multivariate phytochemical space occupied compared to RHA-restorer lines (Figure 2). Both qualitative and quantitative diversity in specialized metabolites is tightly linked to disease and pest resistance in plants (Kessler, 2015; Vear, 2016; Kessler and Kalske, 2018). As such, we can leverage specialized metabolite profile variation and quantitative differences as a proxy for variation in resistance. Our findings suggest that HA-maintainer and RHA-restorer pools may differ in their resistance to various pests and pathogens. Previous work in wild *Helianthus* has found that high concentrations of nonvolatile specialized metabolites predict resistance to both aphids and powdery mildew (Mason et al., 2016). Quantitative increases in specialized metabolites among HA-maintainer lines suggest they may be, on average, more resistant to pests and pathogens than their RHA-restorer counterparts. In the context of industrial hybrid seed production, RHA-restorer and CMS lines are planted in row ratios of 2:6 to 2:14 (Long et al., 2019).

Provided that CMS lines and their associated HA-maintainer lines are phenotypically similar one may surmise that, at the field scale, row ratios may influence the impact of pests or pathogens as increases in plant intraspecific chemotypic diversity in natural settings reduces herbivory (Ziaja and Müller, 2023). Although, to the author’s knowledge, there is no reported impact of row ratio in pest or pathogen damage in hybrid seed production fields, an examination is warranted. In the field, if there is a link between row ratio and line chemodiversity affecting pest or pathogen damage, hybrid seed production may be able to reduce pesticide use by leveraging chemotypically complementary HA-maintainer and RHA-restorer lines during production to increase field chemodiversity. Thus, there may be benefits to a broad examination of multi-line combinations for variation in antagonistic and synergistic impacts on quantitative biotic stress resistance beyond the obvious goals of identifying individual lines with reduced susceptibility. In particular, understanding how underlying plant ecophysiology and specialized metabolism influence the capacity for biotic stress resistance at the individual level and scale to the community level may provide useful insights into potential constraints on cultivar and industrial-scale agricultural systems improvements – including those derived from trait trade-offs and baggage from historical breeding practices.

### Trait Genetic Architecture

We identified many loci associated with several specialized metabolites and observed variable effects among loci, contributing to the growing body of literature documenting functional genomic variation within cultivated sunflower (Masalia et al., 2018; Dowell et al., 2019; Temme et al., 2020). Most specialized metabolites showed an observed oligogenic architecture consisting of few loci scattered across the genome with small effect sizes, which may indicate a more complex architecture not able to be fully described through association mapping and GWA using traditional linear mixed modeling approaches, in line with many GWAS assessments of specialized metabolites (Chan et al., 2011; Kumar et al., 2015; Alves et al., 2020; Chen et al., 2020). Like studies of maize, wheat, and apples, regions associated with specialized metabolites in our study included few genes encoding terminal enzymes in biosynthetic pathways (Kumar et al., 2015; Alves et al., 2020; Chen et al., 2020). However, our study also documented co-localization of genes associated with the perception of pests and pathogens to association with constitutive variation in leaf specialized metabolites.

The low heritability observed for concentrations of most specialized metabolites, coupled with a few metabolites (RT 7.71-HCA, 7.47-UNK, 4.21-UNK, 3.52-UNK, and 8.81-UNK) with large effect SNPs, suggests that much of the variation in constitutive compound production among cultivated sunflower lines may be the product of recent phenotypic convergence (Appendix S2). Recent convergence on an observed phenotype and allelic combination through independent evolutionary routes would explain low heritability, as the variation in the phenotype explained by population parameters explains minor variation compared to the specific SNP, as observed for glucosinolates in wild *Arabidopsis thaliana* (Katz et al., 2021), seed color in *Phaseolus vulgaris* (McClean et al., 2018), and highland adaptation in maize (Takuno et al., 2015). Independent evolutionary routes in this context may be due to introgression and breeding history combinations. While introgression in modern cultivated sunflower shows evidence of linkage drag in course phenotypes like yield (Huang et al., 2023), this observation may be due to increased investment in specialized metabolites, as evidenced by HA-maintainer lines increases in specialized metabolite concentrations relative to RHA-restorers or other open-pollinated varieties and landraces. Thus, population genomic investigations into observed compound variation may lead to the identification of resistance-associated specialized metabolites by examining links between the inheritance of allelic combinations and associations with compound variation in the context of changes to yield. In support of the potential for recent convergence, many regions associated with specialized metabolites were enriched with genes relating to common molecular functions like redox activity and metal binding. Furthermore, our results indicate that variation in the constitutive production of specialized metabolites is linked to variation in perception genes among cultivated sunflower lines. Thus, targeted breeding efforts for constitutive production of specialized metabolites may benefit from modifying perception genes as an alternative to directly focusing on biosynthetic enzymes.

Among leaf ecophysiological traits, there were few significant associations; however, many traits had high heritability values. The polygenic nature of these traits suggests that the detailed genetic architecture of ecophysiology is likely only partially observable through traditional mixed linear model frameworks for this panel (Zhou et al., 2013). However, examination of regions associated with ecophysiological traits and specialized metabolite traits indicated a relationship between transcriptional regulation and responses to oxidative stress.

Further, in line with sunflower breeding targets, fatty acid production and lignin catabolism are associated with many traits, indicating an underlying relationship in this mapping population. Other sunflower studies have demonstrated oligogenic architectures for responses to biotic and abiotic stress, in contrast to the results observed here for constitutive ecophysiological trait variation under high-resource conditions (Mestries et al., 1998; Micic et al., 2004; Masalia et al., 2018; Temme et al., 2020). This pattern suggests that the genetic architecture of constitutive variation in leaf ecophysiology may be more complex than that of plastic or inducible trait responses to stress. Induction of traits that ameliorate stress is in line with current breeding targets towards yield regularity and should be prioritized, provided that reactions to abiotic stresses reduce the diversion of resource investment away from yield over the lifespan of the crop (Vincourt, 2014; Gage et al., 2017; Masalia et al., 2018; Temme et al., 2020). Given our findings of divergence in constitutive ecophysiology between the heterotic breeding groups, future studies should explicitly consider the potential effects of historical breeding practices on such stress-inducible trait variation.

Heterosis is a major goal of hybrid breeding practices in sunflower and many other crops (Goff, 2011; Labroo et al., 2021; Wu et al., 2021). Among the general models to explain heterosis (epistasis, overdominance, and dominance), in sunflower, support for the dominance model of heterosis has been primarily observed for morphological and metabolic traits, indicating that heterotic groups were not selected for overdominance (Encheva et al., 2015; Owens et al., 2019). In expanding models of heterosis beyond allelic heterozygosity, studies have incorporated assessing genomic structural variation and links to heterosis (Song et al., 2020; Lee et al., 2022; Bie et al., 2023; Wang et al., 2023). Considering modern breeding programs, the introgression of landraces and wild species into elite breeding pools may increase genomic structural variation through increased presence/absence variants (PAVs), copy number variants, and other forms of cryptic heterozygosity. In cultivated sunflower, expression complementation of PAVs contributes to heterosis in coarse traits like total biomass and height (Lee et al., 2022). However, PAVs have little effect on more fine-scale ecophysiological and metabolic traits such as oil content and specific leaf area (Lee et al., 2022).

In contrast to direct links, in sunflower, PAVs find utility in genomic prediction, accounting for substantial variation in various traits of sunflower F1 hybrids (Lee et al., 2022). The discrepancy may indicate the utility of PAVs to explain subtle population parameters and not necessarily associations with trait-level variation as opposed to expression complementation. Further, the inclusion of the complexity of structural variation and potential cryptic heterozygosity make association mapping within this population difficult due to the current state of sequencing depth (Lee et al., 2022). However, future work aimed at examining the role of introgression and genomic structural variation in relation to leaf ecophysiological and specialized metabolic variation may benefit from examining explicit syntenic regions to preclude overestimation of effects confounded with cryptic heterozygosity and population parameters (Jaegle et al., 2021; Lee et al., 2022).

### Linkage Disequilibrium

Ours is not the first study to document or suggest the unintentional divergent selection of traits during modern crop breeding and improvement (Breseghello and Coelho, 2013; Shi et al., 2015; Carisse et al., 2017; Vilela et al., 2018). However, we here leveraged trait-associated linkage groups to generate hypotheses of potential processes by which selection may have acted during trait divergence. This suggests that one potential process may involve loci related to self-incompatibility. Interestingly, in contrast to our findings for several traits in linkage with SPH-protein homologs, we did not find nuclear genes related to CMS fertility restoration in any region associated with specialized metabolites or ecophysiological traits. Our findings suggest that observed trait divergence between HA-maintainer and RHA-fertility restorer heterotic groups does not appear to be driven by physical linkage with nuclear CMS fertility restoring loci but rather may be a side effect of different selective pressures within the two breeding groups or even genetic drift due to reduced gene flow between the two germplasm pools.

In crops where the degree of self-compatibility is variable and has a major influence on yield, linkage between regions contributing to self-compatibility and trait-associated genes may result in trait divergence during selection for improved yield (Brouwer and St. Clair, 2004; Dirlewanger et al., 2004; Gandhi et al., 2005; Kardile et al., 2022; Laugerotte et al., 2022). We identified 20% of the SPH-protein homologs within the HA412-HO reference genome in linkage disequilibrium with SNPs associated with specialized metabolites and ecophysiological traits. Associated traits were primarily specialized metabolites, except for leaf dry mass and aspect ratio (important descriptors of leaf size and shape). Evidence from QTL and expression studies provide correlative evidence as to the potential role of SPH-proteins in contributing to the degree of self-compatibility in cultivated sunflower (Burke et al., 2002; Gandhi et al., 2005; Badouin et al., 2017). As one of many potential unexplored molecular mechanisms that may influence the degree of selfed seed production, SPH-proteins’ role in self/nonself recognition and self-compatibility requires mechanistic validation.

Further inquiry into the potential mechanisms of self-incompatibility in cultivated sunflower through reverse genetic approaches (de Graaf et al., 2012; Lee et al., 2020; Lin et al., 2022; Shin et al., 2022) and protein-protein interaction simulations (Ma et al., 2016; Rajasekar et al., 2019) may expand selective targets for cultivated sunflower yield in a mechanistic manner, given the relationship between selfed seed production and yield (Burke et al., 2002; Gandhi et al., 2005; Vleugels et al., 2019). Finally, while cultivated sunflower is largely self-compatible (though variable in the selfed seed production rate), wild populations of *H. annuus* and almost all of the rest of the genus *Helianthus* are self-incompatible. Thus, most SPH-protein homologs should be largely nonfunctional or not expressed in cultivated sunflower. Exploration of the functionality of the SPH-protein family across the genus may provide further insight into the potential roles of this family in mediating self-incompatibility and self/nonself recognition. Lastly, expanding beyond the SPH-protein family, detailing the explicit molecular mechanisms of quantitative and binary self-incompatibility phenotypes is necessary for sunflower and, when paired with population genomic information available, may provide further insights into the mode and tempo of phenotypic divergence in this group.

## Supporting information

All Appendicies

## Acknowledgements

The authors wish to thank Michael Boyd, Caitlin Ishibashi, Seth Bradley, Gaven Meyers, and Erin Clark for contributions to plant growth, leaf sampling, and/or trait data collection. This research was funded in part by an NSF Doctoral Dissertation Improvement Grant (DEB-1404291) to CMM, as well as NSF grant IOS-1122842 to LAD. CMM was supported by a dissertation completion award from the UGA Graduate School during greenhouse data collection. JAD was supported in part by the Bill and Melinda Gates Millennium Foundation and McKnight doctoral fellowship.

## Author Contributions

CMM, AWB, JMB, and LAD designed the experiment. AWB, AJ, RS, and CMM collected data. AWB, LAD, and CMM contributed to phenotypic data analysis, with final phenotypic data analysis performed by JAD. JAD performed genome-wide association mapping and related statistical analyses. JAD wrote the manuscript with input from CMM, AJ, RS, AWB, JMB, and LAD.

## Data Availability

Trait data is available in the supporting information and will be submitted to the Dryad digital repository upon acceptance.

## Supporting Information

**Appendix S1.** Table of common metabolites in the majority of ‘core 12’ lines, box plots of PCA individual loadings, and comparisons of heritability and additive relative effect sizes among traits.

**Appendix S2.** Datasets consisting of SAM panel metadata, Self-incompatibility protein homolog (SPH) genome locations, summary statistics of measured and derived traits overall, summary statistics of measured and derived traits by heterotic group and use type, results of BF analysis for differences between group trait distributions, Genome haplotype block sizes, Summary of GWAS results for all traits, Gene lists for all significantly associated traits, and a list of SNPs and associated with individual and derived traits, and the genome annotation used for GWAS interpretation.

**Appendix S3.** Results and figures for GO-term analyses for individual and derived traits.

