## Supplementary material for "Historic breeding practices contribute to germplasm divergence in leaf specialized metabolism and ecophysiology in cultivated sunflower (*Helianthus annuus*)": All Appendicies: Appendix_S1.docx

**Table S1** Metabolites detected in a majority of the ‘core 12’ lines with HPLC-MS.

**Figure S1** Boxplots of leaf economics spectrum principal components values among HA-maintainer and RHA-restorer lines and Nonoil and Oil for PC2

**Figure S2.** Principal component analysis of the relative ratios of specialized metabolites among 288 cultivated *Helianthus annuus* varieties.

**Figure S3** Boxplots of specialized metabolite peak area PC values among HA-maintainer and RHA-restorer lines and Oil and Nonoil lines for 106 metabolites identified across genotypes, including unique metabolites.

**Figure S4** Boxplots of specialized metabolite peak area PC values among HA-maintainer and RHA-restorer lines and Oil and Nonoil lines for the 32 metabolites identified across at least 200 cultivated varieties

**Figure S5** Comparisons of heritability and additive relative effect sizes for sunflower ecophysiological and specialized metabolite traits.

**Table S1** Metabolites detected in a majority of the ‘core 12’ lines with HPLC-MS. HCA = hydroxycinnamic acid; SQTL = sesquiterpene lactone

| RT (min) | m/z ratio | Prevalence | Putative Identity | Class |
| --- | --- | --- | --- | --- |
| 4.1 | 353.1 | 12/12 | Scopolin | Coumarin |
|  |  |  | 3-O-caffeoylquinic acid (chlorogenic acid) | HCA |
|  |  |  | 4-O-caffeoylquinic acid (crypto-chlorogenic acid) | HCA |
|  |  |  | 5-O-caffeoylquinic acid (neo-chlorogenic acid) | HCA |
| 4.4 | 433.2 | 7/12 | Coreopsin | Flavonoid |
| 4.5 | 583.3 | 7/12 | Flavoxanthin | Xanthophylls |
| 4.7 | 427.2 | 6/12 | Farnesol B (SQTL) | SQTL |
| 4.8 | 357.1 | 11/12 | Atripliciolide angelate | SQTL |
|  |  |  | Eupasserin acrylate 2 hydroxyethyl | SQTL |
|  |  |  | Gardenin B | Flavonoid |
| 4.9 | 613.2 | 11/12 | Citrusin B | Phenlypropanoid |
| 5.0 | 549.2 | 8/12 | Euchretin C | Flavonoid |
| 5.1 | 523.2 | 9/12 | Rhodamine | Xanthene |
| 5.2 | 521.2 | 10/12 | Kankoside A | Terpene glycoside |
| 5.4 | 471.4 | 6/12 | α-amyrin | Triterpene |
|  | 473.3 |  | β-amyrin  Schidigeragenin | Triterpene  Triterpene |
| 6.5 | 341.1 | 12/12 | Calophyllin | Xanthones |
| 6.7 | 519.2 | 12/12 | Physalolactone | Sterol |
| 6.9 | 411.2 | 7/12 | Argophyllin C (formide adduct) | SQTL |
|  |  |  | Orizabin | SQTL |
|  |  |  | Stigmasterol | Phytosterol |
| 7.1 | 515.2 | 12/12 | 3,4-di-O-caffeoylquinic acid | HCA |
|  |  |  | 3,5-di-O-caffeoylquinic acid | HCA |
|  |  |  | 4,5-di-O-caffeoylquinic acid | HCA |
| 7.3 | 399.1 | 8/12 | Angeloylgrandifloric acid | Diterpene |
|  |  |  | Scopolin (formide adduct) | Coumarin |
|  |  |  | 3-O-caffeoylquinic acid (formide adduct) | HCA |
|  |  |  | 4-O-caffeoylquinic acid (formide adduct) | HCA |
|  |  |  | 5-O-caffeoylquinic acid (formide adduct) | HCA |
| 7.7 | 383.2  179.1 | 12/12  12/12 | 5-O-p-coumaroylquinic acid (formide adduct)  Caffeic acid | HCA  HCA |
| 7.9 | 273.0 | 7/12 | Thiarubrine B | Polyacetelyenes |
| 8.0 | 393.1 | 6/12 | Eupasserin trihydroxyangelate | SQTL |
|  |  |  | Tifruticin, 1-hyrdoxy | SQTL |
|  |  |  | Cumambranolide 2 hydroxyethylacrylate | SQTL |
|  |  |  | Cumambranolide angelate | SQTL |
| 8.2 | 505.3 | 8/12 | Euphornin | Diterpene |
| 8.6 | 421.2 | 11/12 | Atripliciolide tiglate 3-hydroxy | SQTL |
|  |  |  | Tifruticin, 15-hydroxy-3-dehydrodeoxy | SQTL |
|  |  |  | Niveusin A, 1,2-anhydrido | SQTL |
| 8.9 | 329.1 | 10/12 | Desmethoxysudachitin | Flavonoid |
|  |  |  | Apigenin, 6,4'-dimethoxy | Flavonoid |
|  |  |  | Jaceosidin (6-methoxy-luteolin 3'-methyl ether) | Flavonoid |
| 9.1 | 183.1 | 12/12 | Salicylic acid (formide adduct) | salicylate |
| 9.2 | 407.1 | 9/12 | Eupaserrin angelate, 14-hydroxy | SQTL |
|  |  |  | Budlein A isobutyrate | SQTL |
|  |  |  | Mollisorin B | SQTL |
|  |  |  | Helieudesmanolide B | SQTL |
|  |  |  | Heliangin | SQTL |
|  |  |  | Leptocarpin | SQTL |
|  |  |  | Niveusin A, 1-methoxy-4,5-dihydro (formide adduct) | SQTL |

**Figure S1** Boxplots of leaf economics spectrum principal components values among HA-maintainer and RHA-restorer lines for A) PC1 and B) PC2 and among Nonoil and Oil for C) PC1 and D) PC2

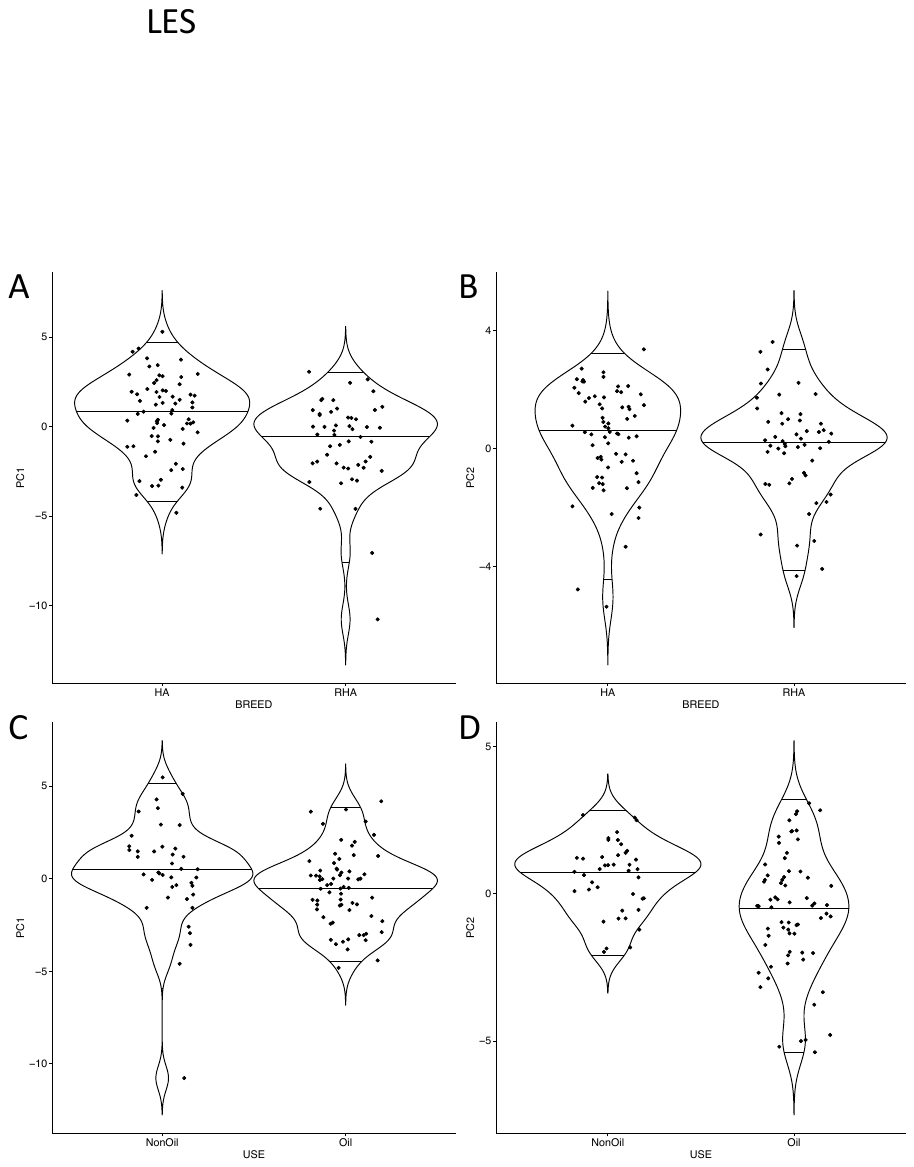

**Figure S2.** Principal component analysis of the relative ratios of specialized metabolites among 288 cultivated *Helianthus annuus* varieties. A) Individual loadings of cultivated *Helianthus annuus* varieties along PC 1 and 2 of the 106 metabolites identifies across varieties, including unique metabolites. B) Individual loadings of cultivated *Helianthus annuus* varieties along PC 1 and 2 of the 32 metabolites found across all varieties. Colors indicate breeding groups, and shapes indicate whether individuals are part of the ‘Core 12’ set of genotypes (circles) representing 50% of the genetic diversity of the SAM population or not (triangles). All breeding groups are from the USDA germplasm repositories except for INRA-HA and INRA-RHA. There are no differences between the loadings of Figure 2 and this figure.

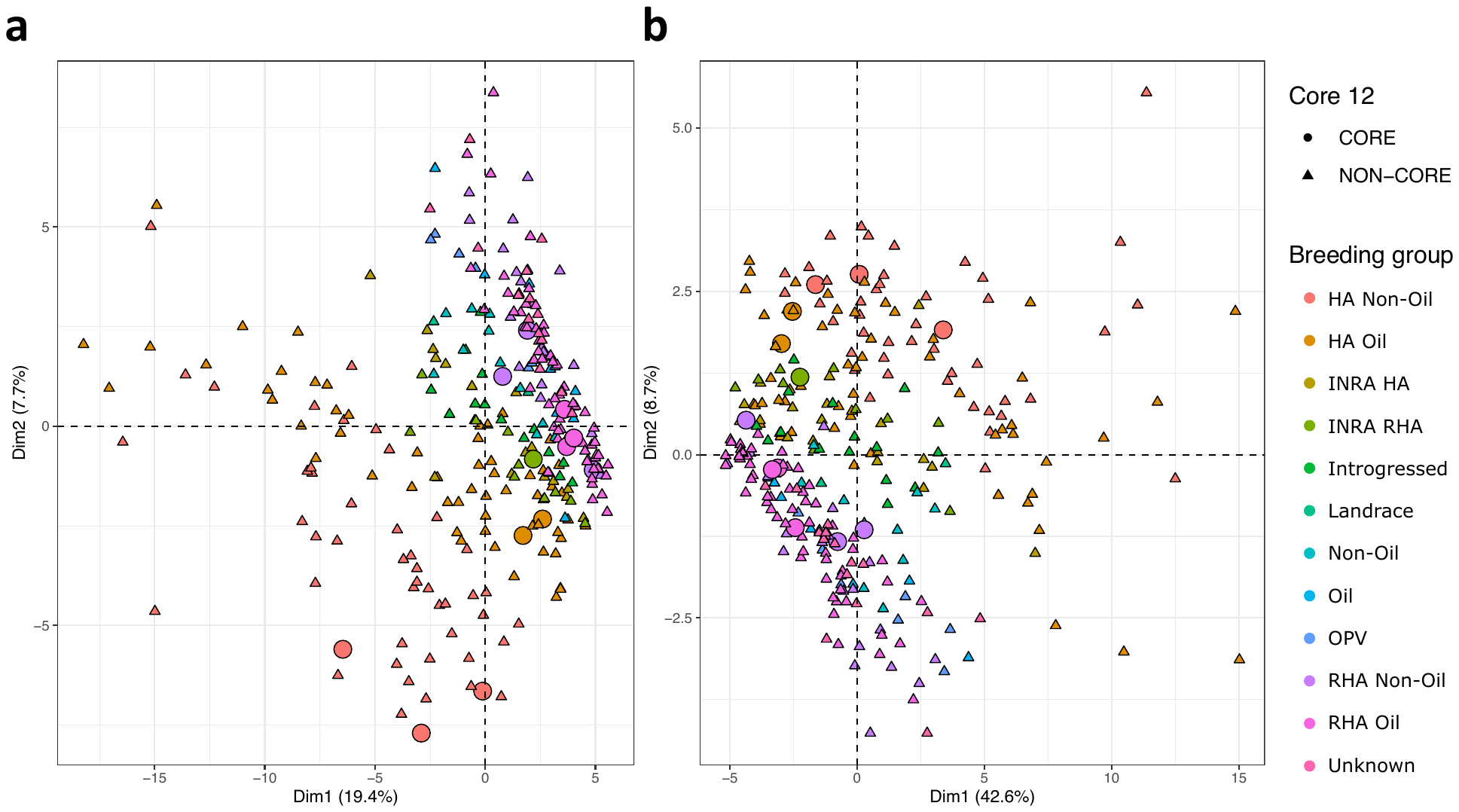

**Figure S3** Boxplots of specialized metabolite peak area PC values among HA-maintainer and RHA-restorer lines for A) PC1 and B) PC2 and among Oil and Nonoil lines for C) PC1 and D) PC2 of the 106 metabolites identified across genotypes, including unique metabolites.

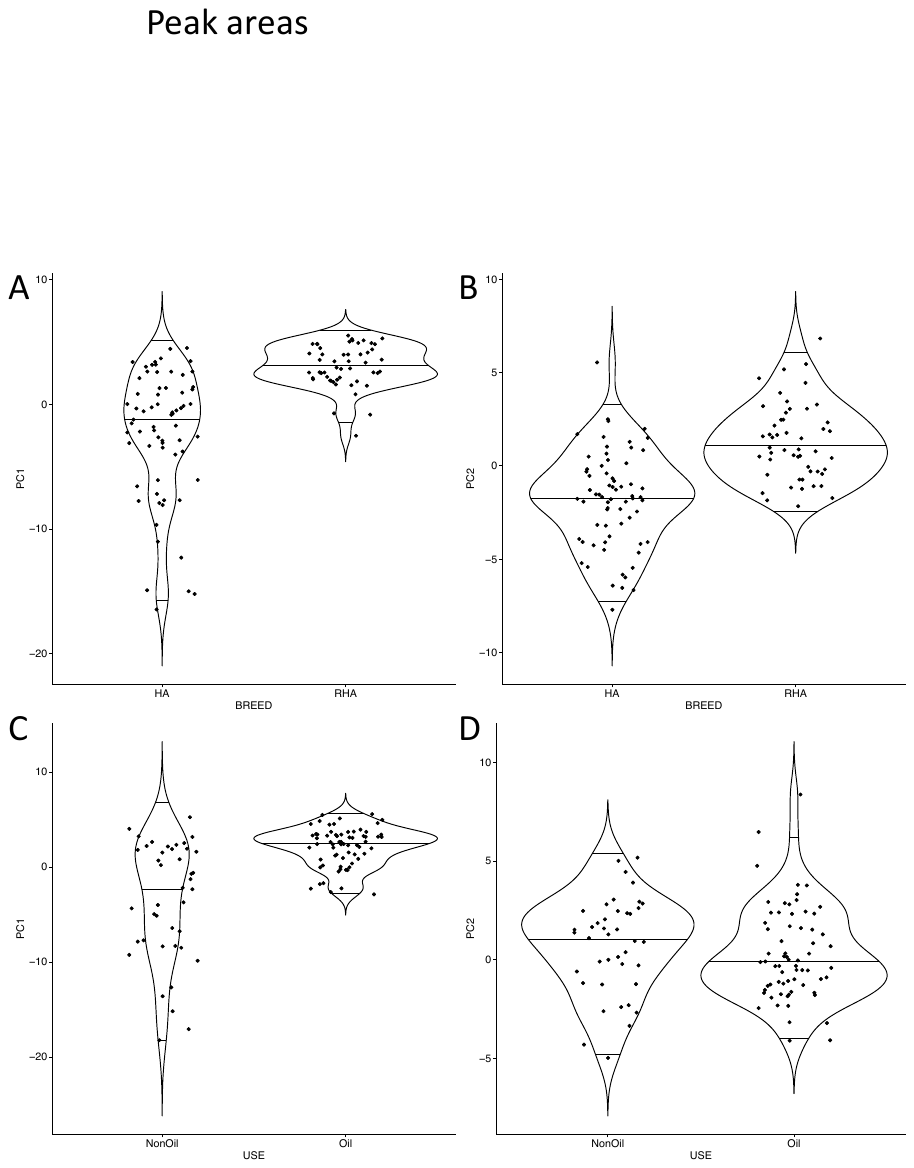

**Figure S4** Boxplots of specialized metabolite peak area PC values among HA-maintainer and RHA-restorer lines for A) PC1 and B) PC2 and among Oil and Nonoil lines for C) PC1 and D) PC2 of the 32 metabolites identified across at least 200 cultivated varieties

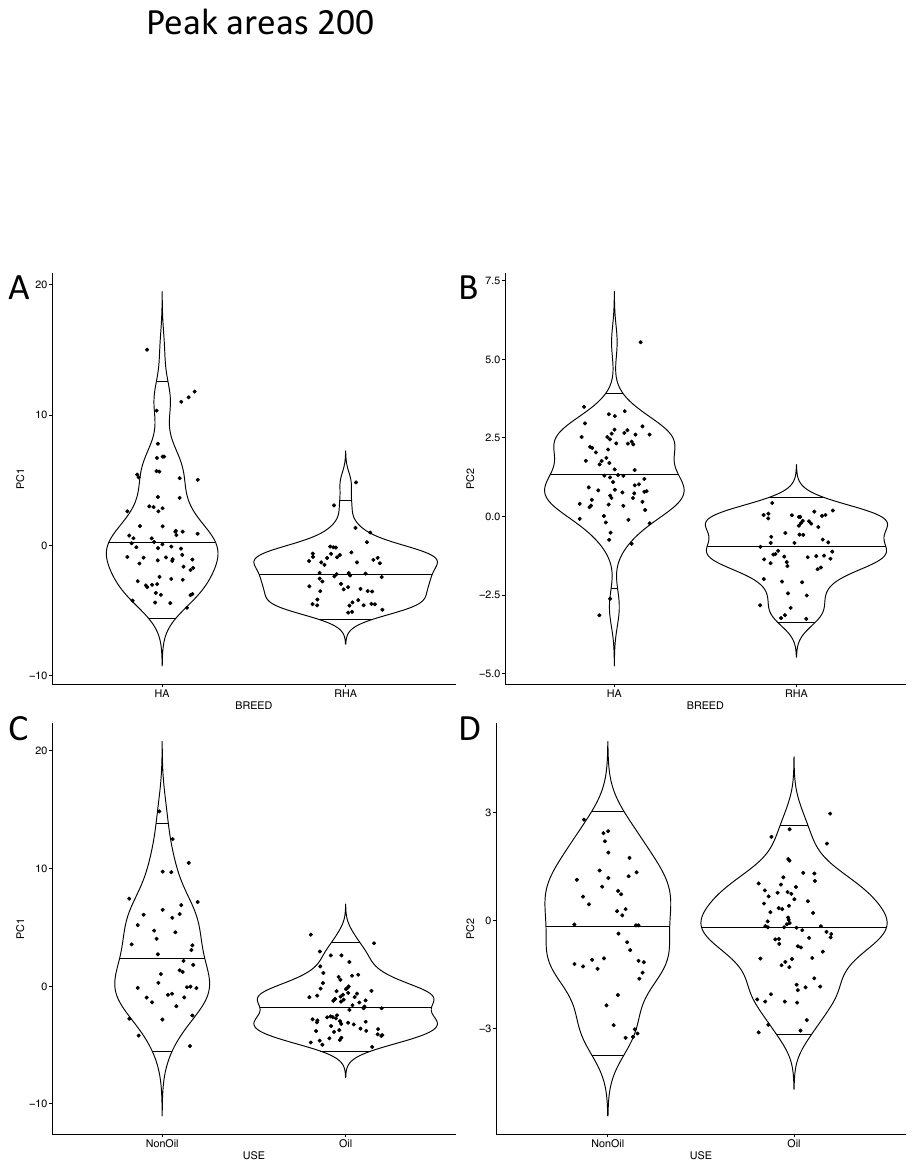

**Figure S5** Comparisons of heritability and additive relative effect sizes for sunflower ecophysiological and specialized metabolite traits. a). Histogram and density curves for all traits and principal components assessed. The dashed black line indicates the average heritability value among all traits and PCs. Red—principal components derived from specialized metabolites, Green—principal components derived from ecophysiological traits, Blue—specialized metabolites, and Purple—ecophysiological traits. b). Additive relative effect sizes for traits with significantly associated SNPs compared to their heritability values. Red—principal components derived from specialized metabolites, Green—specialized metabolite, Blue—Ecophysiological traits.

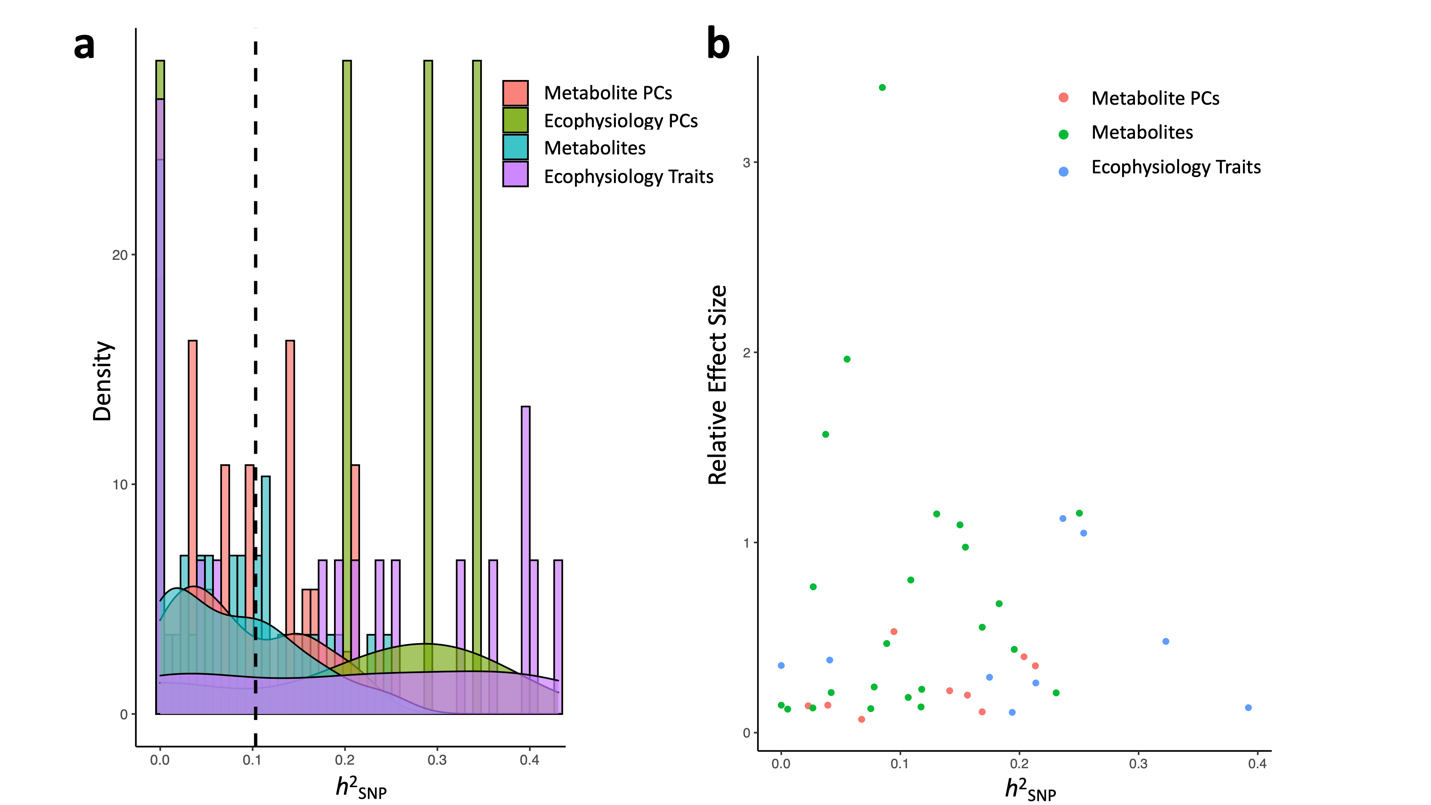
