## Supplementary figures and images for "Historic breeding practices contribute to germplasm divergence in leaf specialized metabolism and ecophysiology in cultivated sunflower (*Helianthus annuus*)"

### AspectRatio_BP.pdf

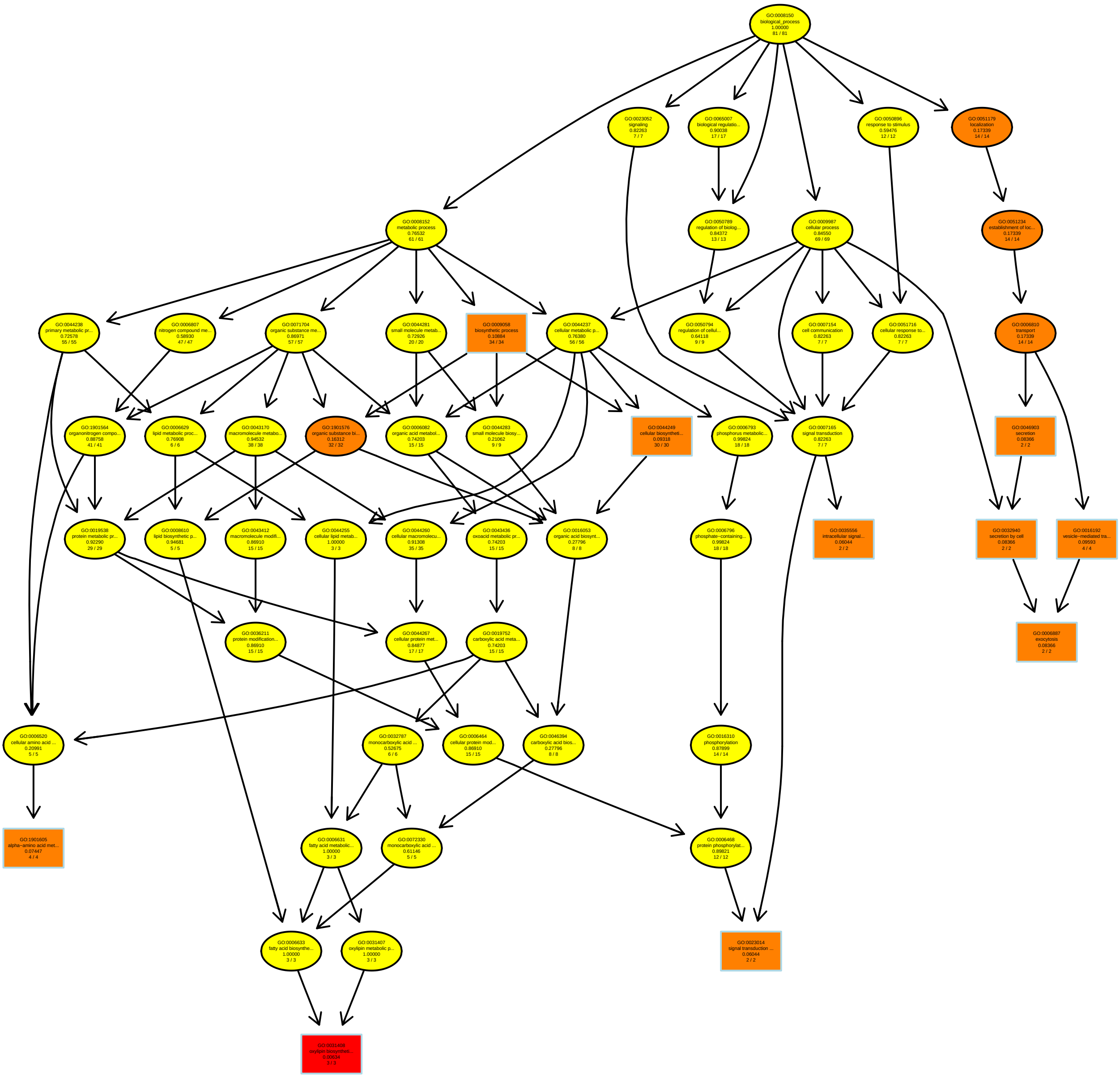

### ChlorophyllContentareabasis_MF.pdf

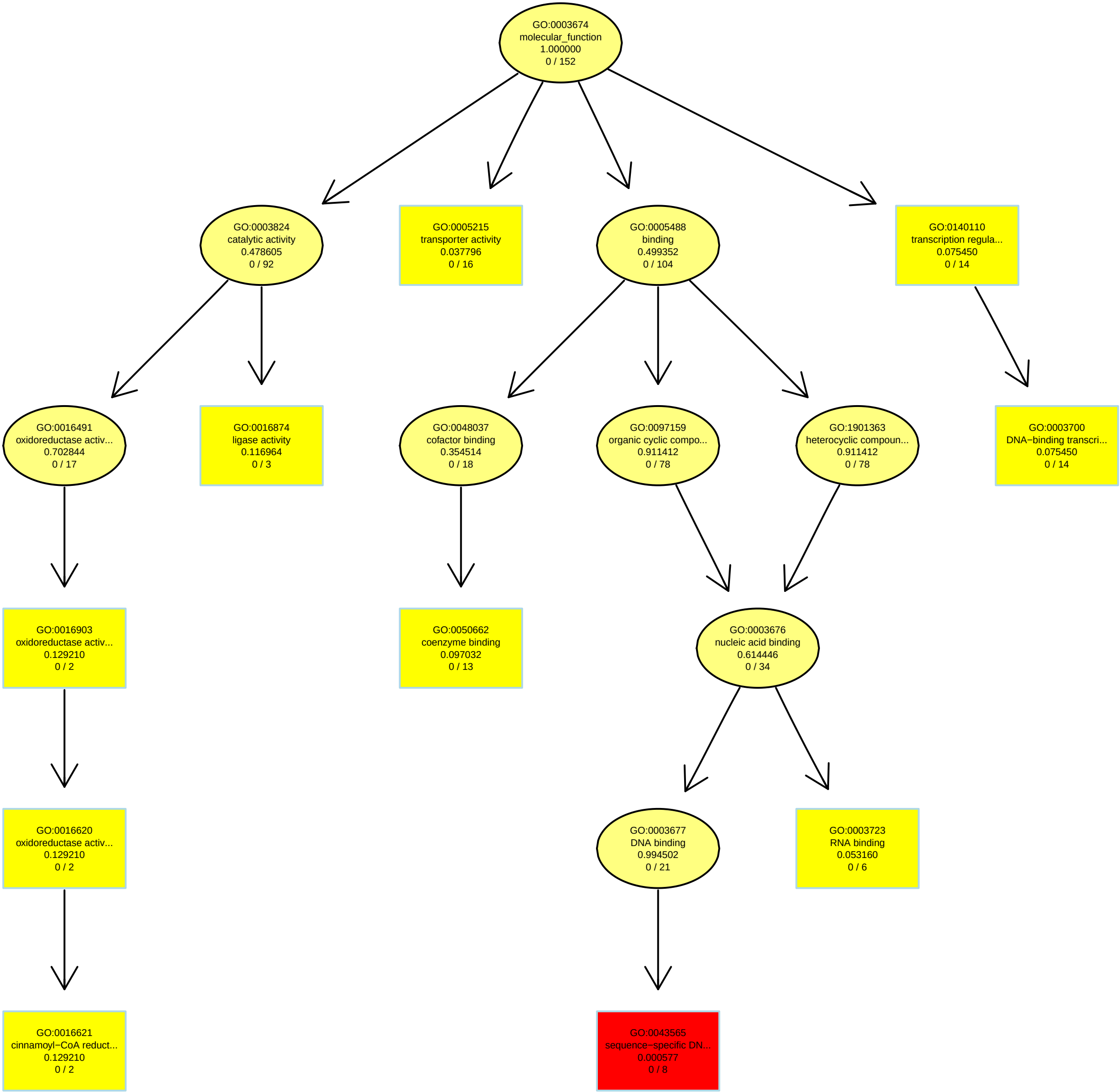

### Circularity_MF.pdf

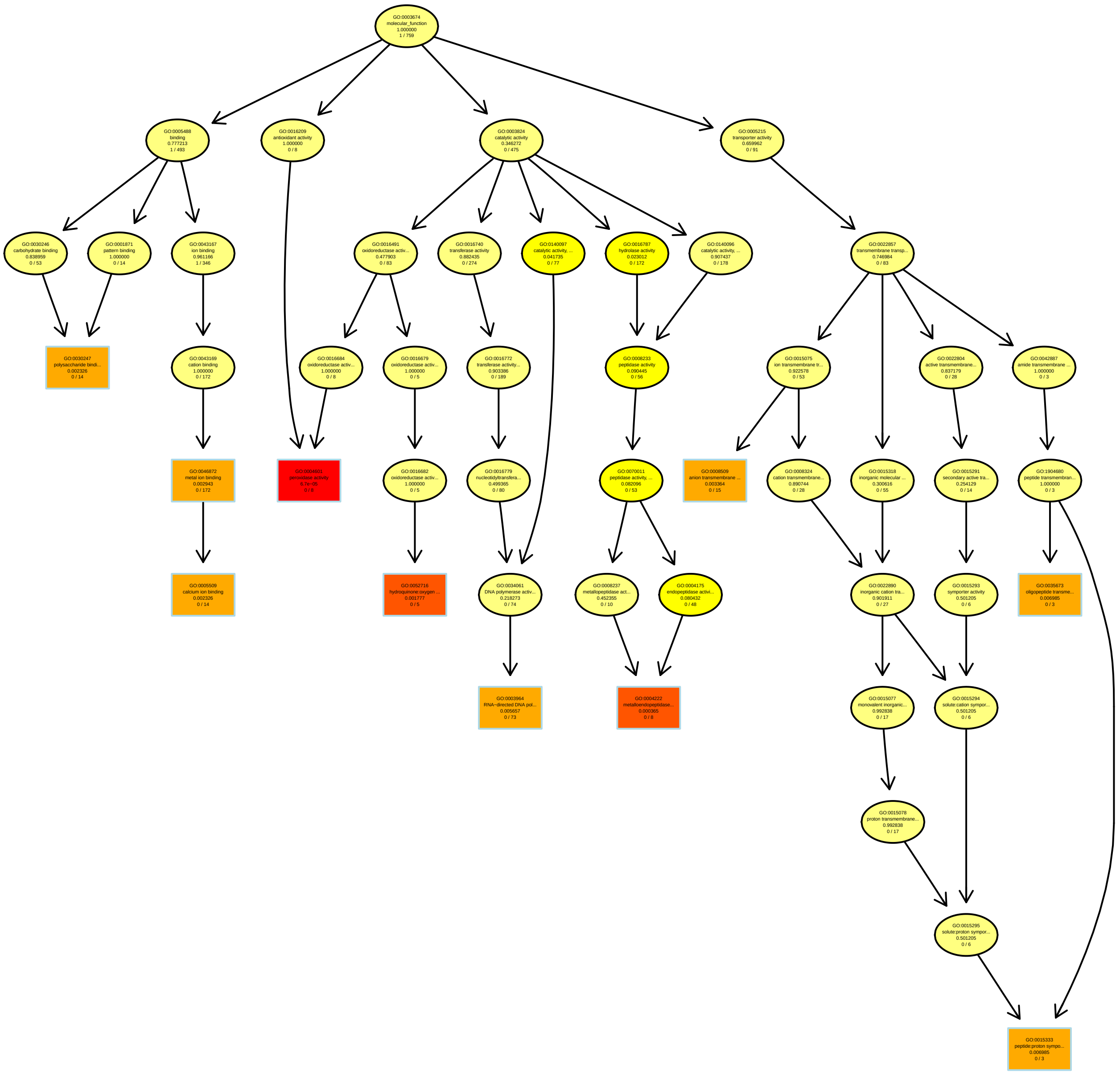

### DryMass_BP.pdf

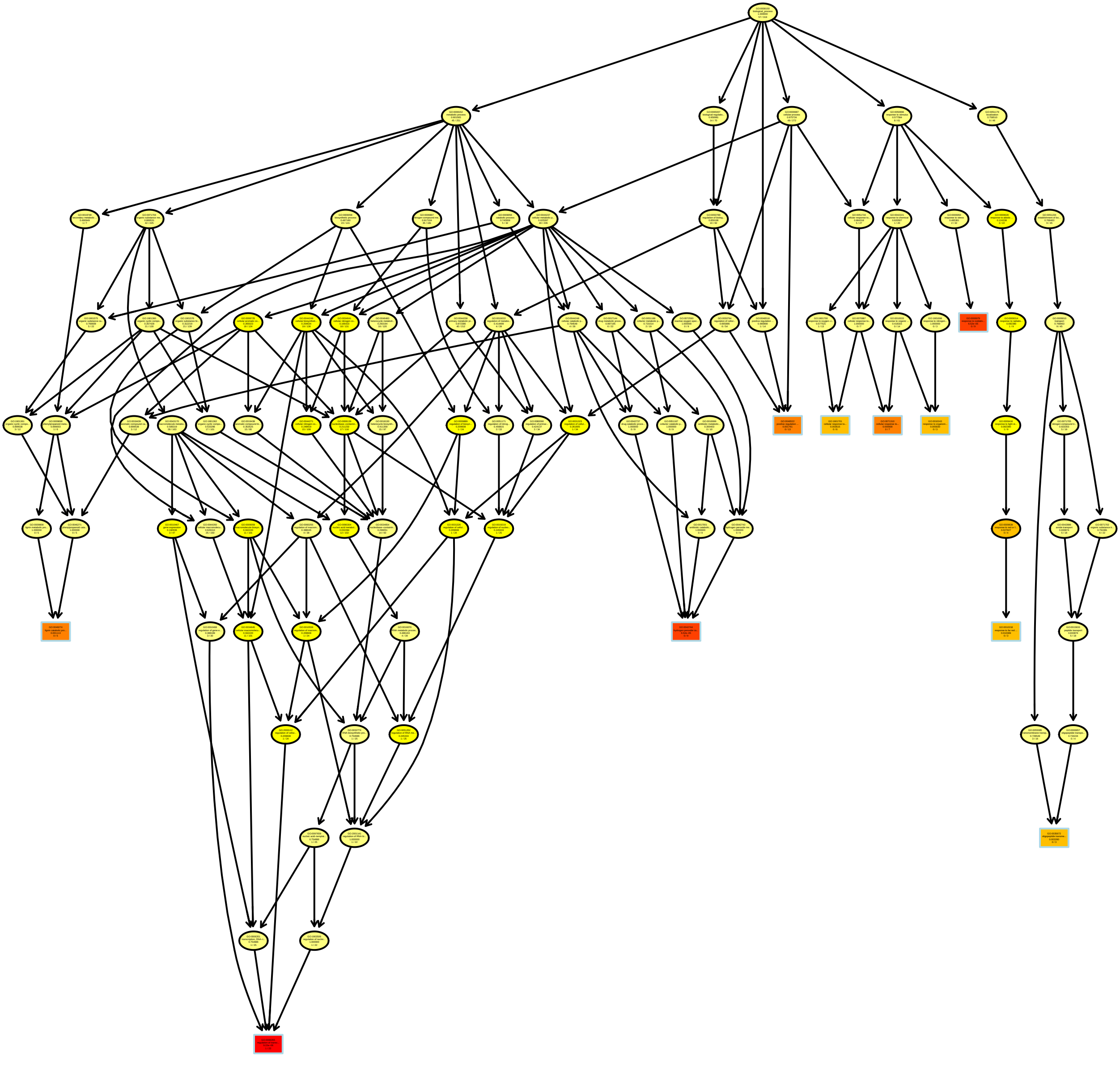

### FLAVPHENPC1_MF.pdf

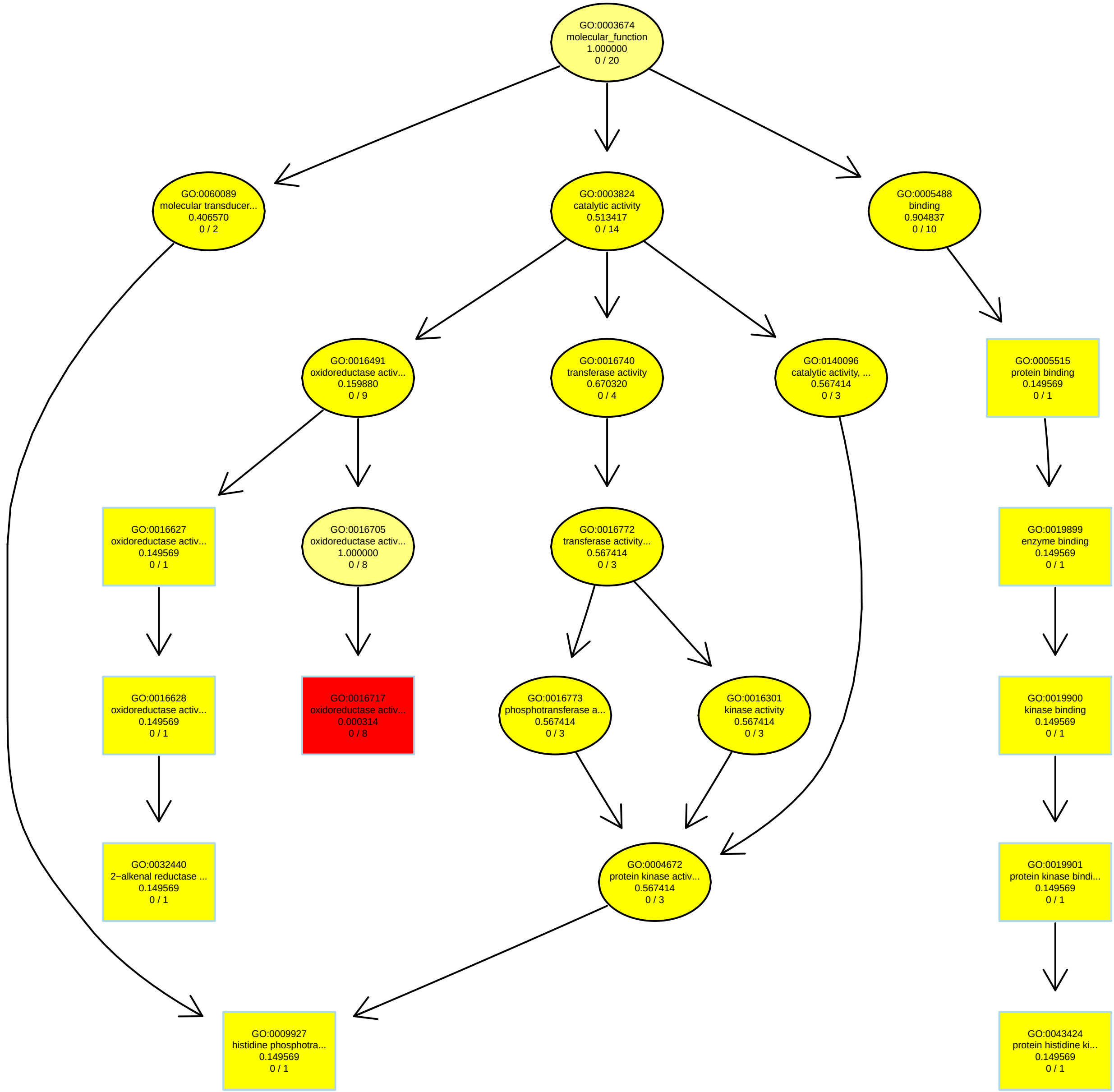

### FLAVPHENPC2_BP.pdf

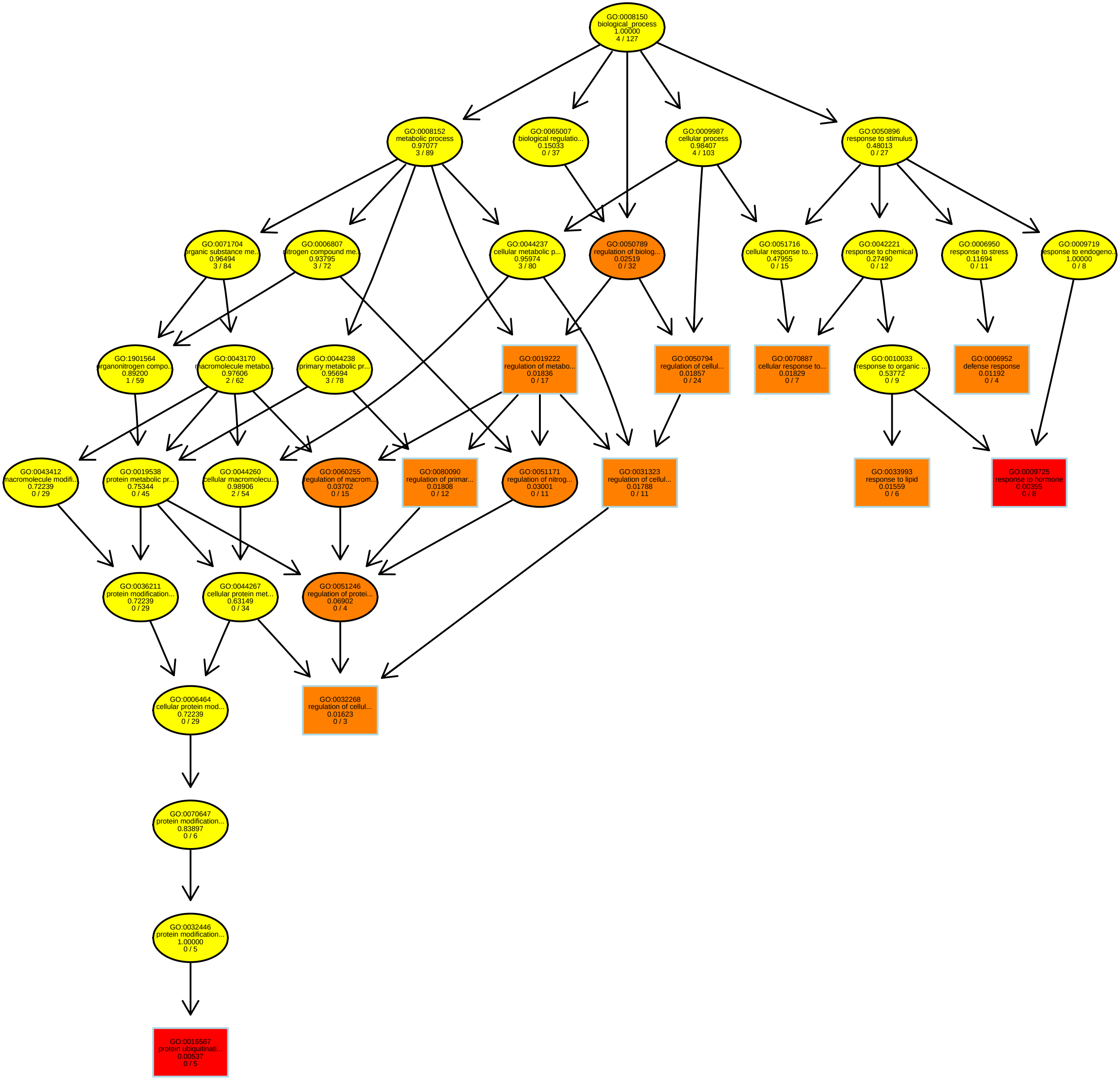

### FreshMass_BP.pdf

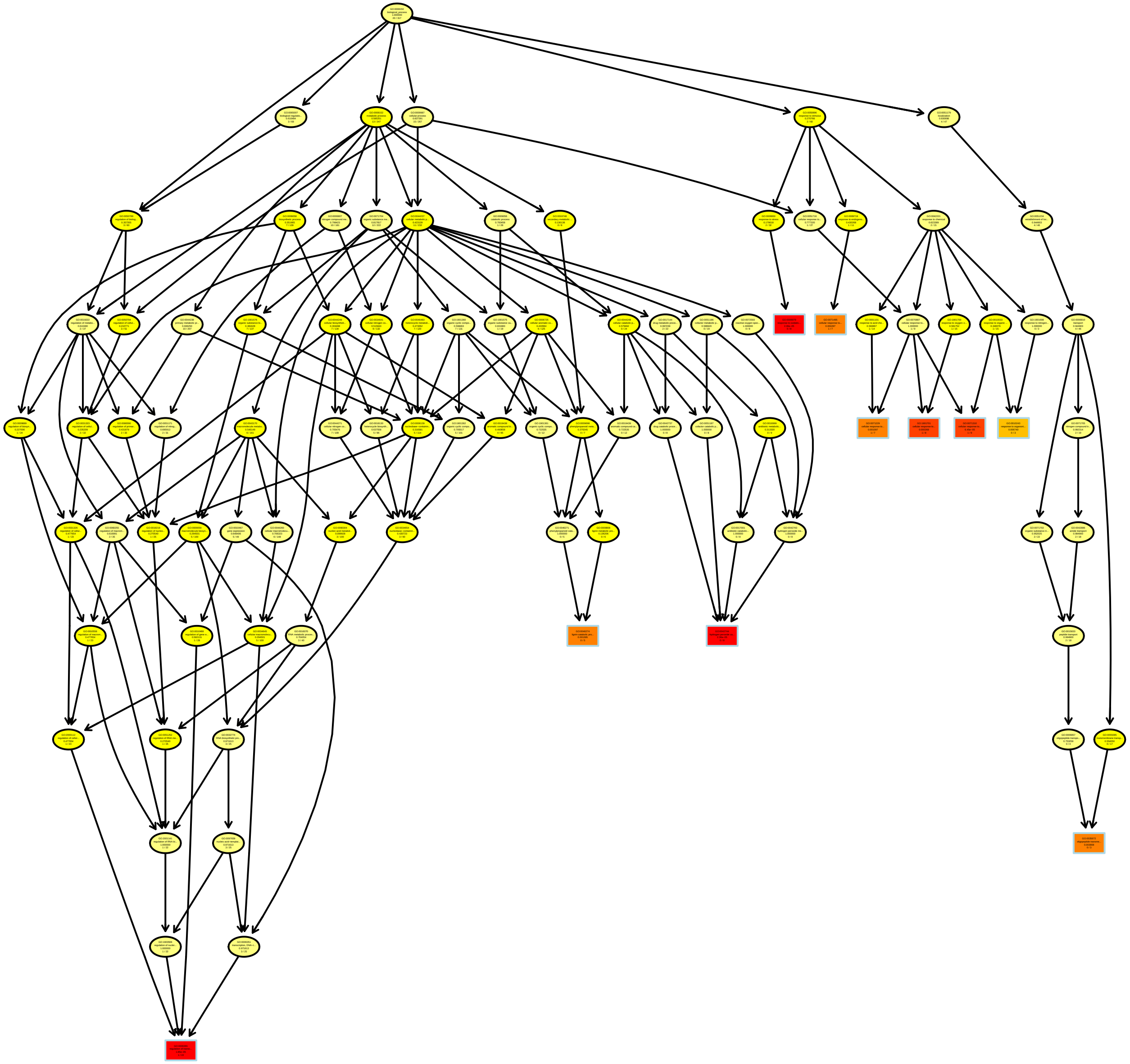

### Global__GOBP.pdf

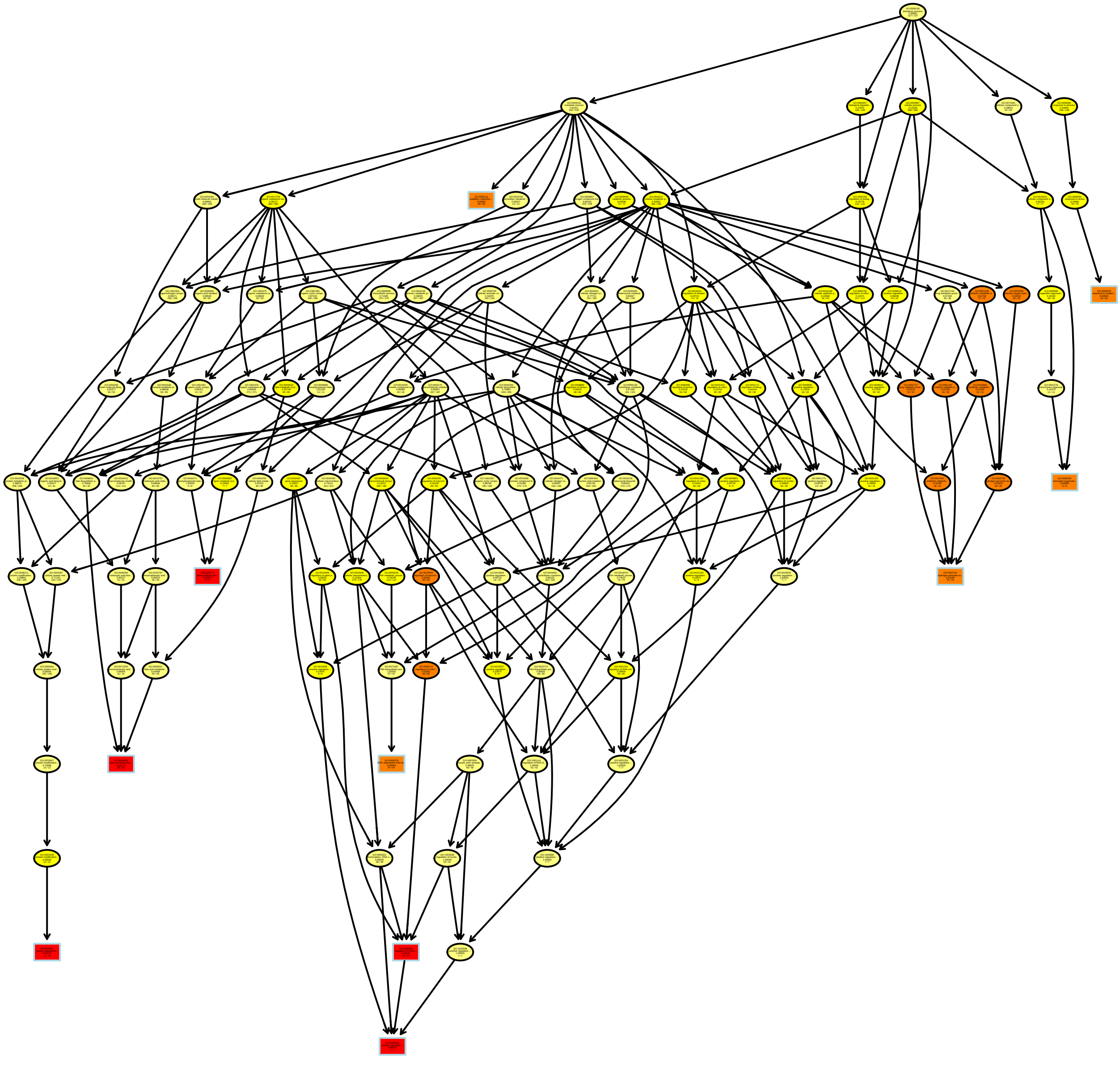

### LaminaDensity_MF.pdf

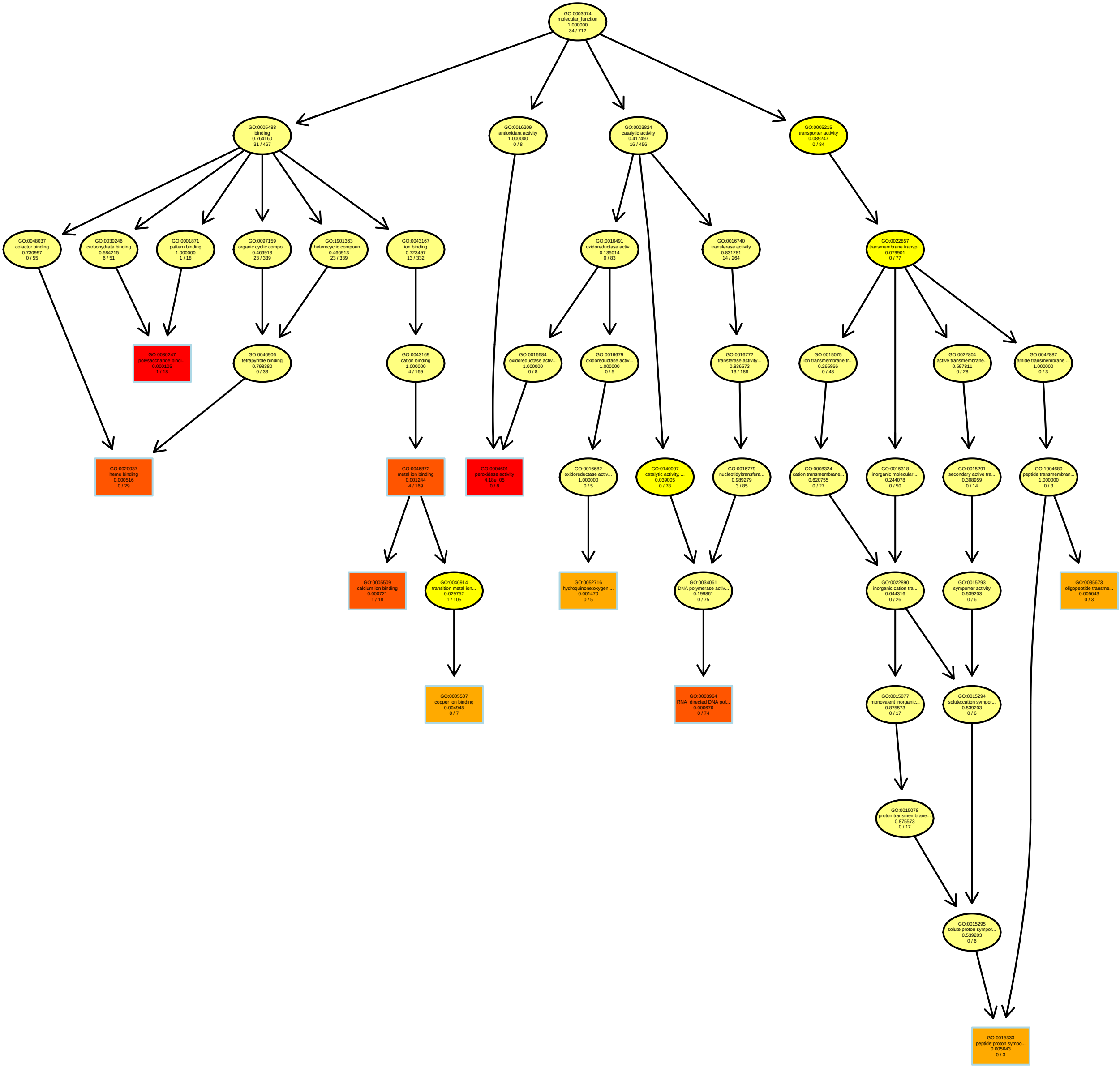

### LaminaThickness_MF.pdf

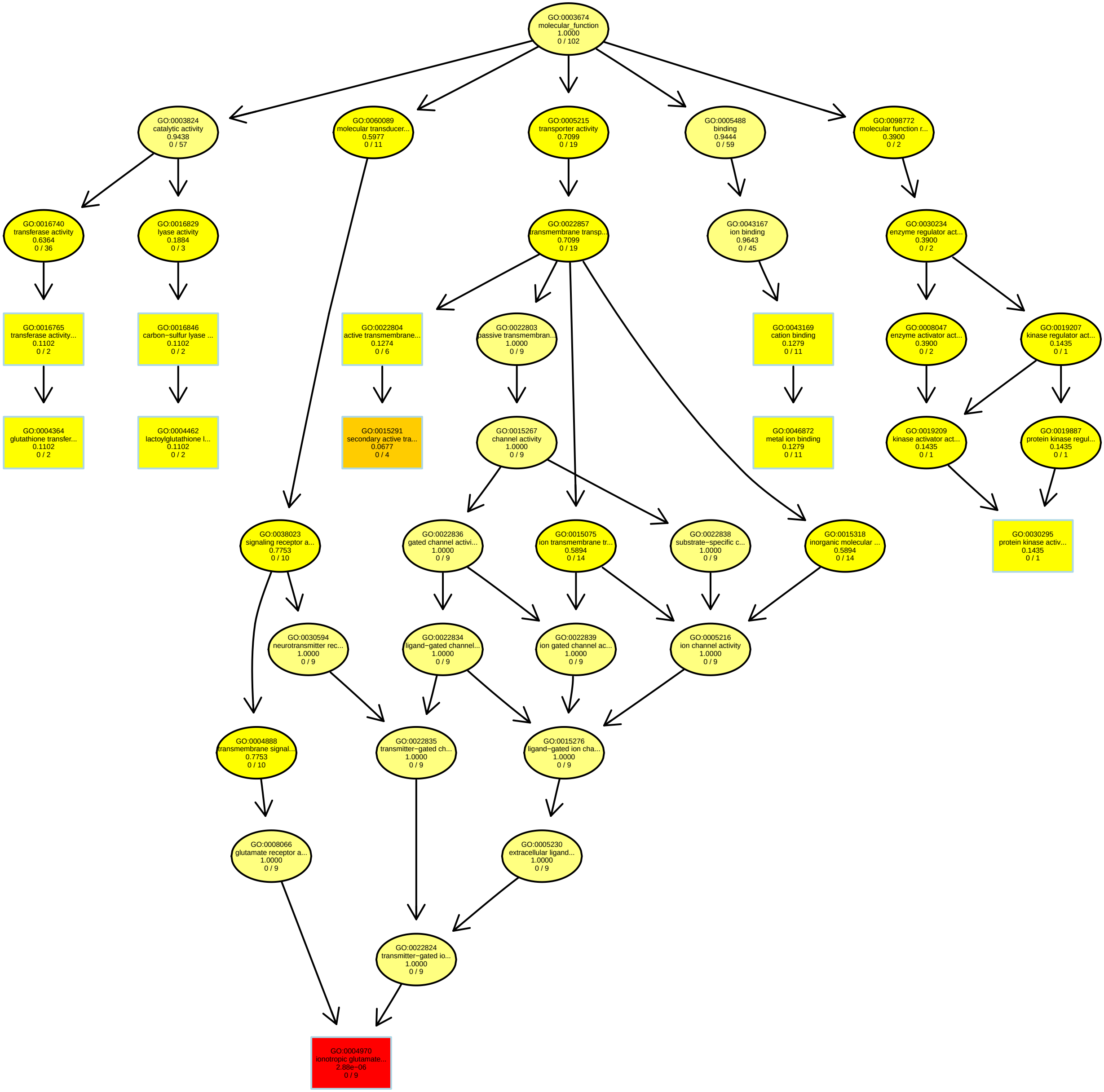

### LaminaToughness_BP.pdf

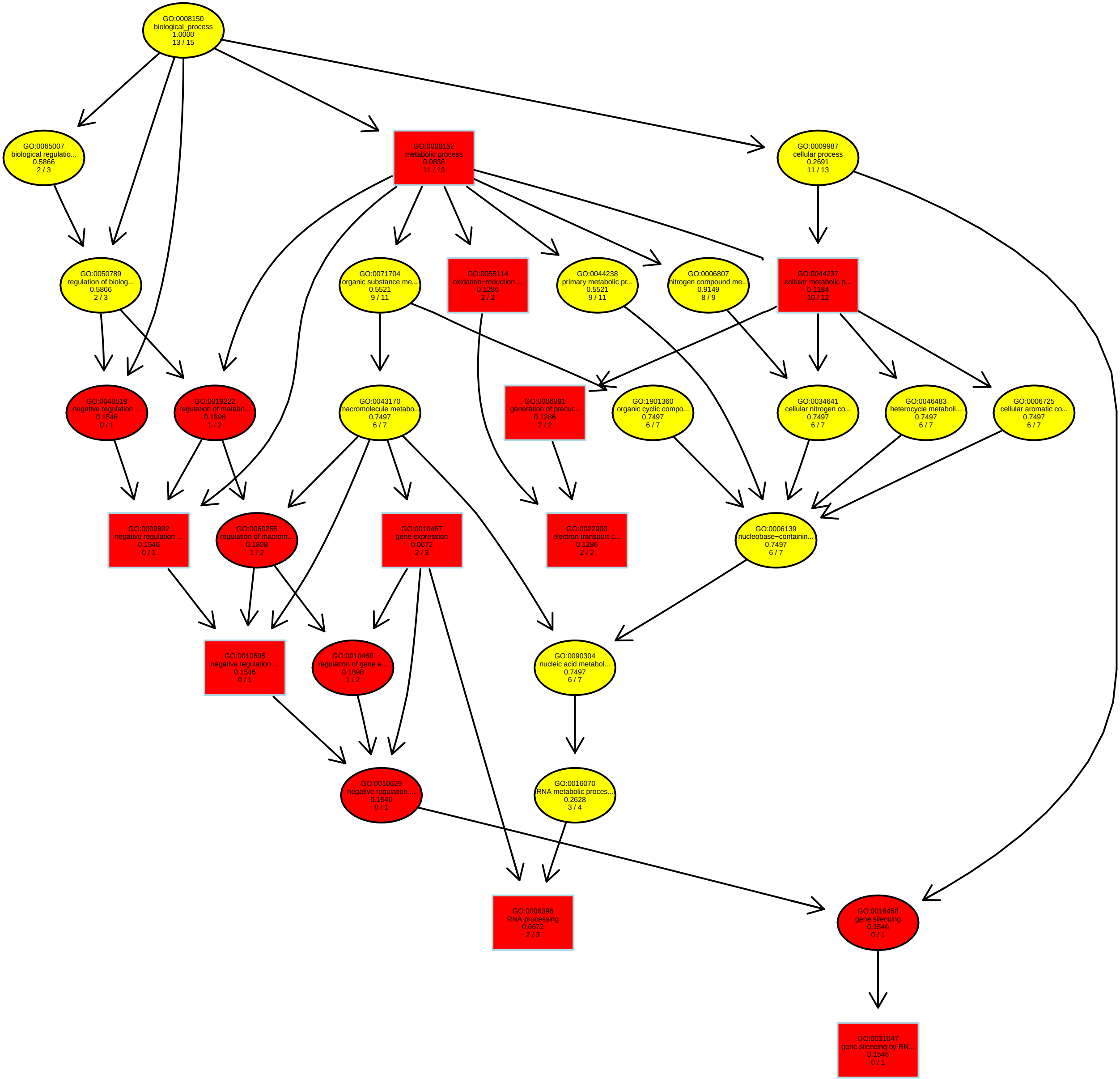

### LeafArea_MF.pdf

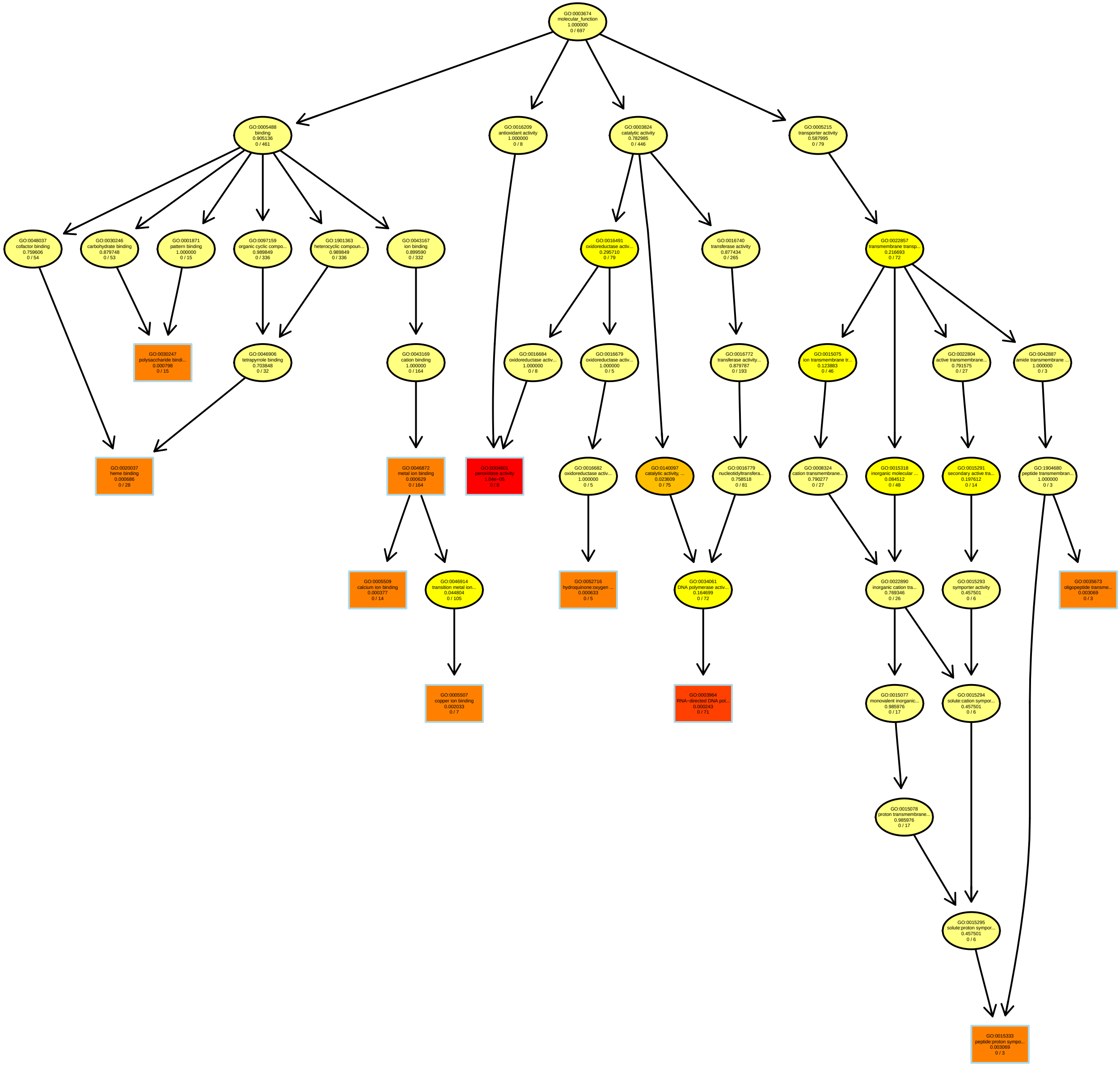

### LESPC2_BP.pdf

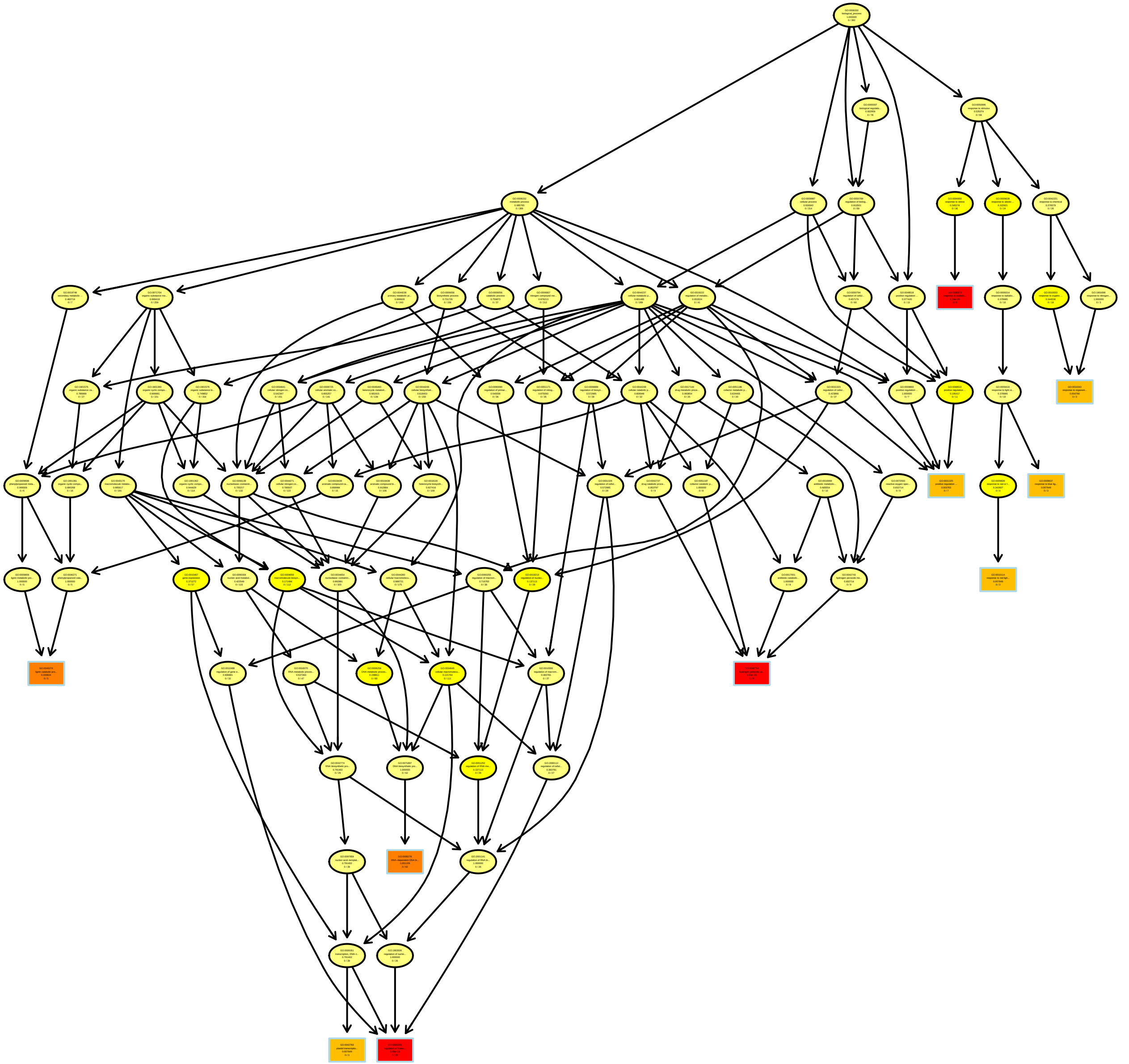

### LESPC3_MF.pdf

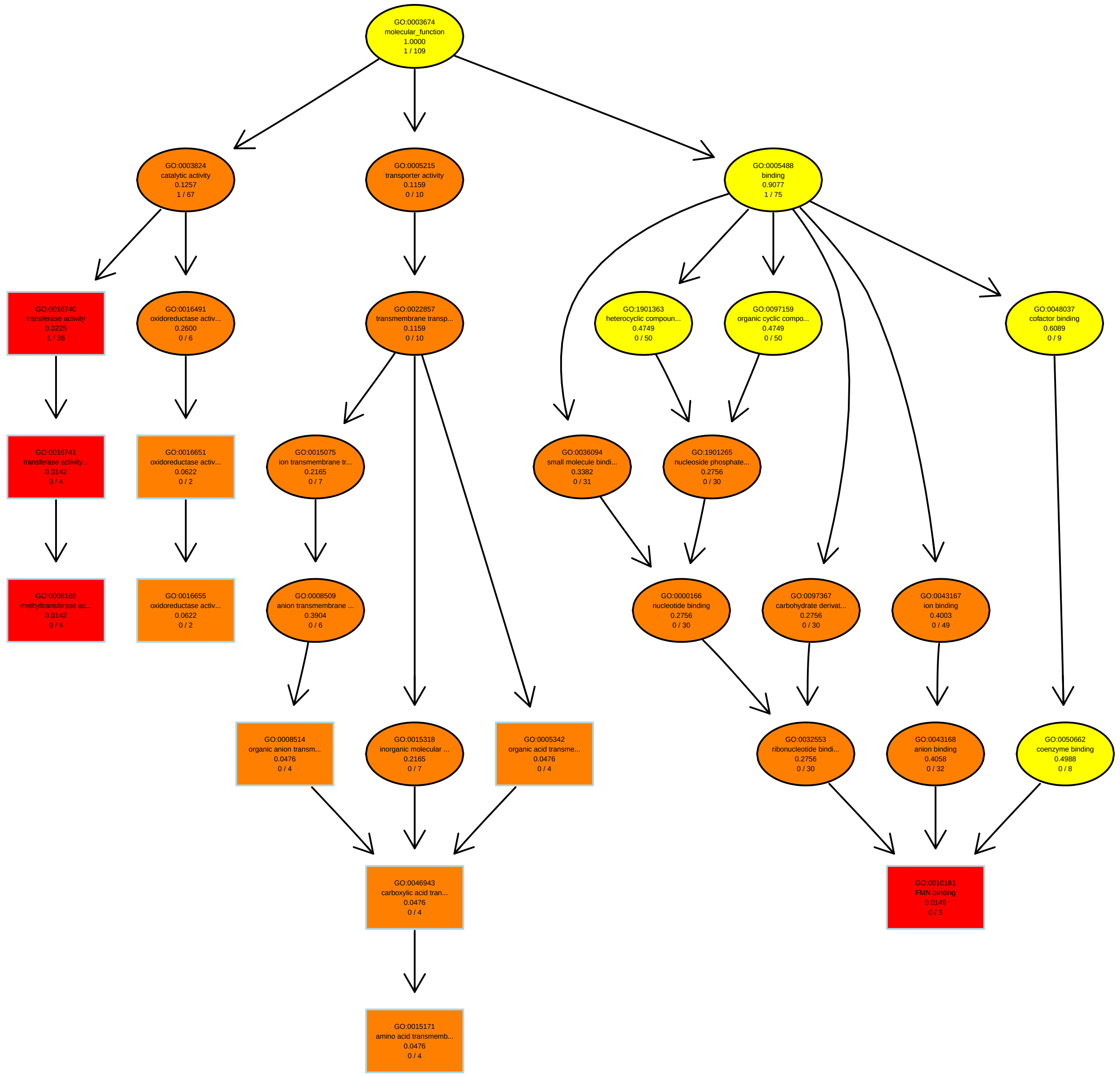

### LMA_BP.pdf

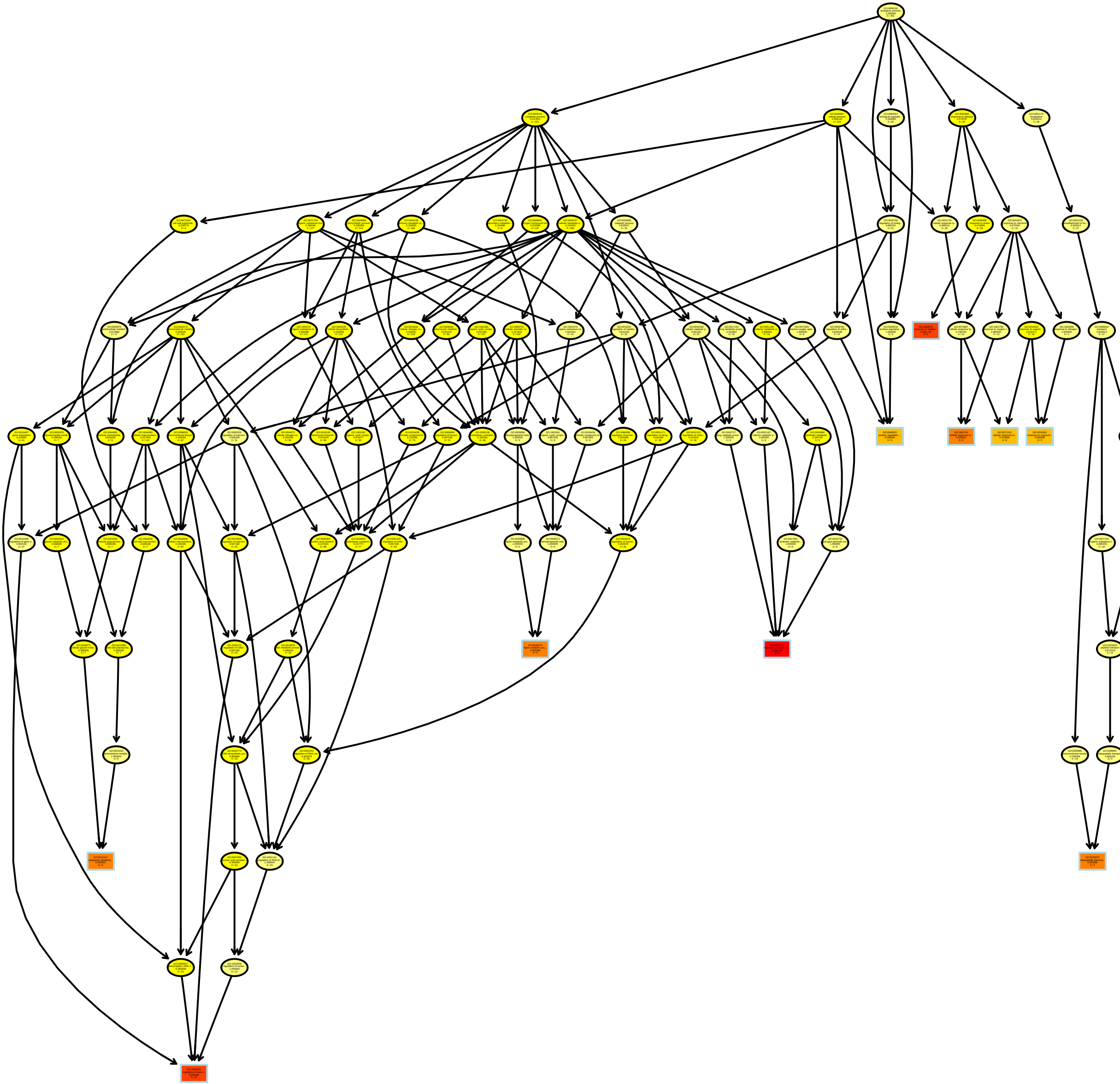

### PeakAreas200PC2_BP.pdf

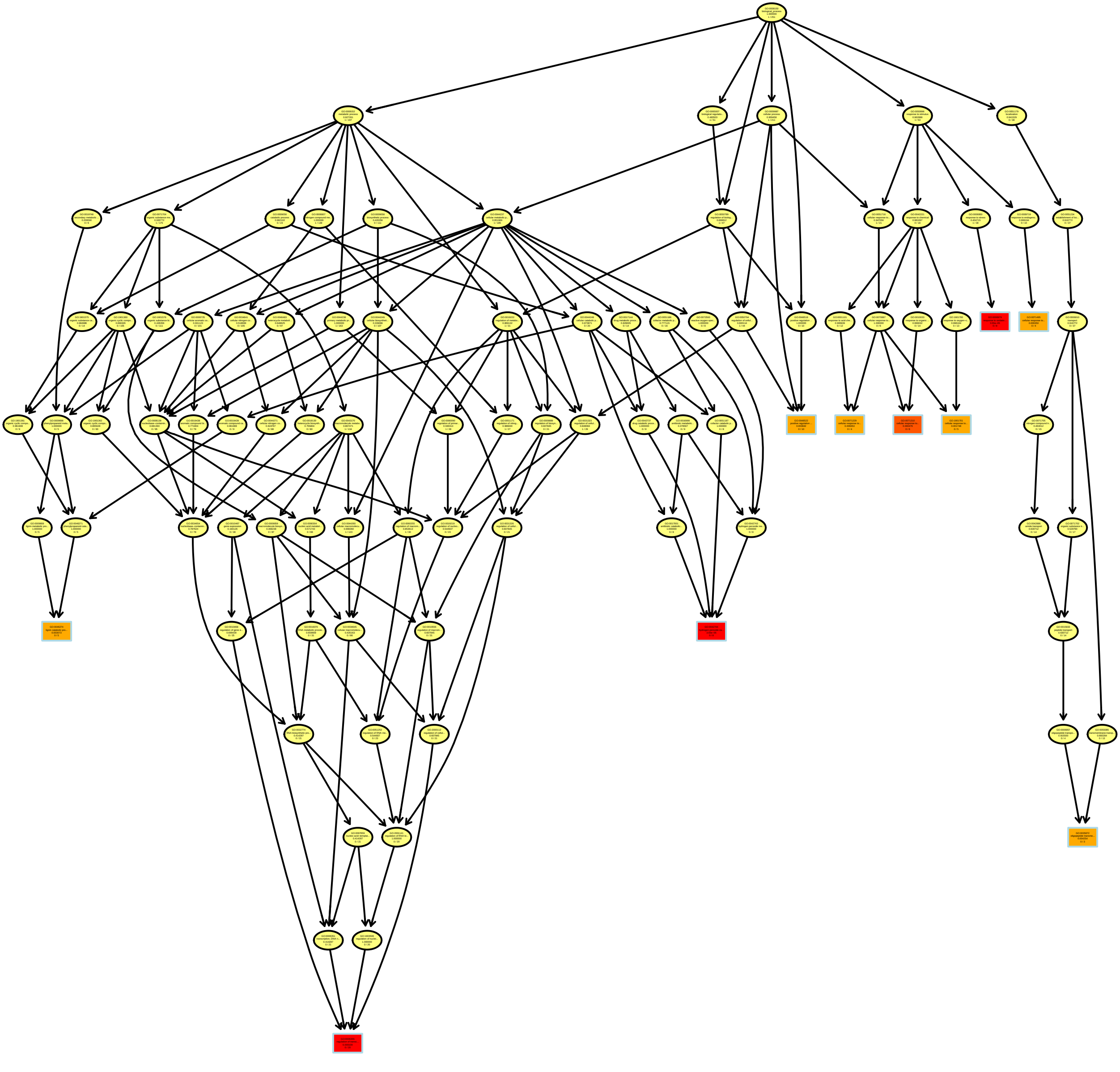

### PeakAreas200PC3_MF.pdf

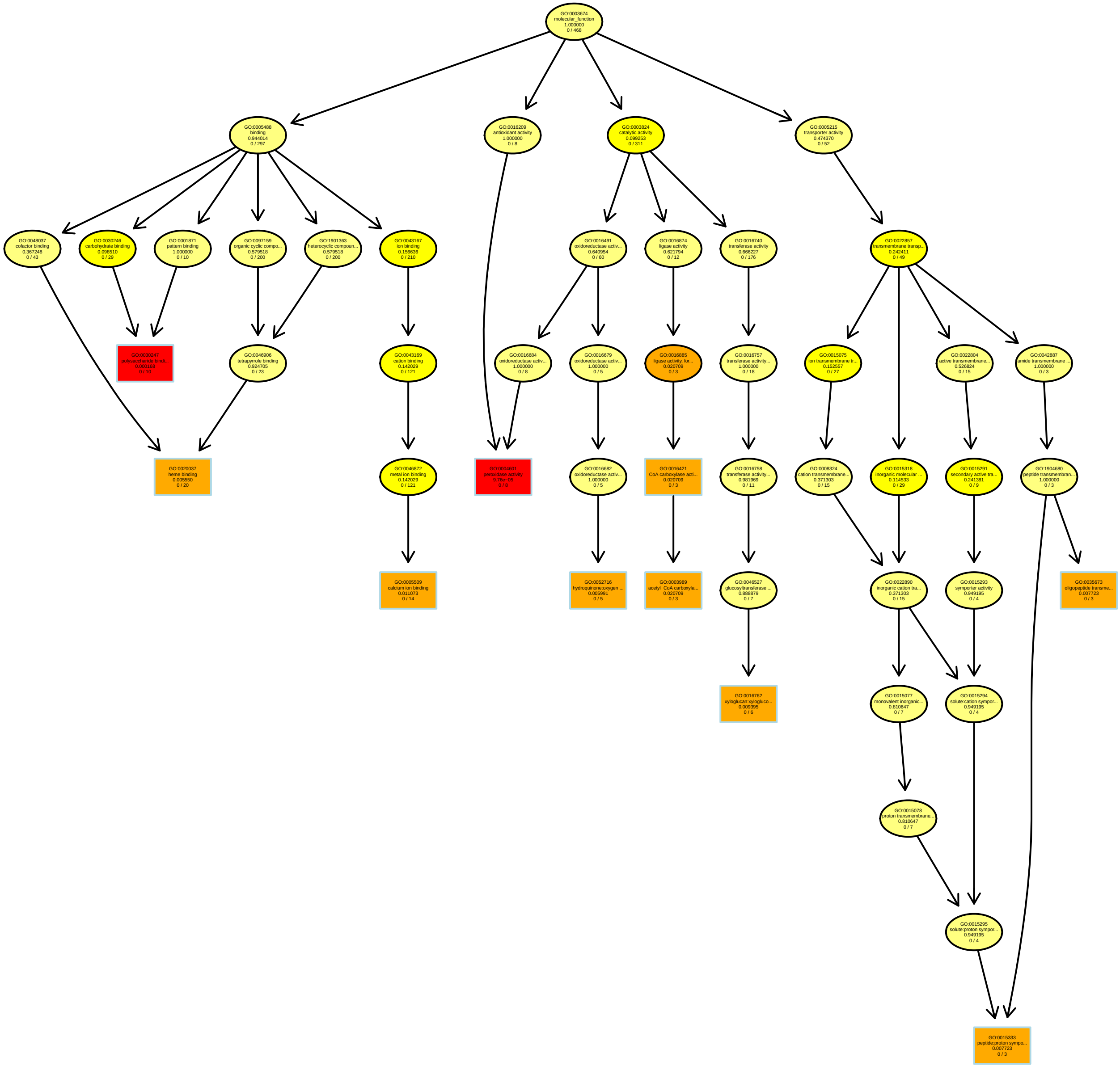

### PeakAreas200PC5_MF.pdf

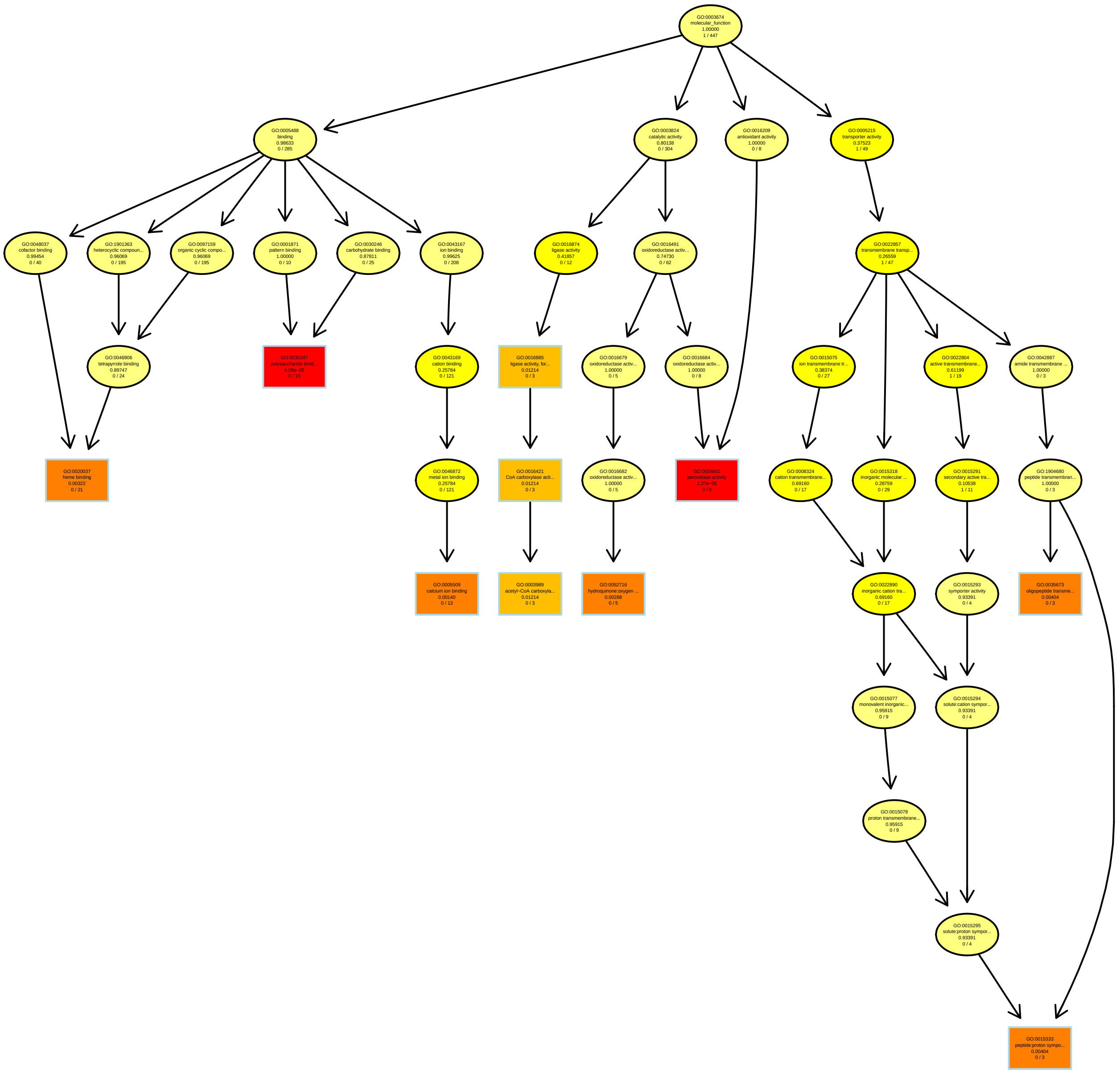

### PeakAreas200PC6_BP.pdf

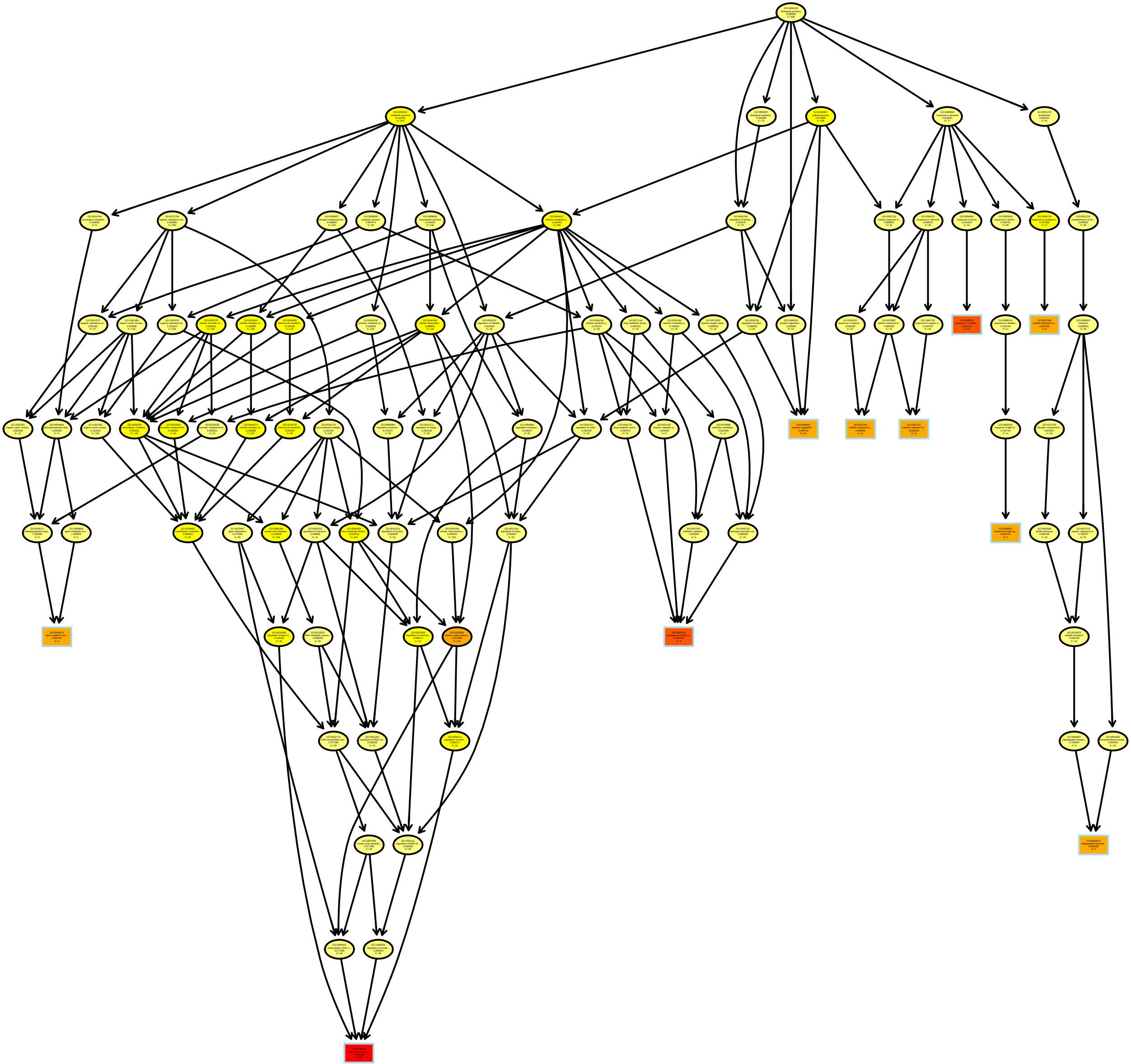

### PeakAreas200PC7_MF.pdf

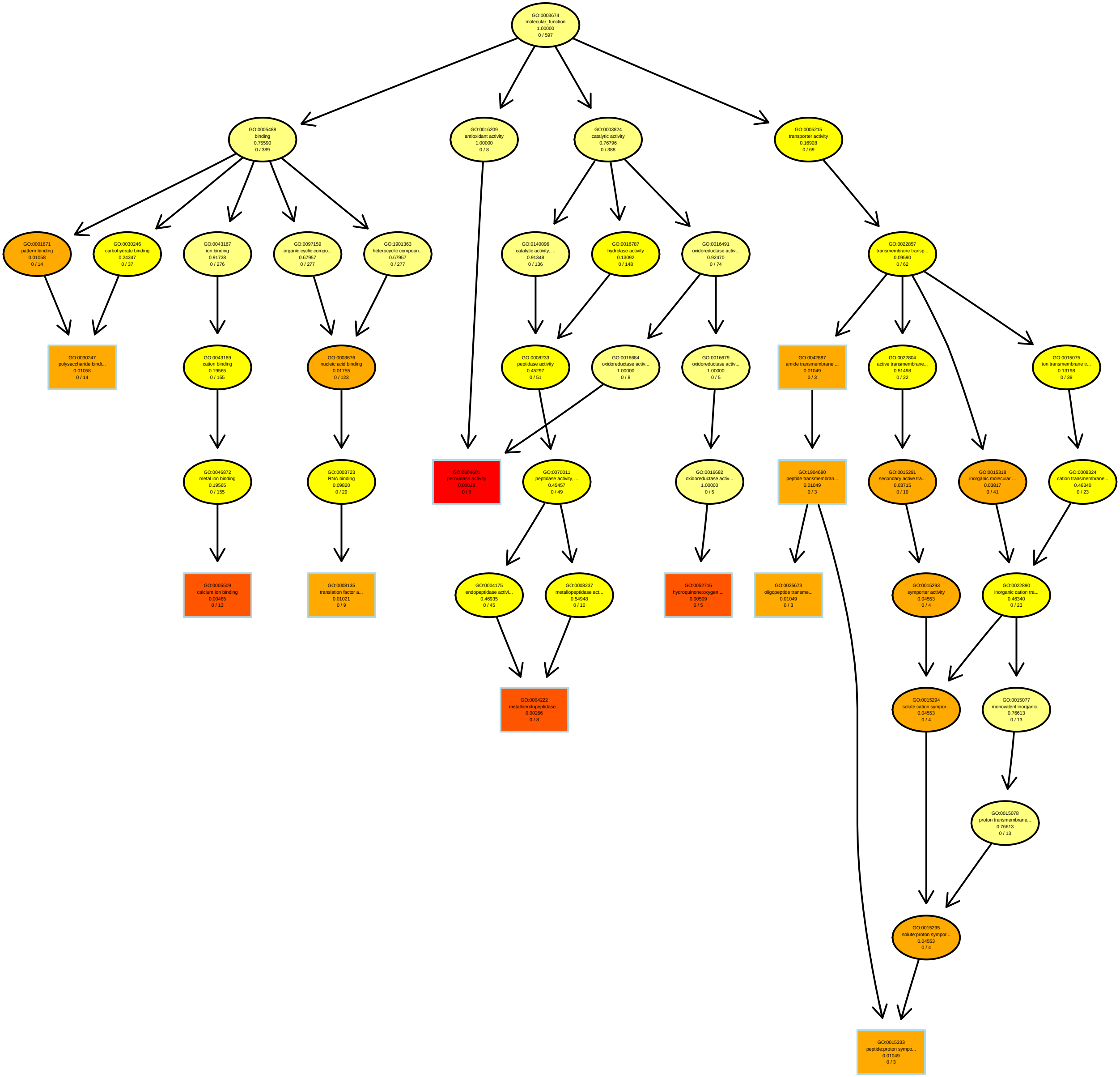

### PeakAreas200PC9_MF.pdf

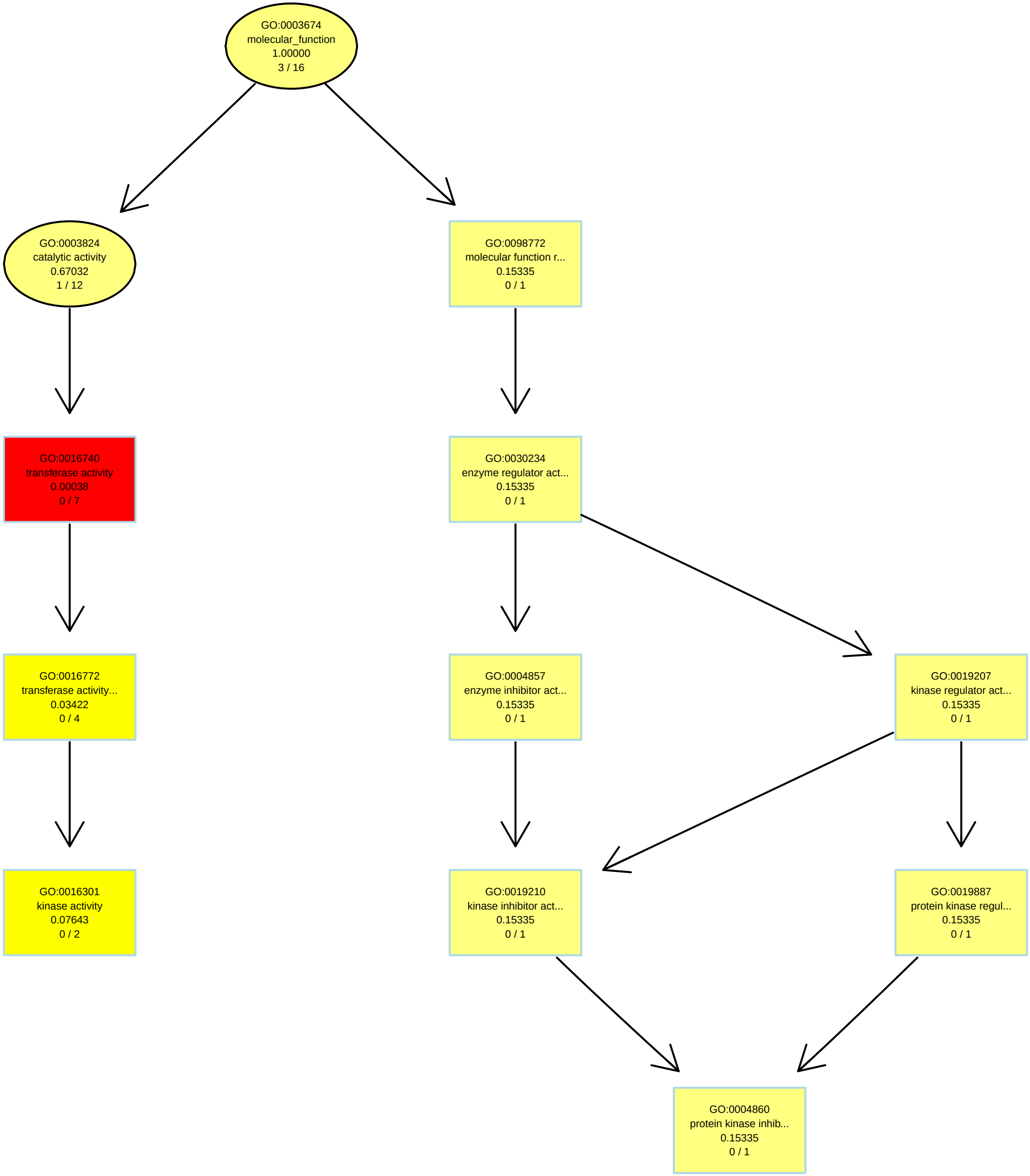

### PeakAreas200PC10_BP.pdf

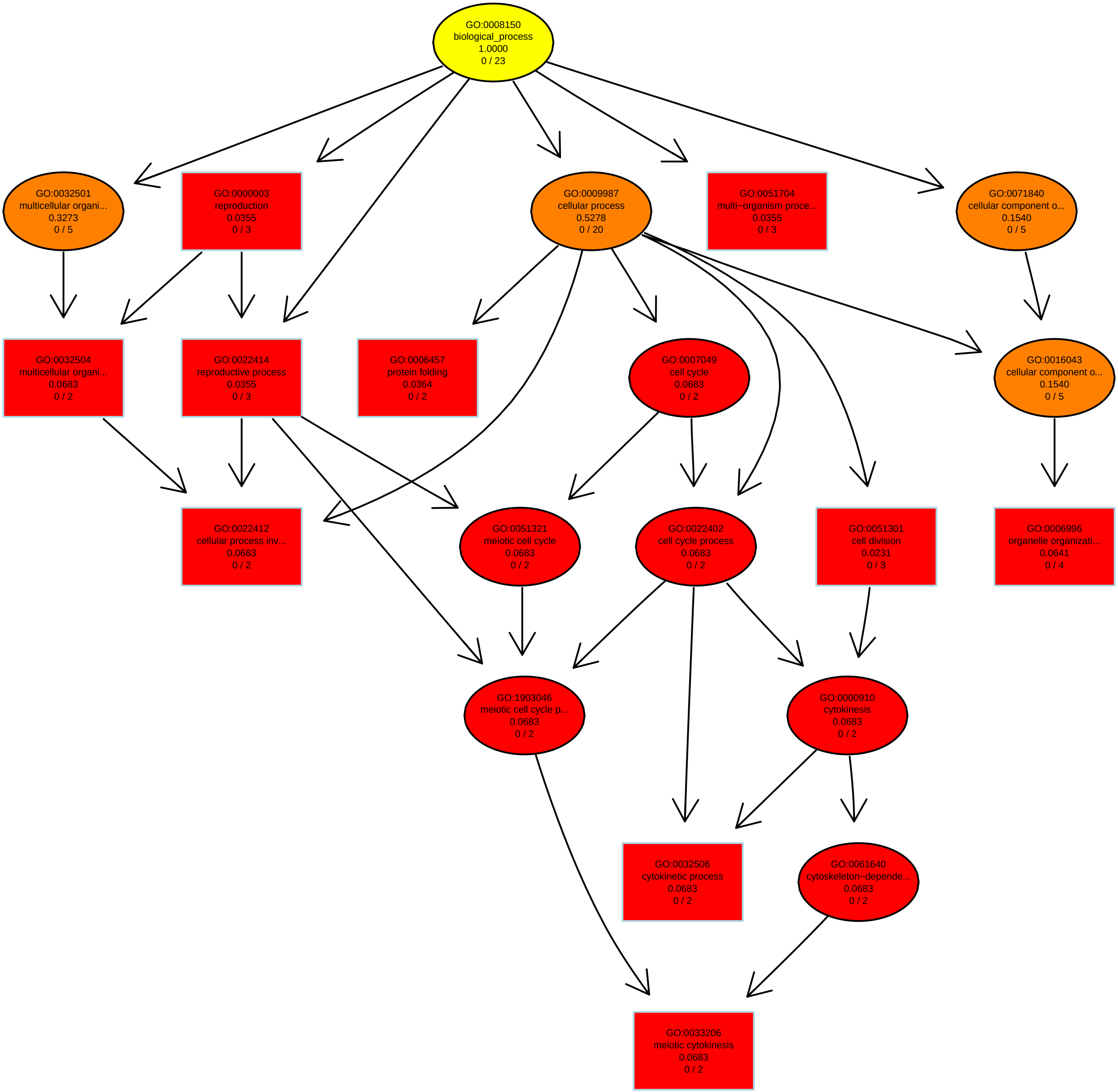

### PEAKAREASPC1_BP.pdf

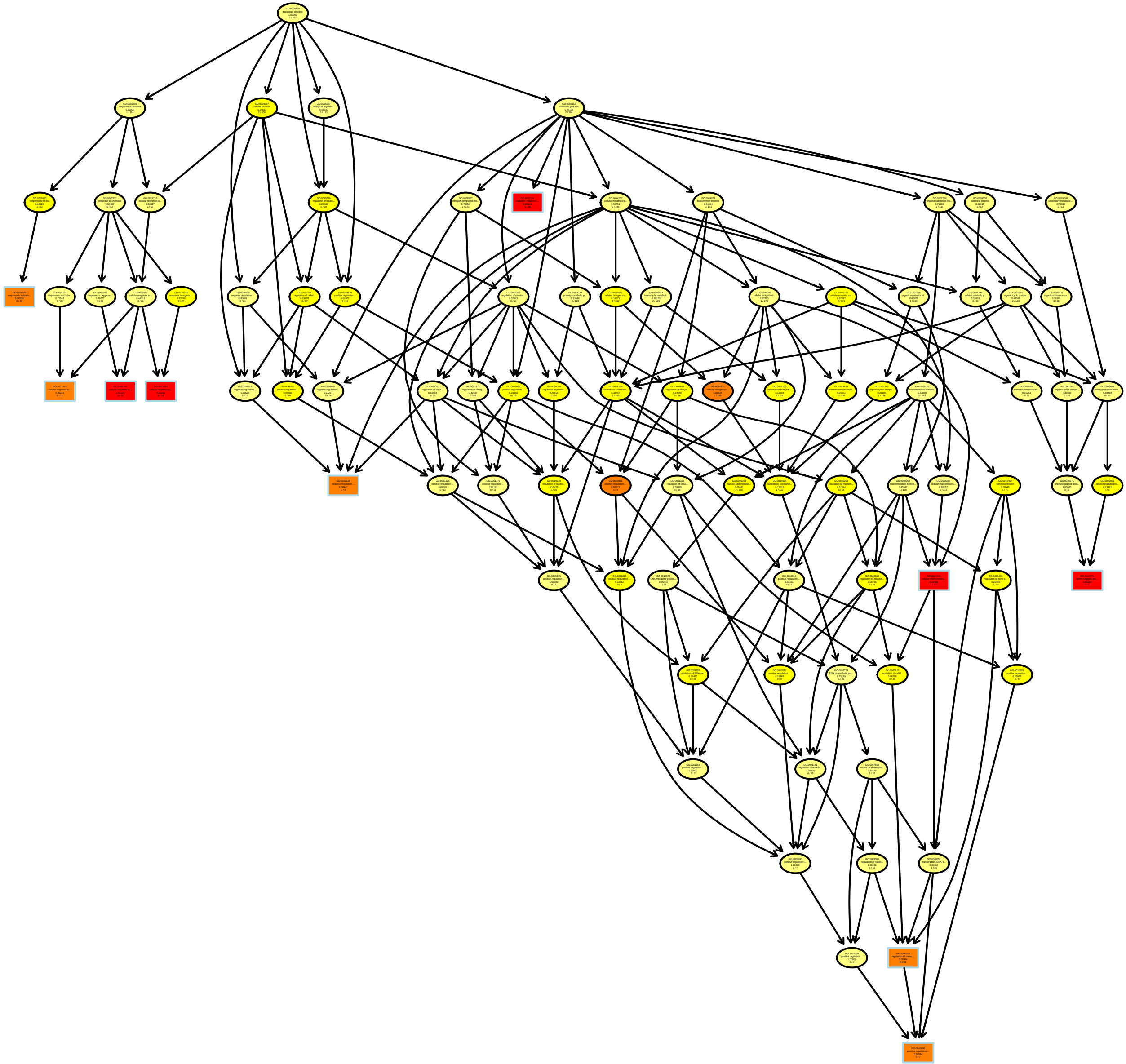

### PEAKAREASPC3_BP.pdf

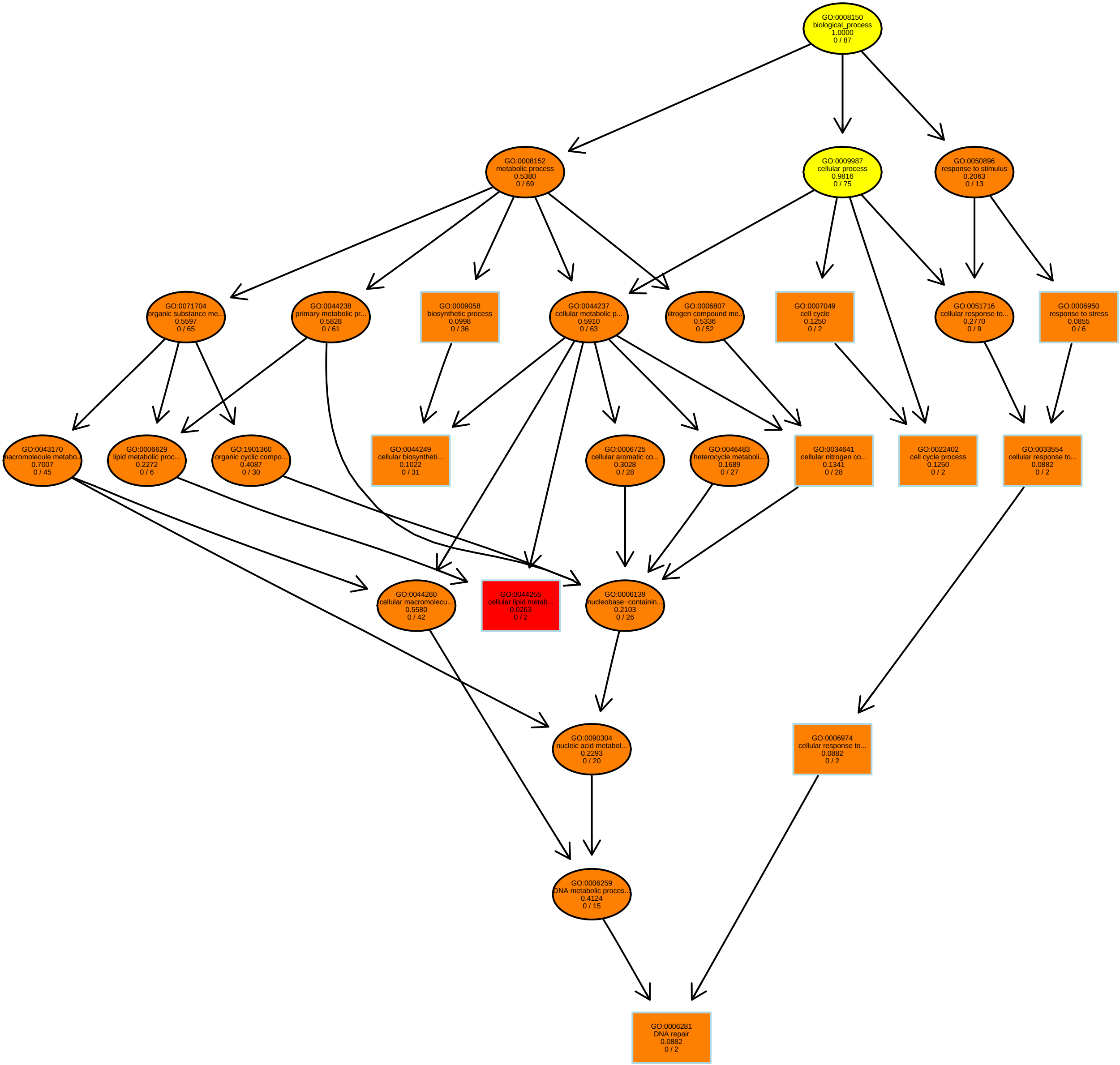

### PEAKAREASPC4_MF.pdf

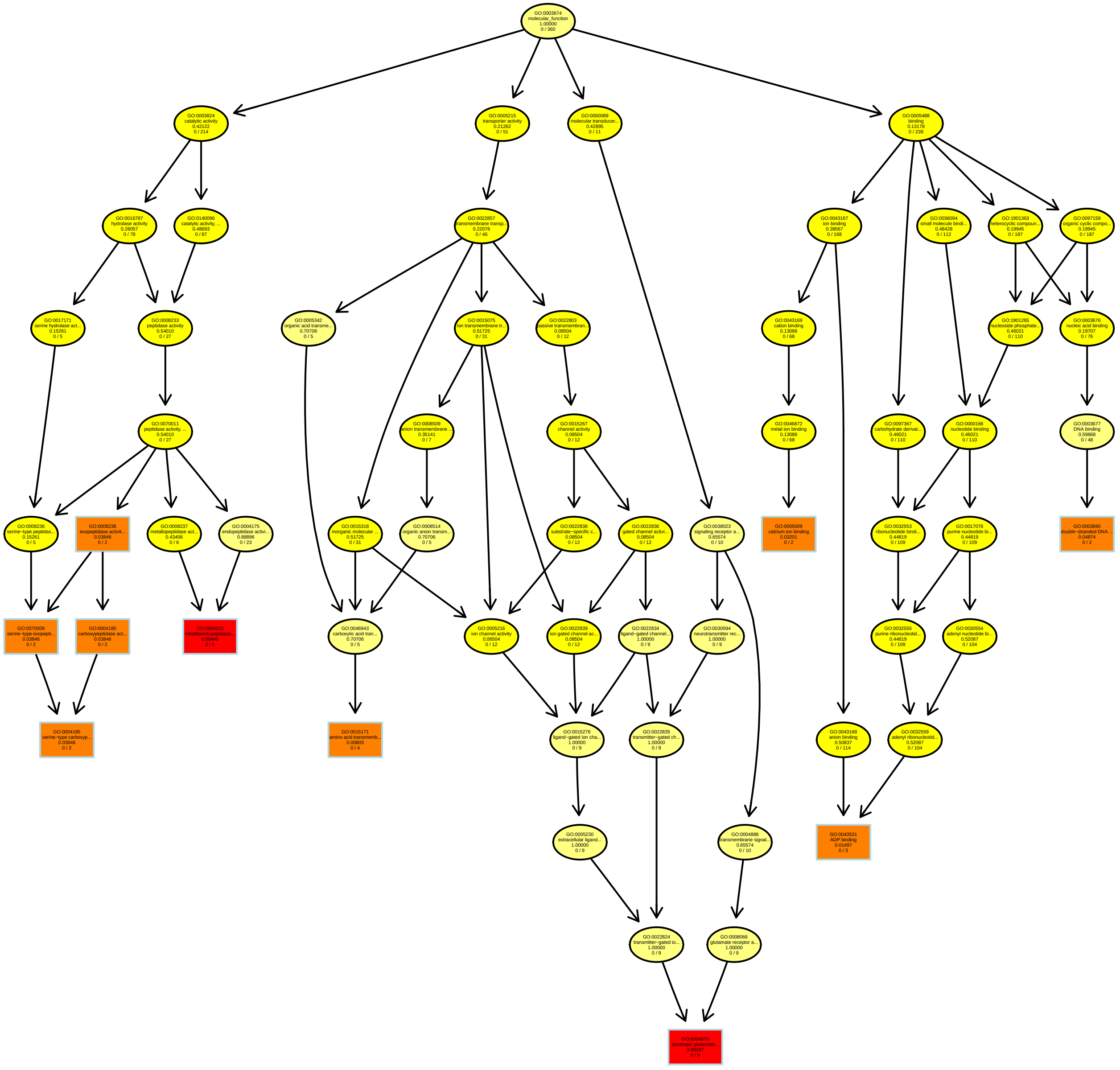

### PEAKAREASPC5_BP.pdf

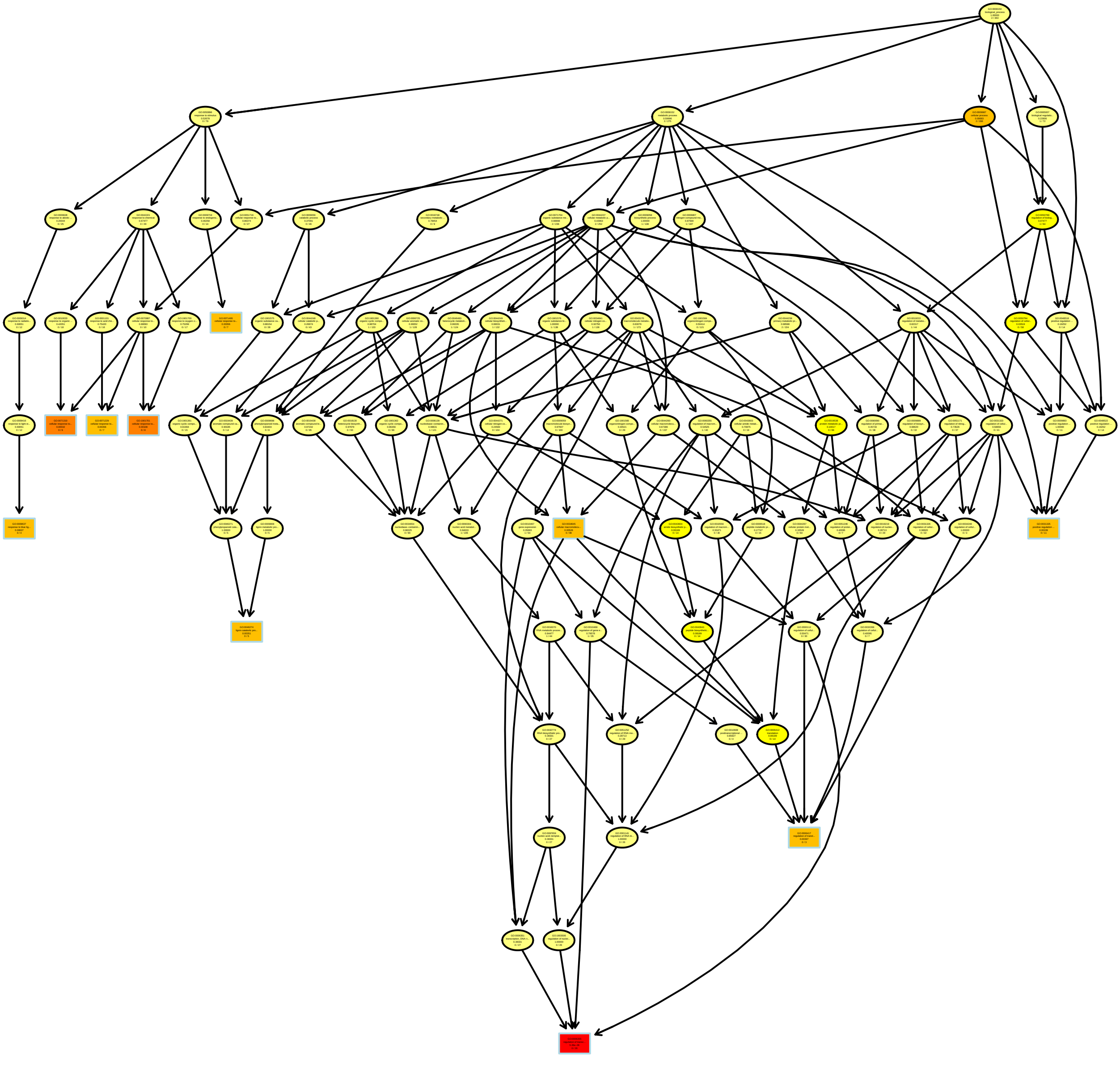

### PEAKAREASPC6_MF.pdf

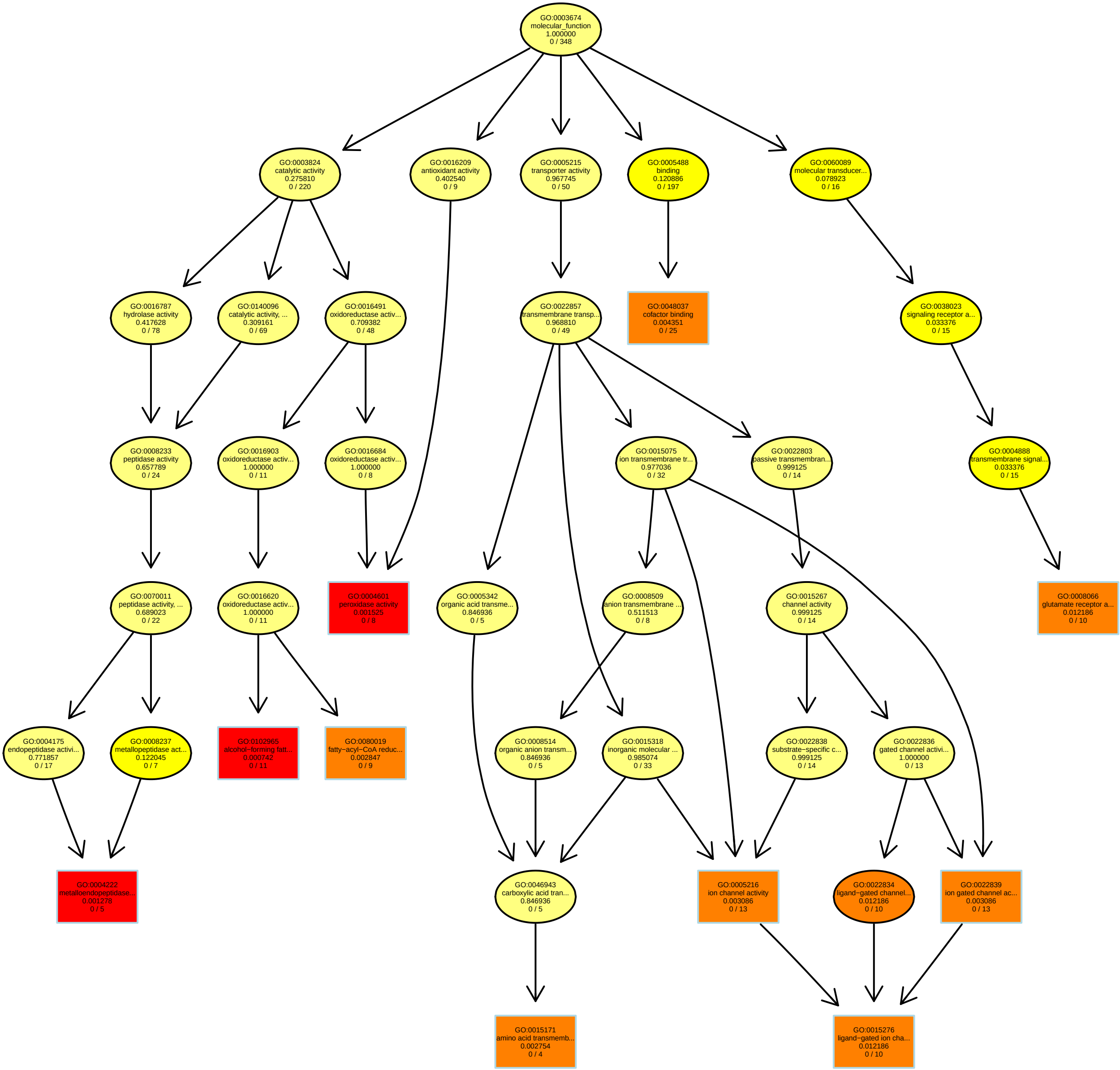

### PEAKAREASPC8_MF.pdf

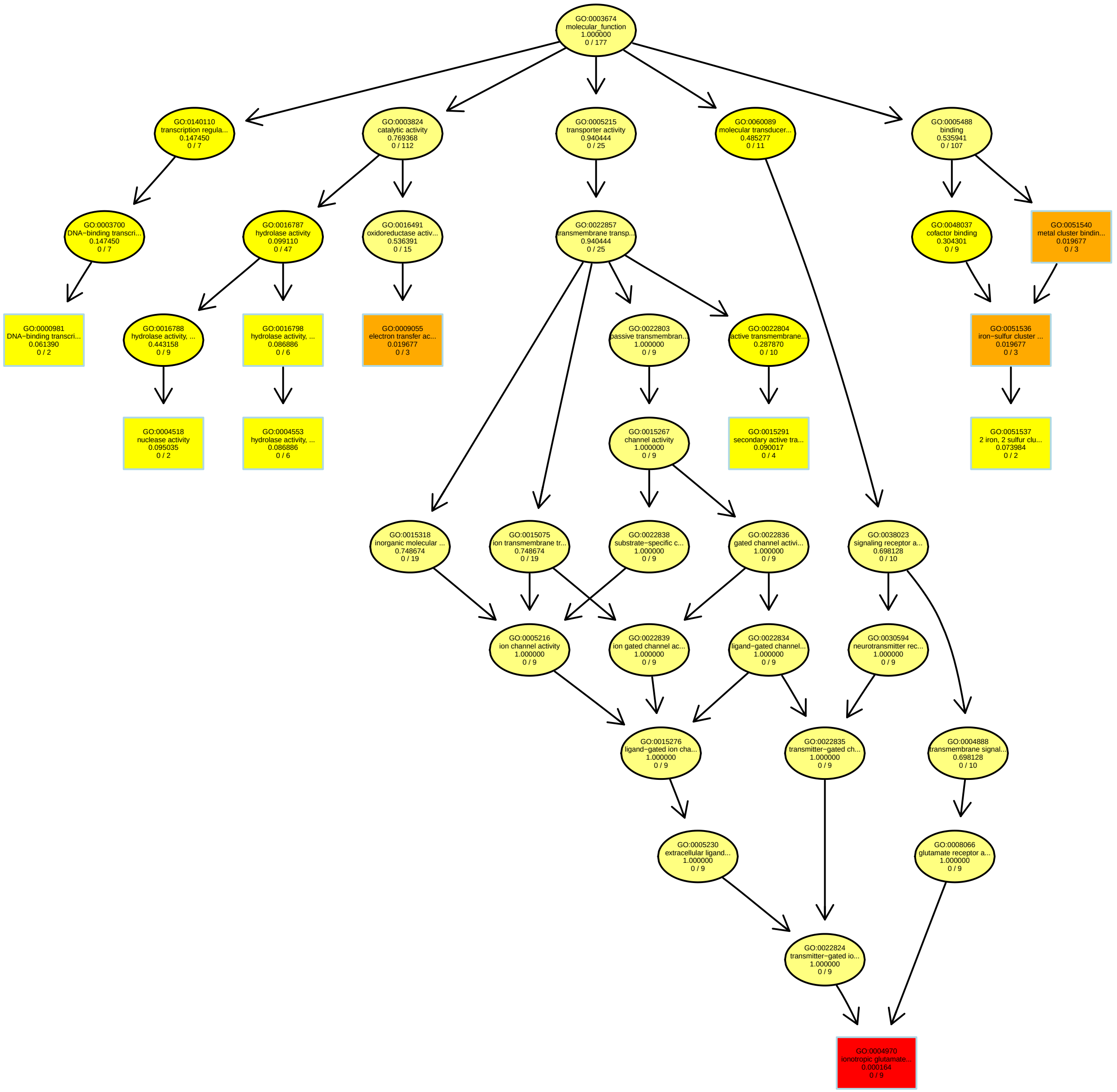

### PEAKAREASPC9_BP.pdf

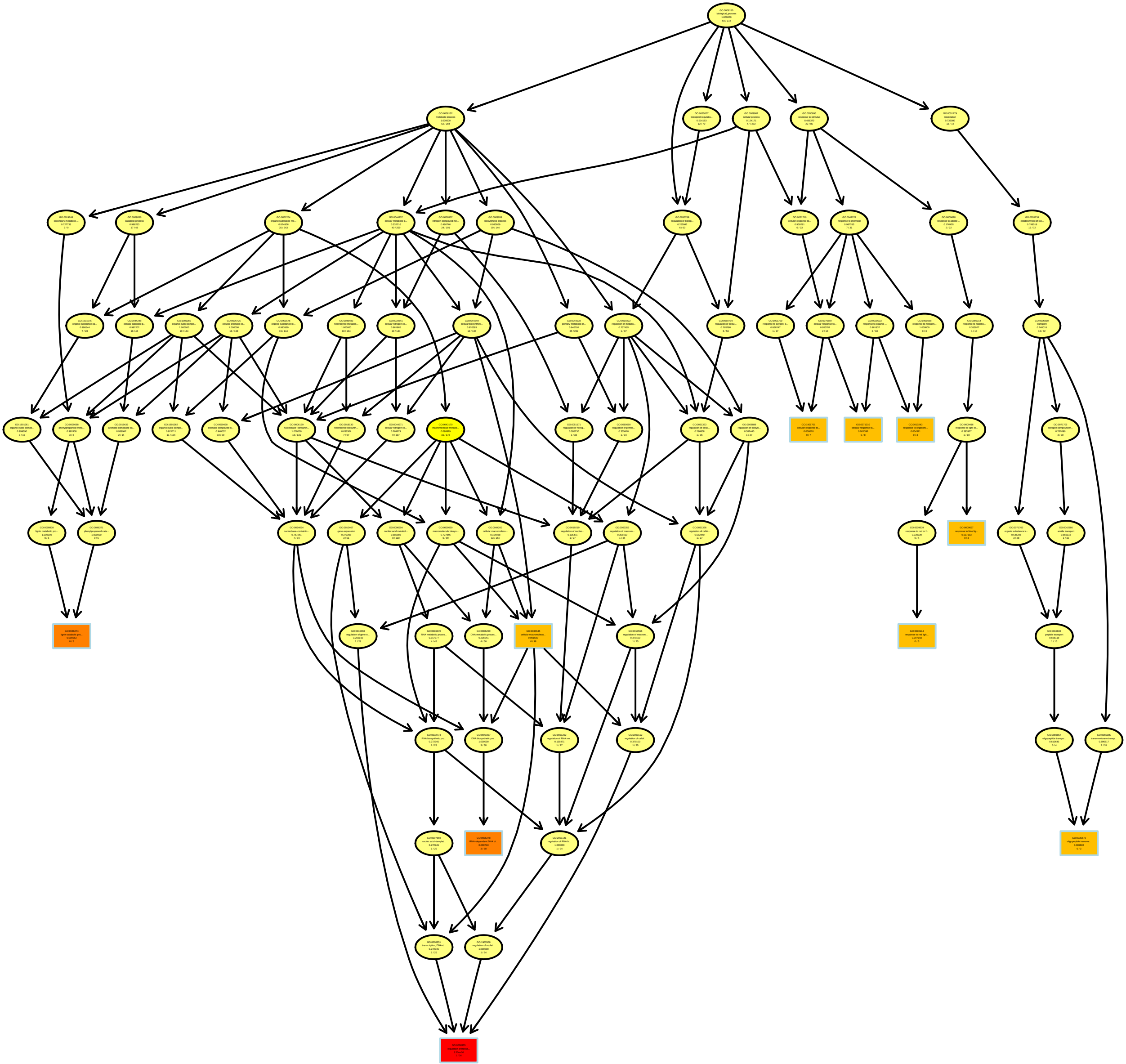

### PEAKAREASPC10_BP.pdf

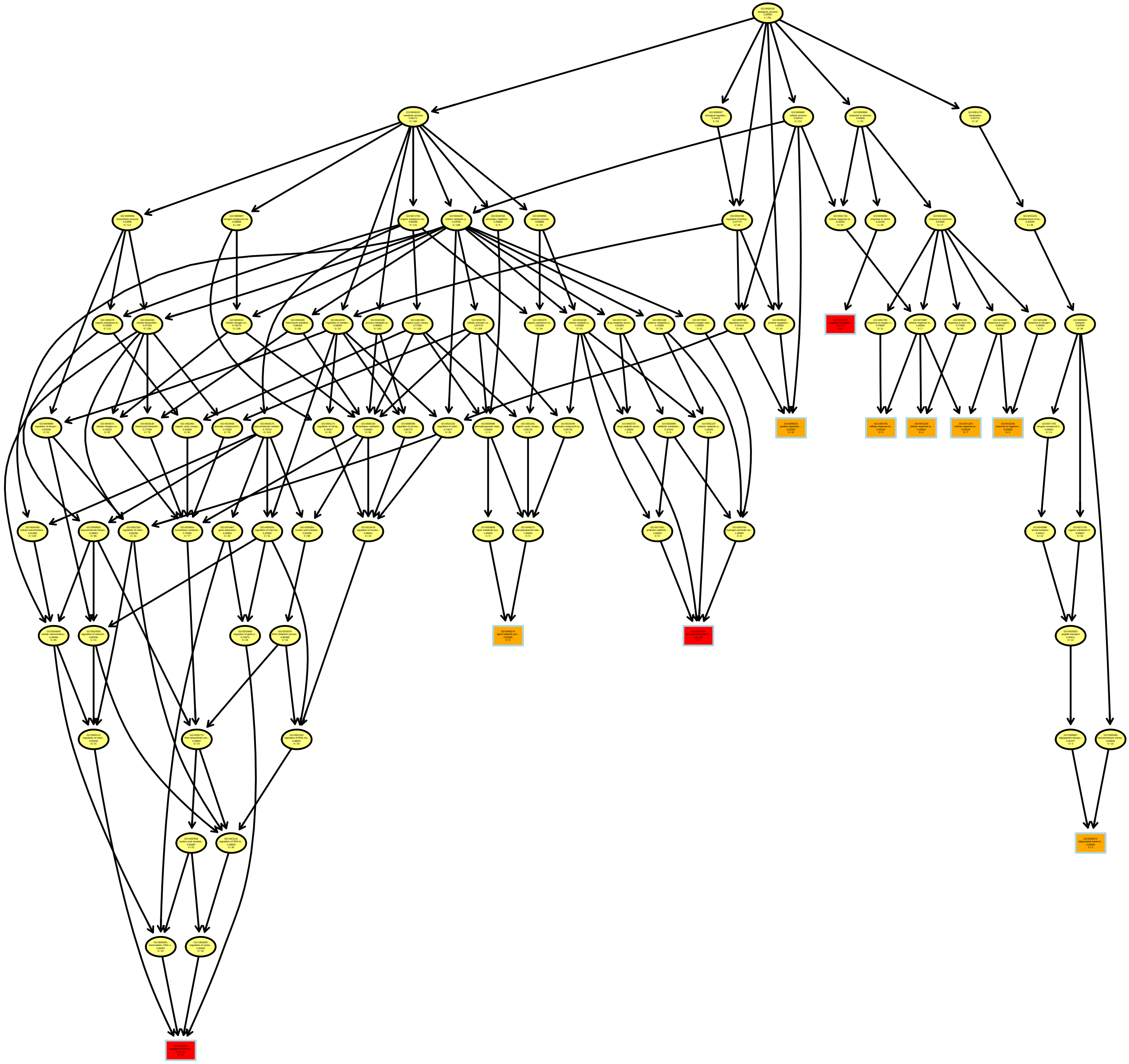

### Perimeter_MF.pdf

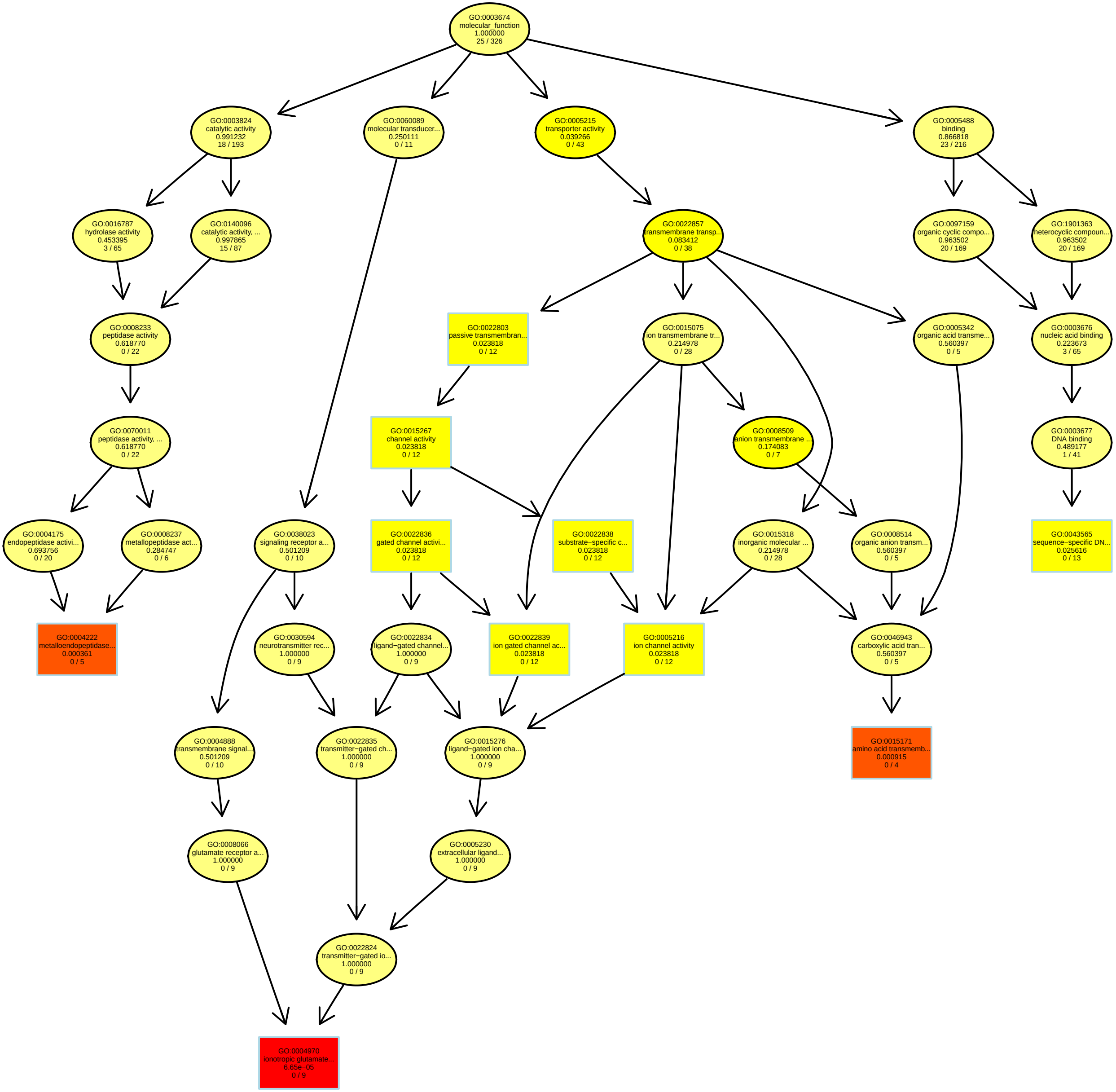

### RelativeRatioRT1.2_MF.pdf

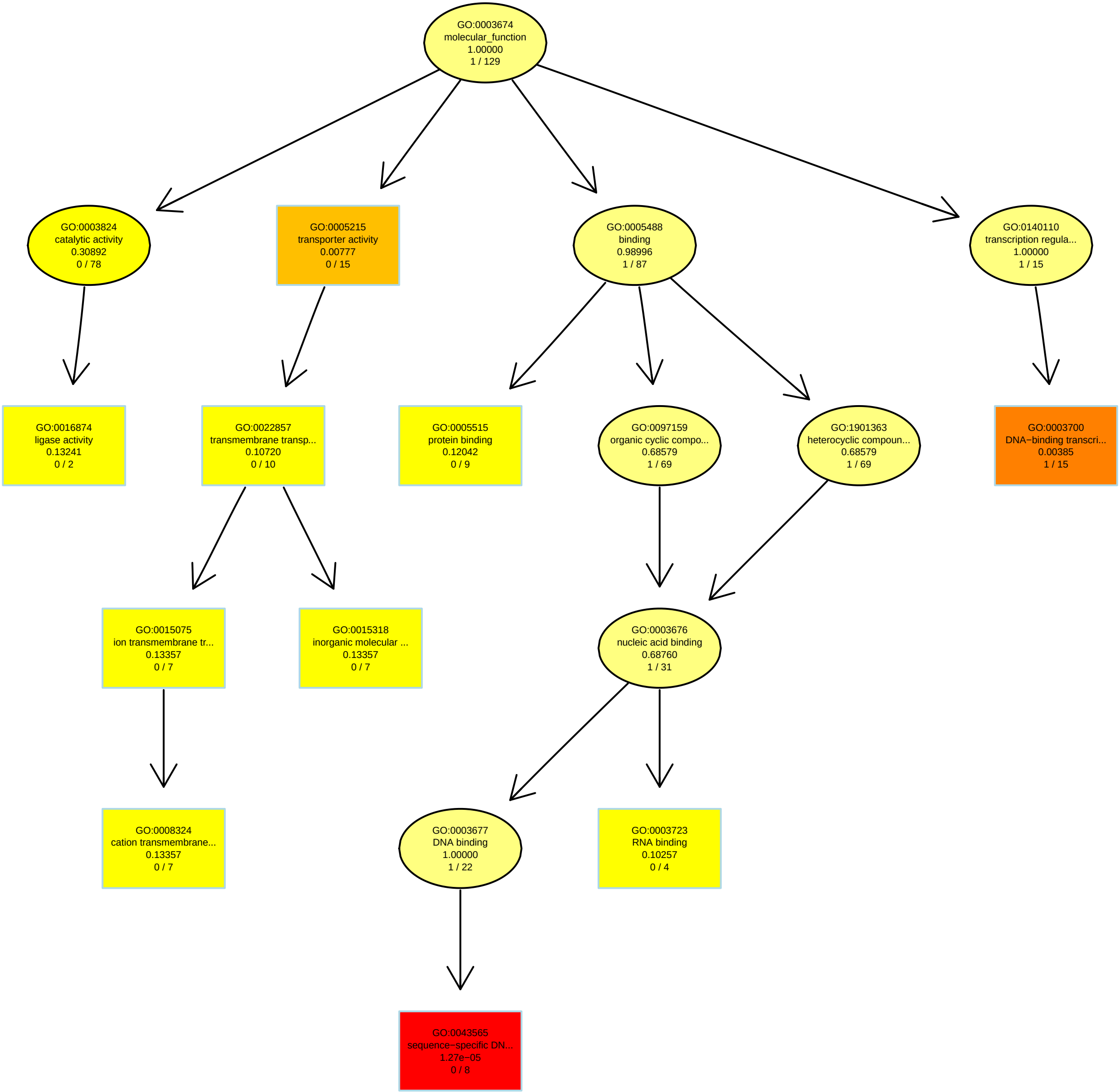

### RelativeRatioRT3.77_BP.pdf

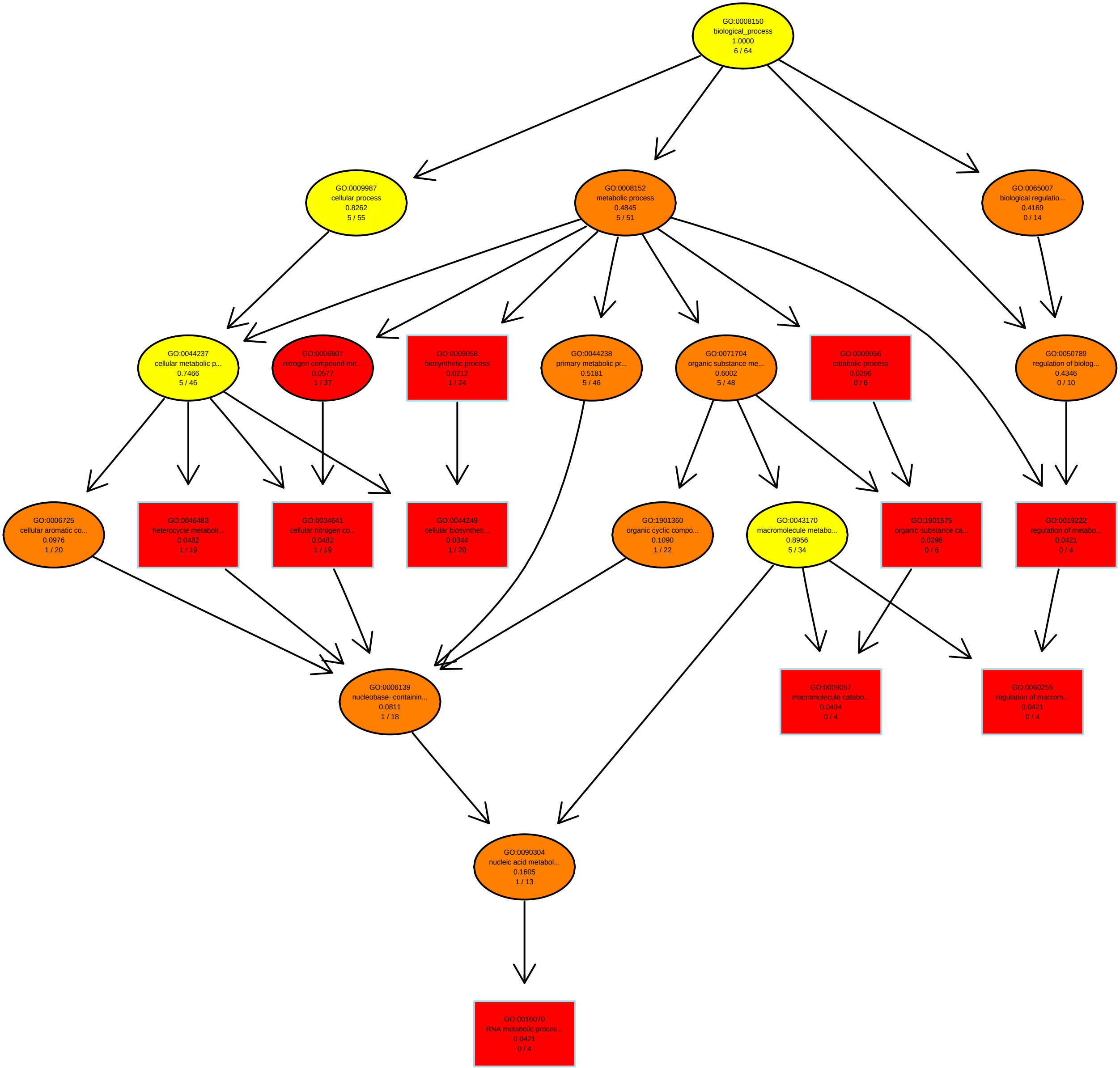

### RelativeRatioRT4.05_MF.pdf

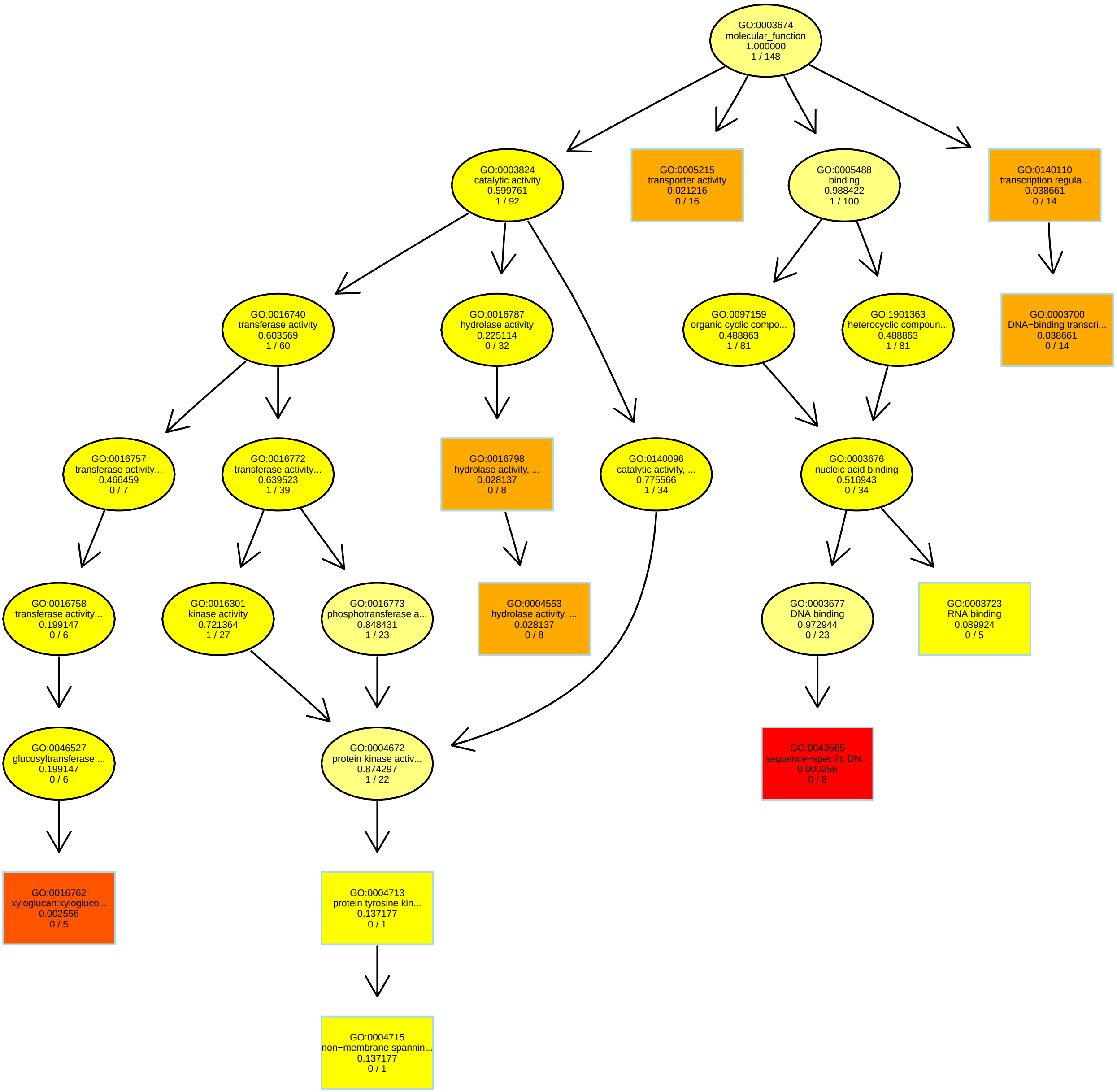

### RelativeRatioRT4.6_BP.pdf

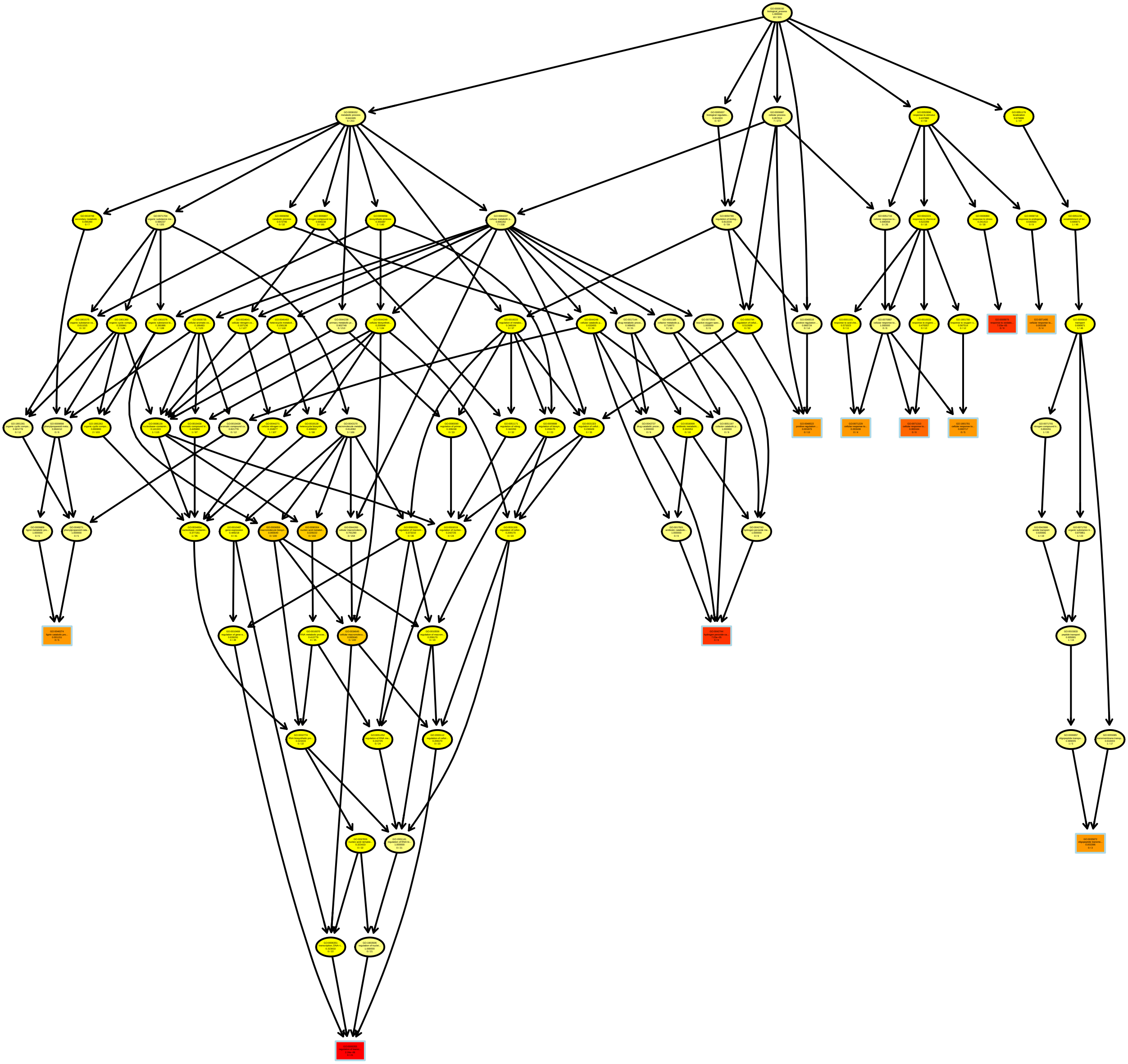

### RelativeRatioRT4.21_MF.pdf

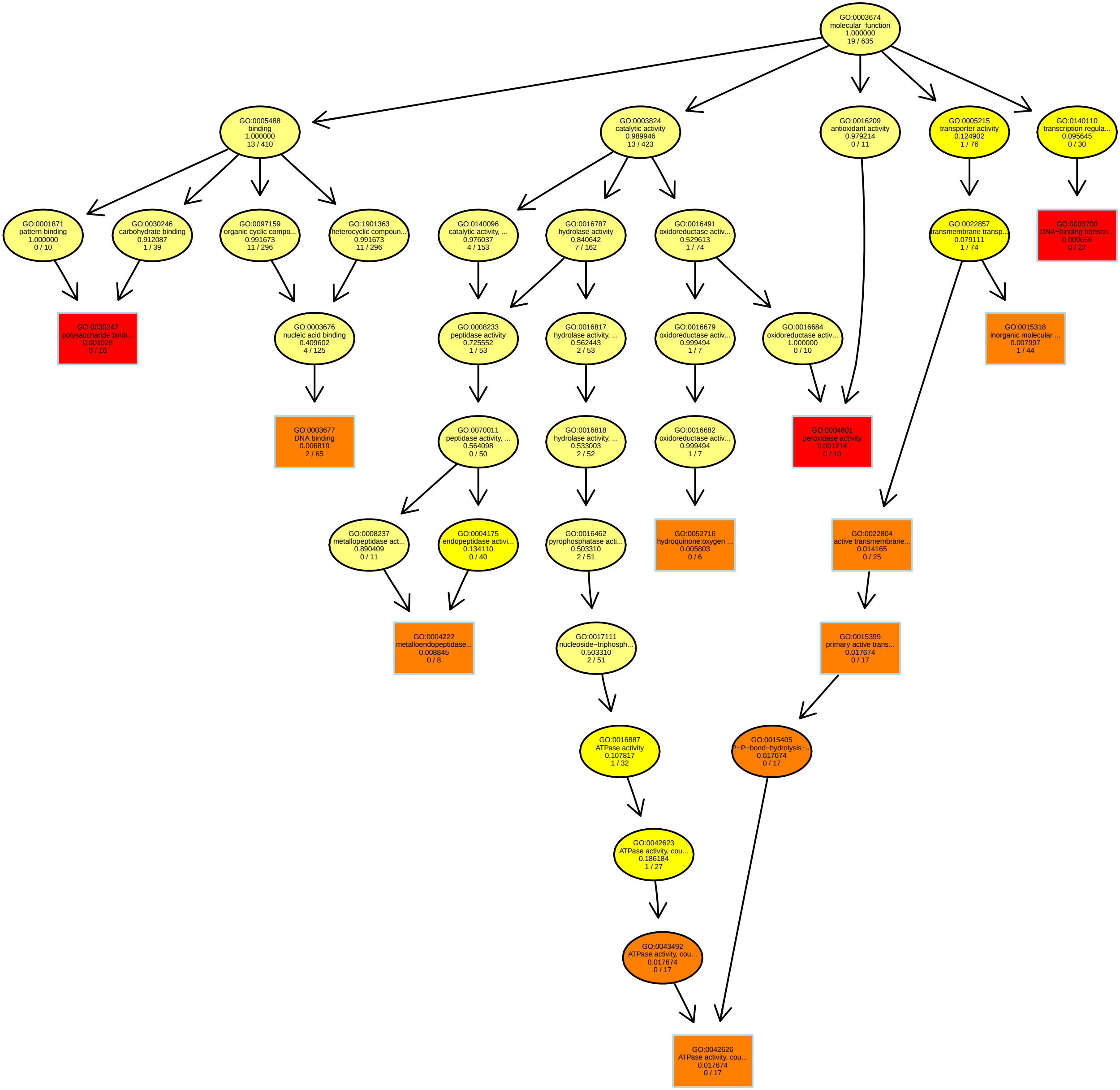

### RelativeRatioRT4.42_MF.pdf

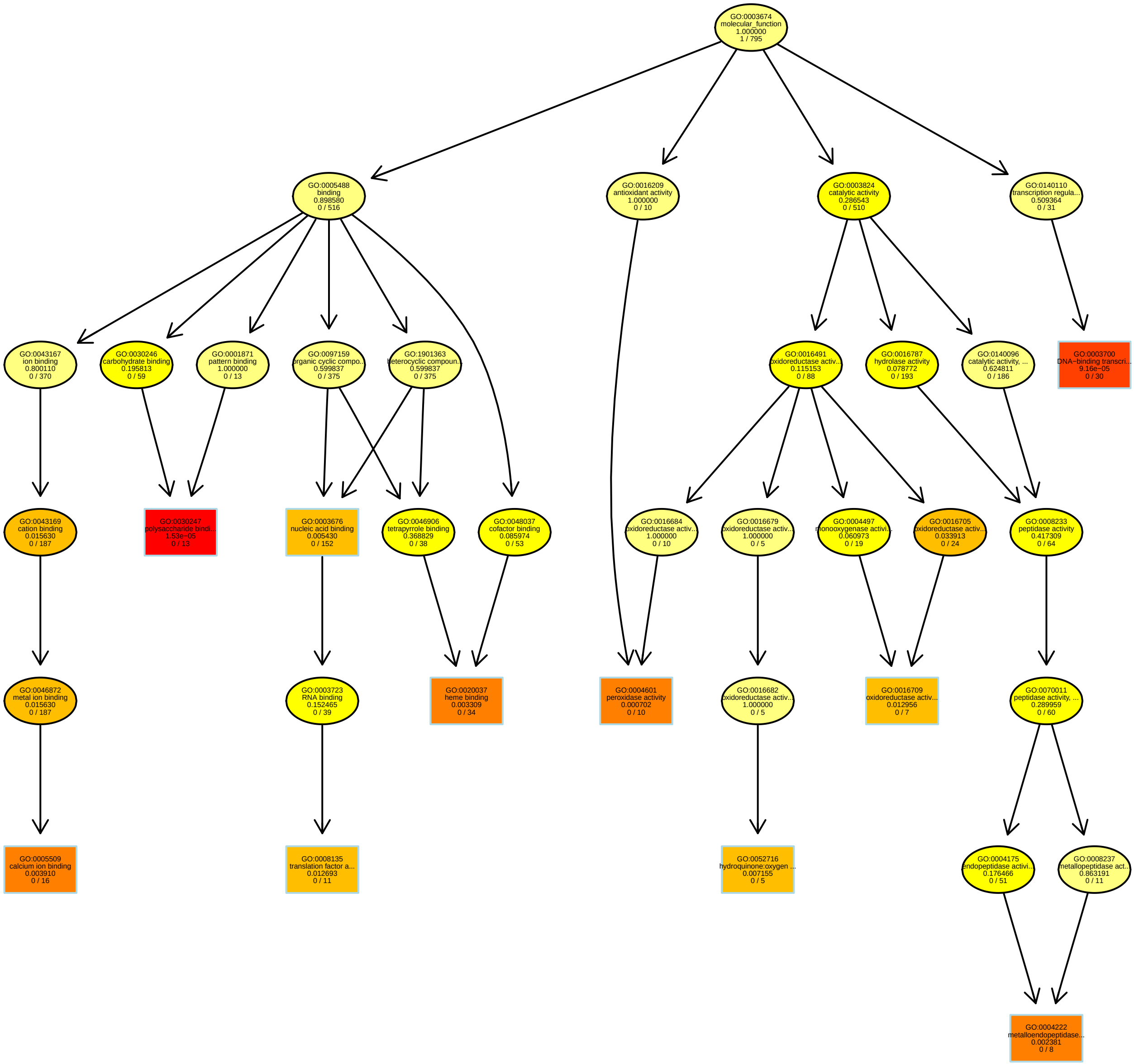

### RelativeRatioRT4.55_BP.pdf

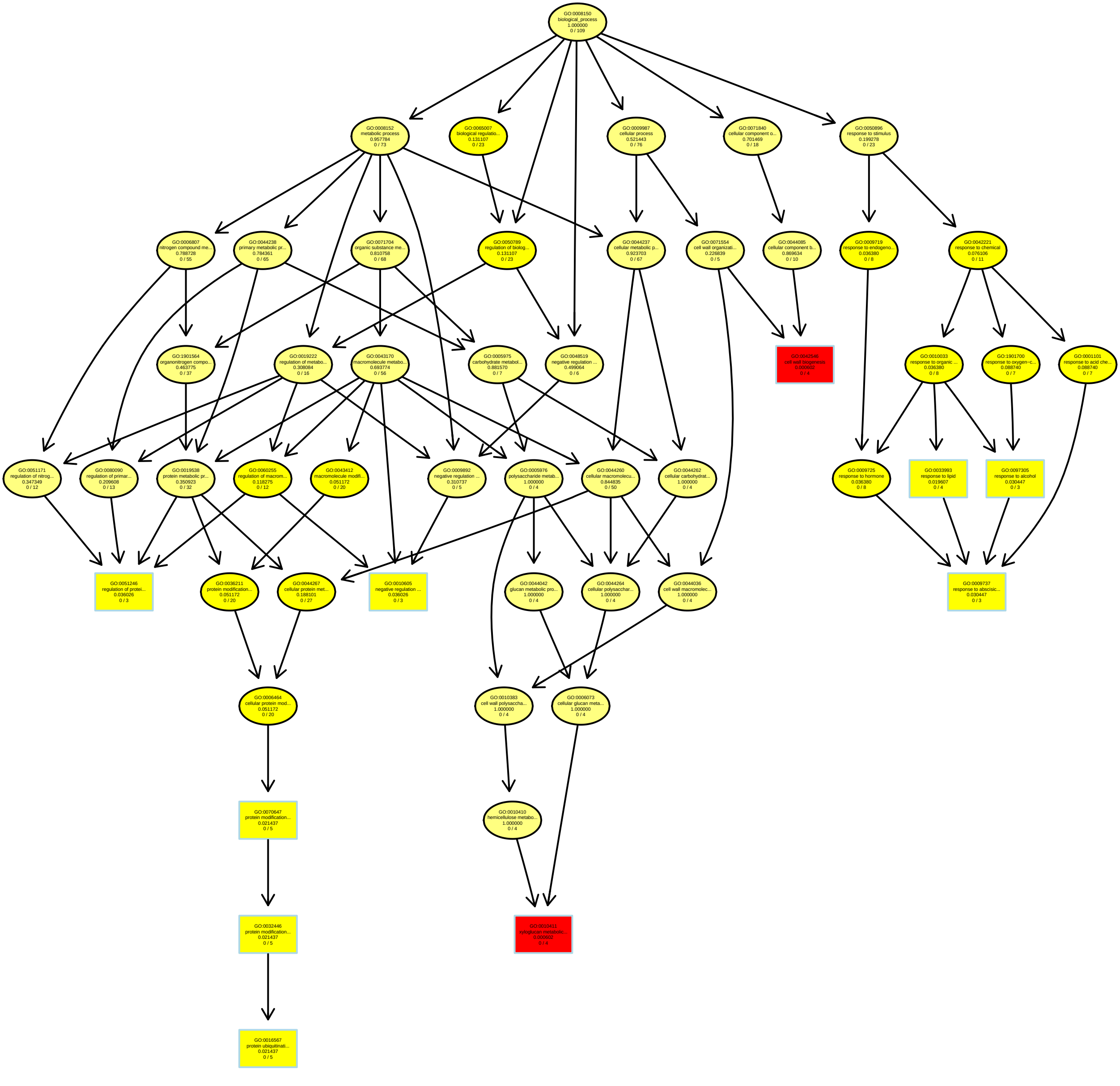

### RelativeRatioRT4.87_MF.pdf

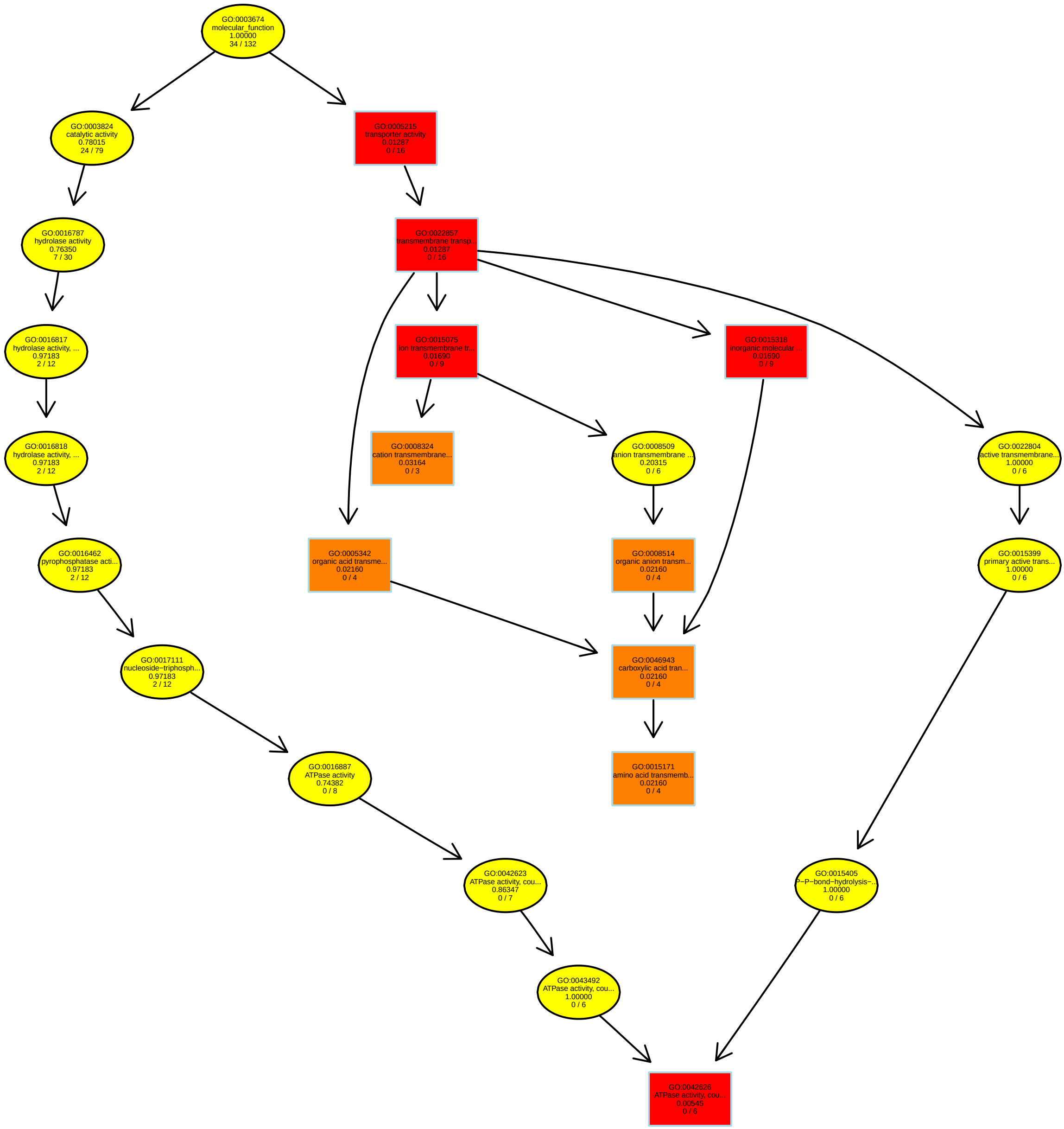

### RelativeRatioRT5.53_BP.pdf

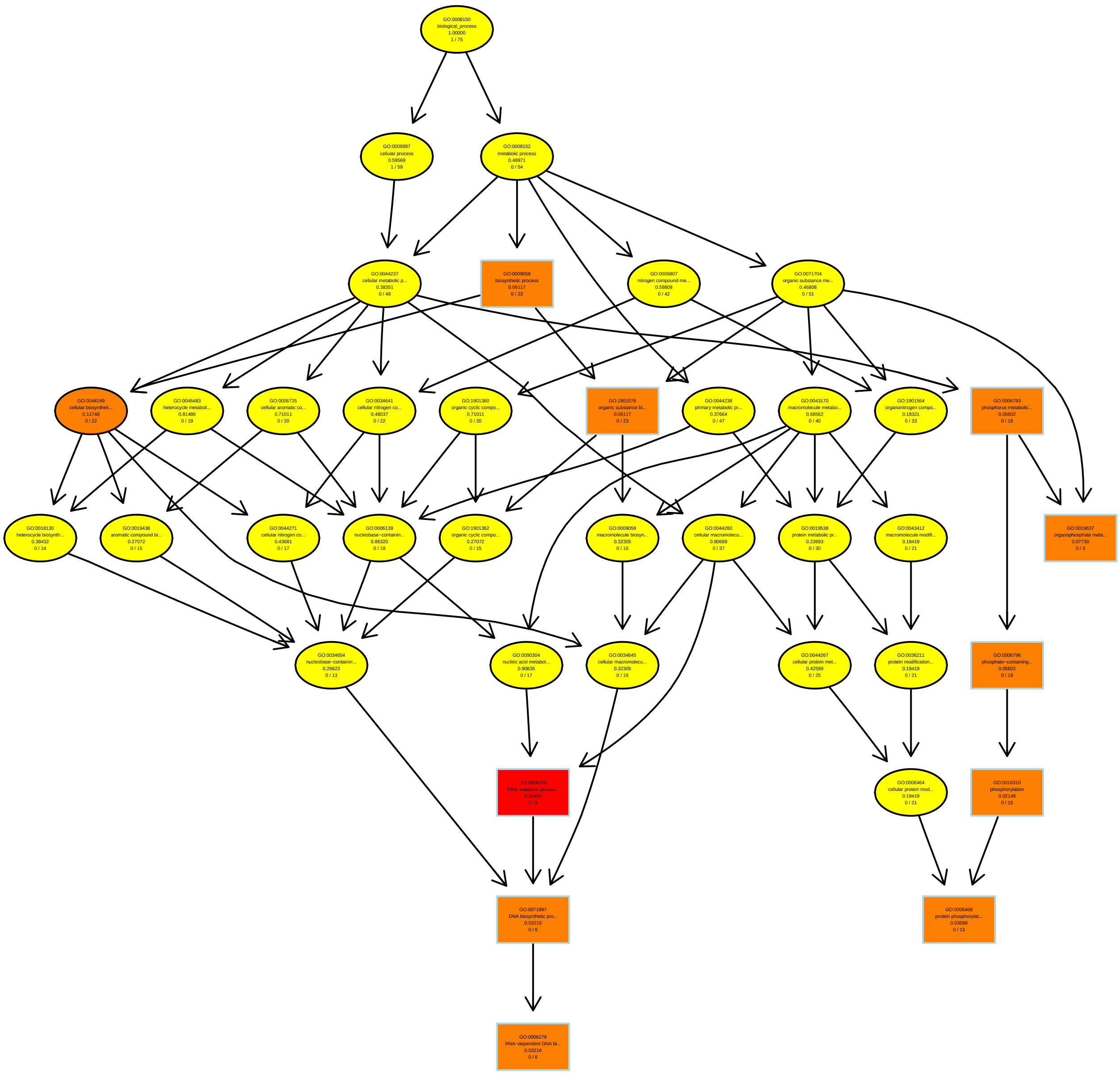

### RelativeRatioRT5.89_MF.pdf

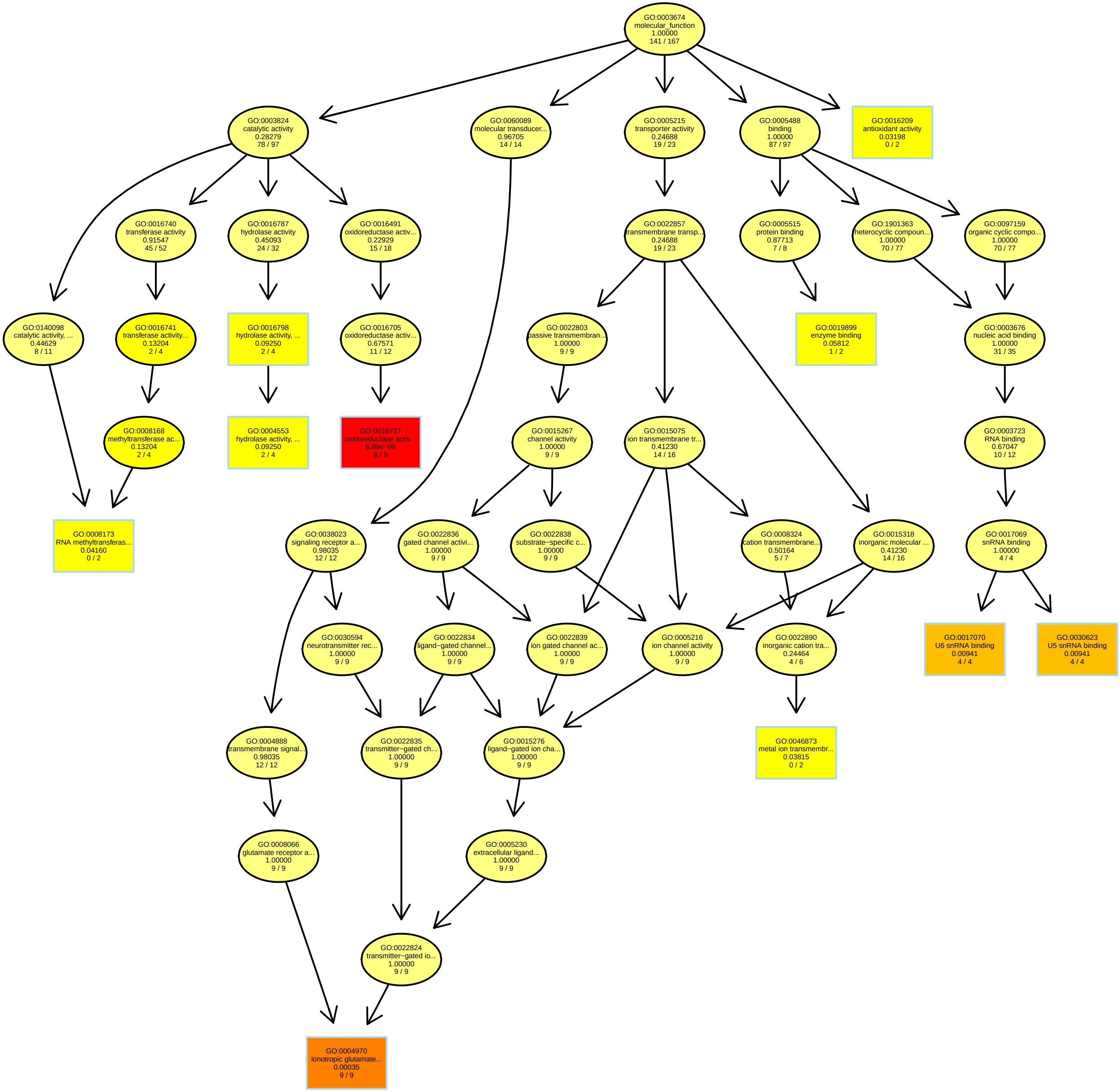

### RelativeRatioRT6.37_BP.pdf

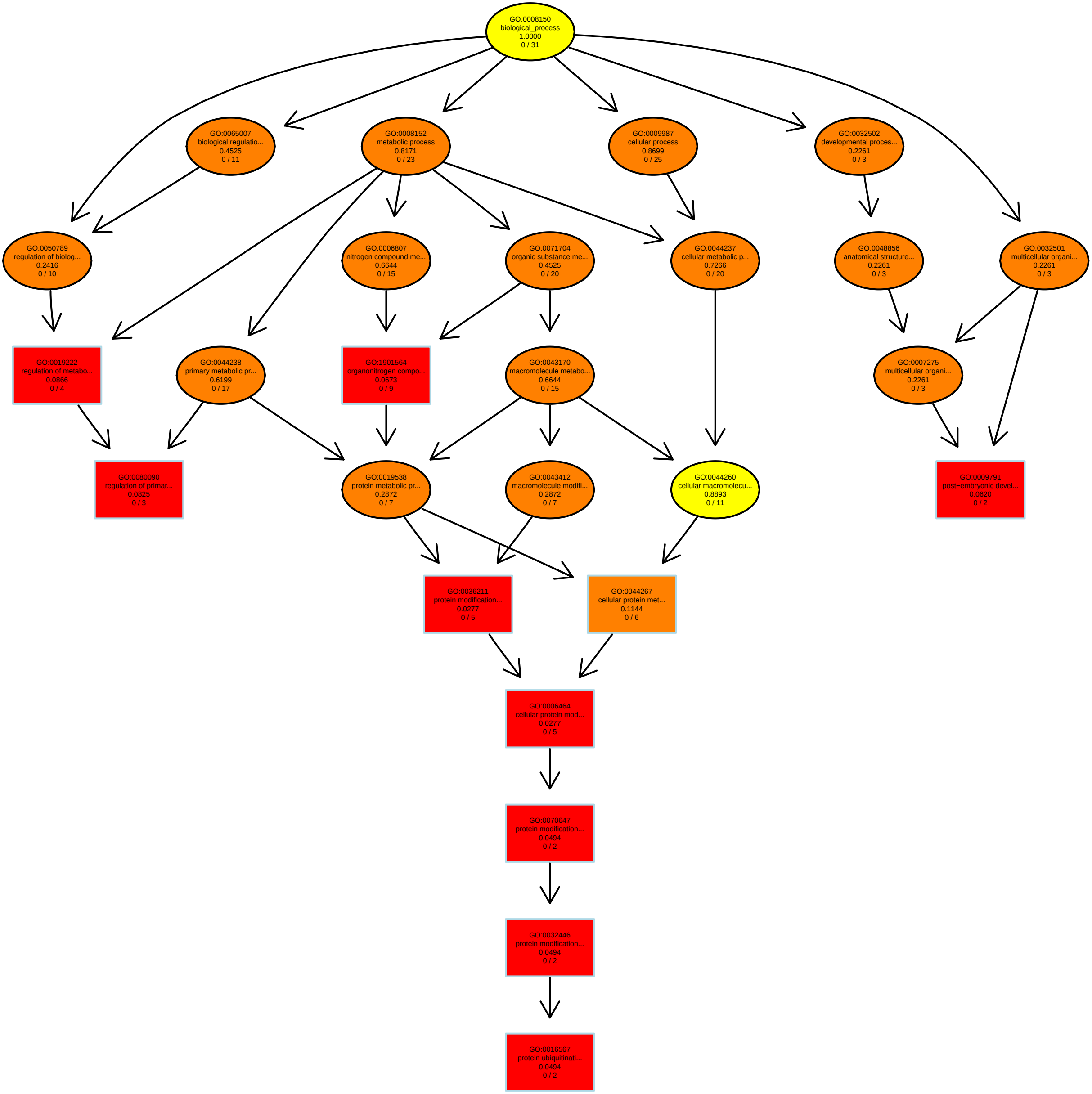

### RelativeRatioRT7.28_MF.pdf

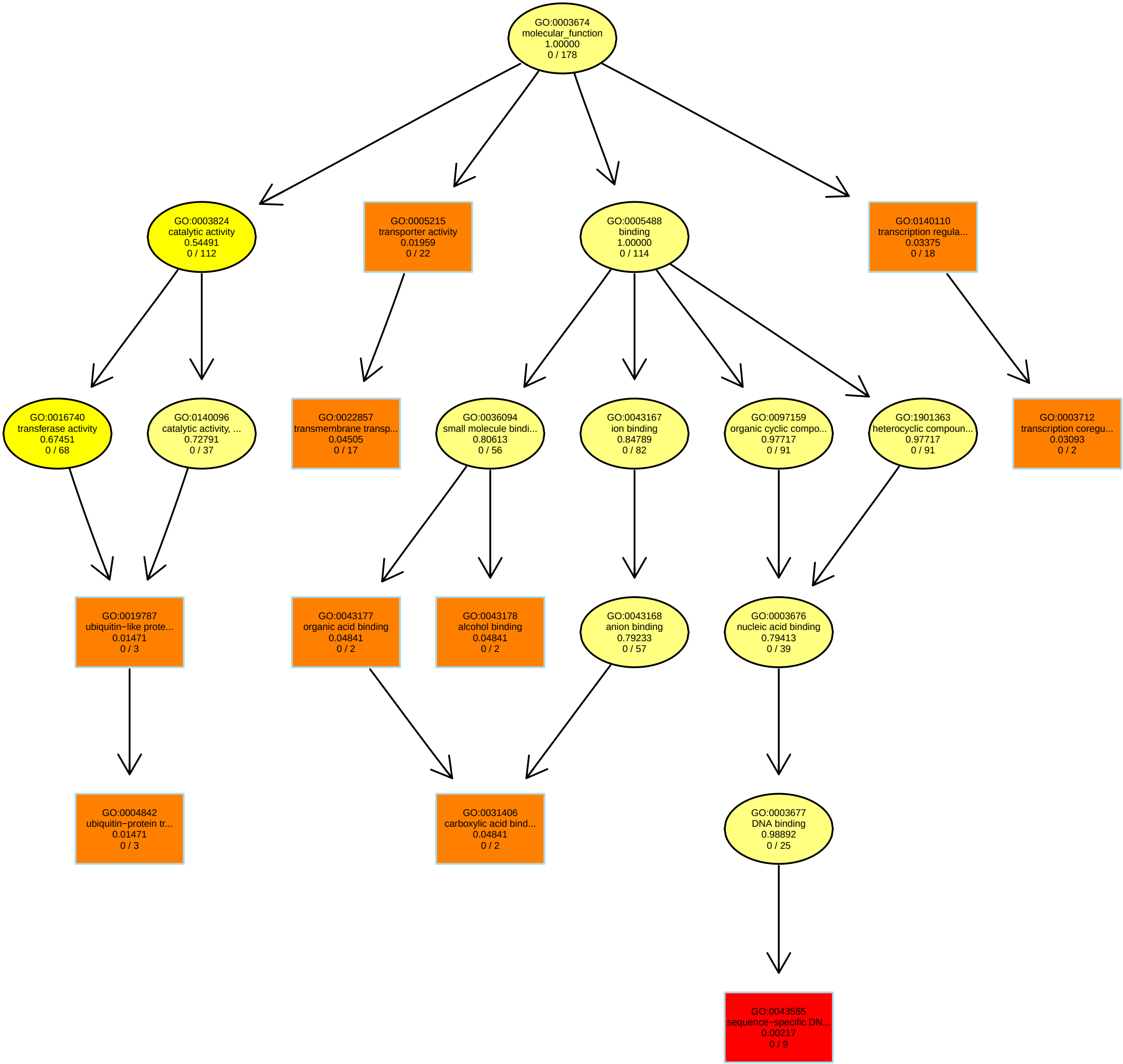

### RelativeRatioRT7.47_MF.pdf

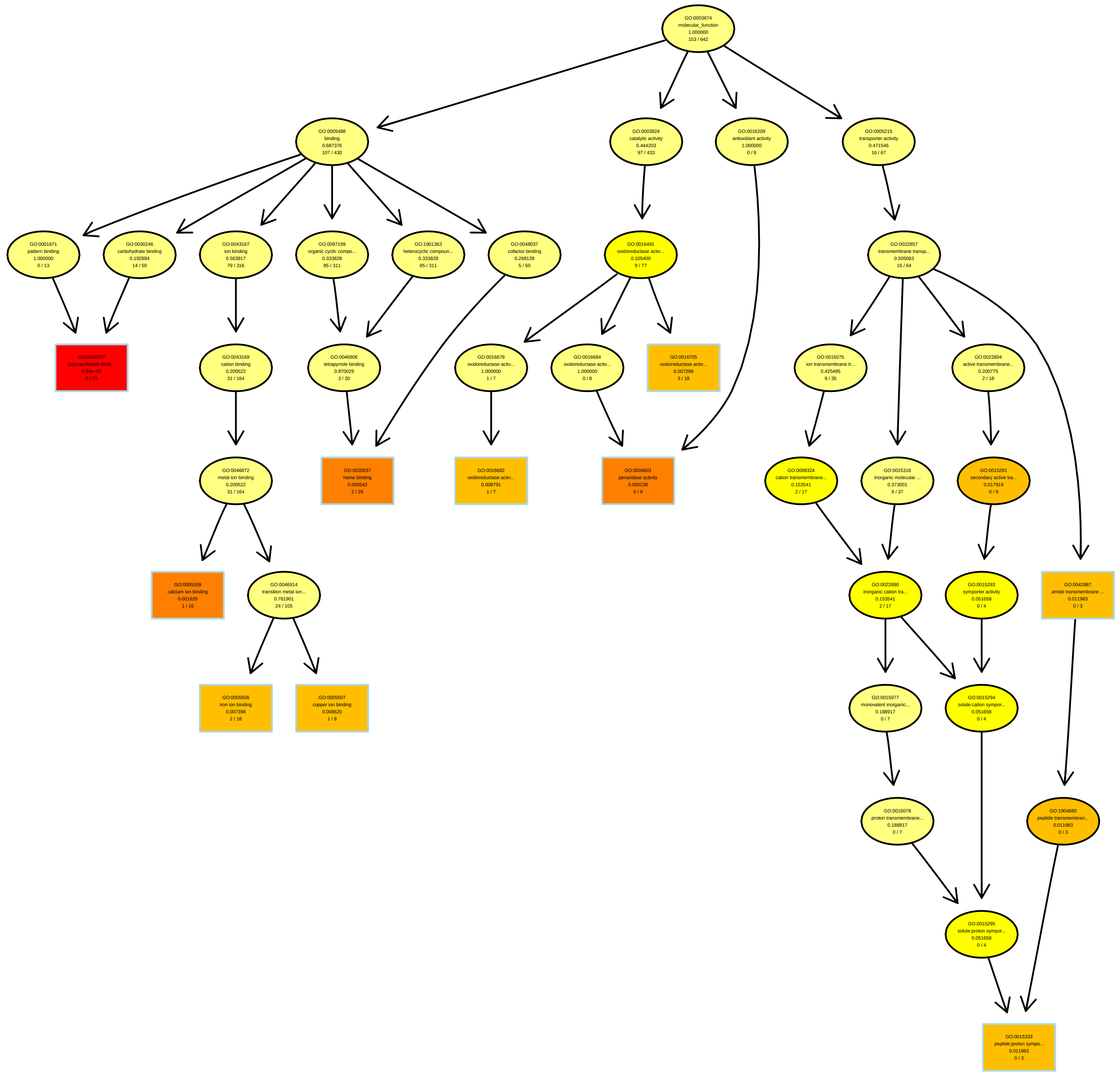

### RelativeRatioRT7.71_MF.pdf

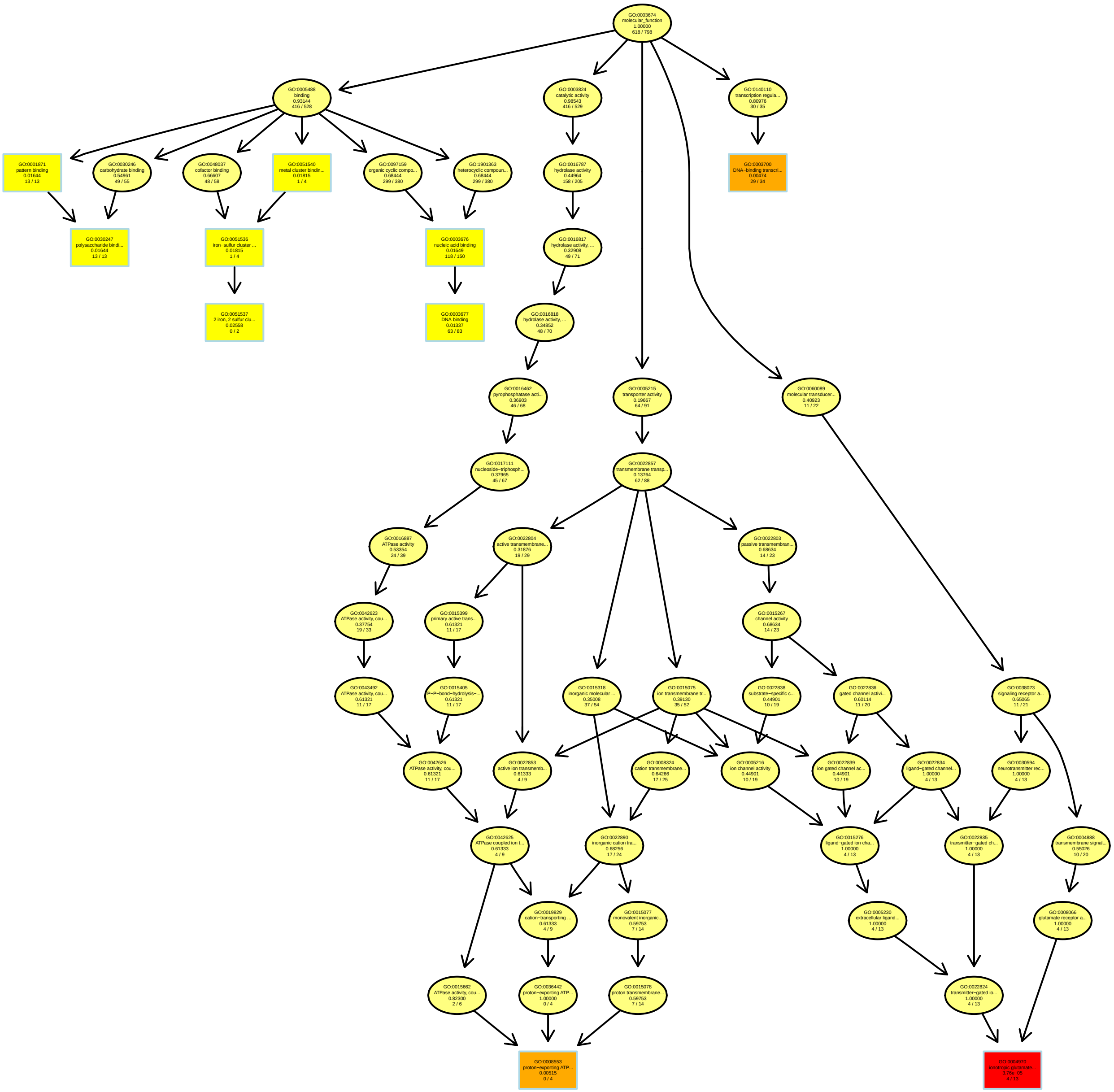

### RelativeRatioRT7.93_BP.pdf

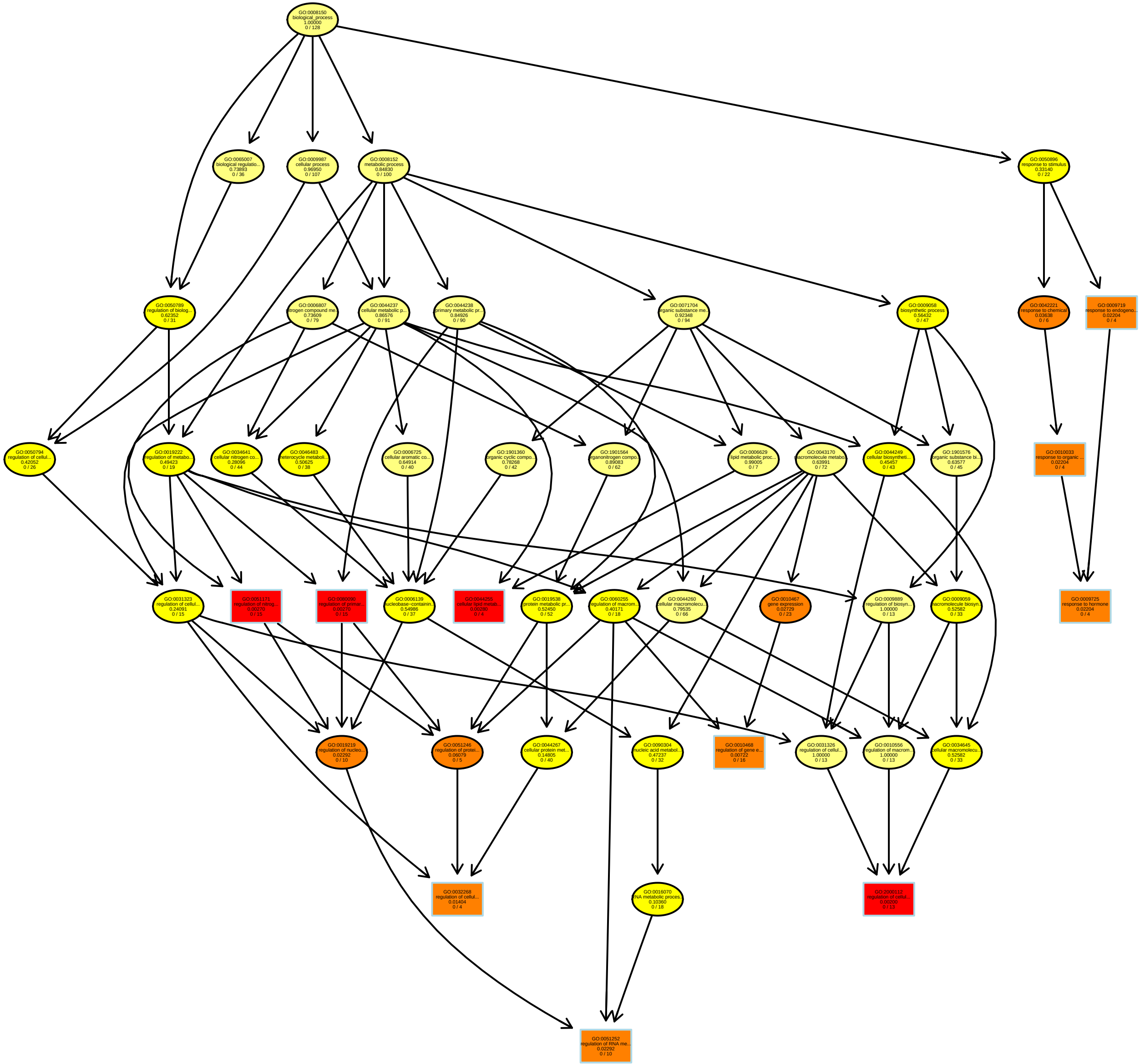

### RelativeRatioRT8.11_MF.pdf

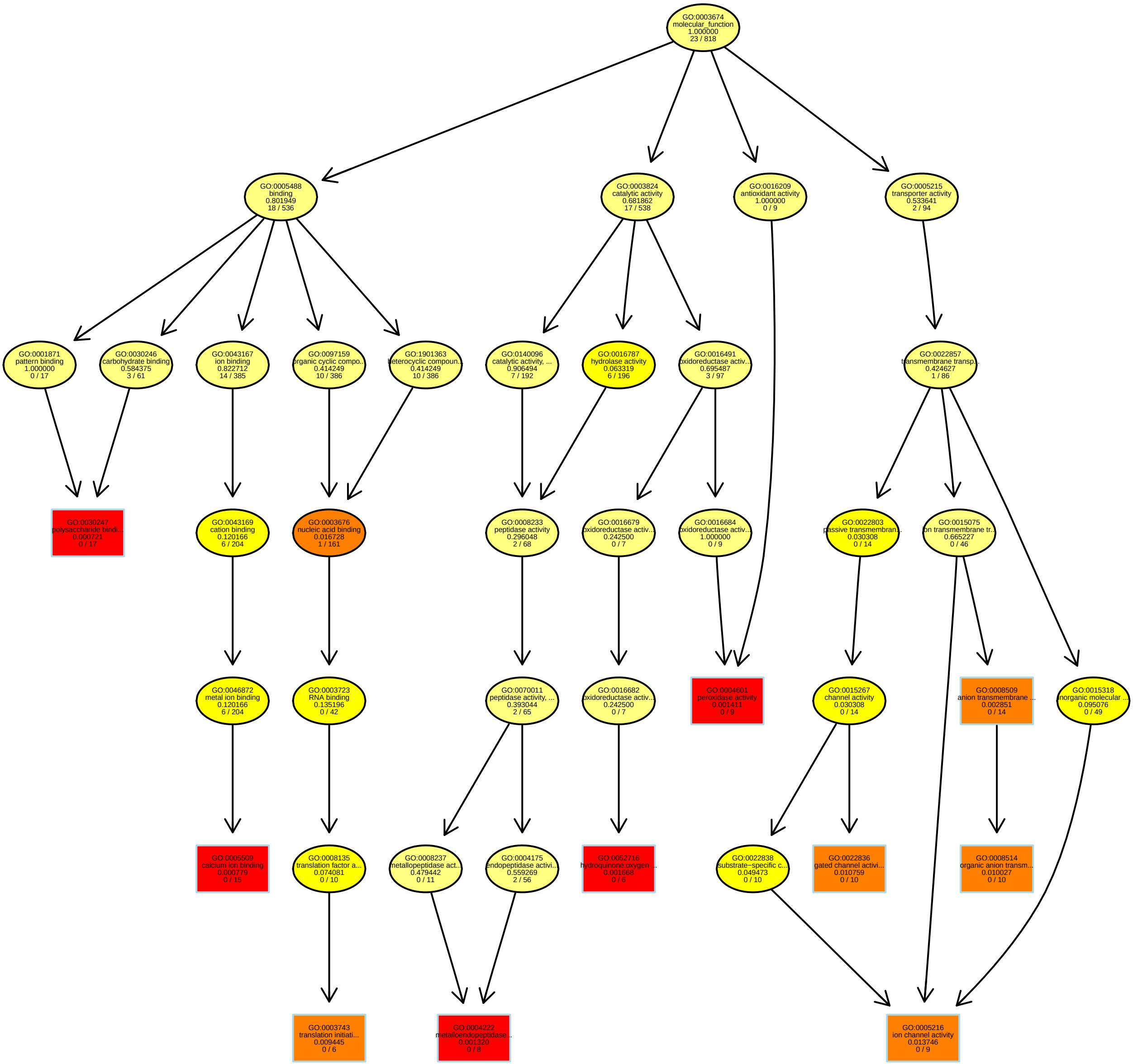

### RelativeRatioRT8.38_BP.pdf

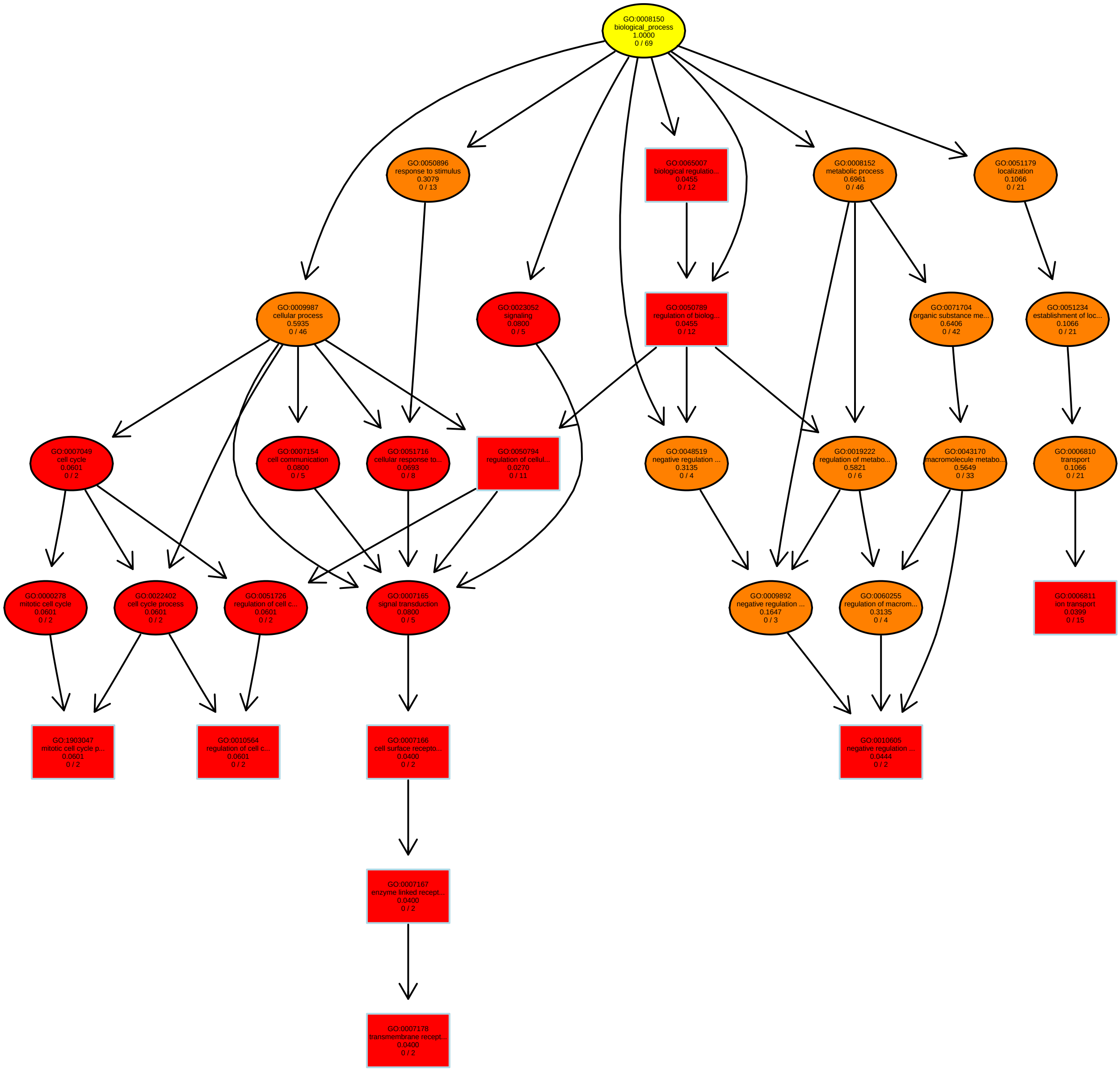

### RelativeRatioRT8.51_BP.pdf

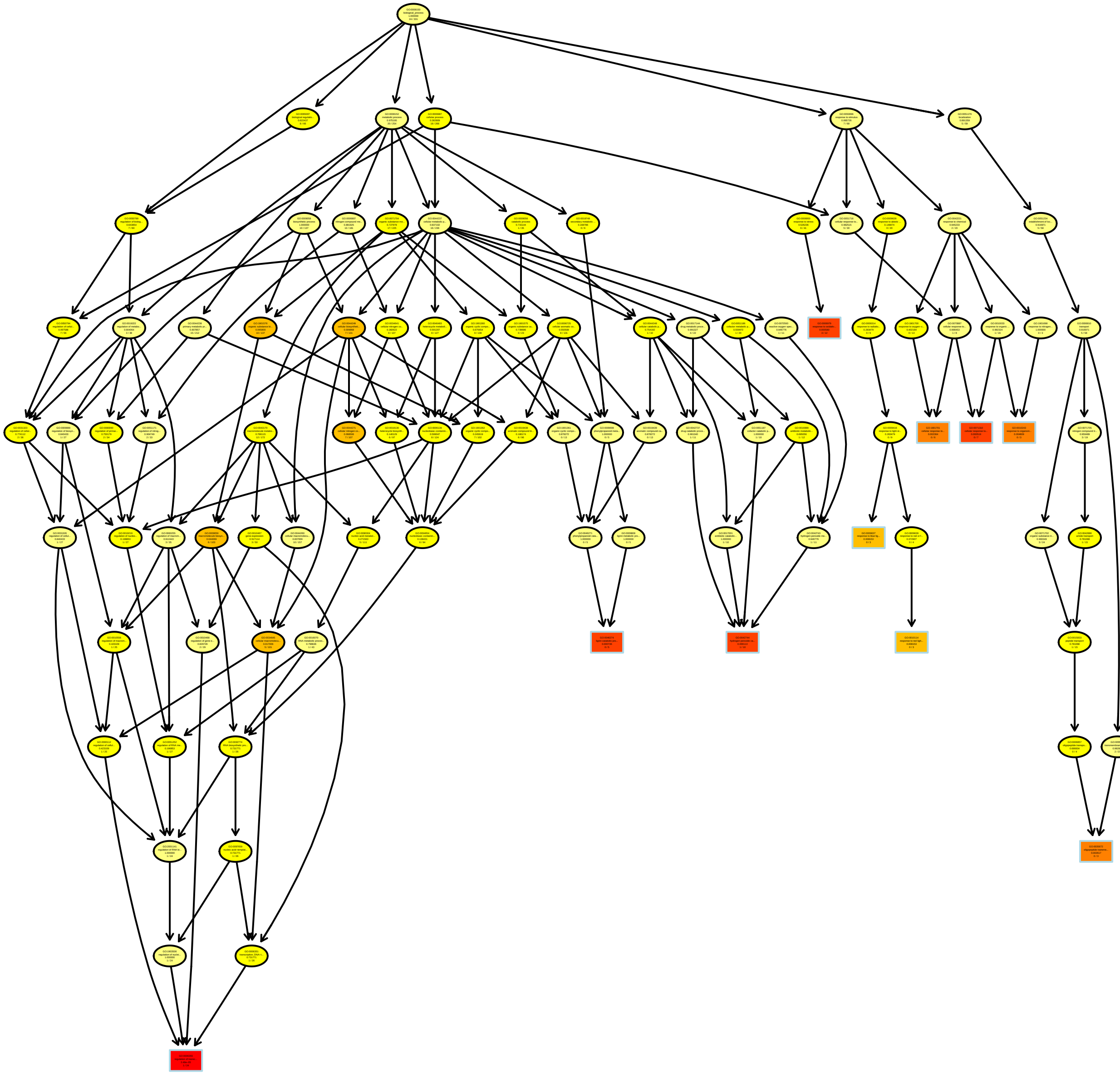

### RelativeRatioRT8.69_BP.pdf

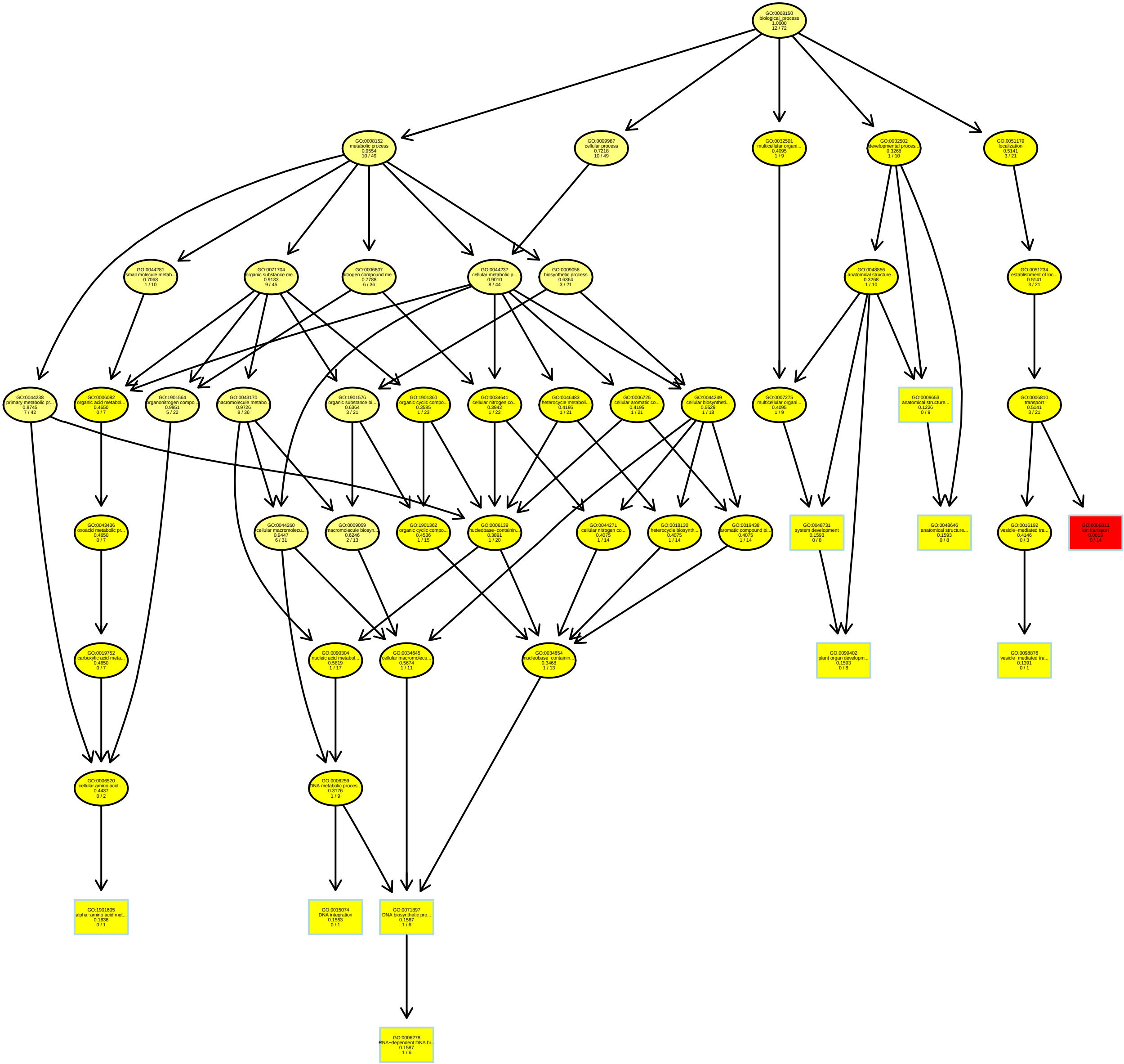

### RelativeRatioRT8.81_MF.pdf

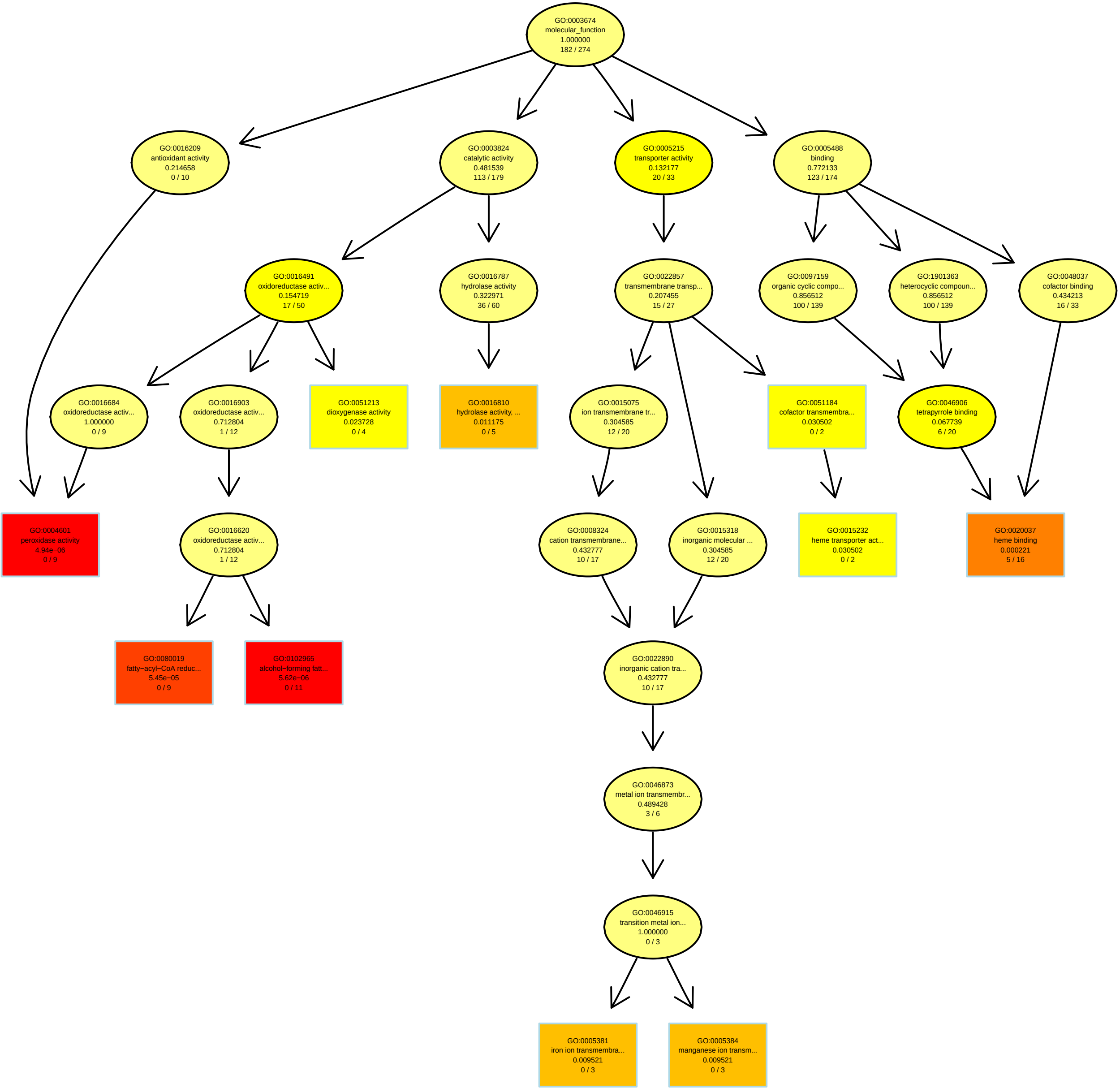

### RelativeRatioRT8.89_MF.pdf

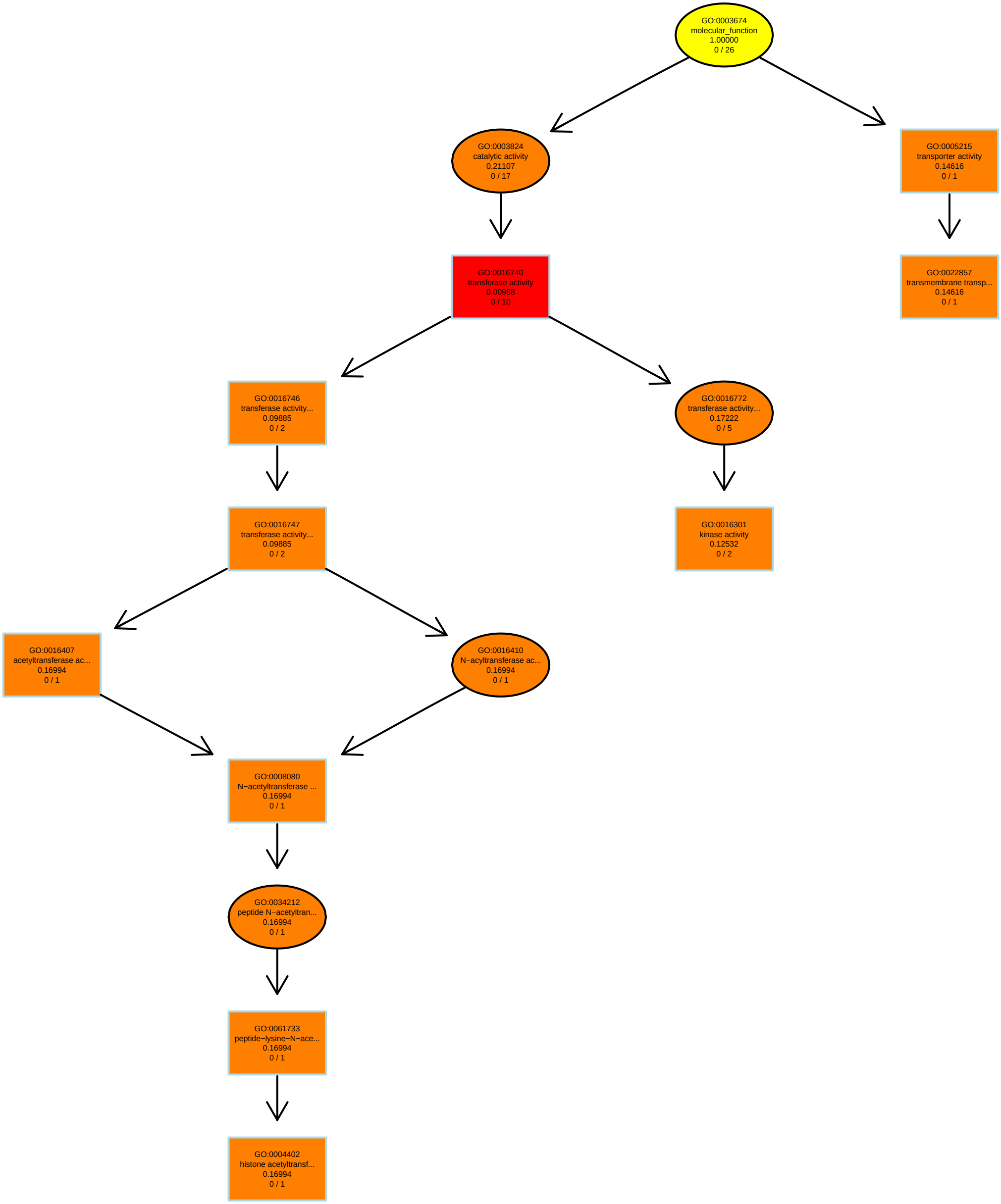

### RelativeRatioRT11.78_MF.pdf

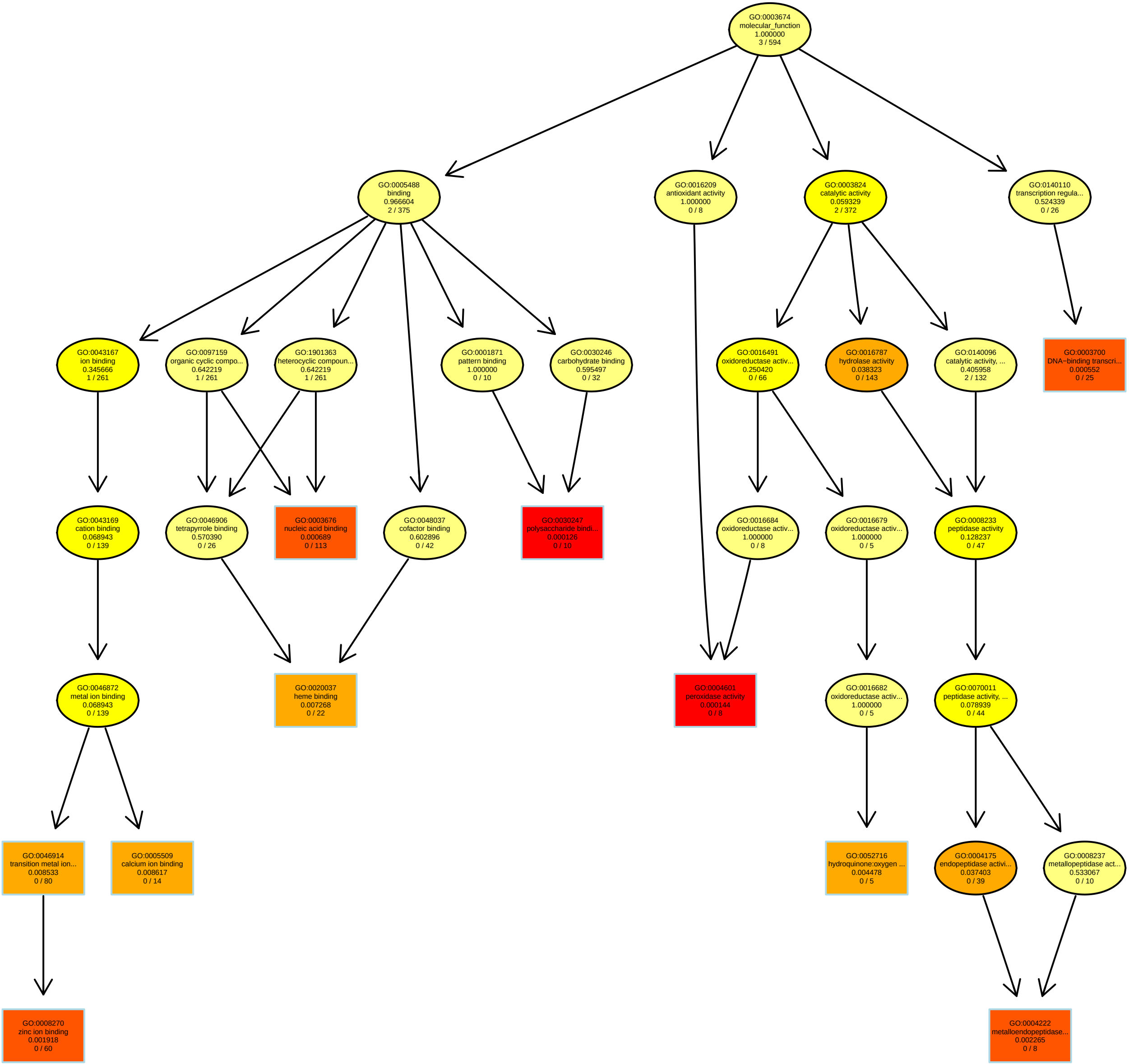

### RelativeRatios200PC1_MF.pdf

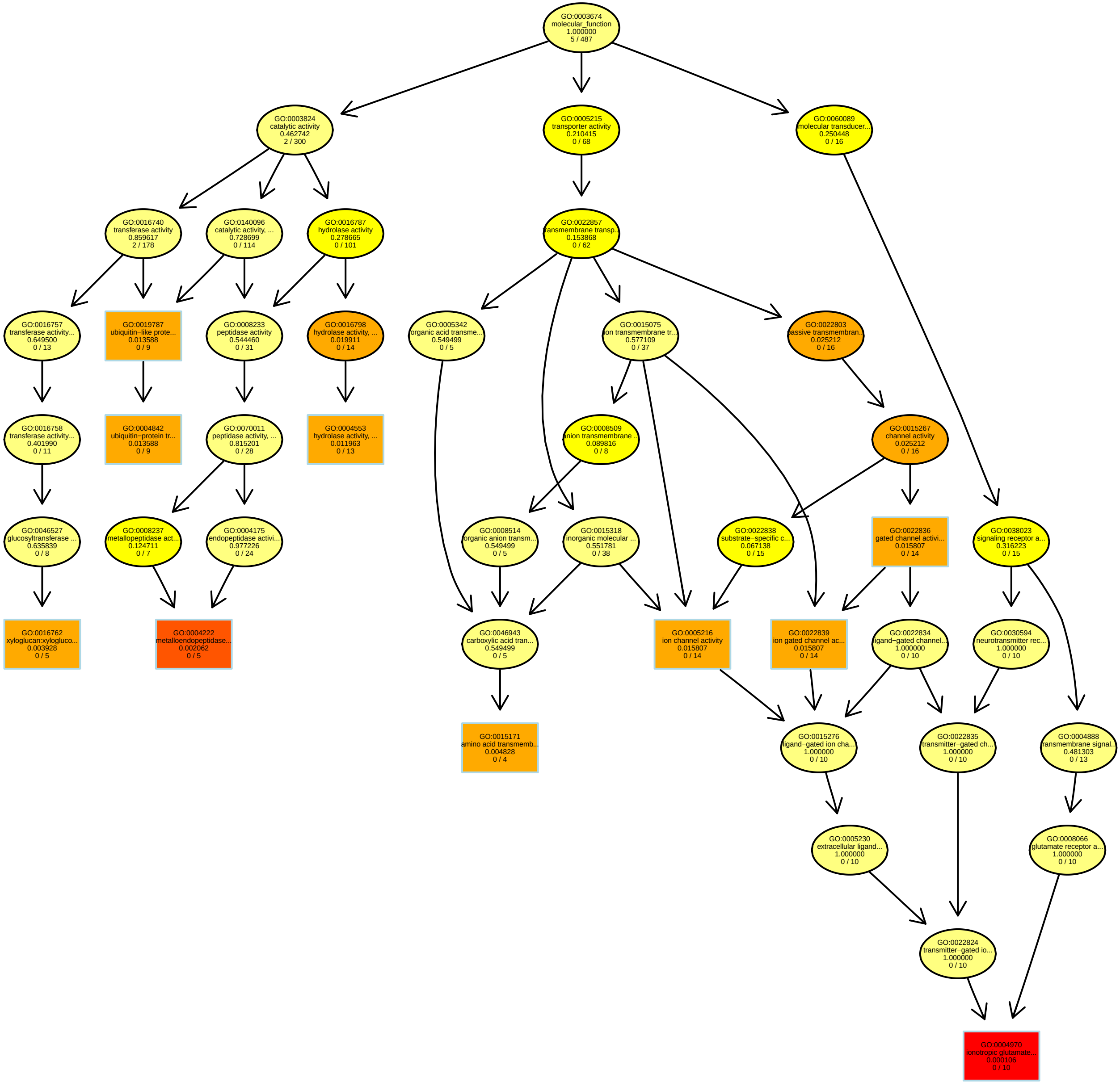

### RelativeRatios200PC2_BP.pdf

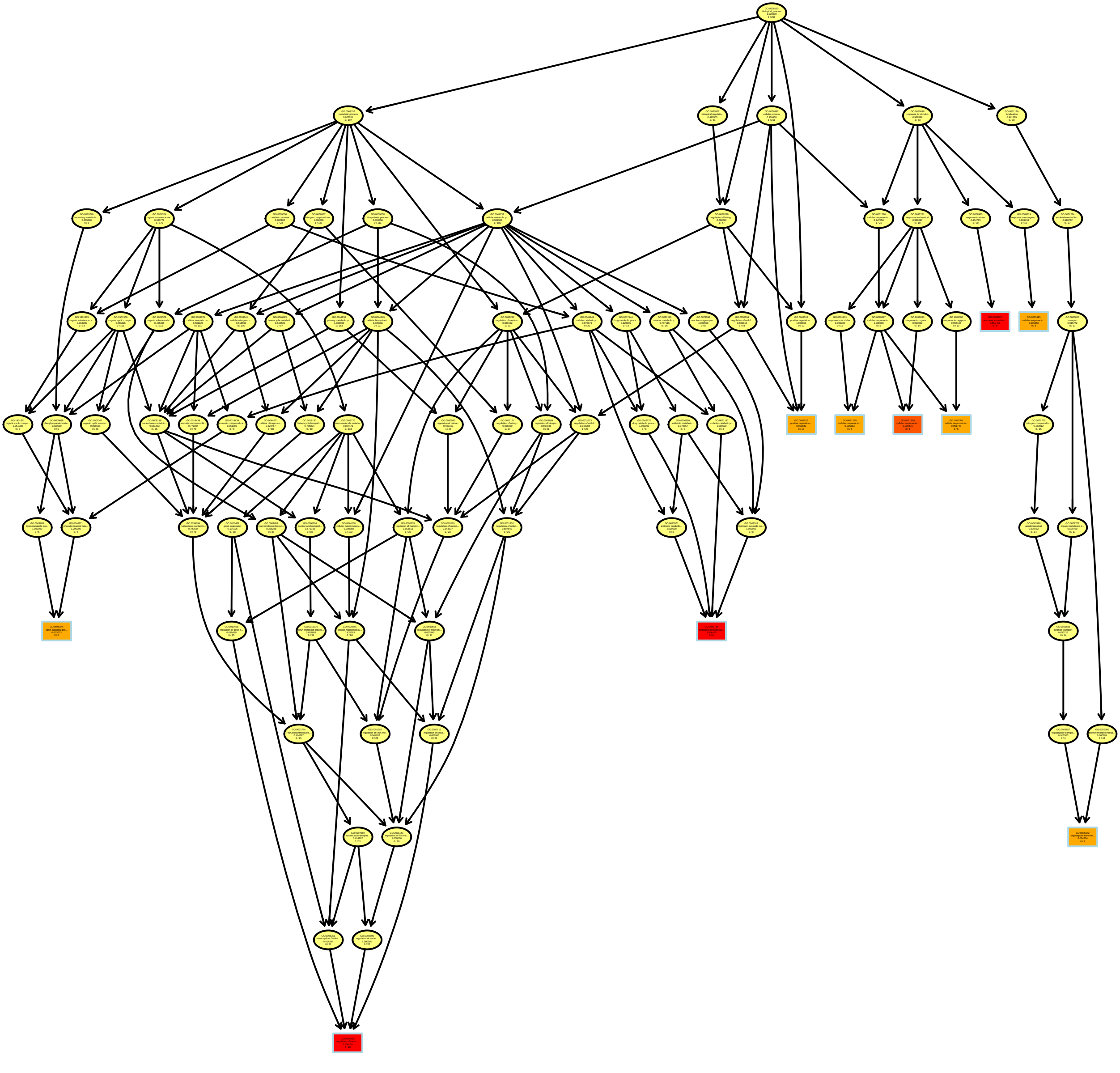

### RelativeRatios200PC3_MF.pdf

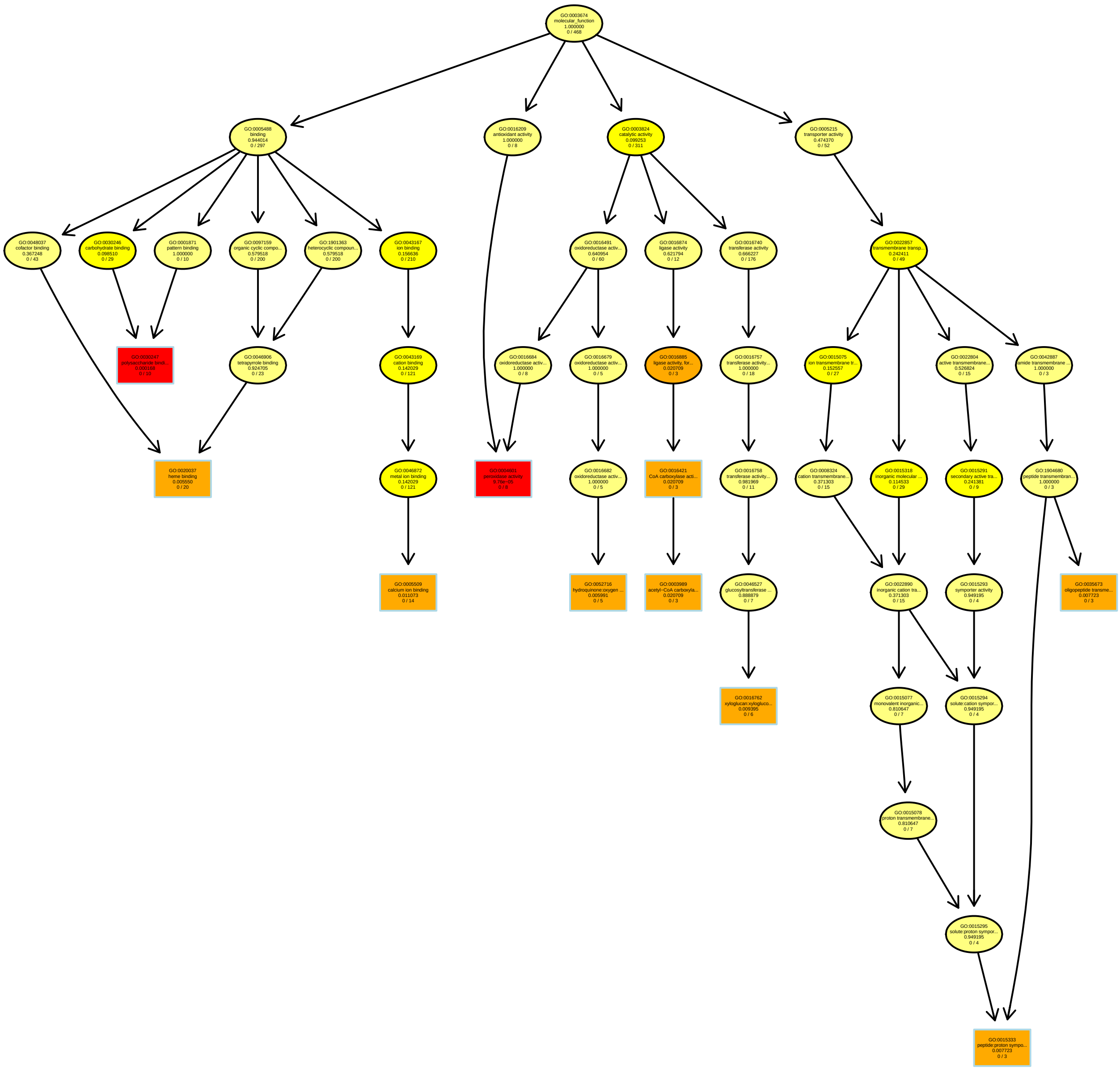

### RelativeRatios200PC4_BP.pdf

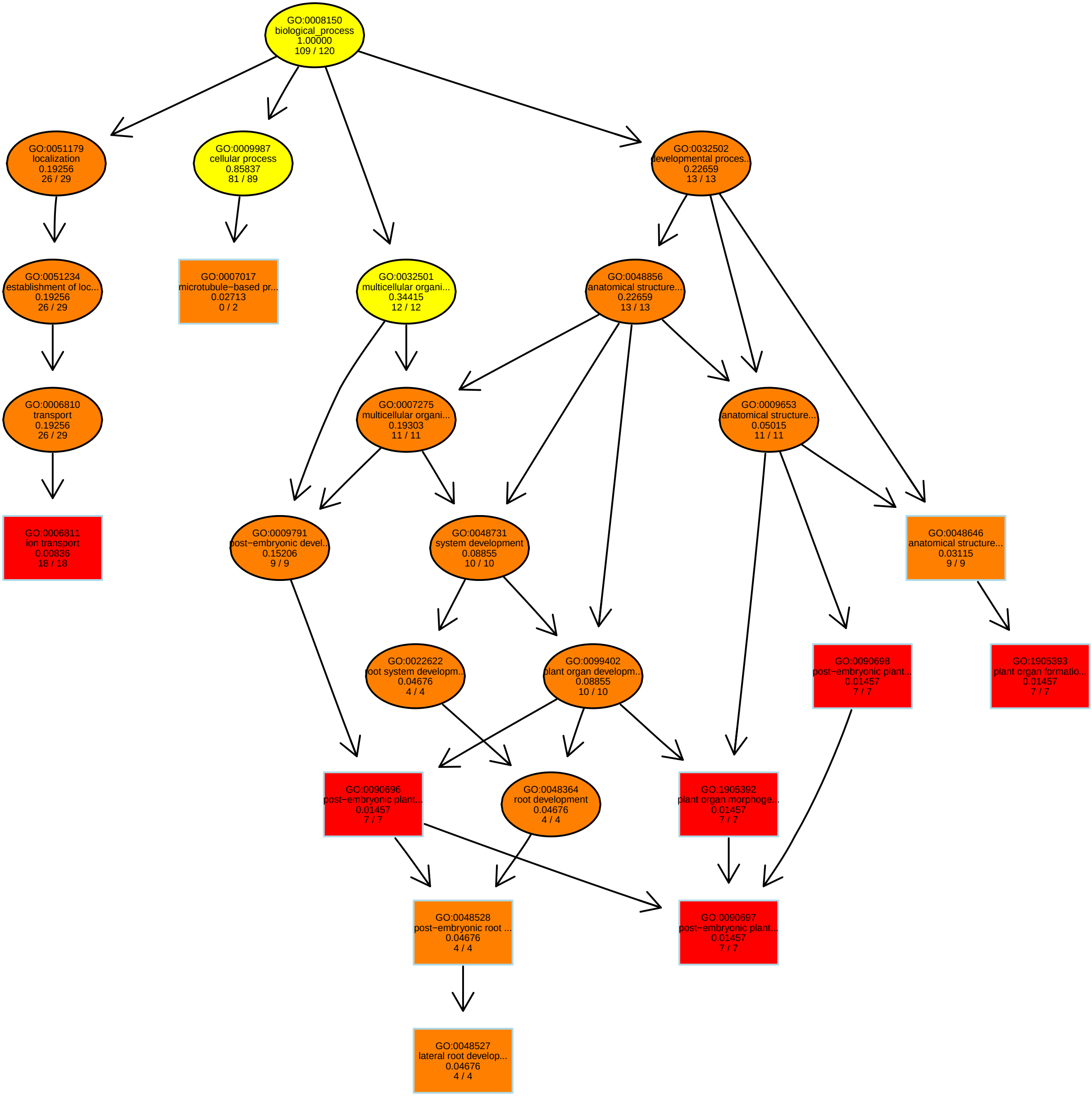

### RelativeRatios200PC6_BP.pdf

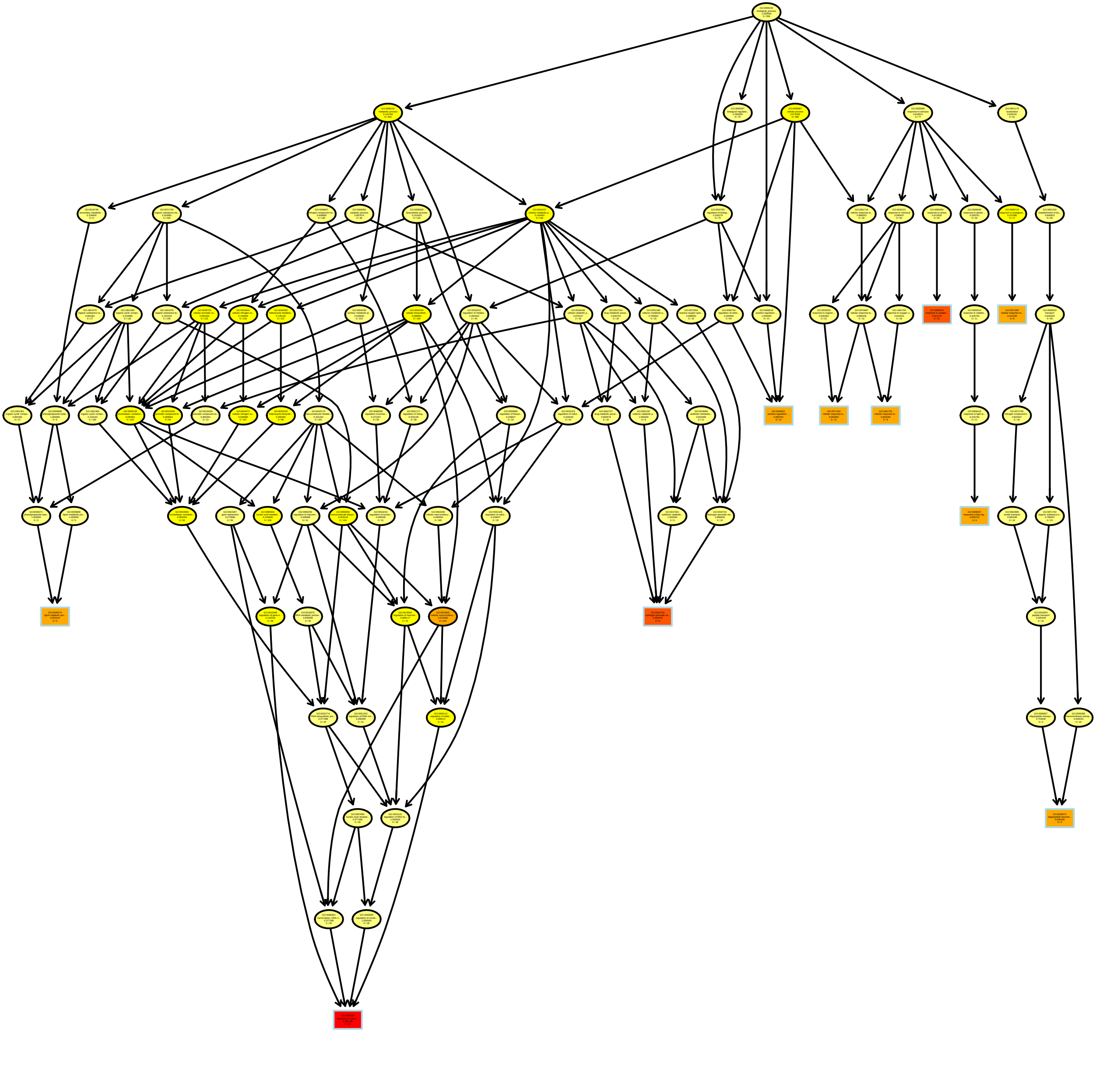

### RelativeRatios200PC7_MF.pdf

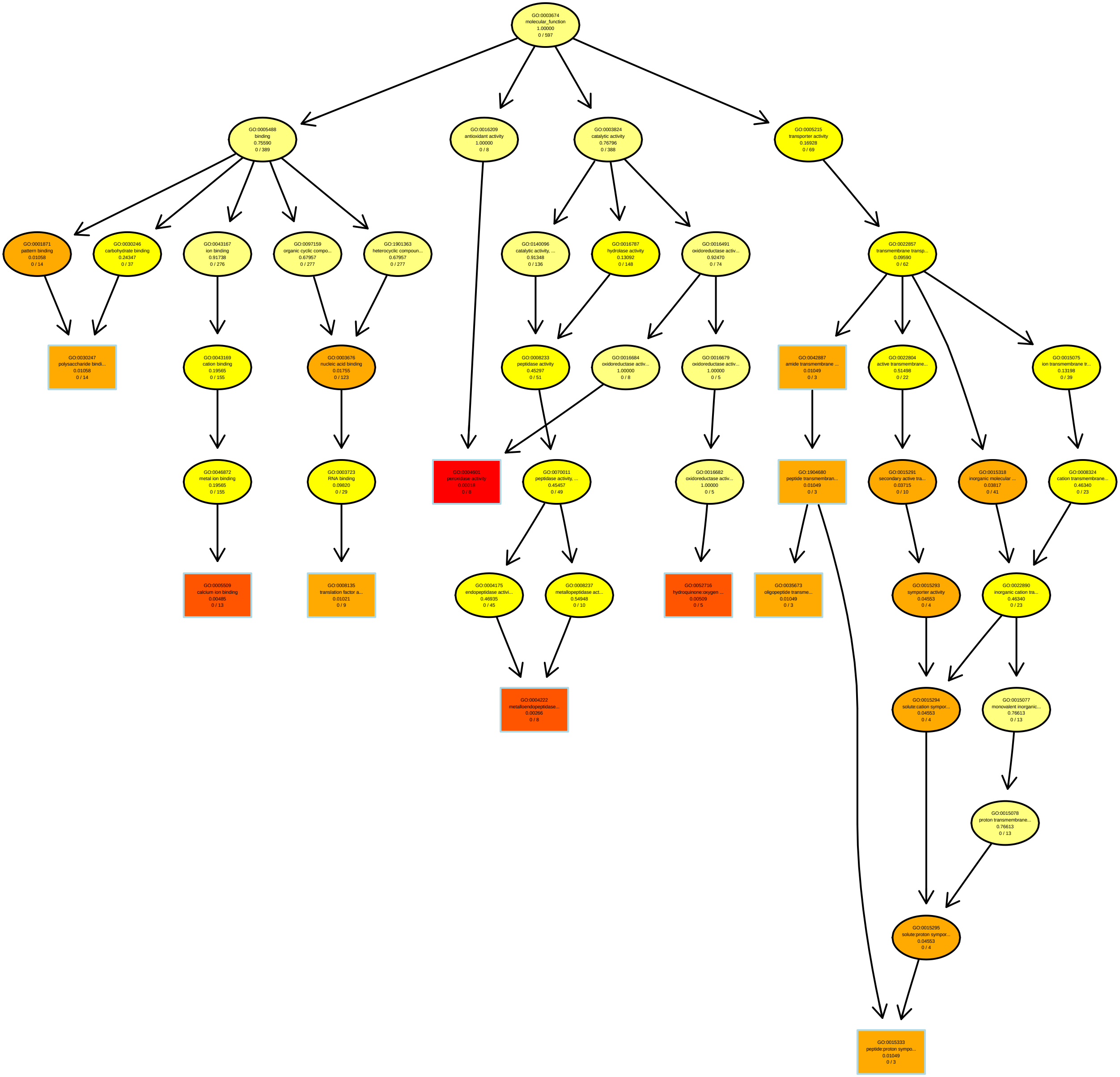

### RelativeRatios200PC8_BP.pdf

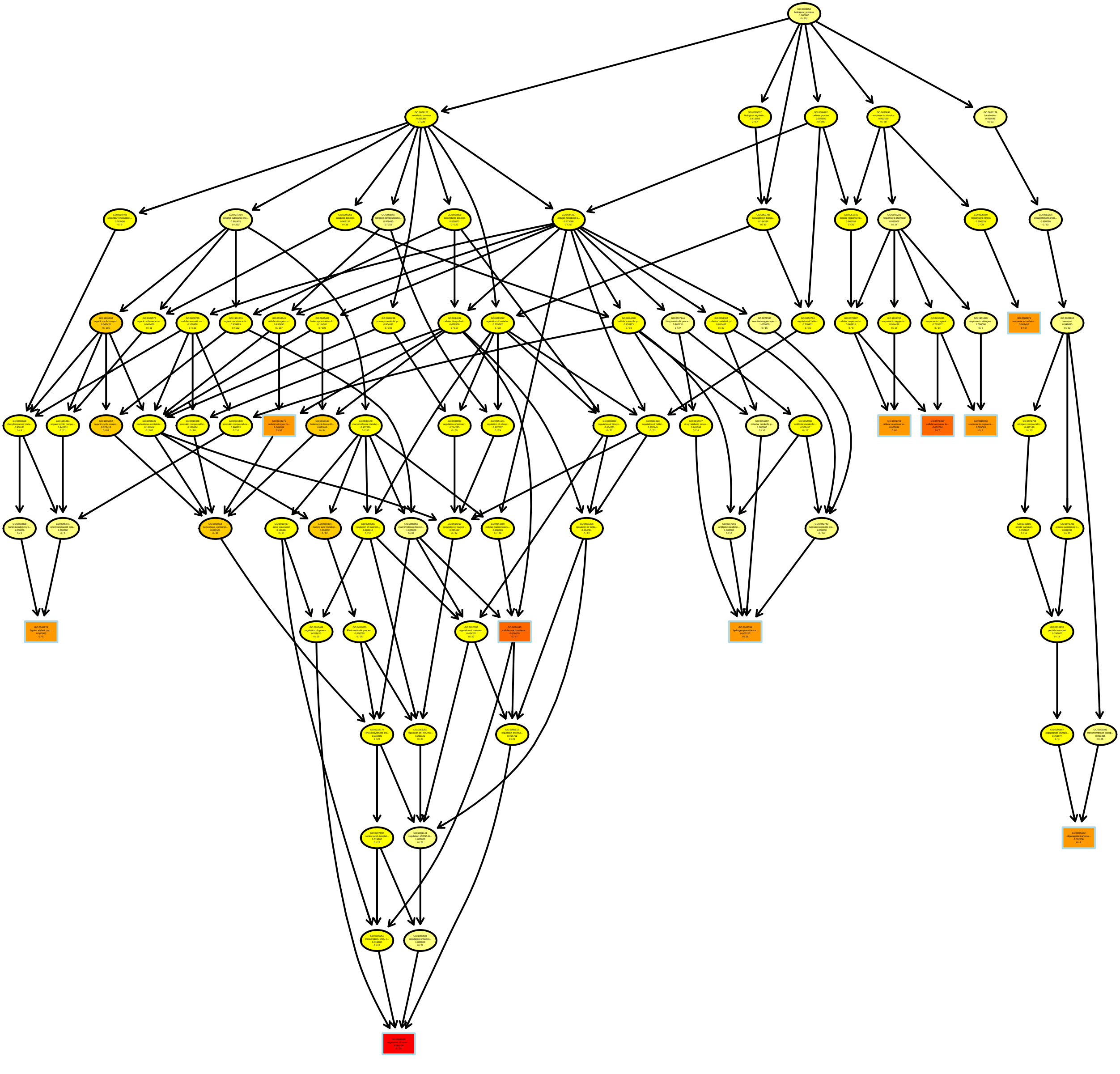

### RelativeRatios200PC10_MF.pdf

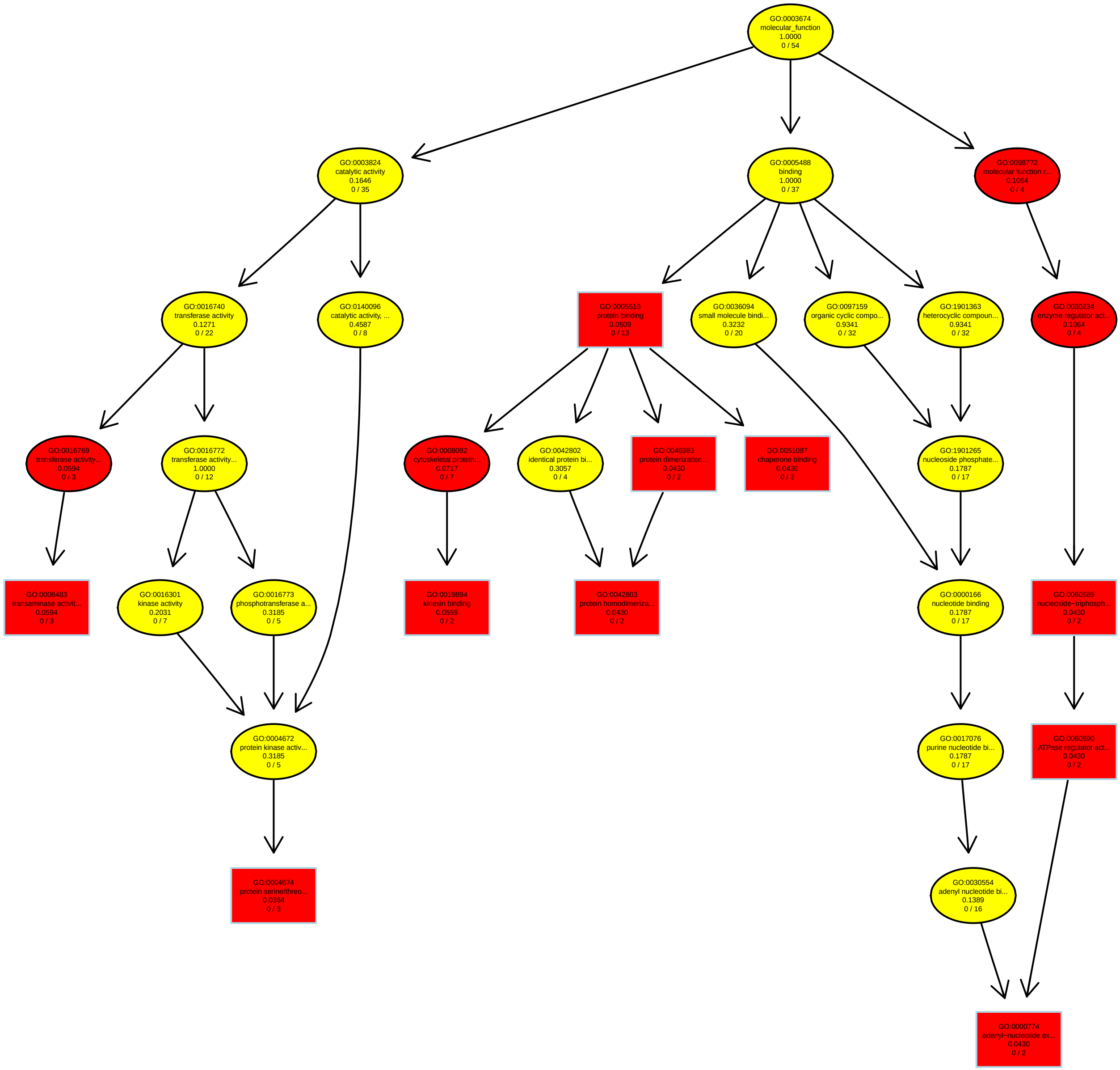

### RelativeRatiosPC2_MF.pdf

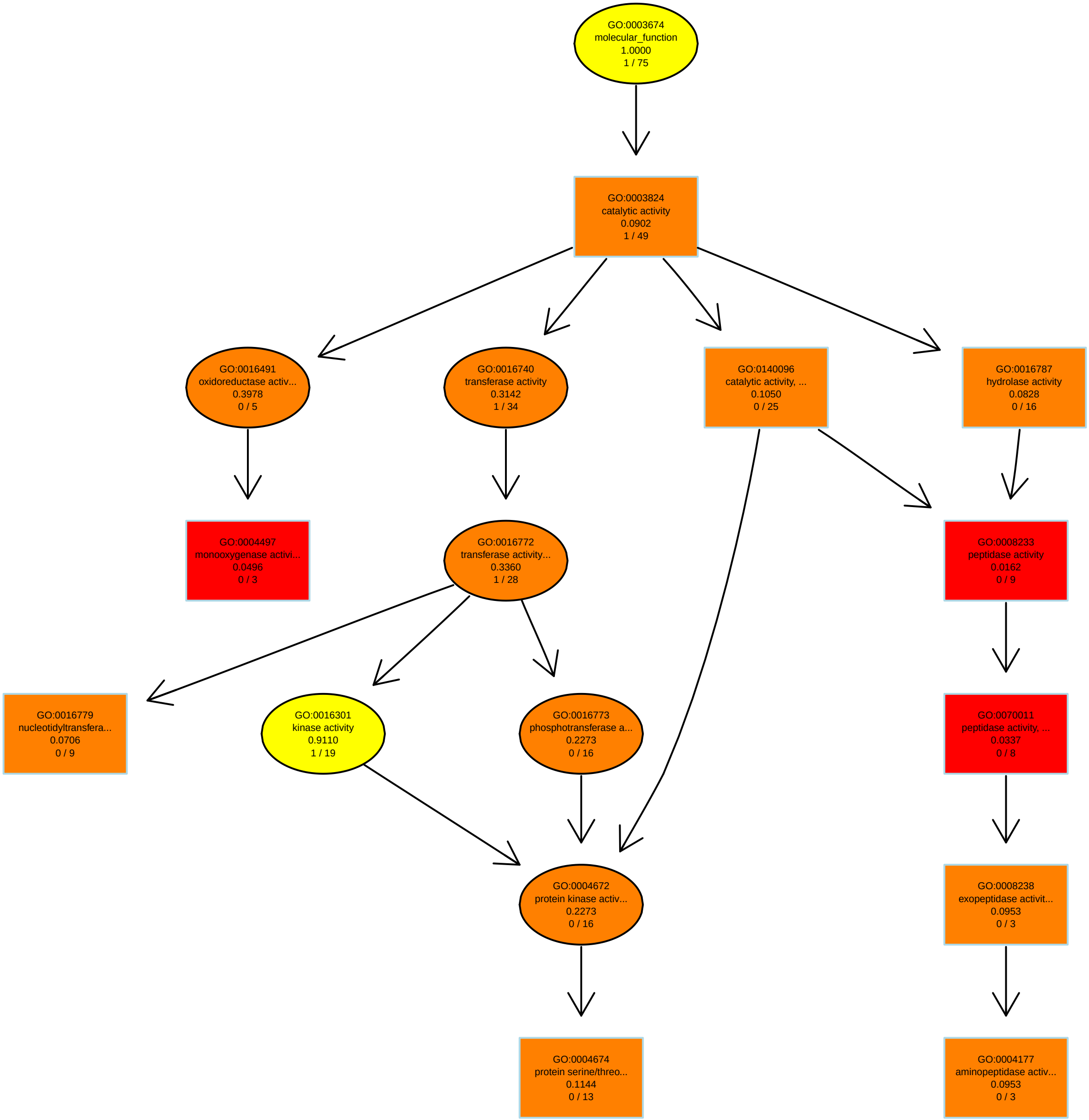

### RelativeRatiosPC3_BP.pdf

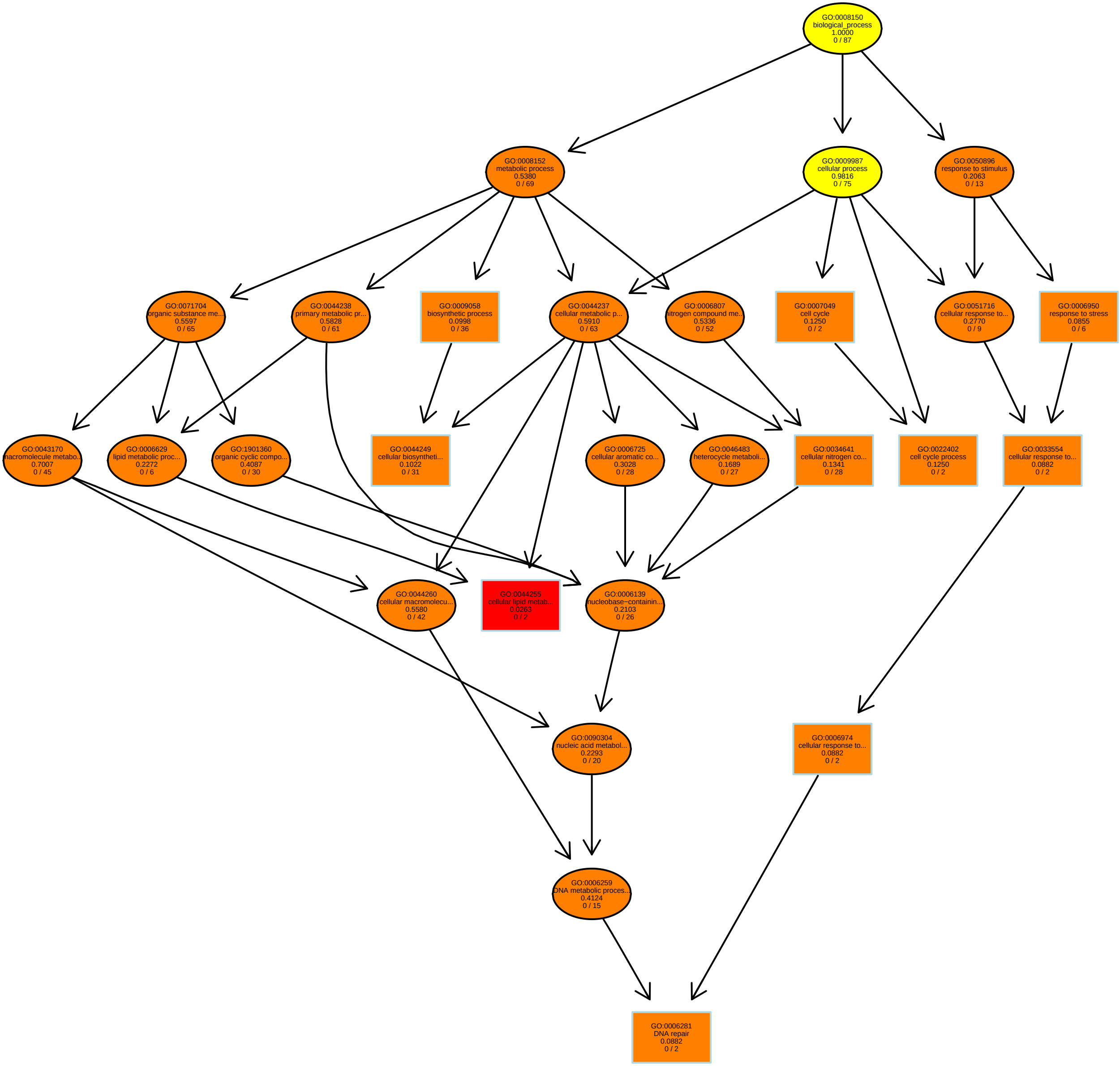

### RelativeRatiosPC5_BP.pdf

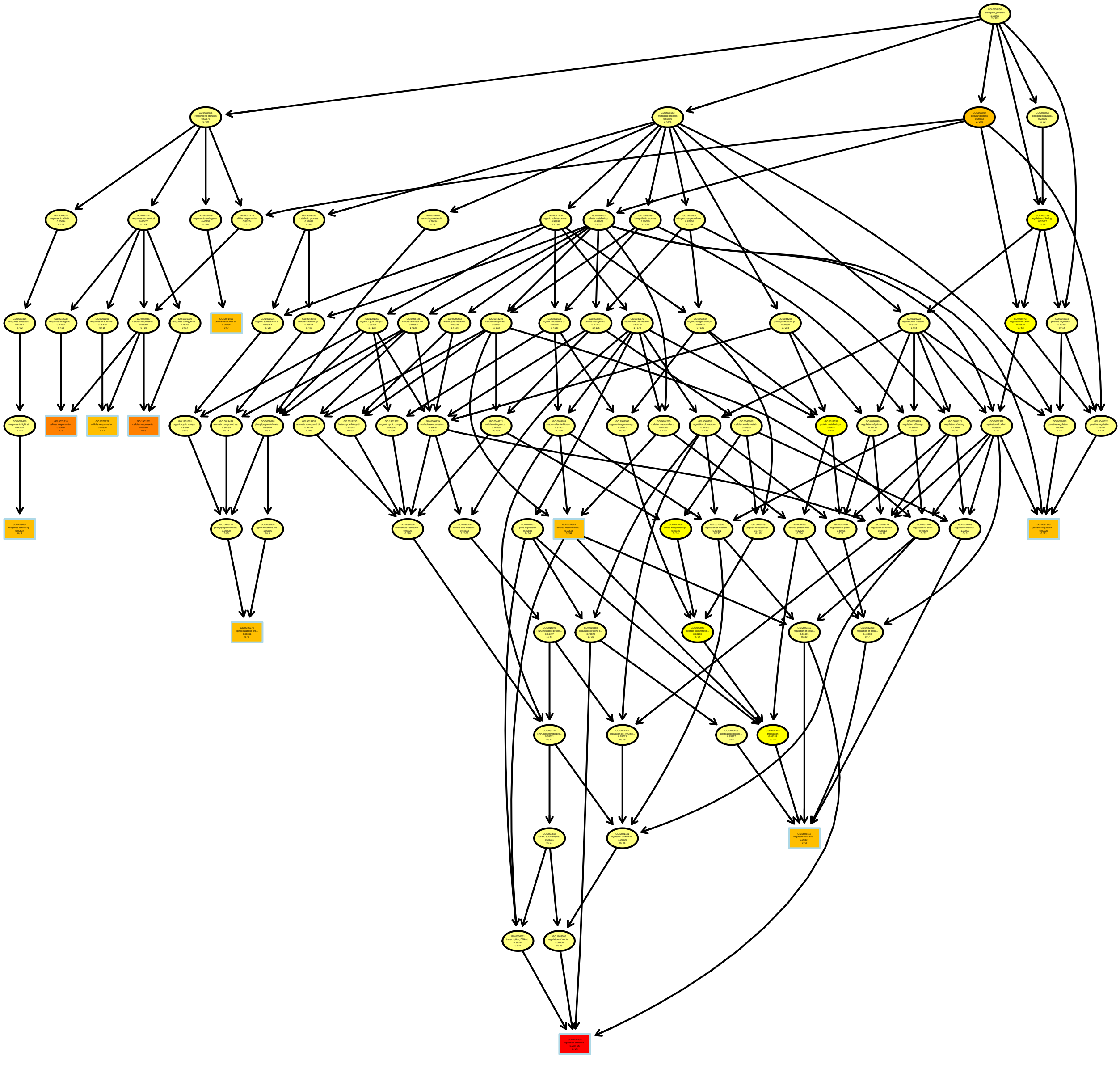

### RelativeRatiosPC6_MF.pdf

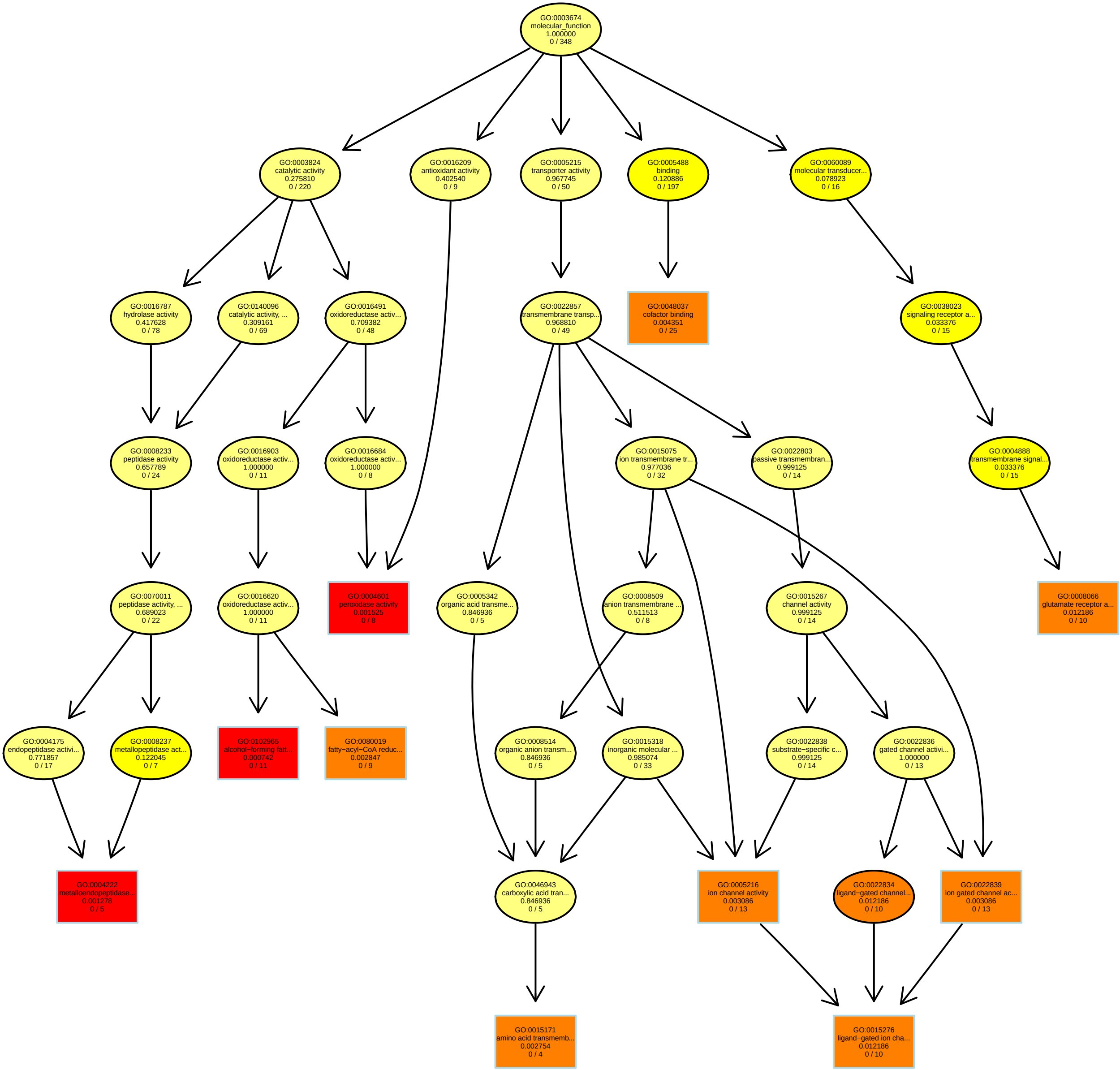

### RelativeRatiosPC7_BP.pdf

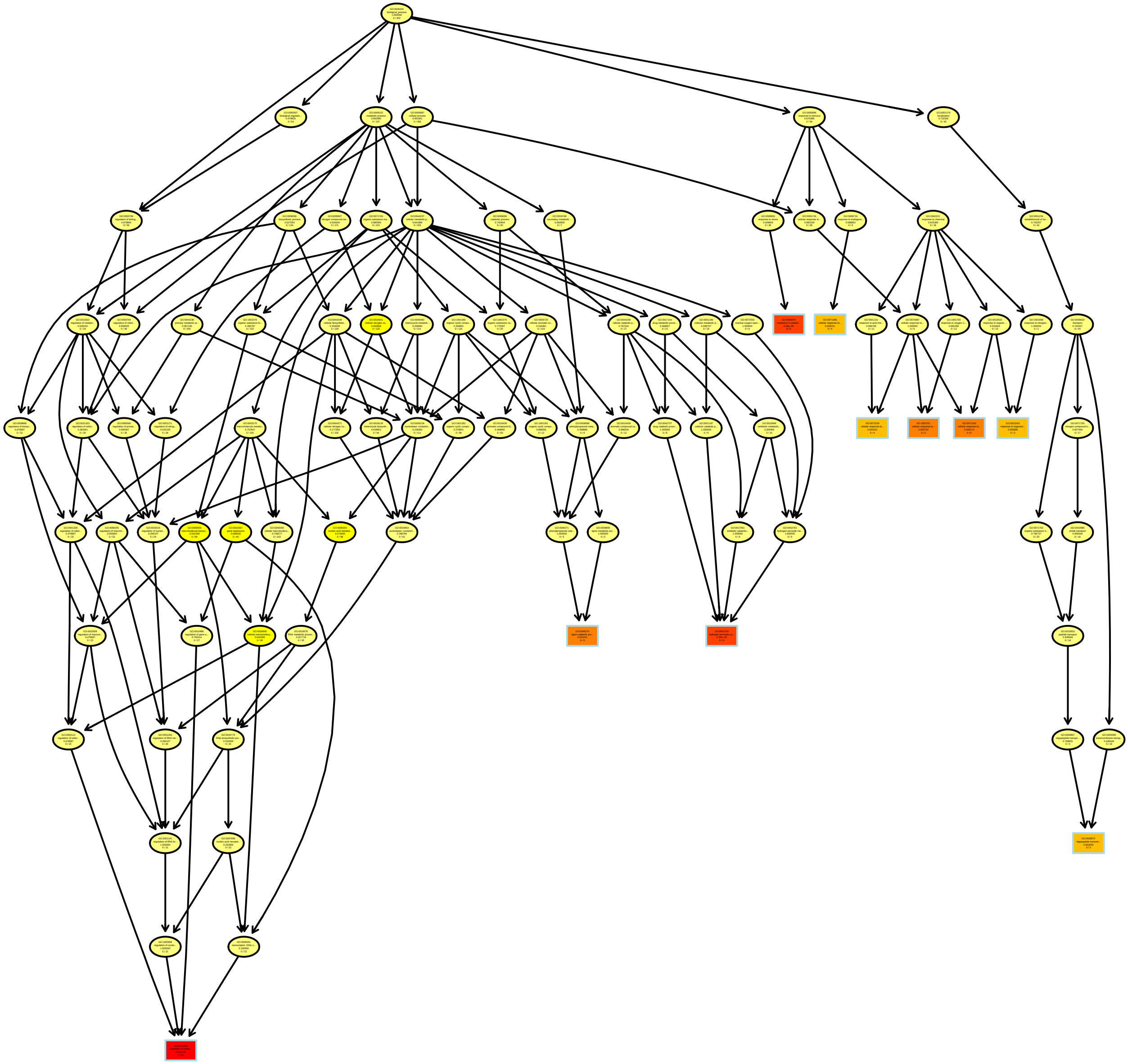

### RelativeRatiosPC9_BP.pdf

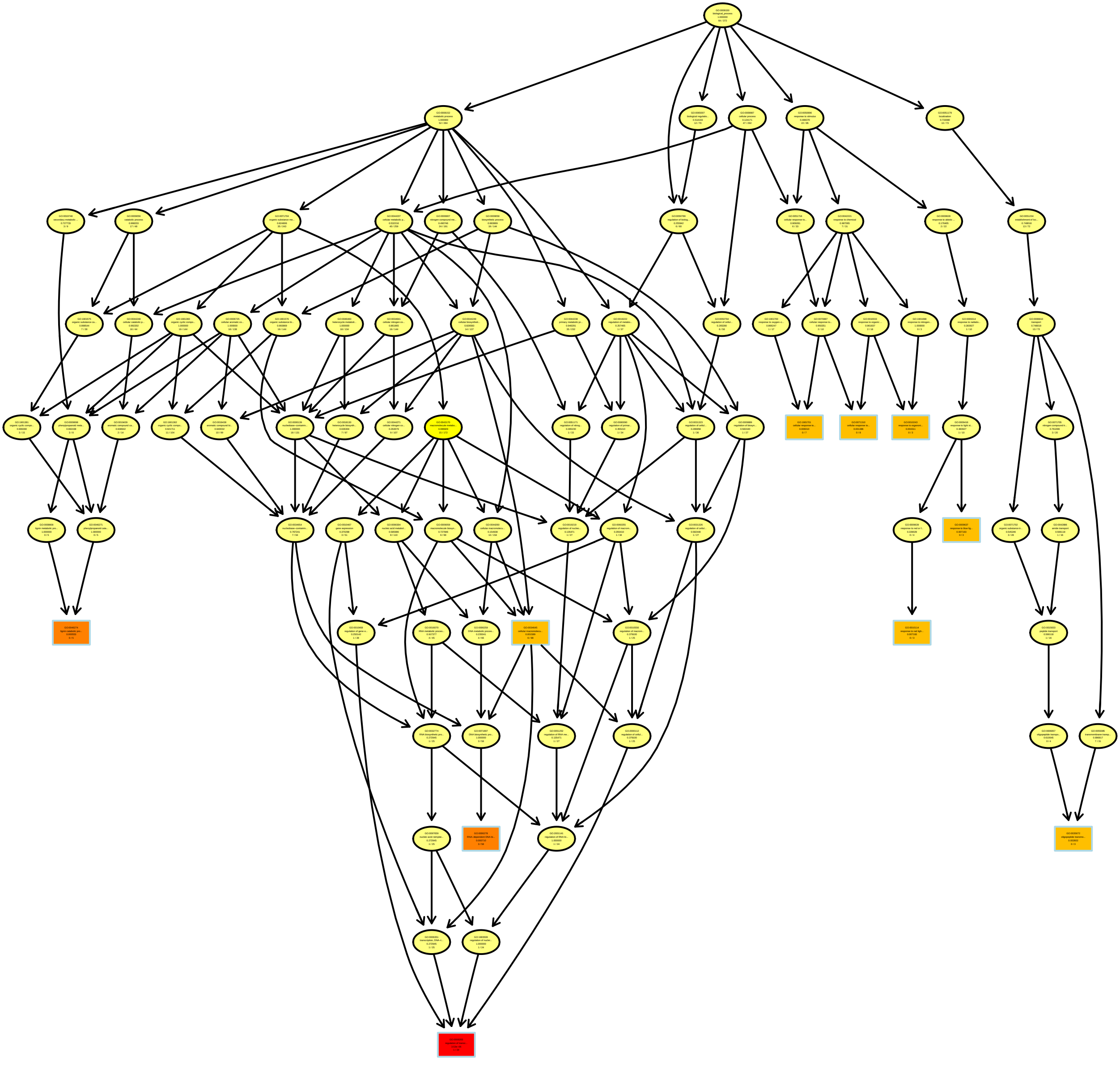

### RelativeRatiosPC10_MF.pdf

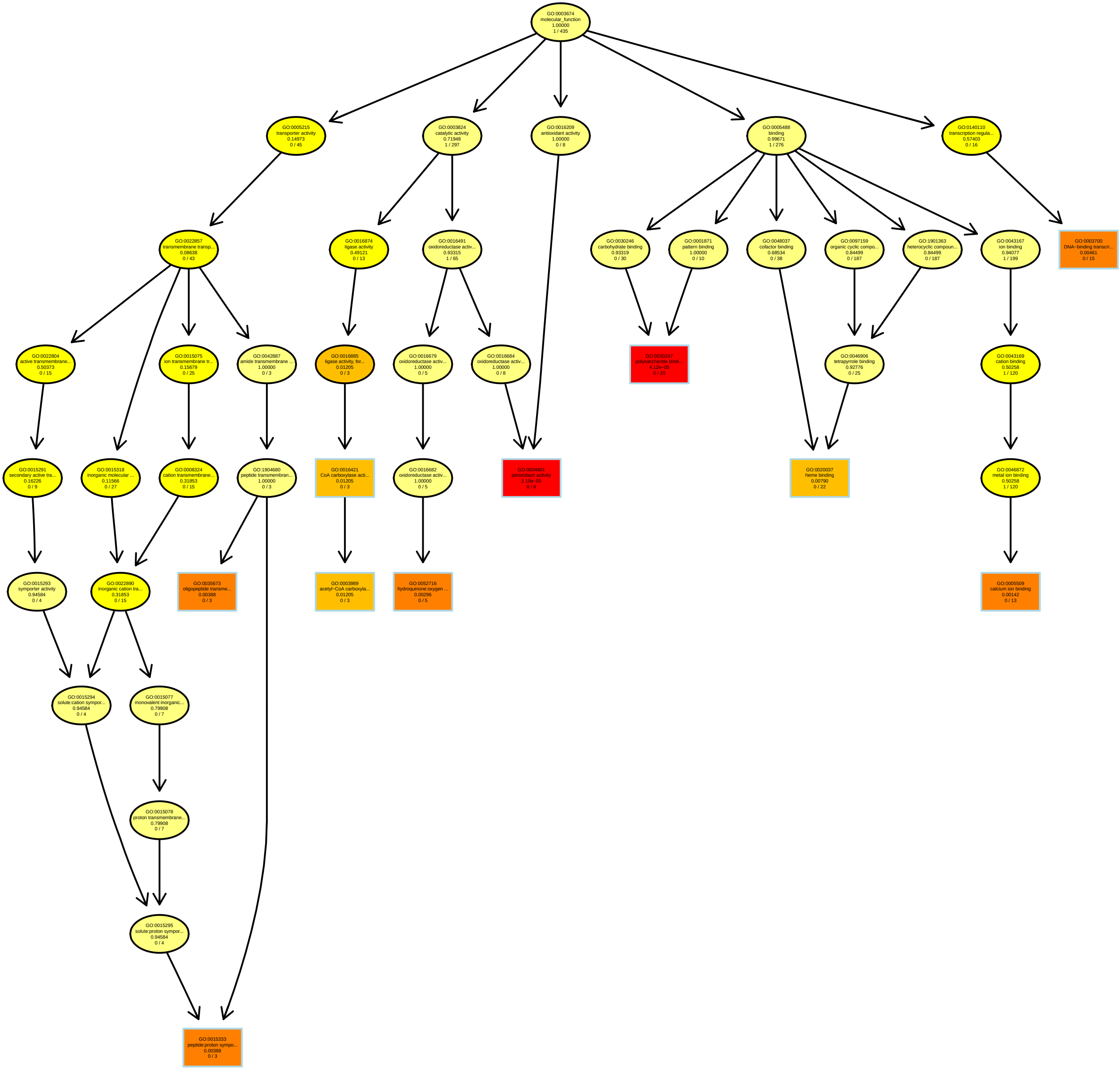

### RT1.2_BP.pdf

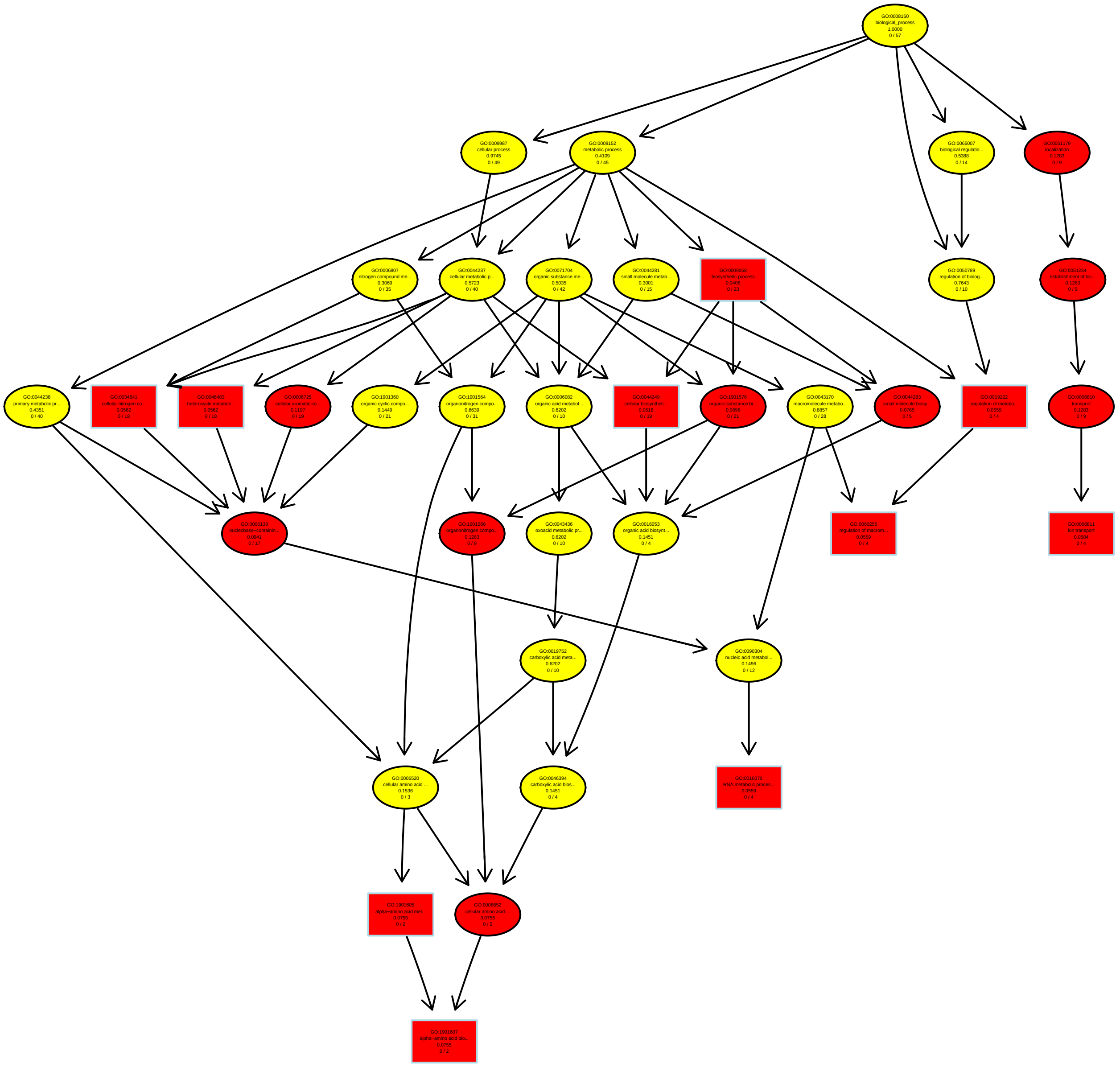

### RT3.3_BP.pdf

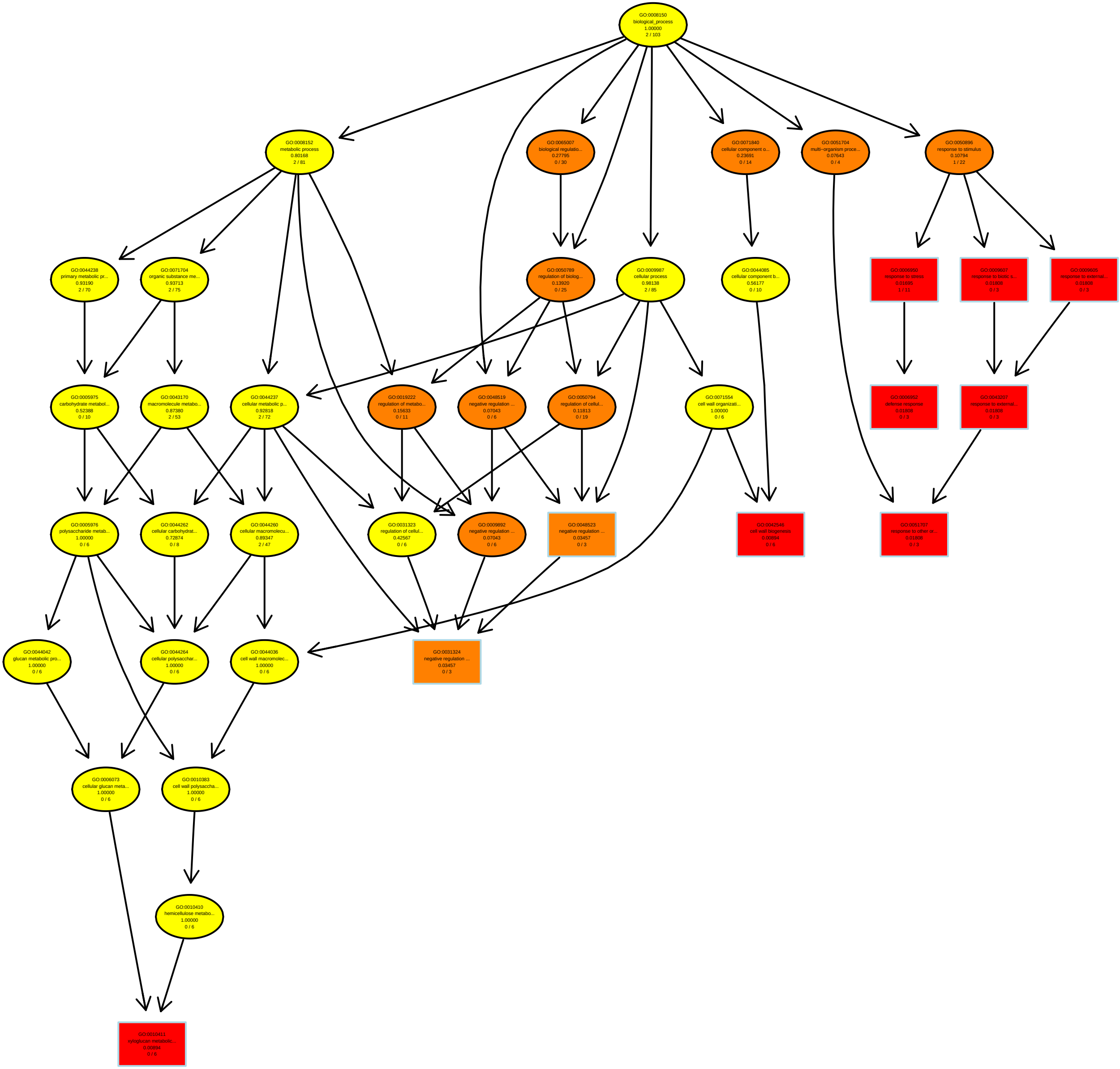

### RT3.52_MF.pdf

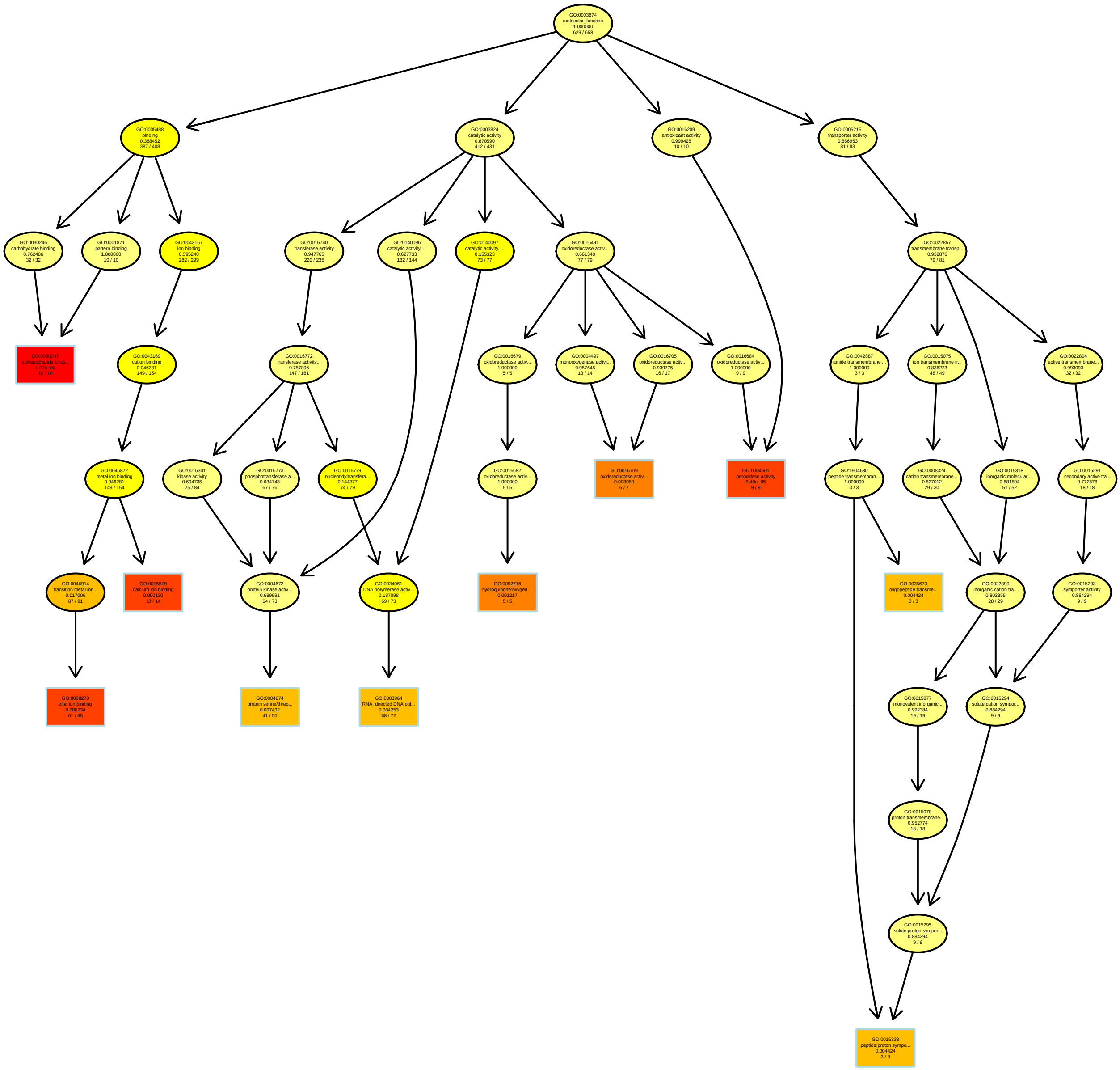

### RT3.77_BP.pdf

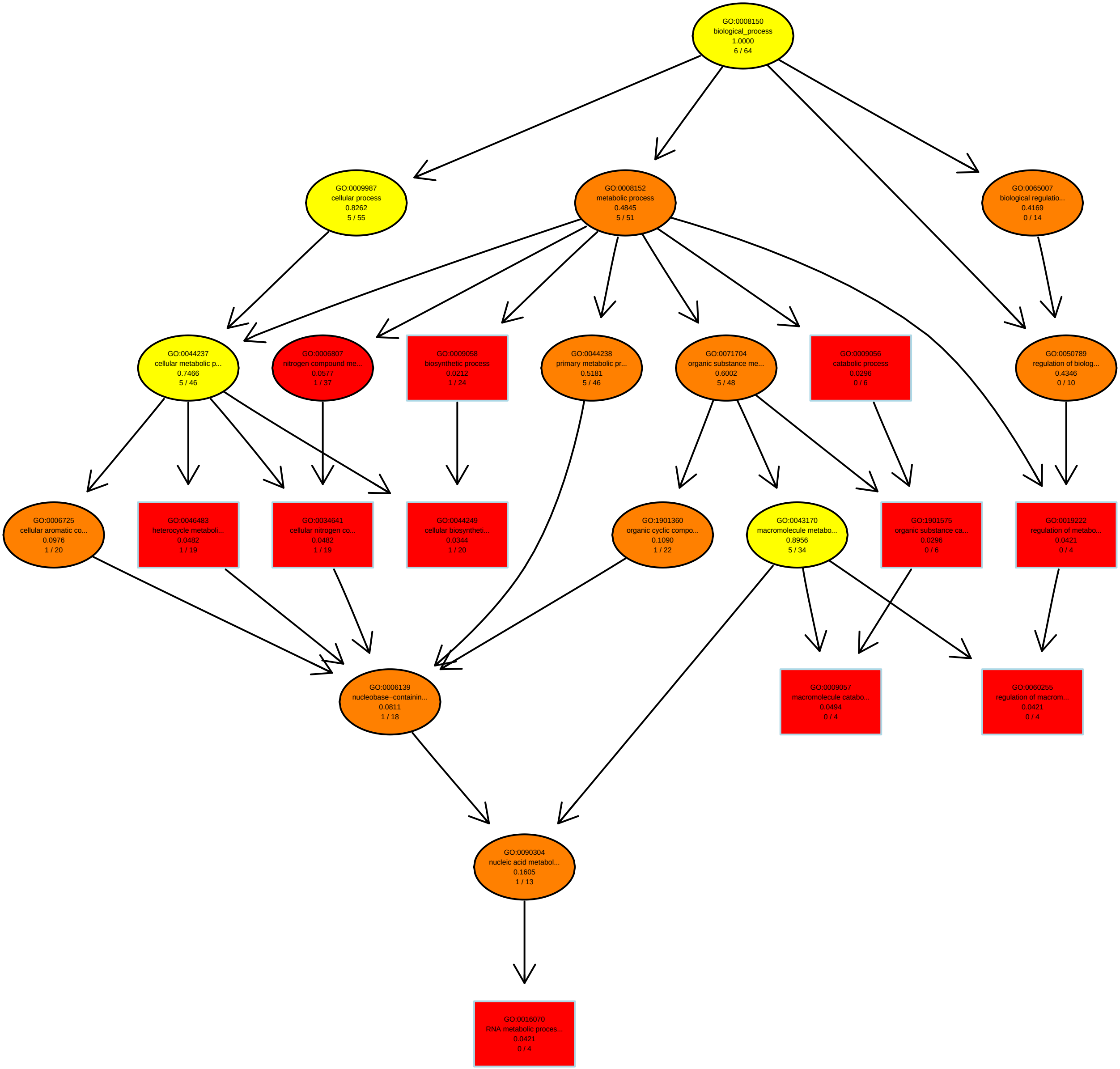

### RT4.6_MF.pdf

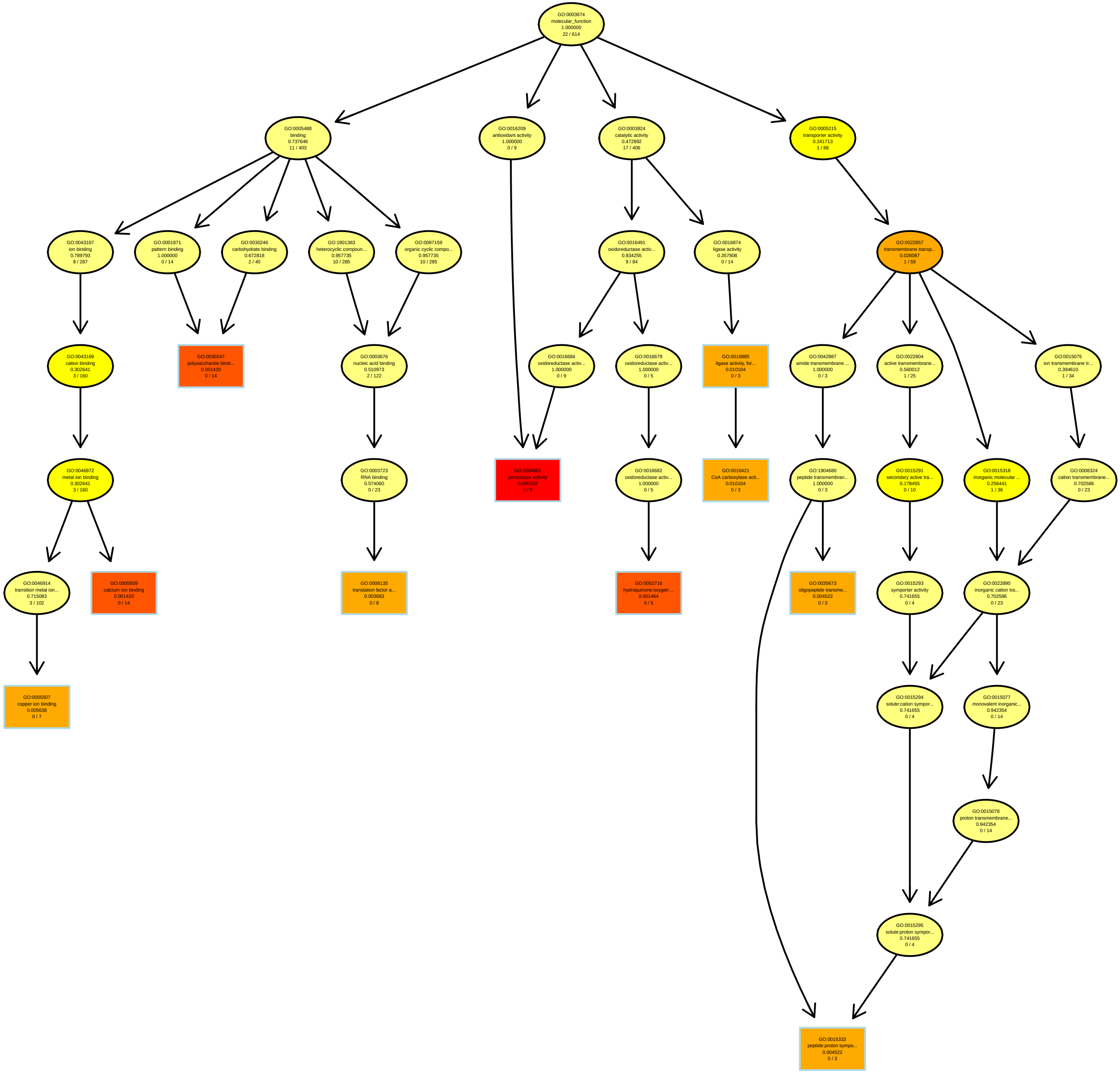

### RT4.98_BP.pdf

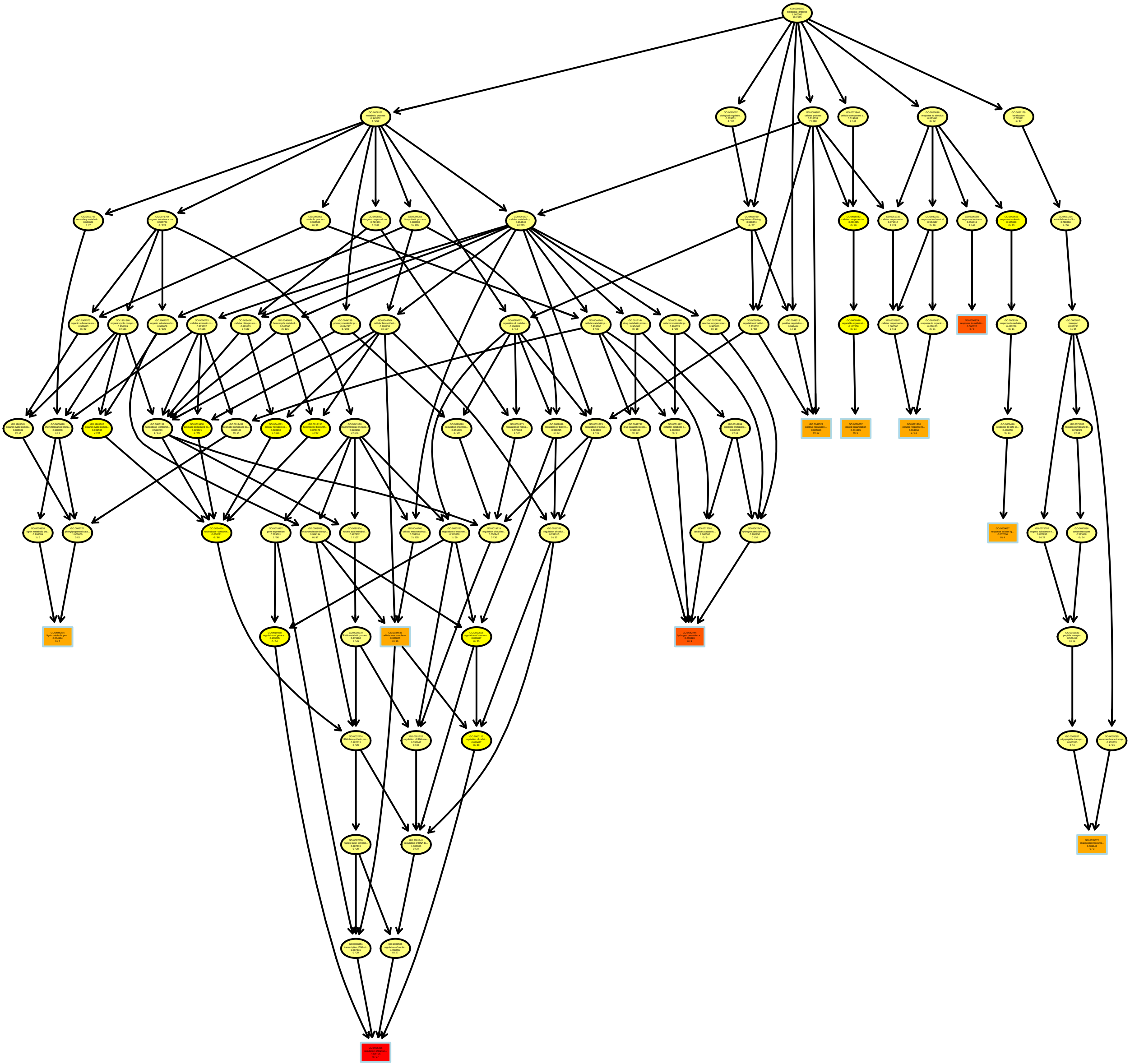

### RT5.28_BP.pdf

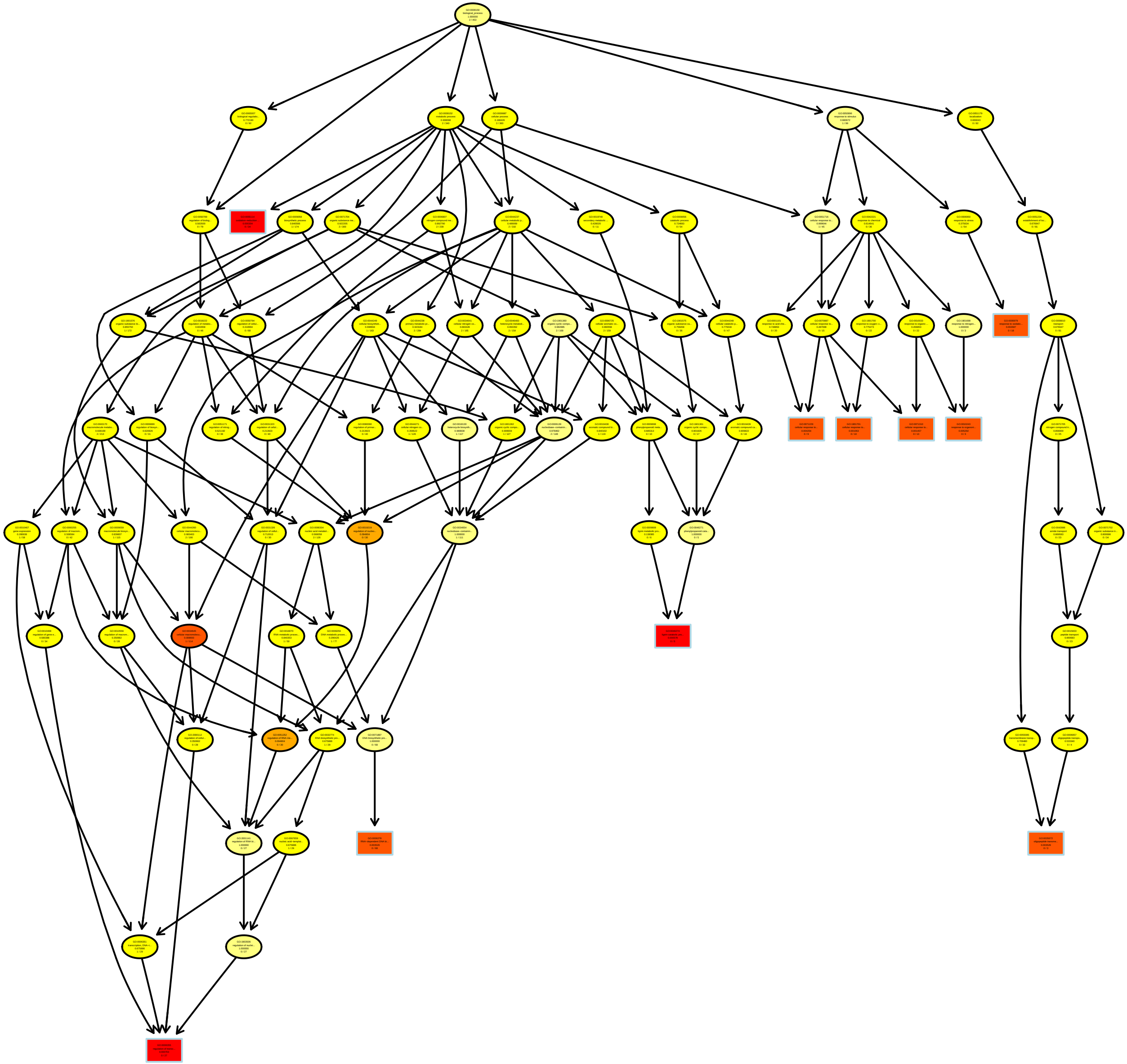

### RT5.89_MF.pdf

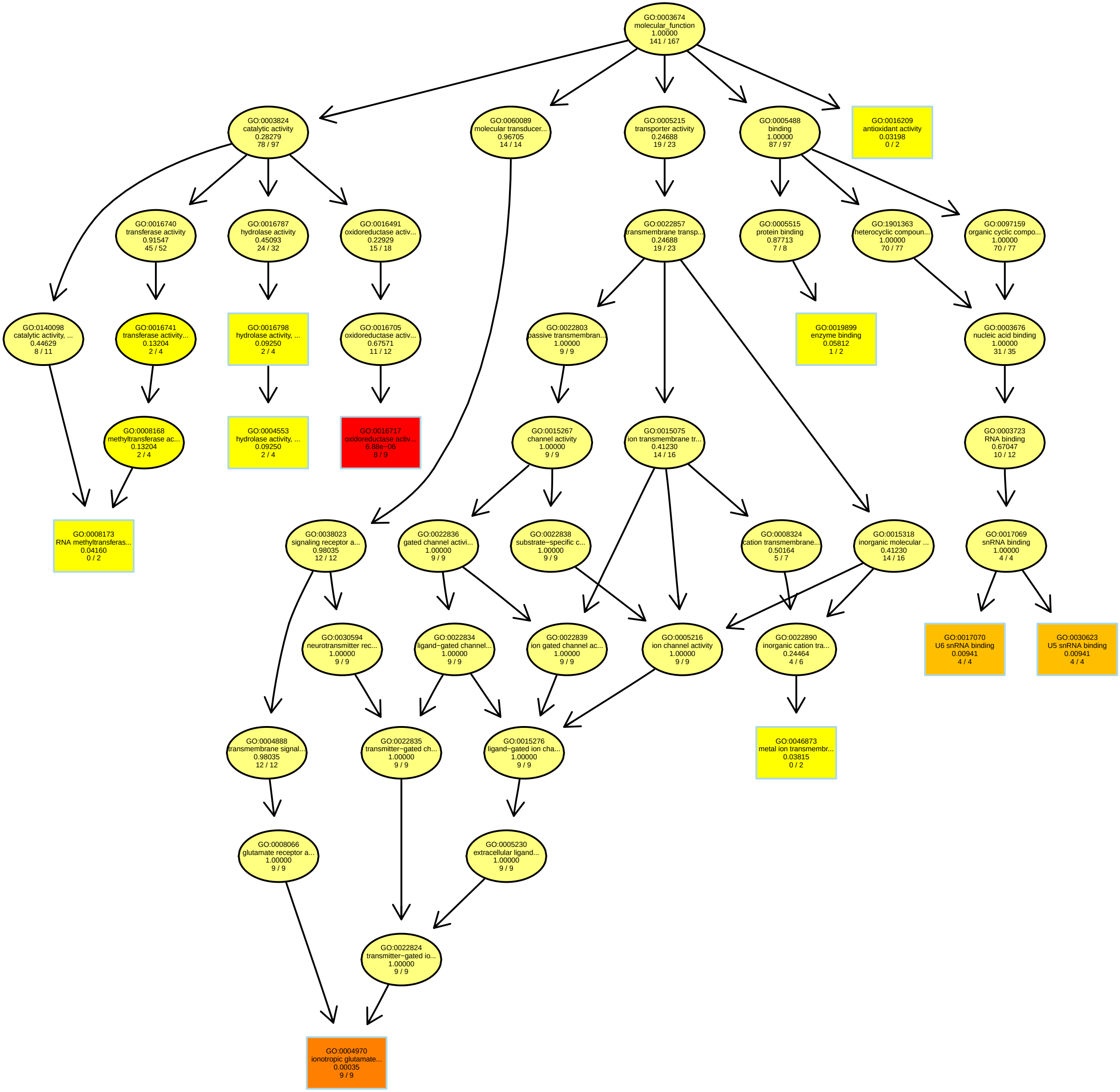

### RT7.1_MF.pdf

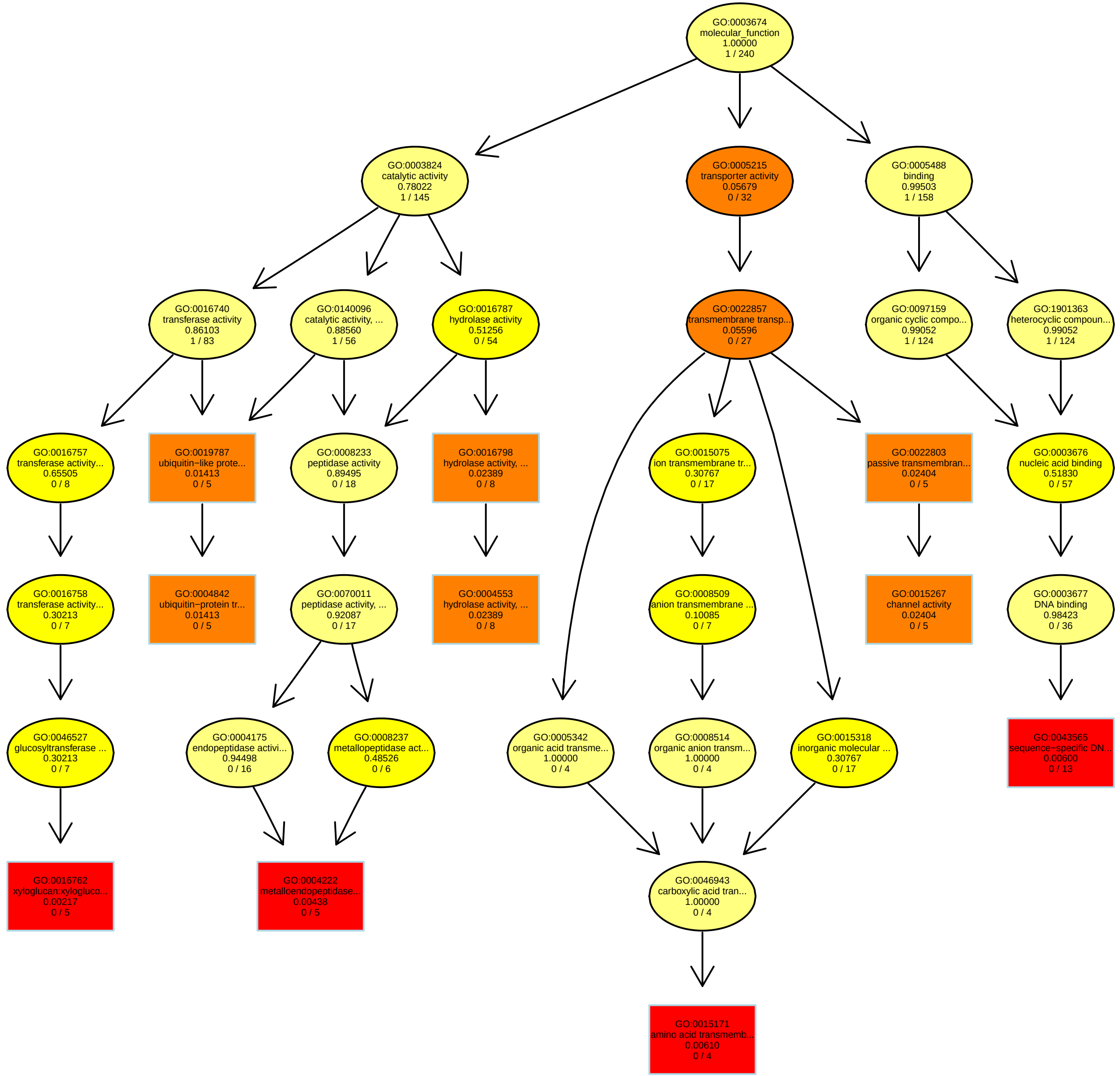

### RT7.28_MF.pdf

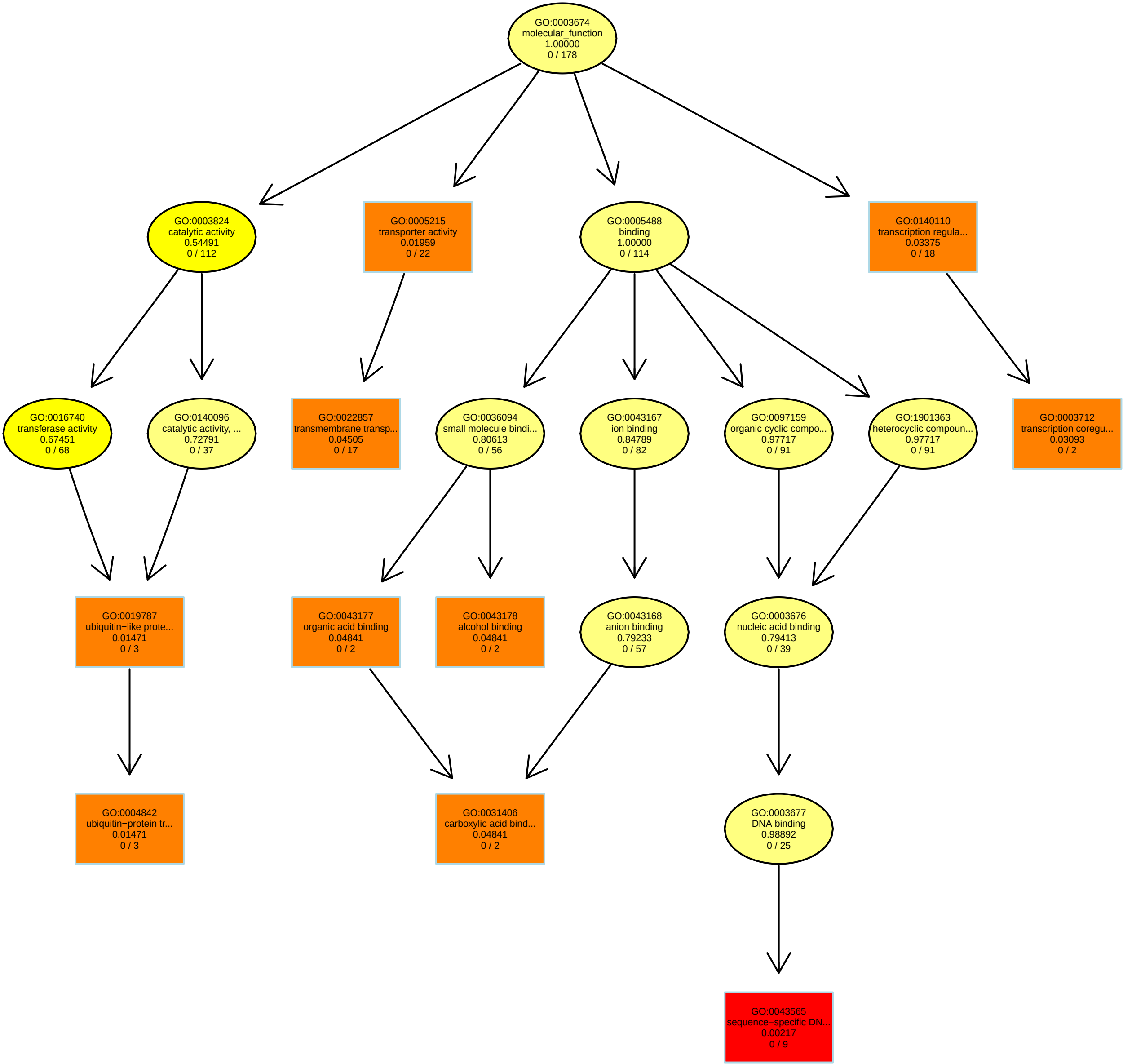

### RT7.47_MF.pdf

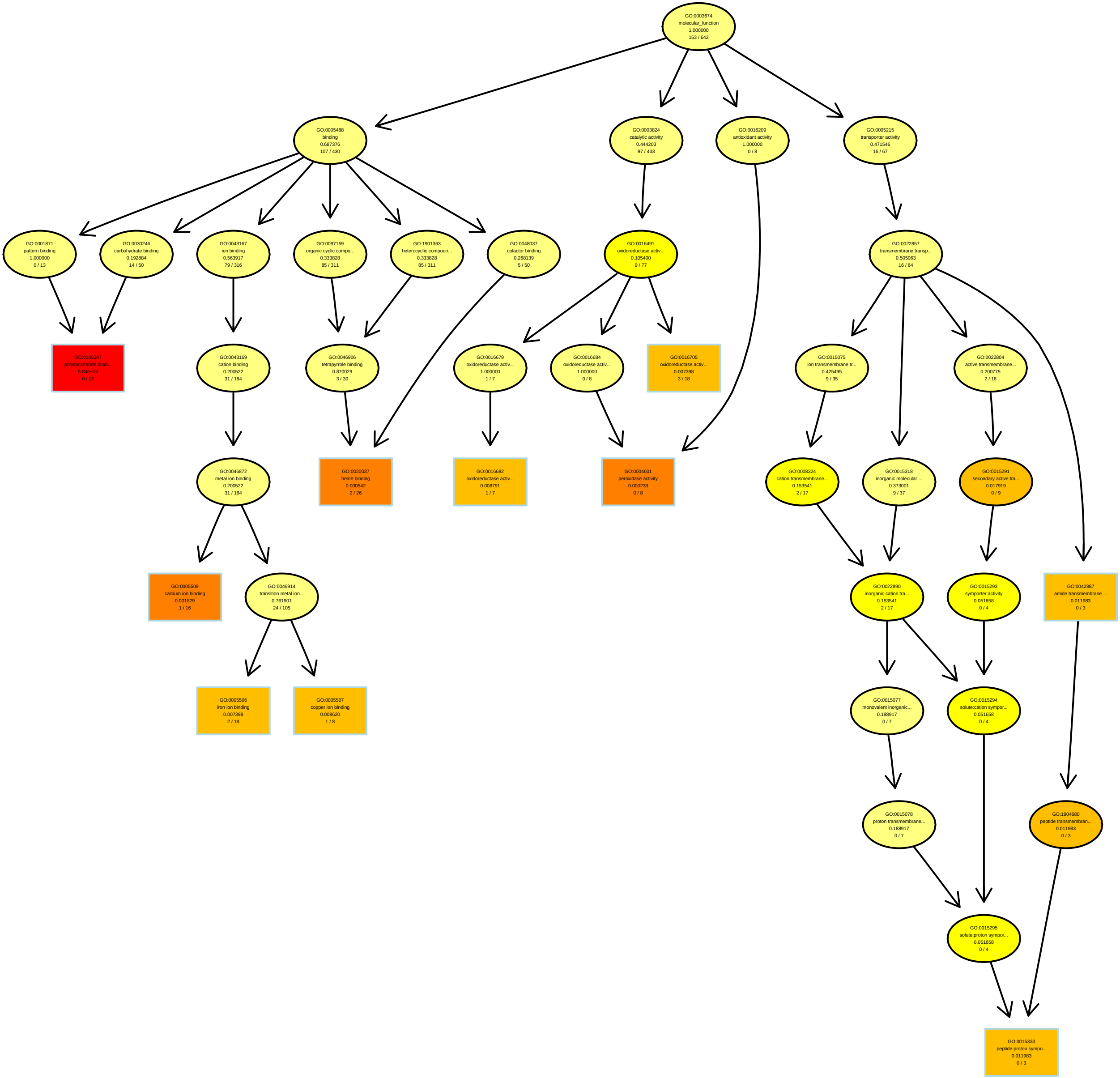

### RT7.58_BP.pdf

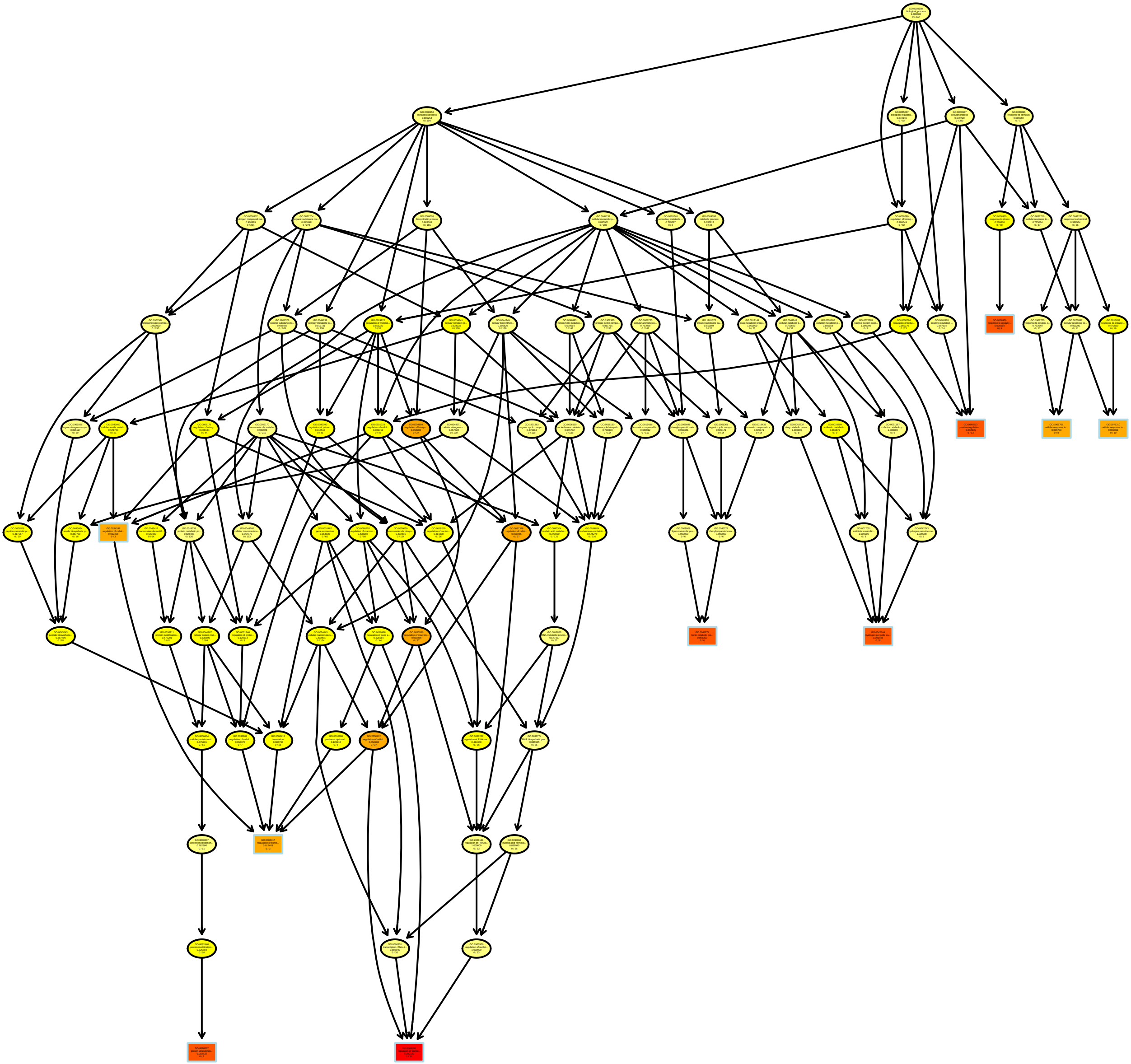

### RT7.71_MF.pdf

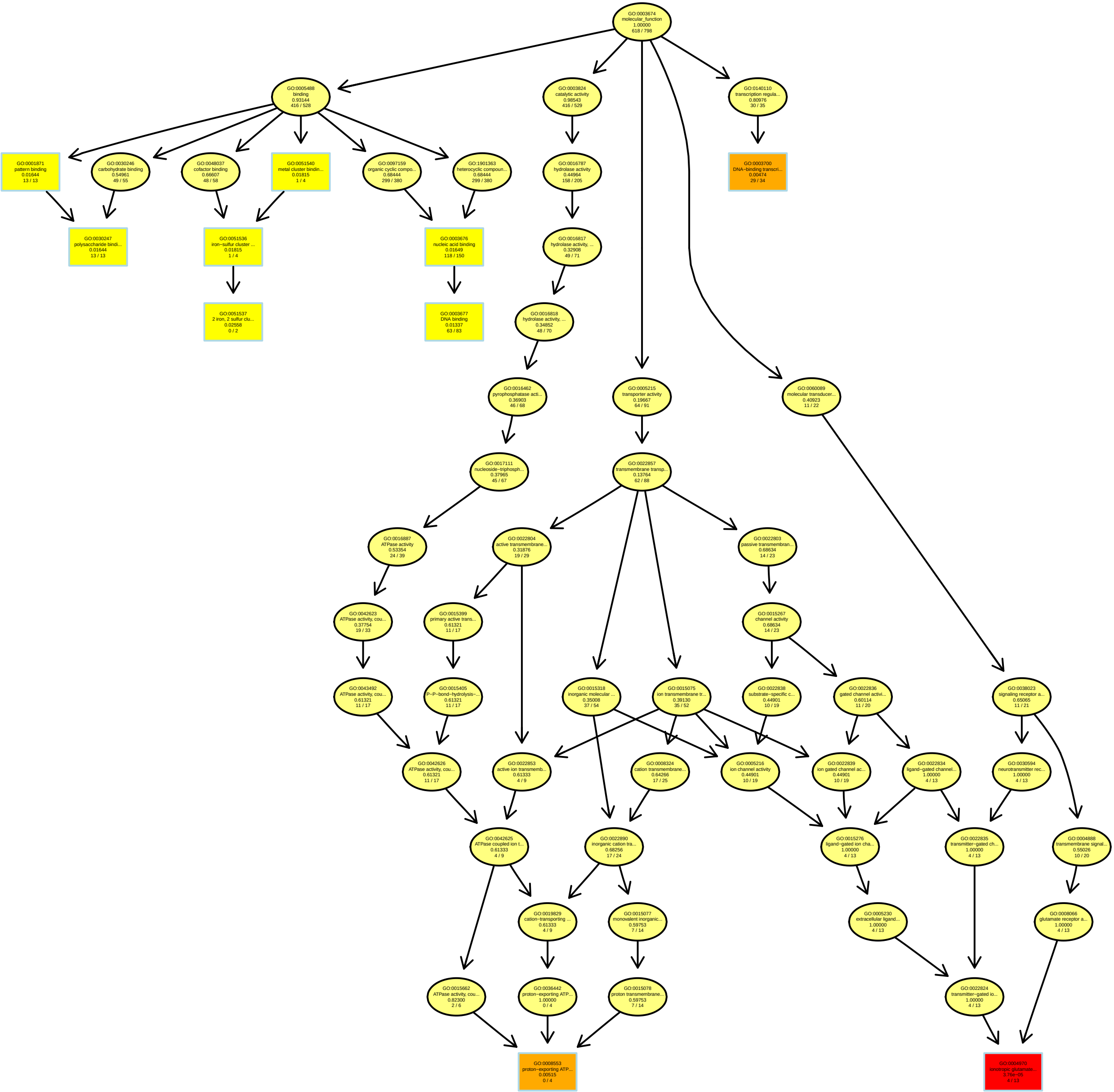

### RT7.83_BP.pdf

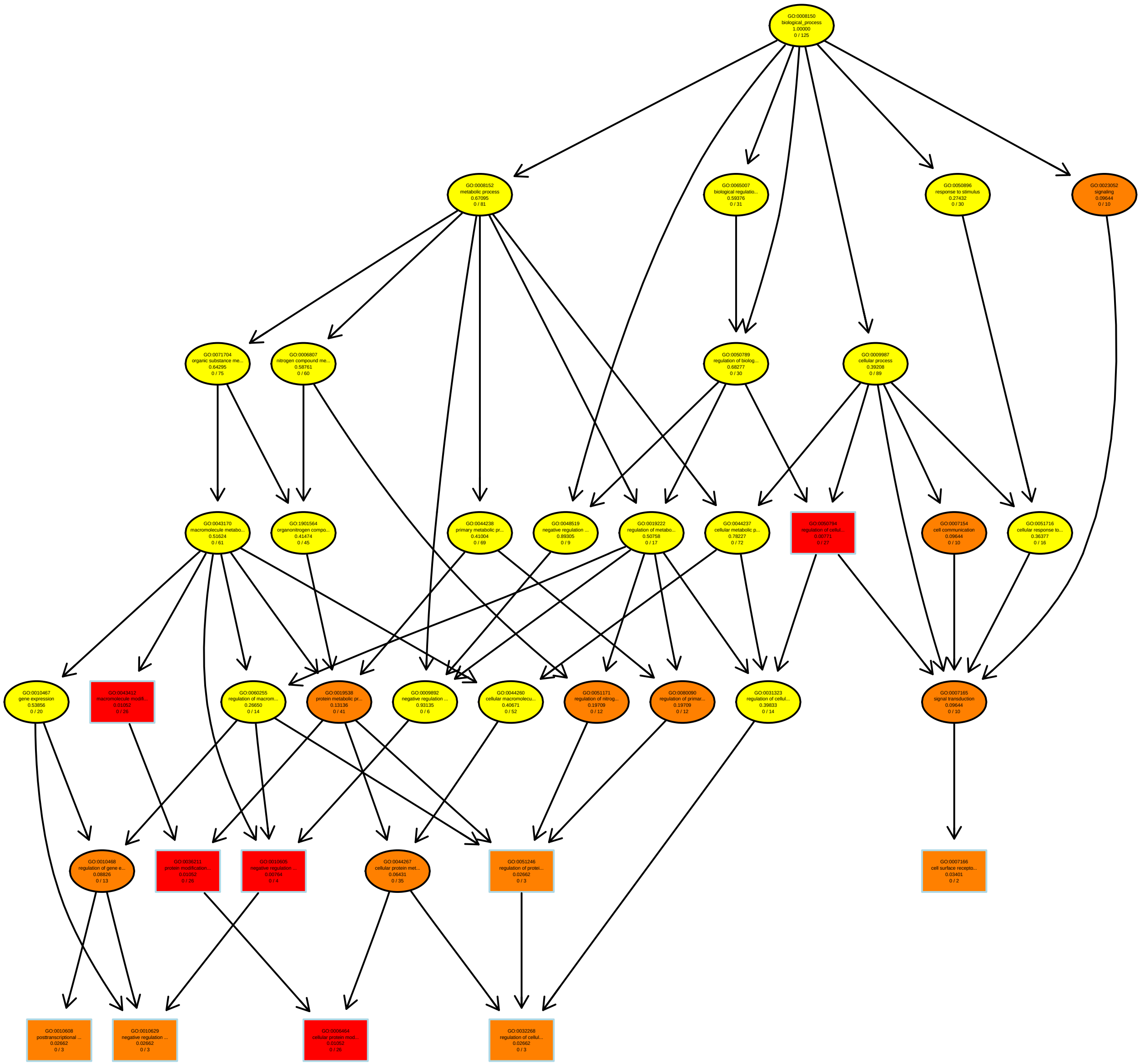

### RT7.93_BP.pdf

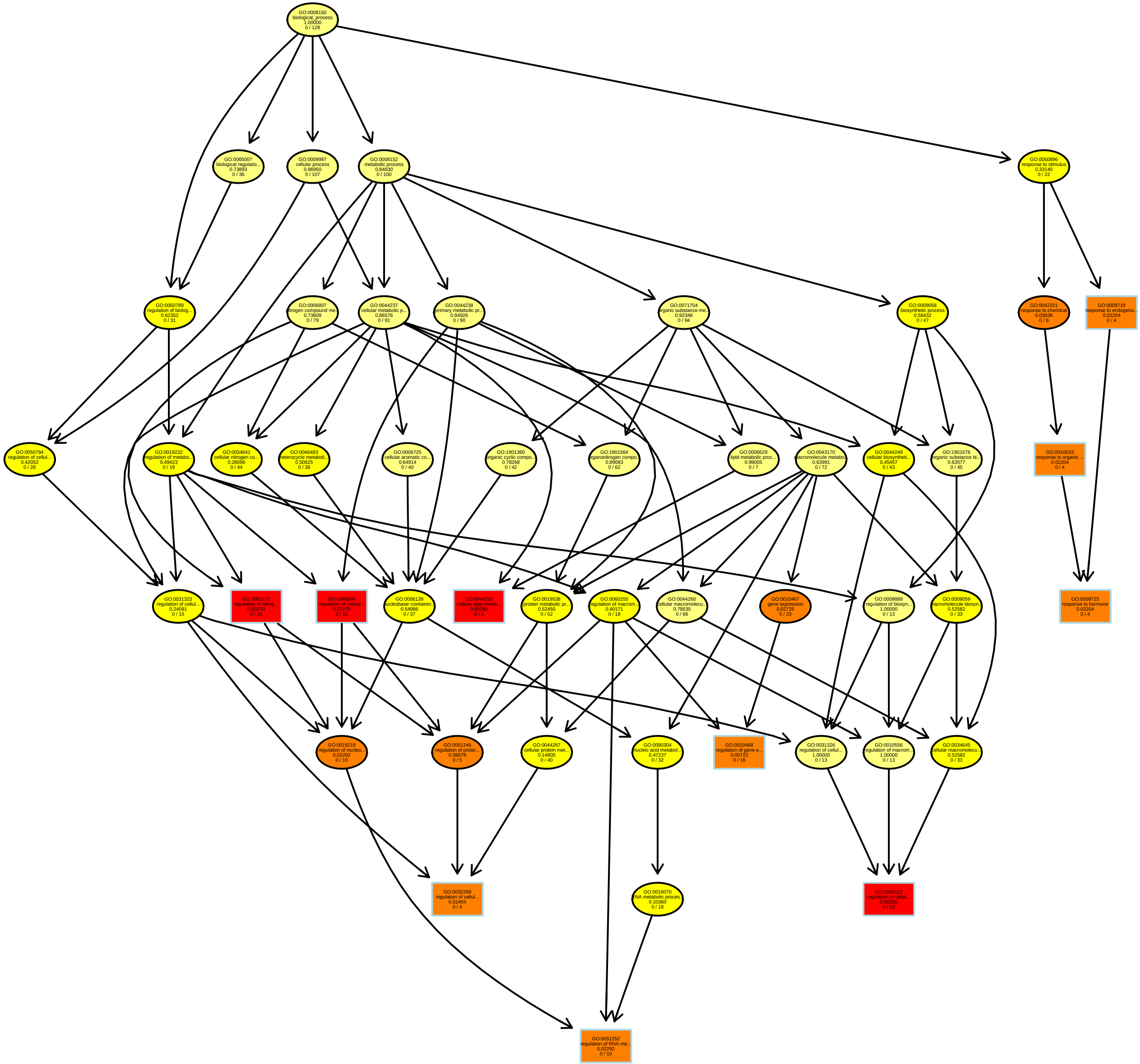

### RT8.3_BP.pdf

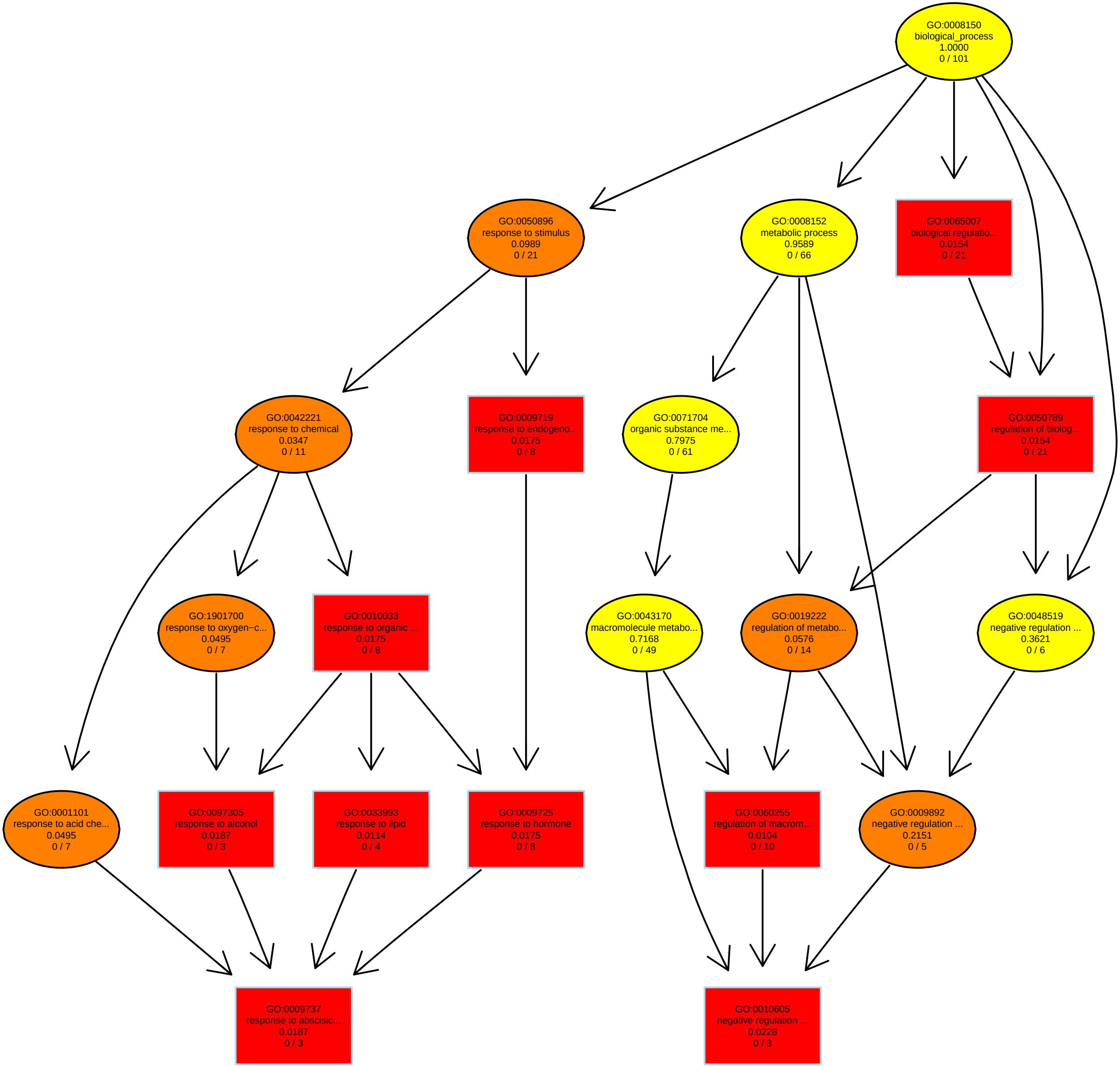

### RT8.21_MF.pdf

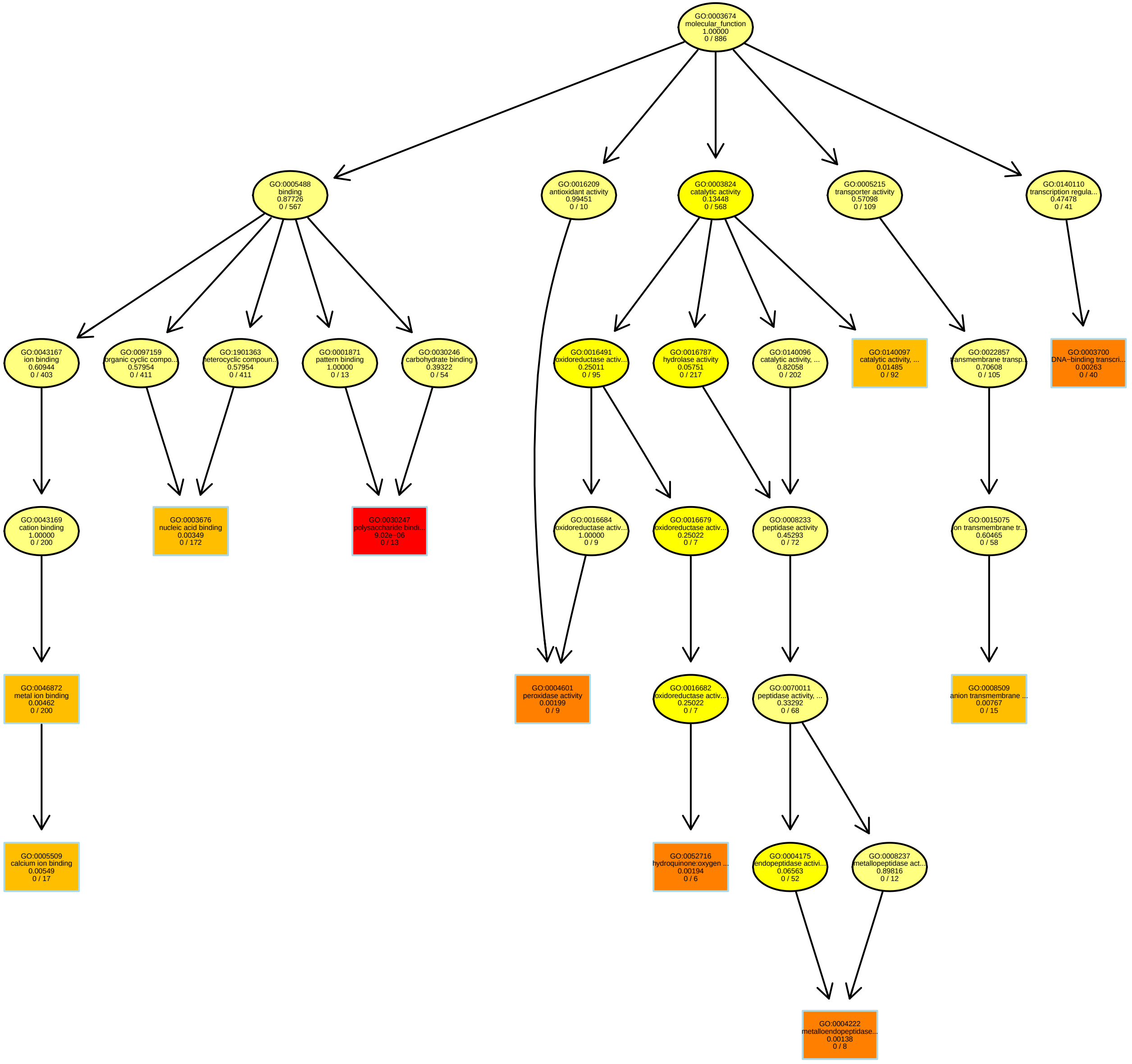

### RT8.38_BP.pdf

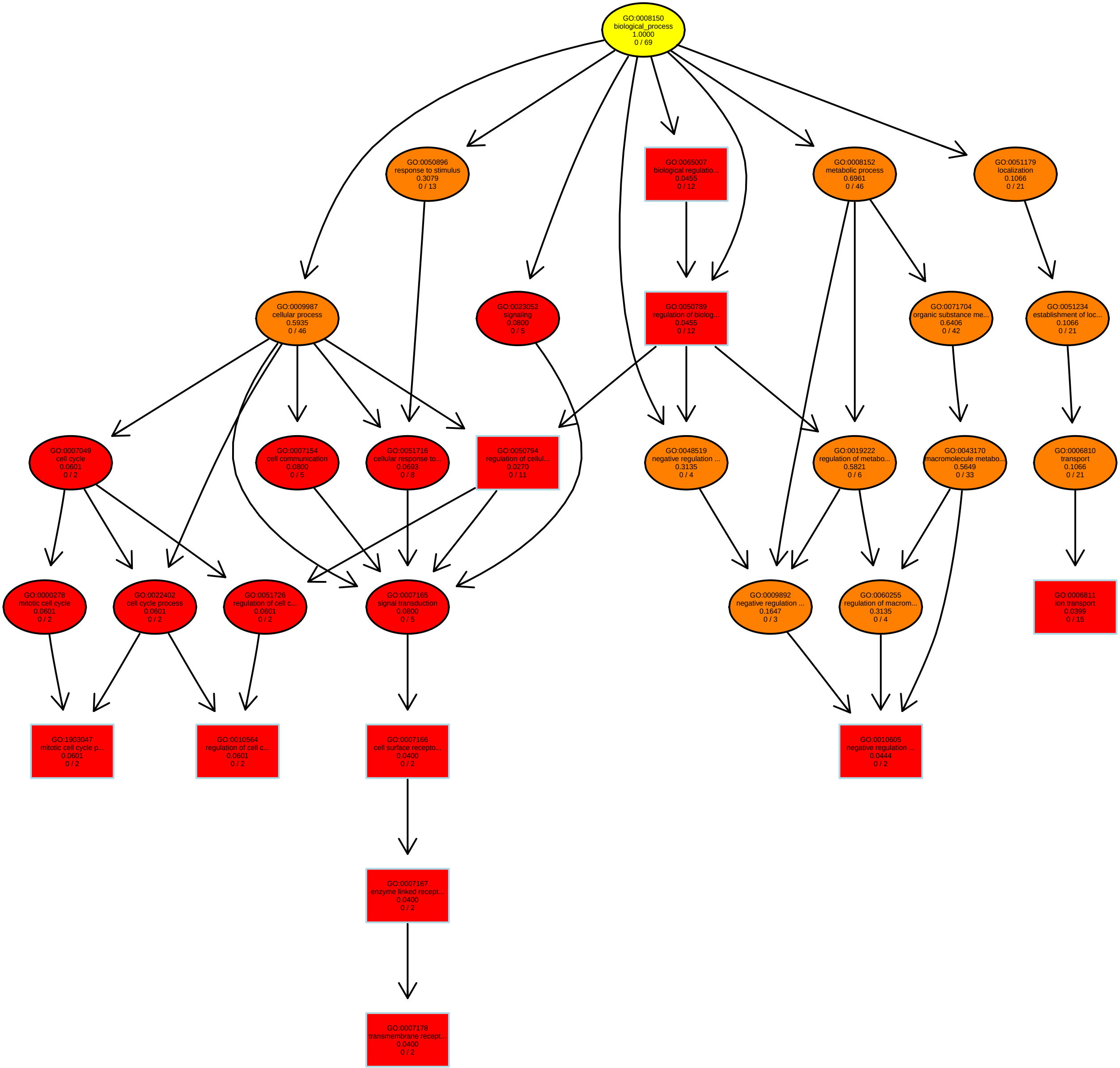

### RT8.51_BP.pdf

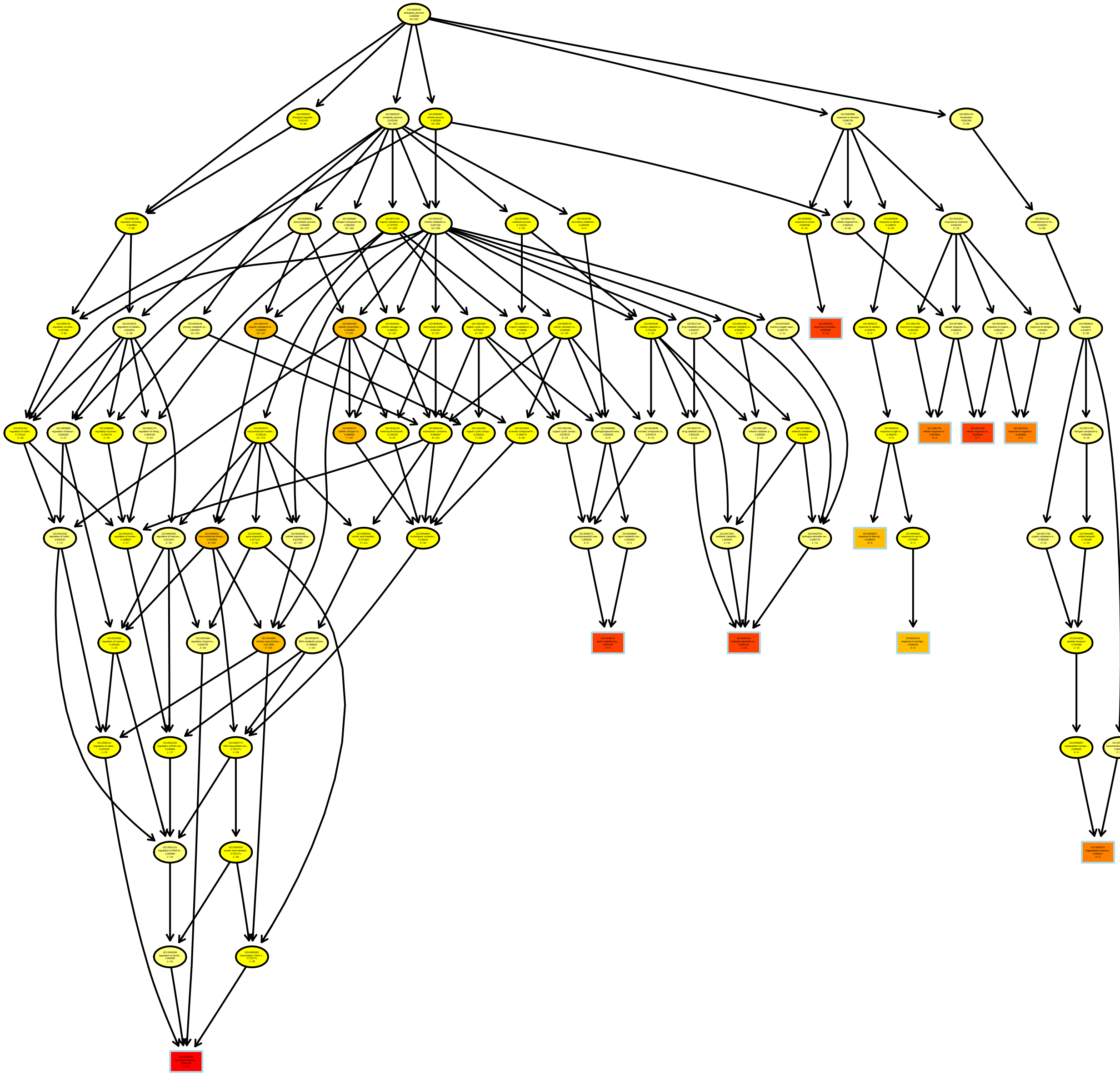

### RT8.89_MF.pdf

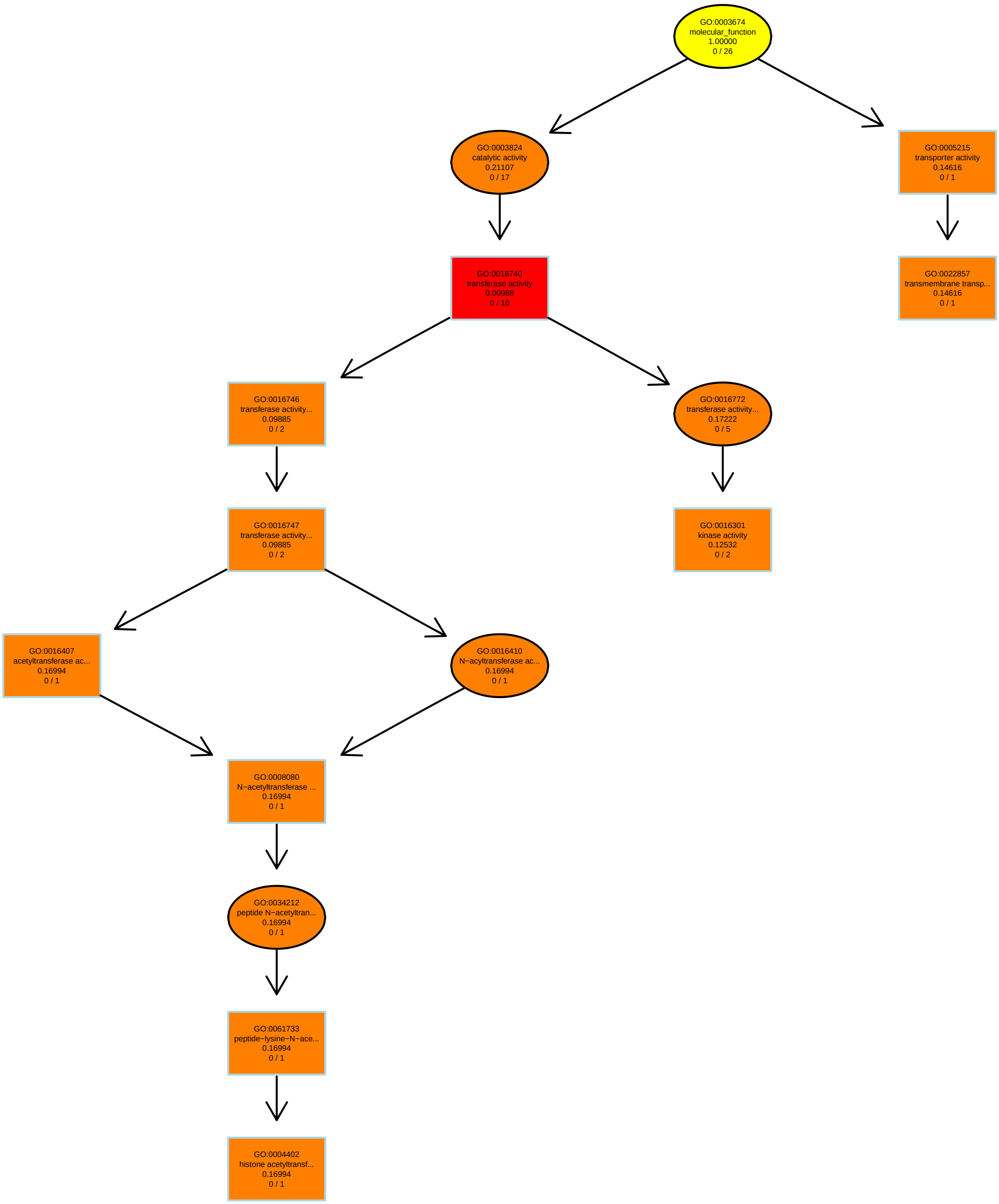

### RT11.78_MF.pdf

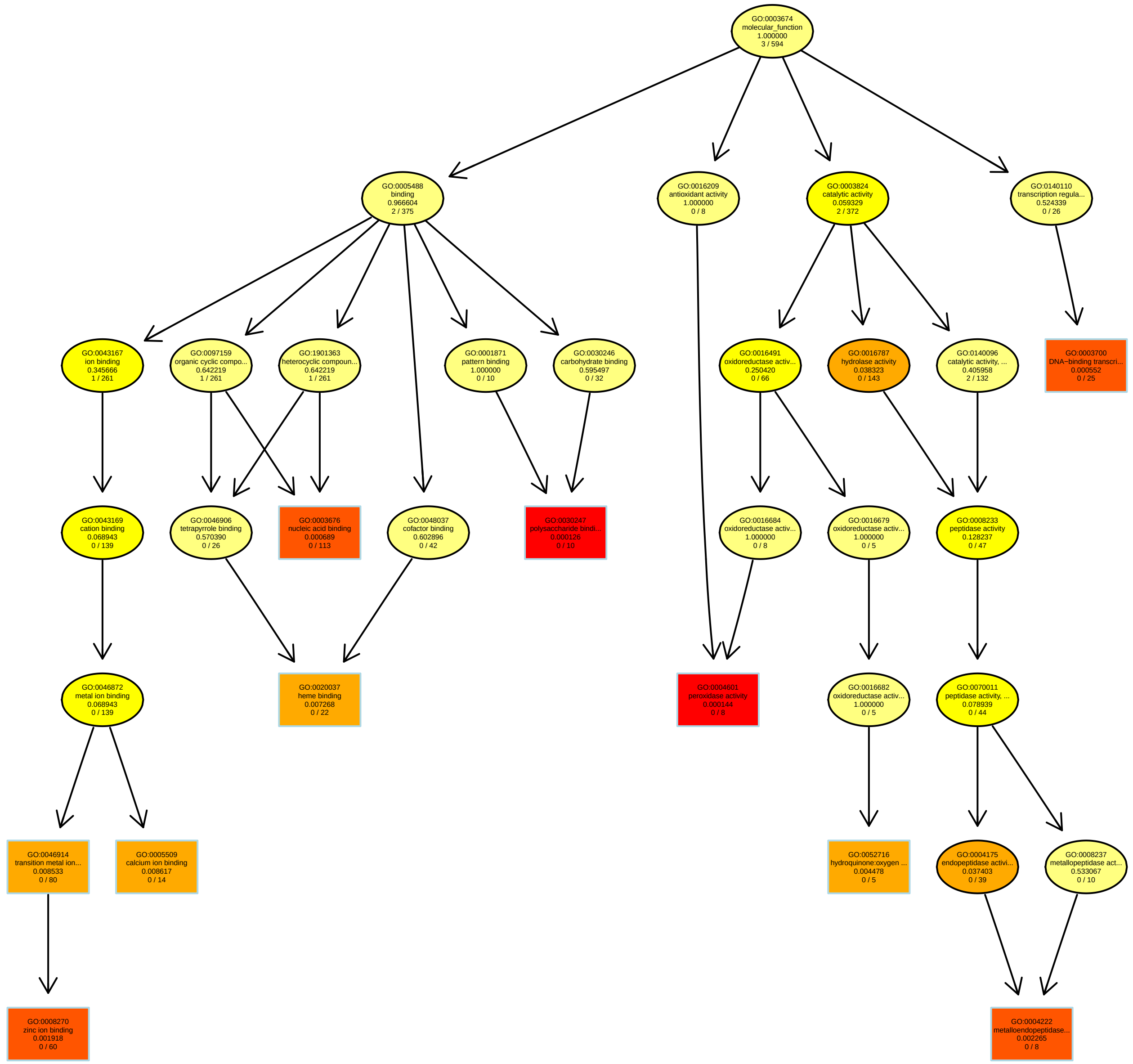

### Solidity_MF.pdf

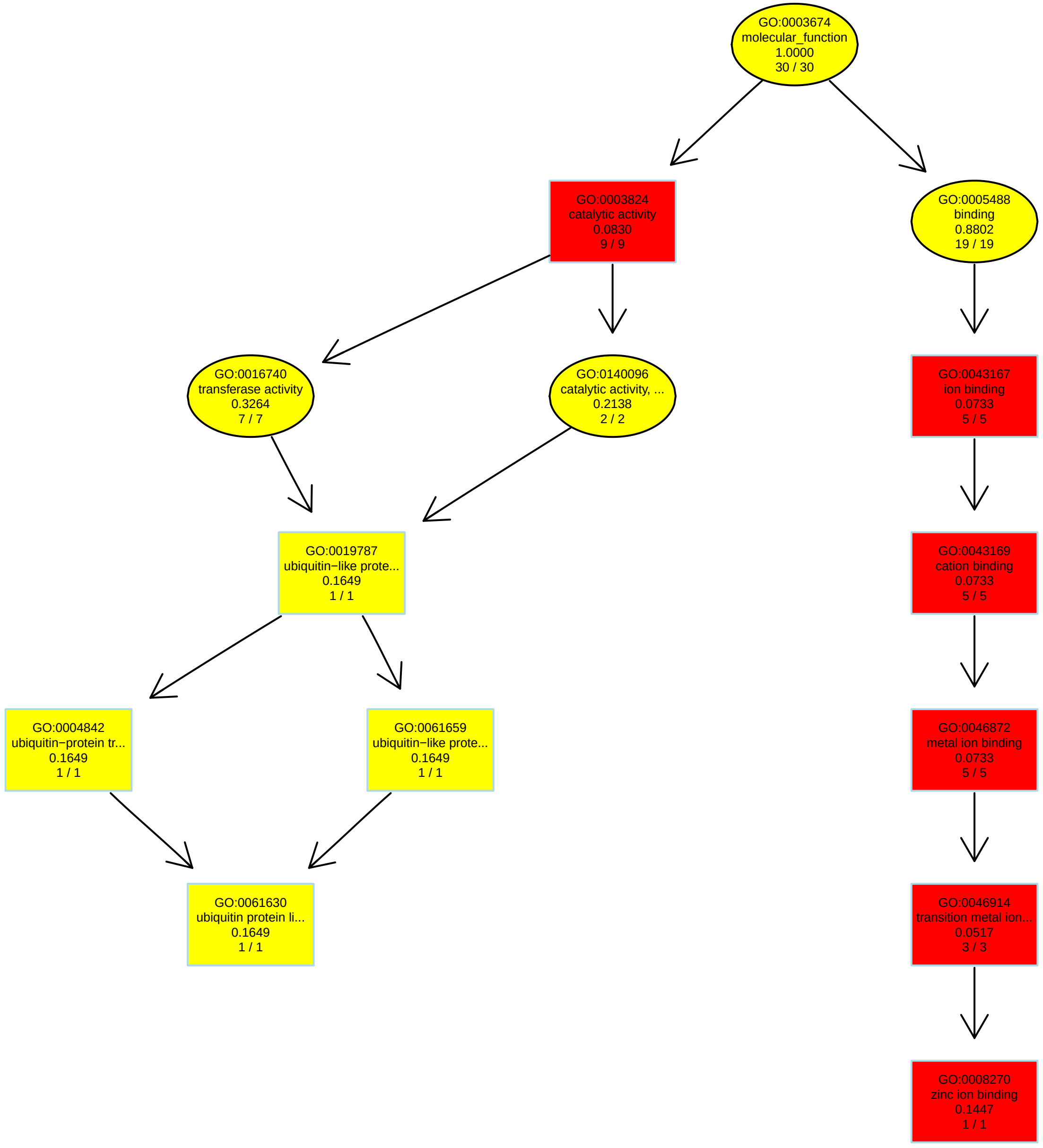

### TotalPhenolics_BP.pdf

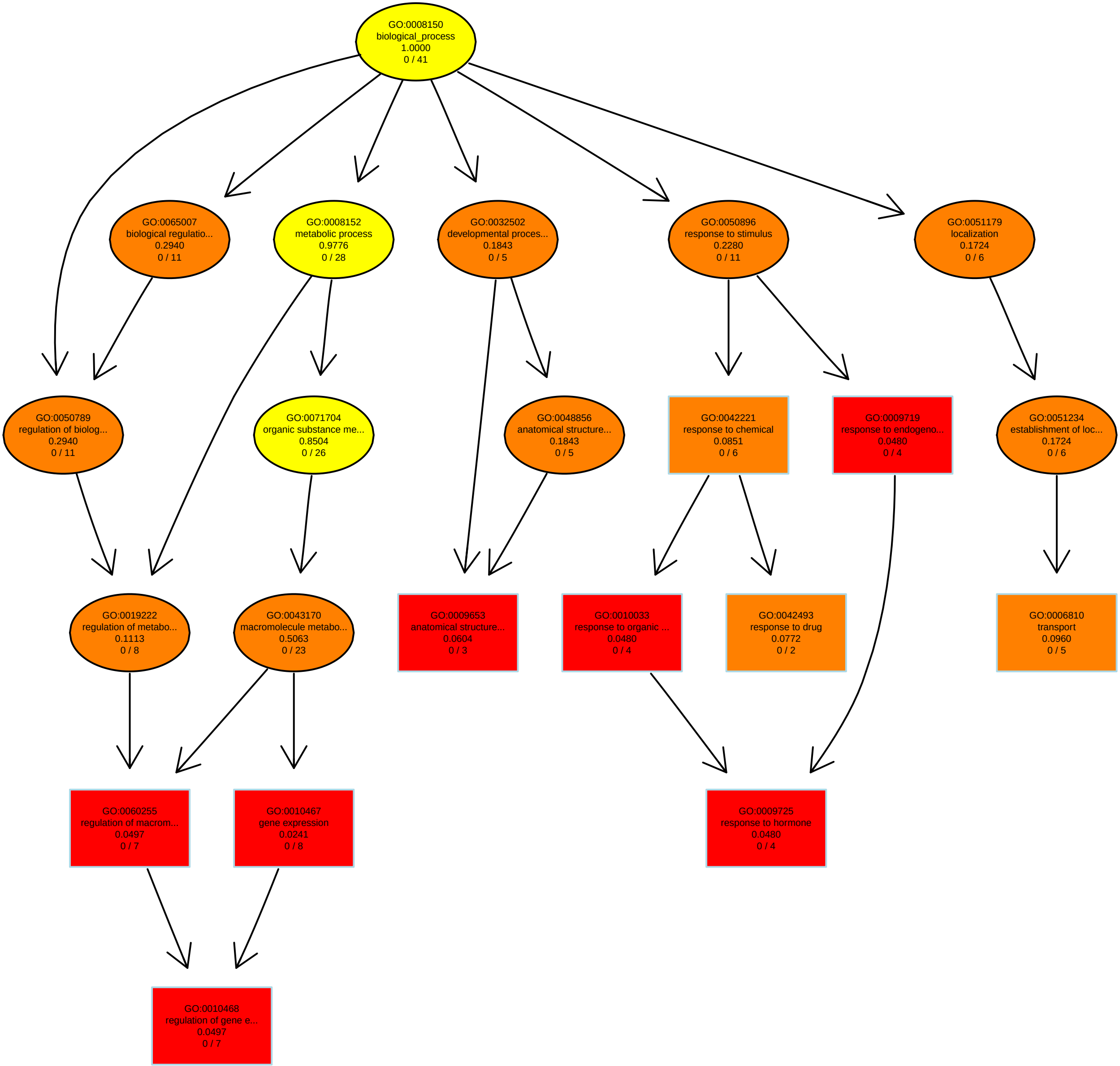

### WaterContent_BP.pdf

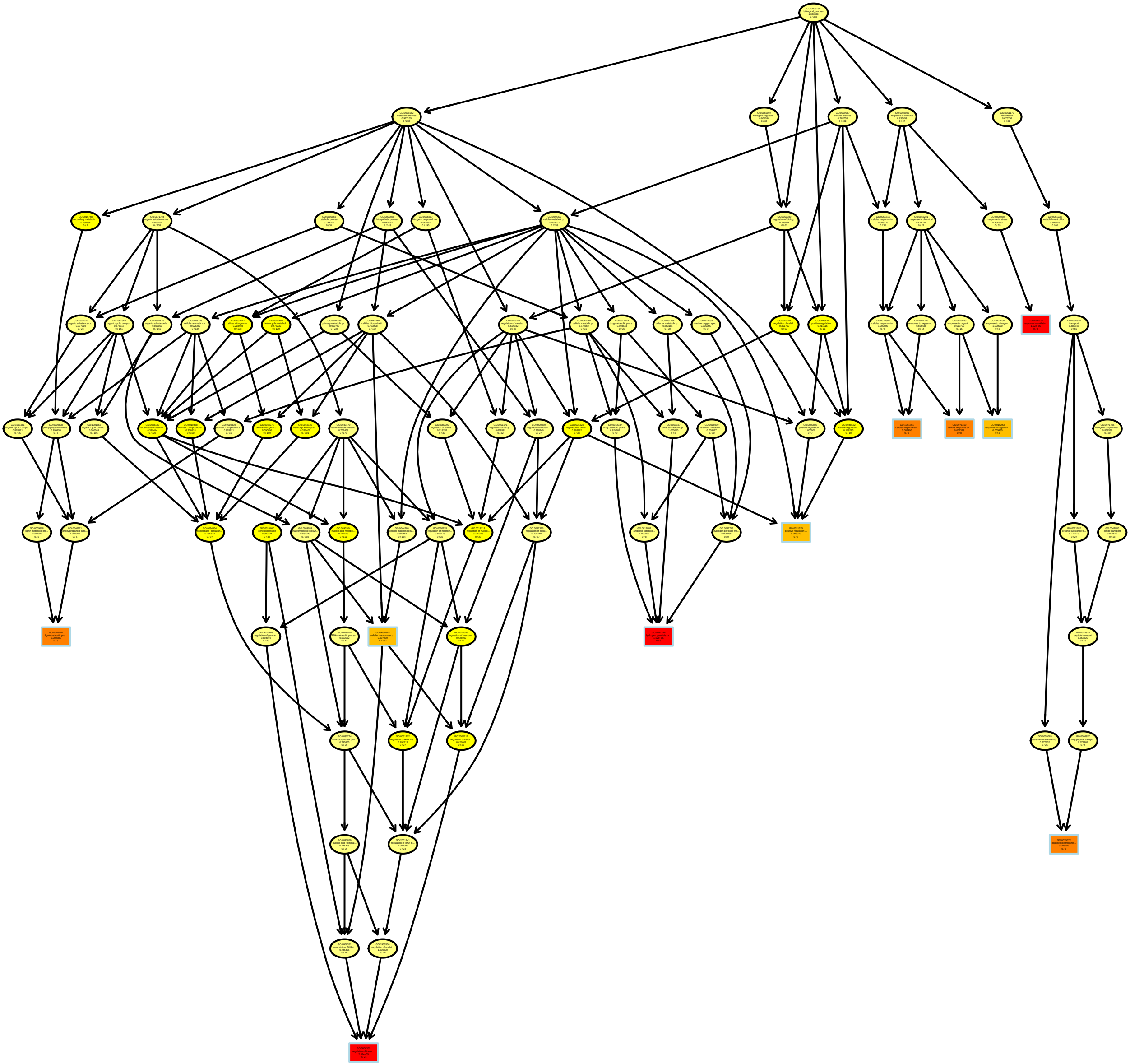
